# Landscape profiling of PET depolymerases using a natural sequence cluster framework

**DOI:** 10.1101/2024.04.01.587509

**Authors:** Hogyun Seo, Hwaseok Hong, Jiyoung Park, Seul Hoo Lee, Dongwoo Ki, Aejin Ryu, Hye-Young Sagong, Kyung-Jin Kim

**Affiliations:** KNU Institute for Microorganisms, Kyungpook National University, Daegu 41566, Republic of Korea; School of Life Sciences and Biotechnology, BK21 FOUR KNU Creative BioResearch Group, Kyungpook National University, Daegu 41566, Republic of Korea; Institute of Biotechnology, CJ CheilJedang Co., Suwon-si, Gyeonggi-do, 16495, Republic of Korea; Zyen Co., Daegu, 41566, Republic of Korea

## Abstract

Since the demonstration that rapid polyethylene terephthalate (PET) decomposition using enzymes is feasible, a number of efficient depolymerases have been reported with the aim of resolving the plastic pollution issues. However, sporadic studies on enzymes with PET hydrolysis activity hinder the understanding of the distribution of potential PETases hidden in nature’s repertoire, and subsequently, the identification of potent enzymes. Here, we present the clustering of 1,894 PETase candidates, which include the majority of known PETases, and describe their profiling. An archipelago landscape of 170 lineages shows distribution of 289 representative sequences with features associated with PET-degrading capabilities. A bird’s-eye view of the landscape identifies three highly promising yet unexplored PETase lineages and two potent PETases, Mipa-P and Kubu-P. The engineered Mipa-P^M19^ and Kubu-P^M12^ variants exhibit both high PET hydrolysis activity and thermal stability. In particular, Kubu-P^M12^ outperformed the engineered benchmarks in terms of PET depolymerization in harsh environments, such as with high substrate load and ethylene glycol as the solvent. Our landscape framework and the identified variants assist in the understanding of how biological processes respond to solid-state and non-natural PET plastics.

## Introduction

Microbial decomposers can exhibit the ability to break down newly encountered material through closely functioning genes, with subsequent mutations that improve the process recorded in their genetic information. The isolation of *Ideonella sakaiensis*, which utilizes polyethylene terephthalate (PET) as its sole carbon and energy source (*1*), illustrates this type of evolution at work, which may explain the occurrence and ecological accommodation of natural polymers. However, evolutionary processes progress very slowly and inconsistently, making them difficult to perceive in real time. For this reason, although identifying natural phenomena is critical to biological research, enzyme candidates that have evolved to mediate naturally inefficient reactions, such as the breakdown of plastics, remain hidden in nature’s repertoire.

Biocatalysts have been widely investigated as a means to break down the ester bonds of PET and regenerate its monomers (*2*, *3*). The isolation of microorganisms and extracellular proteins capable of degrading synthetic polyester, such as cutinase from the fungus *Humicola insolens* (HiC) (*4*) and hydrolases and cutinases from *Thermobifida* spp. (*5*, *6*, *7*), has raised the possibility of biological PET degradation using α/β structure hydrolytic enzymes. Other PET hydrolases, such as leaf compost cutinase (LCC) (*2*, *8*) and PETase from *I*. *sakaiensis* (IsPETase) (*1*) have also been identified through homology searches and characterization in a genetic pool of esterolytic sample. In recent years, extensive research has been conducted to increase the PET hydrolytic ability of these enzymes through various methods, including rational engineering (*2*, *9*, *10*, *11*), computational and machine learning-based engineering (*12*,

*13*, *14*), and directed evolution (*15*).

There have been significant efforts to find enzyme templates using sequence information. For example, Joo et al. classified a group of type IIb PET hydrolases with key residual features of IsPETase in a phylogenetic tree of 69 sequences (*16*) that included mesophilic PET hydrolases from *Rhizobacter gummiphilus* (RgPETase) and the *Burkholderiales* bacterium RIFCSPLOWO2_02_FULL_57_36. Similarly, Danso et al. manually selected 13 sequences for biochemical analysis among 504 PET hydrolase candidates retrieved from BLAST search using the seed sequences prepared using HMMER and previously identified PET hydrolases such as PET2, PET5, PET6, and PET12 (*17*). In a search for thermotolerant enzymes, Erickson et al. selected 74 sequences that contained 37 sequences with PET hydrolysis activity using various tools such as a hidden Markov model (HMM), optimal growth temperature data, and a machine learning model ThermoProt, ultimately finding 23 novel PET hydrolases (*18*). Recently, Hong et al. reported the discovery of a PETase from *Cryptosporangium aurantiacum* (CaPETase) with both high thermostability and activity at ambient temperatures from 10 unknown sequences (*19*).

Nevertheless, these one-by-one and/or HMM-based mining approaches are limited in that they cannot see the entire fitness landscape, which provides a comparative perspective on the evolutionary success of naturally occurring PETases. Thus, a more comprehensive framework based on sequence mapping on the landscape is necessary to fully understand groups of genes and discover superior PET depolymerases. In the present study, we report the generation of a visual map of protein homolog lineages using public sequence databases to utilize the natural diversity of the evolutionary success in PETase development. We extracted 2,064 non-redundant protein sequences grouped as polyesterase-lipase-cutinase in the ESTHER classification (*20*), and generated 1,894 unique categories with 170 clusters. Primary and secondary sampling on the map revealed a total of 107 potential PETases and identified sequence lineages with highly promising proteins through a bird’s-eye view. We then engineered these proteins using sequence information from the PETases discovered in this study and successfully developed a variant that can be utilized under harsh industrial depolymerization conditions. This analysis of global diversity and the engineering of multiple templates represents a groundbreaking method for identifying potential enzymes with specific functions and engineering their sequences.

## Results

### Clustering of PETase candidates

To organize the view of fitness landscape of PETase candidates, we developed a protein sequence clustering and indexing method named PHILOSOPI (<u>P</u>rotein <u>H</u>omolog Islands <u>L</u>andscape <u>O</u>rganization for <u>S</u>equence <u>O</u>rigin and <u>P</u>rofile Indexing) (Fig. 1). For the library construction, we first searched for a group of enzymes homologous to the 26 reported sequences with PETase activity and constructed an initial set of 25,418 non-redundant sequences (table S1). This set was distributed across all blocks of the ESTHER database and contained various families of α/β hydrolases, including polyesterase, lipase, tannase, fungal cutinase, type-B carboxylesterase, and fold-3 domain-containing esterase. Because 19 reported PET hydrolases with relatively high activity were clustered in the polyesterase-lipase-cutinase family, we reduced the initial set to a subset library with 2,064 sequences (Fig. 1). The library mostly contained sequences derived from microorganisms in the phyla *Actinomycetota* and *Proteobacteria*, which accounted for approximately 83.5% and 12.5% of the sequences, respectively (fig. S1). The lowest pairwise sequence identity in the library was 14.18% (fig. S2). Because the library was expected to cover the major potential sequences with a solid polyester depolymerization function, we referred to it as the ester-based plastic hydrolase library.

**Figure 1.**
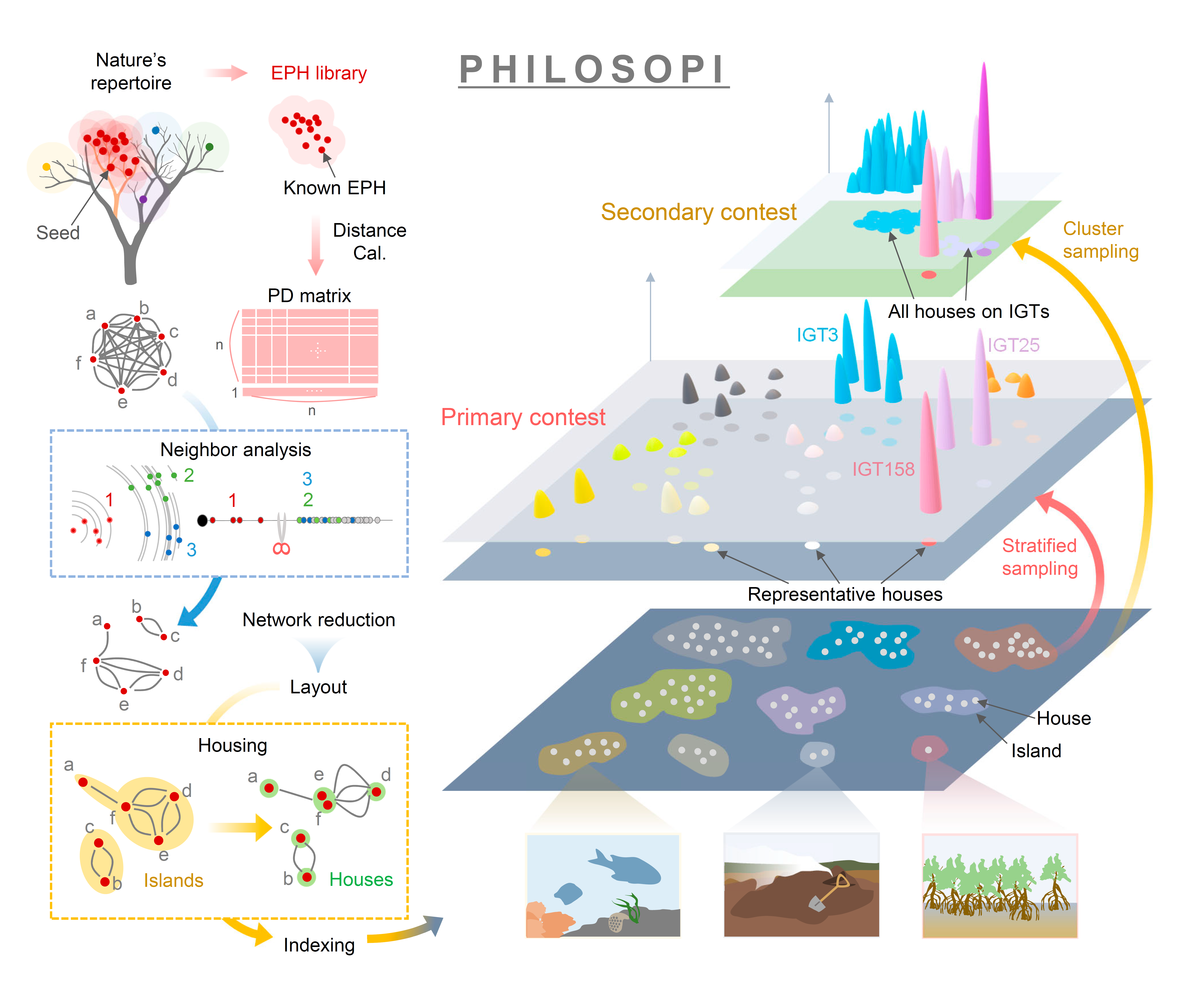
Construction of ester-based plastic hydrolase (EPH) library and PHILOSOPI procedures. PHILOSOPI analysis proceeds as follows: library construction, networking, and landscaping. The EPH library is prepared using the seed sequences with search ranges individually applied to PSI-BLAST cutoff. For networking, pairwise distance calculation generates a distance matrix, and neighbor analysis (materials and methods) in PHILOSOPI reduces the network edges for clustering and visualization. The branches of protein lineages in the variation (evolutionary) tree are presented by landscaping. In the separate islands, housing the ‘sequence’ nodes into new ‘house’ nodes provides indexes of island and house, and makes the subsequent feature profiling easier. The primary and secondary contests were held by stratified and cluster samplings, respectively, on the map.

Because the number of sequences of the library was still too large to explore experimentally, the network analysis using PHILOSOPI classified the sequences of the library into 170 clusters using Wilbur and Lipman distances (*21*) and a heuristic algorithm of nearest neighbor methods (Fig. 1; figs. S3 and S4). The clustering reduced the number of network edges from 2,064^2^ to 14,889 and the mean and standard deviation of the distance from 0.556 and 0.162 to 0.081 and 0.074, respectively (fig. S5). The 2,064 sequences in the library were also merged into 1,894 nodes of sequence origins in the PHILOSOPI network by considering differences in an extra N-terminal region including signal sequences and cross-referencing with both nucleotide sequencing data and the UniProtKB database. To help imagine the archipelago landscape of the nodes’ origination, we refer to each of the clusters and nodes as an island (I_1_-I_170_) and a house (H1-H1894), respectively. The number of houses on each island was varied widely; for example, the largest island (I_1_) contained 689 houses, while 23 islands had only one house (Fig. 2; fig. S5). It is worth noting that three well-studied PET hydrolase groups were clearly represented in a specific island archipelago: I_20_ for *Thermobifida* enzymes (assigned to Group 7 by Erickson et al.) containing TfCut2 (*22*) and *T. cellulosilytica* cutinase (Thc_Cut2) (*23*); I_33_ for type IIb enzymes containing IsPETase (*1*) and RgPETase (*24*); and I_120_ for extreme thermophilic enzymes containing LCC (*8*) and bacterium HR29 PETase (BhrPETase) (*25*) (Fig. 2). Other reported enzymes were located on 22 islands (Fig. 2), resulting in 145 of 170 islands remaining *terra incognita*.

**Figure 2.**
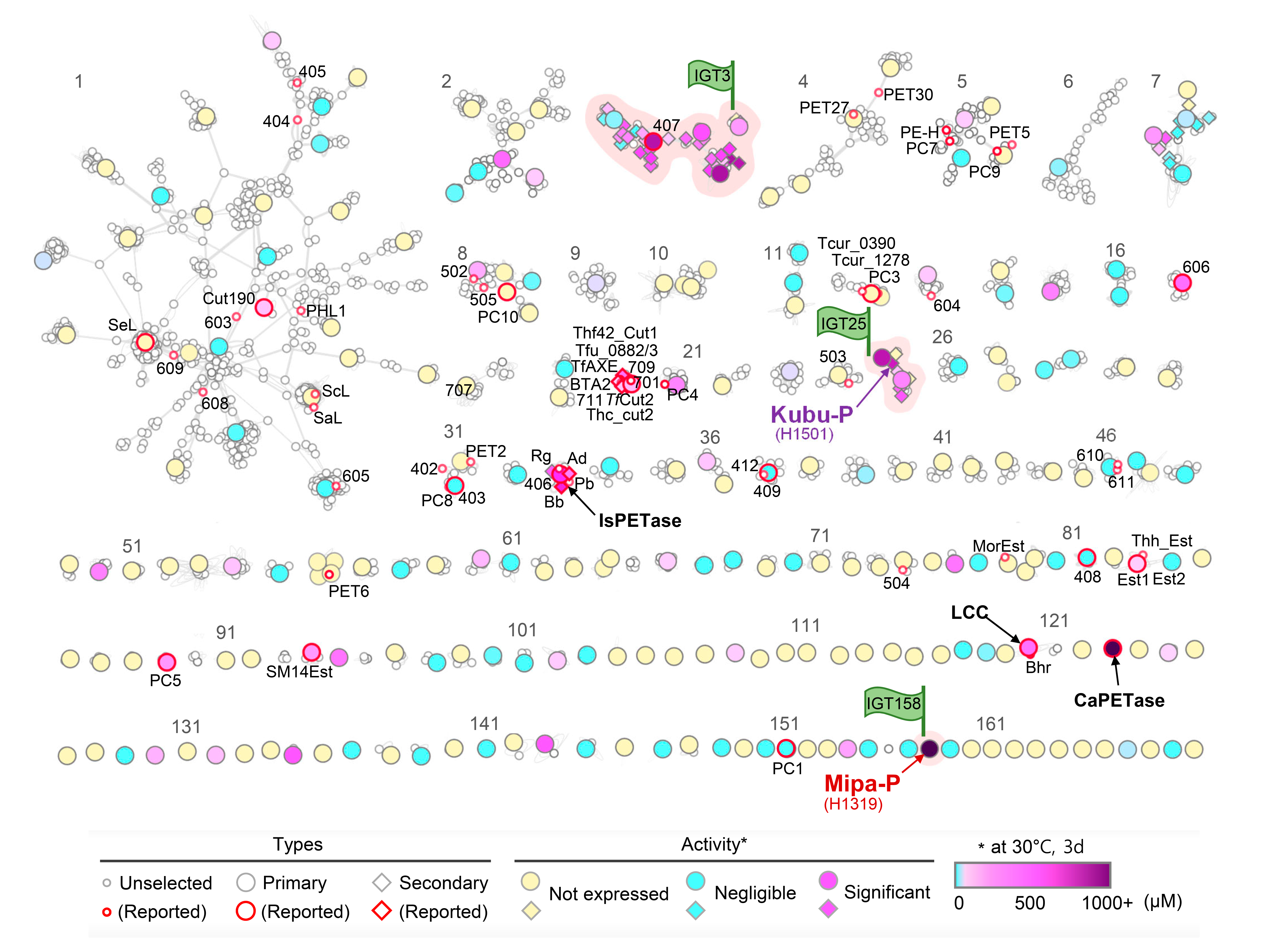
Landscape profiling of PETase candidates. Archipelago map of 170 islands is presented. The 170 islands are labeled with their numbers. Each node represents a house. Large circles and diamonds indicate houses in the primary and secondary contest, respectively. The tested houses are colored by mean PET-degrading activities at 30 °C, and edges were shown as light-green color. The three IGTs are highlighted with pink boxes and purple flags and the two potent PETases Mipa-P and Kubu-P are labeled with their house numbers. Houses of benchmarks are indicated by arrows and labeled accordingly.

### Primary contest for PETases using stratified sampling

A primary contest for PETases was held using representative houses from the 164 islands. Because islands with a larger number of houses tend to have a higher mean distance (Fig. 2 and fig. S5), stratified sampling was conducted proportional to the network density on each island, leading to a total of 232 houses from 164 islands being selected for the primary contest. When the representative houses were expressed in an *E. coli* strain, many houses were not successfully produced at a level suitable for experimentation, thus only 105 houses were retained for participation in the primary contest. The 105 houses contained 13 benchmarks, including the highlighted PET hydrolases such as IsPETase (I_33_-H1262), LCC (I_120_-H971), Thc_Cut2 (I_20_-H6), and CaPETase (I_123_-H1110). The contest was conducted based on evaluation criteria for three key factors: monomer product release at 30 °C for PET hydrolysis activity, the melting temperature (Tm) for thermal stability, and enzyme production titer (Fig. 3A; fig. S6 and S7). Surprisingly, 58 houses (55.2%) exhibited detectable amounts of released monomer, of which 34 (32%) had activity levels higher than Est2 (*26*), the PET hydrolase with the lowest activity among the active benchmarks. More surprisingly, of the houses exhibiting PET hydrolysis activity, five houses exhibited higher activity than IsPETase (Fig. 3A). The highest activity was found in H1319 from I_158_, which showed even higher activity than CaPETase (Fig. 3A), the PETase with the highest activity at 30 °C reported to date (*19*). The thermal stability of the tested houses was then assessed by measuring the differential scanning fluorimetry Tm values of the proteins, and the Tm values of 94 houses were successfully determined (Fig. 3A; fig. S7). There were 15 thermophilic houses with Tm values above 60 °C, and the top six houses included four benchmarks, such as LCC (86.2 °C), Thc_Cut2 (69.5 °C), 407 (67.7 °C) (*18*), and CaPETase (64.9 °C), which indicates that the previous studies for PETase have been directed towards thermophilic enzymes. The protein production titer values exhibited a large variation; SM14est (*27*), CaPETase, Thc_Cut2, and Est2 were in the top quartile, while PC8 (*19*), 408 (*18*), LCC, 606 (*18*), and IsPETase were in the bottom quartile (fig. S6).

**Figure 3.**
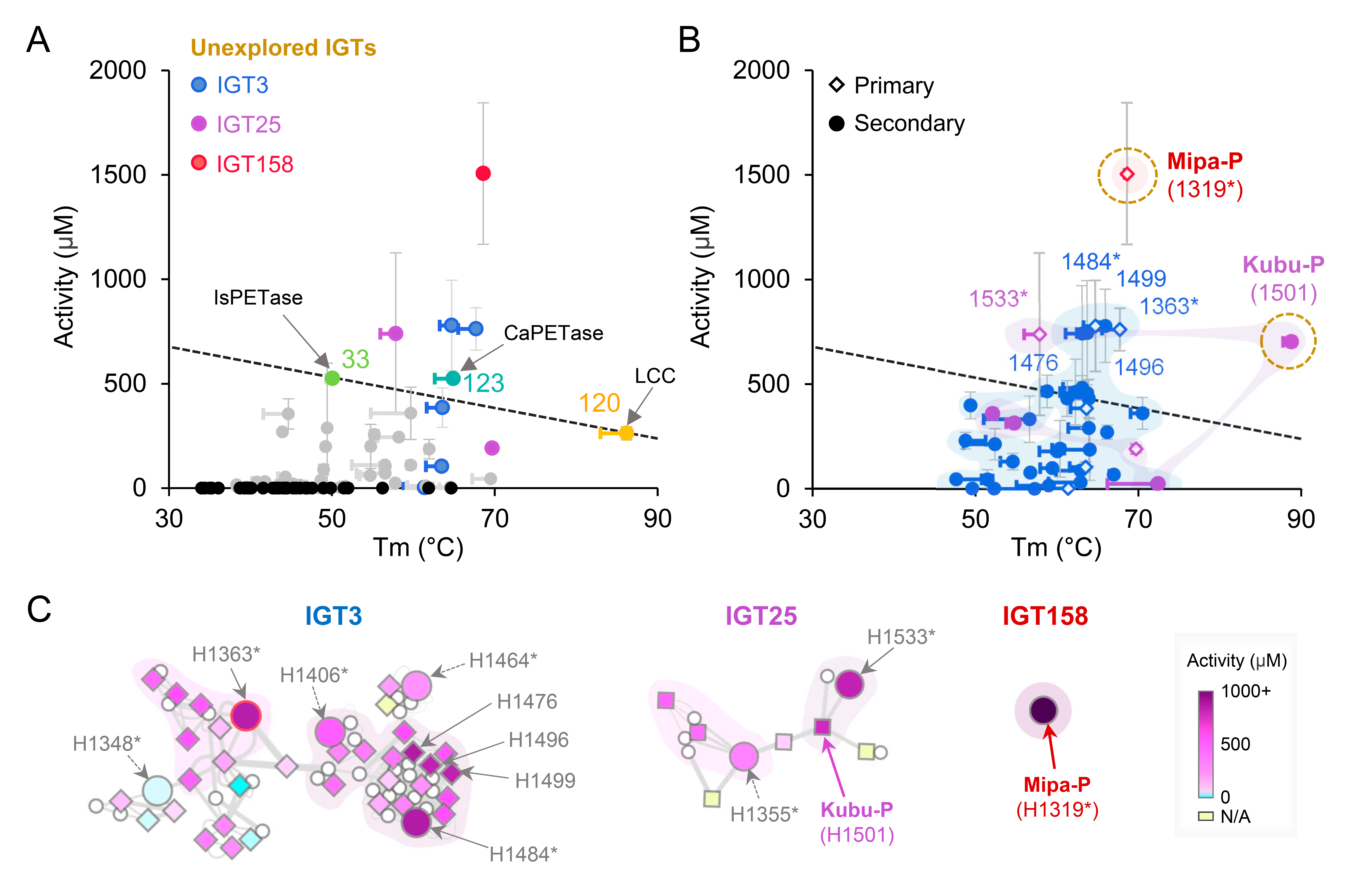
PETase contest. (**A**) The primary contest using stratified sampling. Houses releasing detectable monomers from PET and those with ignorable activity are shown as gray and black circles, respectively. The SD values of the activity data are presented as bars□(n = 3). The Tm values are shown as a line between Tm_D_ (car) and Tm_B_ (circle). IsPETase, LCC, and the houses on the three IGTs, are colored by each color scheme. (**B**) The secondary contest using cluster sampling. Houses on IGT3, IGT25, and IGT158 are shown in activity-Tm plot. The eight qualified houses are labeled, and the representative houses in the primary contest are distinguished by the marker shape and adding an asterisk in label. The shadows represent edge connections between the houses. (**C**) Network of the three unexplored IGTs. The figure is drawn as in Fig. 1B. The qualified houses are indicated by solid-line arrows and the houses in the primary contest but not selected as qualified are by dashed-line arrows.

The primary contest for PETases suggests that there are more enzymes acting on PET in natural sequences than originally suspected, which allowed islands containing houses satisfying both high PET hydrolysis activity and thermal stability to be identified. Based on a connection line between IsPETase and LCC in an activity-Tm plot, five houses, including H1484 and H1363 from I_3_, H1533 from I_25_, H1110 from I_123_, and H1319 from I_158_, were selected as qualified PETase candidates (Fig. 3A). As a result, the four islands containing the qualified representative houses (I_3_, I_25_, I_123_, and I_158_) were identified as distinguished lineages of sequence evolution, and we flagged these islands as “Island’s Got Talent” (IGT). Of the four IGTs, IGT123, which contains one house (H1110; CaPETase), was the only island that has previously reported, and the remaining three IGTs (I_3_, I_25_, and I_158_) have been unexplored to date. H1319 from IGT158 exhibited the highest activity at 30 °C among the tested houses and also had a high thermal stability with a Tm_B_ of 68.6 °C (Fig. 3A and fig. S7).

### Secondary contest for PETases using cluster sampling

The three unexplored IGTs (IGT3, IGT25, and IGT158) contained varying numbers of houses on each island. In particular, IGT3 and IGT25 contained a total of 65 and 13 houses, respectively, while IGT158 had only one house (H1319). In order to identify and search the most talented house on IGT3 and IGT25 and assess the representativeness of the house selected in the primary contest on each island, we conducted a secondary contest for PETases using cluster sampling and selected 33 and 6 additional houses from IGT3 and IGT25, respectively (Fig. 3B), which covered more than half of the untested houses of those islands. To further assess the representativeness of the houses on each island, we also included 5 and 3 houses from I_20_ and I_33_, which are well-known islands containing TfCut2 and IsPETase, respectively, and 10 houses from I_7_, an island with low activity and a medium number of houses, in the secondary contest. Overall, a total of 57 houses from 5 islands participated in the secondary contest. The success rate of protein production increased dramatically compared to the primary contest, with 53 houses (92.9%) producing at a level suitable for the experiment (fig. S6), indicating that the protein expression patterns of the houses were quite similar within an island. The PET hydrolysis activity and Tm values were also significantly higher, with 49 houses (92.4%) exhibiting detectable amounts of released monomer, and 30 houses (56.6%) having Tm values above 60°C (fig. S7). Interestingly, houses from IGT3 and IGT25 had a sample mean for PET hydrolysis activity or Tm values that were significantly higher than those from I_20_ and I_7_, or I_33_, respectively (fig. S8). These results suggest that the representative houses selected for the primary contest were reasonable and that PET degradation talent is connected within each island. However, we also observed disparities in regional characteristics within individual islands. In IGT3, local regions with H1484 or H1499 had high activity, while those with H1348 tended to be relatively weak (Fig. 3C). Similarly, IGT25 was also divided into small local regions with somewhat different activity or Tm values (Fig. 3C and fig. S9). These observations indicate that, although the houses on each island share relatively similar enzymatic characteristics, the PETase candidates could not be completely separated into islands with unique characteristics, emphasizing the importance of density-based stratified sampling in enzyme screening.

As a result of the secondary contest, four additional qualified houses (H1501 from IGT25 and H1476, H1496, and H1499 from IGT3) were identified, approaching a final total of eight qualified houses, including the four houses found in the primary contest (H1319 from IGT158, H1533 from IGT25, and H1363 and H1484 from IGT3) (Fig. 3, B and C).

### Two highly talented houses and comparison with benchmarks

To select the most talented house among the eight qualified houses discovered from the primary and secondary contests, we conducted the final evaluation contest by measuring the PET hydrolysis activity of these houses at various temperatures using amorphized post-consumer PET powder (*19*). Of the eight houses, two (H1319 and H1501) exhibited outstanding PET hydrolysis activity across the range of temperatures tested; H1319 demonstrated the highest activity from low to moderate temperatures and H1501 exhibited the highest activity at high temperatures (fig. S10 and S11). Taken together, of 158 houses that participated in the three contest rounds, H1319 from IGT158 and H1501 from IGT25 were selected as the most talented houses with a balanced combination of PETase characteristics, including high catalytic ability and high thermal stability. Because the genes coding for H1319 and H1501 are originated from microbial strains of *Micromonospora pattaloongensis* DSM 45245 (*28*) and *Kutzneria buriramensis* DSM 45791 (*29*), we refer to these PET depolymerases as Mipa-P and Kubu-P, respectively.

We compared the enzymatic characteristics of Mipa-P and Kubu-P with those of the benchmarks CaPETase, IsPETase, and LCC. For a detailed comparison, we investigated initial rate and depolymerization extent as indicators of PET hydrolysis speed and the thermal-acidic tolerance of the enzymes. In particular, the apparent initial rate and the depolymerization extent were calculated by measuring the PET hydrolysis rate during the initial 3 h and the total amount of monomers released after 168 h, respectively, at temperatures ranging from 30 to 70 °C (Fig. 4A). In these experiments, initial rate tended to increase with higher temperatures, but started to decrease beyond a certain temperature close to Tm for Mipa-P, CaPETase, and IsPETase (Fig. 4B). Importantly, this pattern of increase for initial rate according to the temperature was similar for all enzymes, indicating that enzymes that are generally more active at low temperature will still exhibit more rapid activity at higher temperatures within their tolerance range.

**Figure 4.**
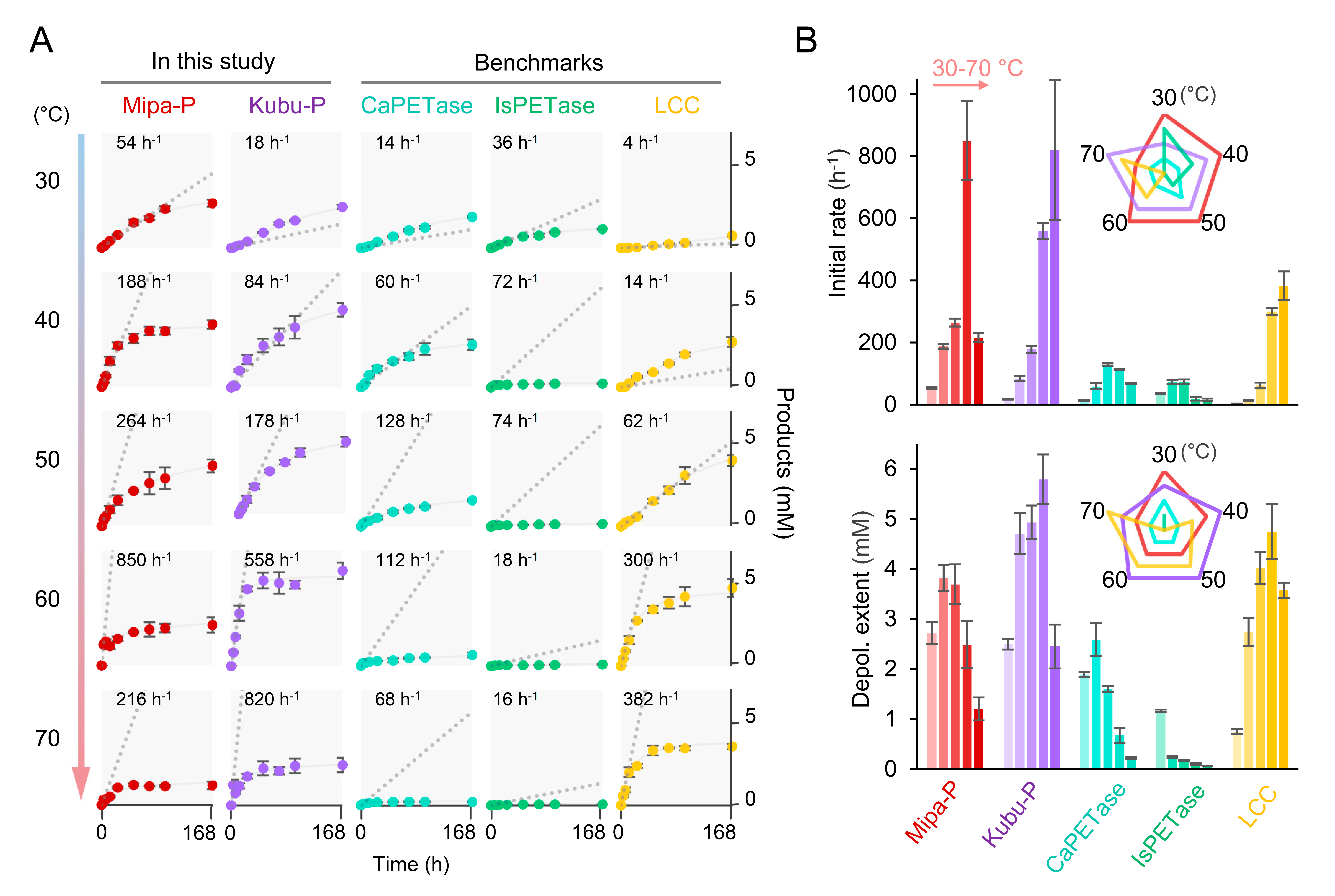
Comparison of Mipa-P and Kubu-P with benchmarks. (**A**) The initial rate and depolymerization extent of Mipa-P and Kubu-P are compared with those of benchmarks. Time course activities of the enzymes are measured at temperatures ranging from 30 to 70 °C. The initial rates correspond to the apparent velocity for 3 h and are indicated by a gray dashed line with their values. Data are presented as mean values□±□SD (n = 3). (**B**) Initial rate and depolymerization extent of enzymes along temperatures. Radar charts of initial rate and depolymerization extent ranks along temperatures by enzymes are also displayed. The rank 1 to 5 are shown with axes in direction towards the center.

Of the five PETases tested, Mipa-P exhibited the most outstanding initial rate across all temperatures except 70 °C, where it was outperformed by Kubu-P, which has a much higher Tm than Mipa-P (Fig. 4, A and B). Kubu-P also had a much higher initial rate than the benchmarks at all temperatures. In terms of depolymerization extent, Kubu-P exhibited the best performance at most temperatures, followed by Mipa-P and LCC, which had the highest depolymerization extent at 30 and 70 °C, respectively (Fig. 4, A and B). LCC generally exhibited slow-and-steady characteristics, with the lowest depolymerization extent at 30 °C, but this increased with higher temperatures, reaching the highest depolymerization extent at 70 °C among the PETases tested (Fig. 4B). These results indicate that the two PETases, Mipa-P and Kubu-P, discovered in the PETase contests described in the present study are potent PETases that demonstrate a high decomposition speed and duration when compared to benchmarks.

### Mipa-P^M19^ and Kubu-P^M12^ variants

Although Mipa-P and Kubu-P were shown to be more potent PETase templates than the benchmarks, their PET degradation ability remained insufficient. To understand their molecular features as engineering frames, we determined their crystal structures (table S2). Structural alignment with the benchmark enzymes CaPETase, IsPETase, and LCC and mutagenesis experiments revealed active site similarity and residual uniqueness near the Serine-Histidine-Aspartate catalytic triad (figs. S12 to S16). Mipa-P and Kubu-P share most of the local sub-network structures of CaPETase (*19*) but do not have the wobbling tryptophan mechanism of IsPETase (30) (figs. S12 and S16; text S1). However, the Mipa-P and Kubu-P templates do not appear to have sufficiently evolved an optimal structure for *in-vitro* depolymerization (fig. S14), suggesting that there is room for the introduction of artificial mutations to these templates.

We consequently sought to improve the thermal stability of these enzymes while maintaining or increasing their activity levels. Candidates for engineering points were generated through cross-template information for the various houses characterized in the present study. Mutation points exhibiting positive effects were applied to wild-type or intermediate variants generated during the engineering process to produce the Mipa-P^M19^ and Kubu-P^M12^ variants with enhanced properties (figs. S17 and S18). Mipa-P^M19^ and Kubu-P^M12^ contained a total of 19 and 12 amino acid substitutions, respectively, including single amino acid mutations and disulfide bond introductions (Fig. 5A). The thermal stability of these two variants was significantly higher, with Mipa-P^M19^ achieving a Tm of 93.5 °C, representing an increase of 22.9 °C compared with wild-type counterpart (figs. S17 and S19). For Kubu-P^M12^, its Tm_B_ was not measurable within the temperature range of 0-99.9 °C at a scanning rate of 0.1 °C·s^-1^, indicating that its Tm_B_ may be higher than 99.9 °C (figs. S18 and S19). Both the initial rate and depolymerization extent of Mipa-P^M19^ and Kubu-P^M12^ were also significantly higher at both 50 and 70 °C compared with their wild-type versions (fig. S20), illustrating the enhanced enzymatic properties of these variants.

**Figure 5.**
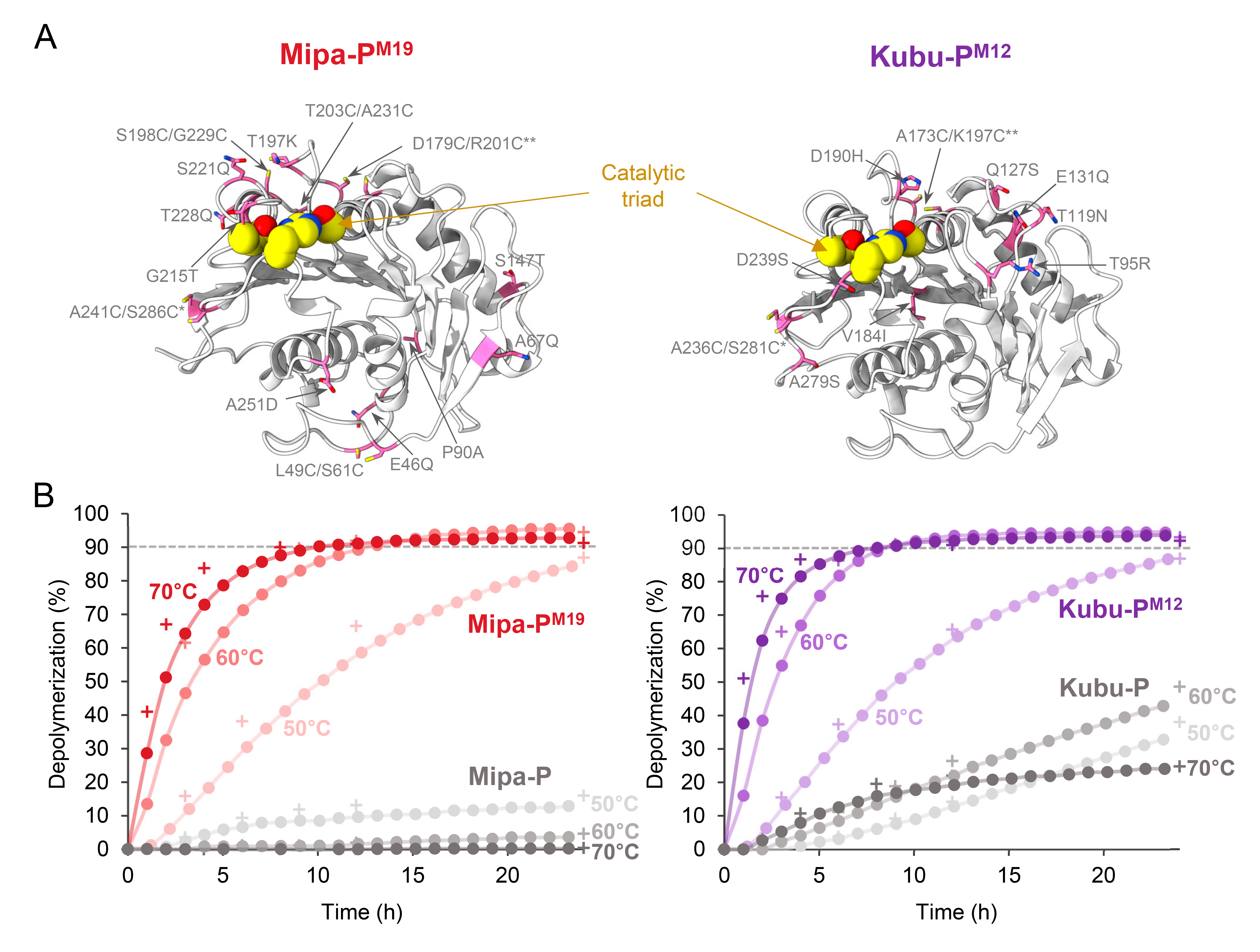
Performances of Mipa-P^M19^ and Kubu-P^M12^. (**A**) Crystal structures of Mipa-P^M19^ and Kubu-P^M12^. The structures are shown as gray cartoon models, and the mutation points are highlighted by sphere models and labeled. (**B**) Comparisons of Mipa-P^M19^ and Kubu-P^M12^ with their wild-type counterparts. The PET depolymerization experiments are performed in a pH-stat bioreactor. The graphs of linked circles or crosses indicate degree of depolymerization calculated by monomer release (HPLC sampling) or ester bond cleavage (NaOH consumption), respectively. The level of 90% degree of depolymerization is indicated by a gray dashed line.

To investigate the performance of Mipa-P^M19^ and Kubu-P^M12^ in a pH-stat reaction that allows complete depolymerization (*2*), we compared them with their wild-type counterparts in a bioreactor with reaction conditions of 2% (w/w) PET powder and 3 mg_enzyme_·g ^-1^ enzyme at 50-70 °C. The improved performance of these two variants in the agitated pH-stat bioreactor was more striking than in the closed reaction (Fig. 5B). While both wild-type enzymes achieved less than 50% depolymerization after 24 h, the engineered variants degraded more than 90% of the PET within 24 h at all temperatures tested (Fig. 5B). Mipa-P^M19^ was slightly faster than Kubu-P^M12^ at 50 °C after 12 h but, at 60 and 70 °C, the depolymerization rate of Kubu-P^M12^ was higher, with depolymerization of 51.2% and 75.8% after 1 and 2 h at 70 °C, respectively (Fig. 5B). We also tested the two engineered benchmarks, Hot-PETase (*15*) and LCC-ICCG (*2*), under the same bioreactor conditions. Hot-PETase could not achieve a final degree of depolymerization of 90% or higher at any of the temperatures tested, while LCC-ICCG achieved a final degree of depolymerization of 90% at 60 and 70 °C, but could not reach 90% at 50 °C, possibly due to its lower depolymerization activity than the Mipa-P^M19^ and Kubu-P^M12^ (fig. S21). These results indicate that the two engineered variants, Mipa-P^M19^ and Kubu-P^M12^, have excellent PETase characteristics, with Kubu-P^M12^ in particular exhibiting both the highest depolymerization activity and final degree of depolymerization of the PETase variants tested.

### High tolerance to high PET loads of Kubu-P^M12^

Industrial conditions often represent a harsh environment for enzymes due to the high temperatures, high substrate loads, and use of organic solvents to maximize the productivity (Fig. 6A). In particular, it has been reported that enzymes with a higher activity than LCC-ICCG under mild conditions exhibited a low final degree of depolymerization when a high substrate loading of 16.5% (w/w) is used (31). Thus, we evaluated the tolerance of Mipa-P^M19^ and Kubu-P^M12^ to high substrate loads with reaction conditions of 20% (w/w) PET and 1 mg_enzyme_·g ^-1^ at 70 °C in comparison with LCC-ICCG. Of the three engineered variants, Kubu-P^M12^ showed the highest degree of depolymerization, reaching 45.2% in the first hour, approximately 1.5 and 3.5 times higher than Mipa-P^M19^ and LCC-ICCG, respectively (Fig. 6B). Kubu-P^M12^ also achieved a degree of depolymerization of 90% within 8 h. However, the activity of Mipa-P^M19^ slowed after 2 h, and the final decomposition rate at 24 h remained at approximately 40% (Fig. 6B). LCC-ICCG maintained its activity more consistently than Mipa-P^M19^, but its final depolymerization rate at 24 h was still limited to 80% (Fig. 6B). The difference in the PET degradation behavior of Kubu-P^M12^ and LCC-ICCG was confirmed using two biologically independent bioreactor trials. Under harsher conditions, with a PET loading of 30% (w/w) and an enzyme supplement of 0.58 mg_enzyme_·g ^-1^, the difference in performance between Kubu-P^M12^ and LCC-ICCG was even greater. Kubu-P^M12^ depolymerized 90% of the PET in 24 h, while LCC-ICCG achieved less than 40% depolymerization after 48 h (Fig. 6C). Based on these results, we conclude that Kubu-P^M12^ is an excellent PETase variant in terms of both depolymerization speed and duration when compared to other engineered PETase variants.

**Figure 6.**
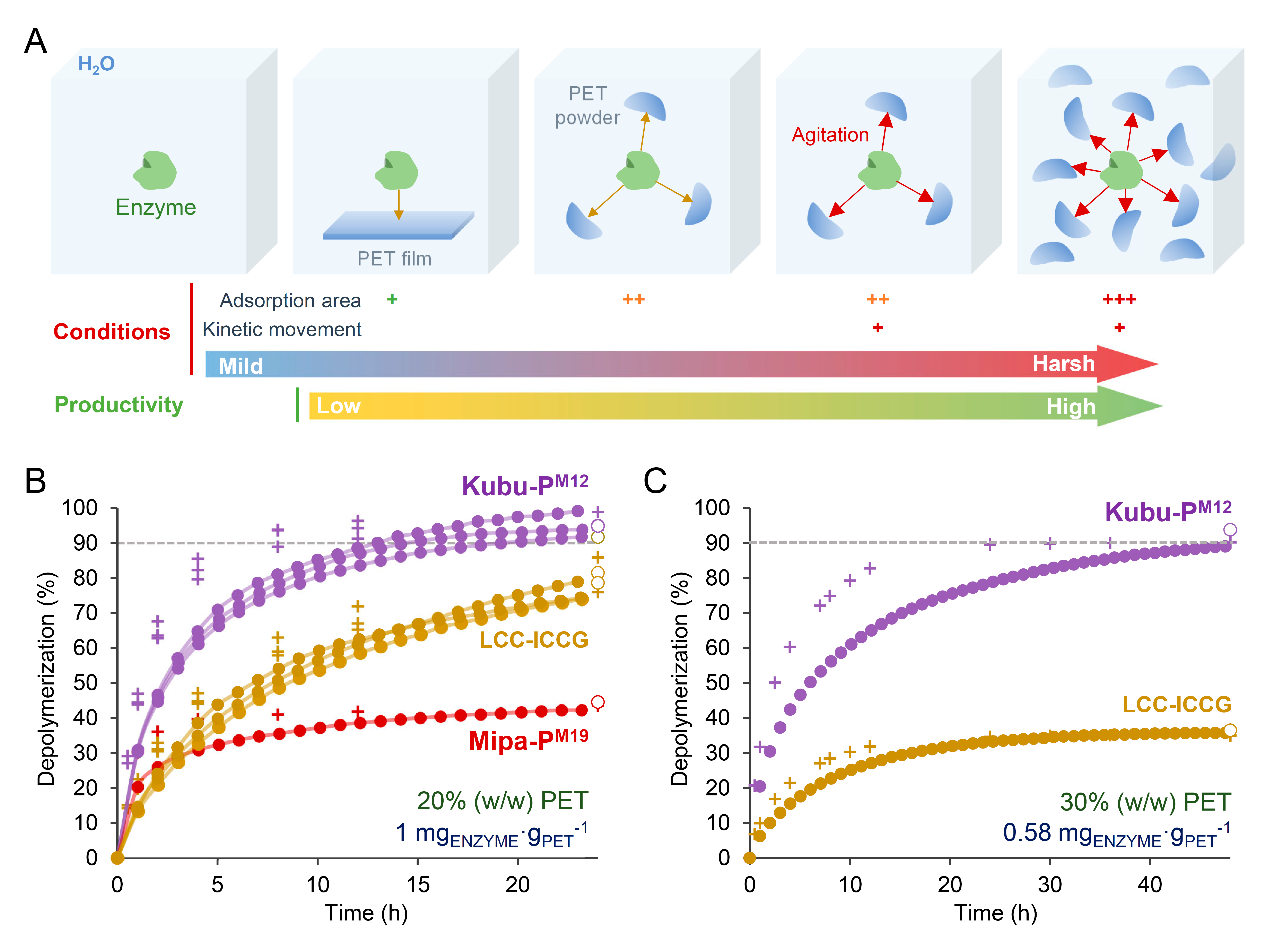
Kubu-P^M12^ in high PET loads. (**A**) Schematic diagram of conditional effects on productivity. Pulverization and high substrate load increase the material’s surface area that will enhance protein adsorption onto the surface. Agitation in a reactor also increases the kinetic energy of the space, enhancing adsorption and dissociation events for a specific time. These conditional effects can increase space-time yield but make the environment harsher for protein. (**B**-**C**) Depolymerization performance by the enzymes in high substrate load conditions. The bioreactor operation with PET loads of 20% (**B**) and 30% (**C**) are drawn as in Fig. 5B. The final weight loss for each reaction is also shown as a white circle.

### Enzyme-catalyzed PET glycolysis using Kubu-P^M12^

We then tested whether Kubu-P^M12^ could maintain its structure in ethylene glycol (EG) for enzyme-catalyzed PET glycolysis (*32*), which can produce bis(2-hydroxylethyl) terephthalate (BHET) via the reaction of the hydroxyl-groups in EG with the tetrahedral intermediate formed during the common de-acylation reaction (Fig. 7A). Thus, PET degradation experiments were conducted in a water-depleted EG (>95%) solvent at 40°C using the PETase variants Mipa-P^M19^, Kubu-P^M12^, LCC-ICCG, and Hot-PETase. Significant levels of BHET as the main product (>98%) were observed with the use of Mipa-P^M19^, Kubu-P^M12^, and LCC-ICCG after incubation for 24 h, while Hot-PETase produced a negligible amount of BHET (Fig. 7B; figs. S22 and S23). In particular, Kubu-P^M12^ exhibited the highest glycolysis activity, with a more than 4-fold higher activity compared to Mipa-P^M19^ and LCC-ICCG, which exhibited similar activity to each other (Fig. 7B). When the tolerance of these variants to EG was investigated at various temperatures, Mipa-P^M19^ and Kubu-P^M12^ generally maintained their glycolysis activity at 40 and 50 °C for 10 days in EG, while LCC-ICCG was more susceptible to EG (Fig. 7B). These observations indicate that Mipa-P^M19^ and Kubu-P^M12^ had much higher tolerance to EG than the benchmarks. When enzyme-catalyzed PET glycolysis of Mipa-P^M19^ and Kubu-P^M12^ was tested in the bioreactor, the use of Kubu-P^M12^ resulted in the highest production of BHET reported to date producing approximately 21 mM BHET with a monomer fraction purity of 98%, which corresponded to a degree of depolymerization of 8% within 48 h at 40 °C (Fig. 7C). These results demonstrate that Kubu-P^M12^ satisfies the characteristics required for enzyme-catalyzed PET glycolysis, such as tolerance to EG, high temperatures and high PET loads, PET decomposition activity, and enzyme reactivity to EG as a second substrate, thus demonstrating the potential for use in enzyme-catalyzed PET glycolysis for the biological recycling of PET.

**Figure 7.**
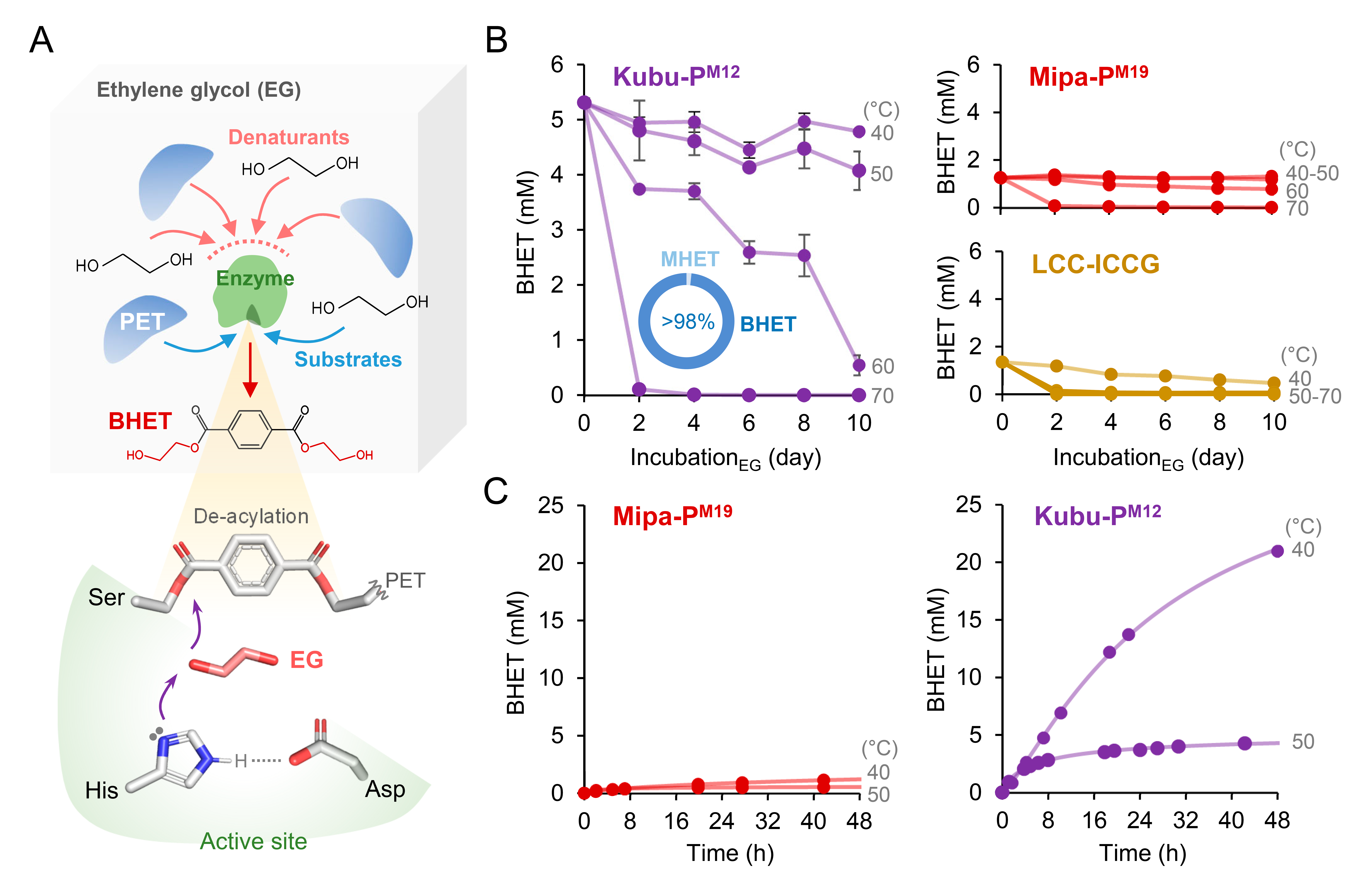
Use of Kubu-P^M12^ in enzyme-catalyzed PET glycolysis. (**A**) Schematic diagram of enzymatic glycolysis and reaction mechanism at the de-acylation step. Under the ethylene glycol (EG) solvent, both PET and EG can act as denaturants and substrates of the enzyme. (**B**) Enzyme-catalyzed PET glycolysis activity and inactivation patterns of the three enzymes by EG. The mol fraction of BHET, mono(2-hydroxyethyl) terephthalate (MHET), and negligible TPA at 1-day reaction are shown in a donut chart. The temperature condition is labeled at the right side. Data are presented as mean values□±□SD (n = 3). (**C**) Enzyme-catalyzed PET glycolysis using Mipa-P^M19^ and Kubu-P^M12^ in a bioreactor scale. The temperature condition is labeled as in (B).

## Discussion

In this study, we navigated a natural landscape of ester-based plastic hydrolases using a sequence homology network of PHILOSOPI. Although known esterases outside of this family were not considered due to their relatively weak activity (*18*, *33*), the natural PETase screening based on accumulated sequence data in our study was geographically extensive and widest reported to date. The available isolation source data from BioSample provides the environmental origins of many potential PETases, such as the rhizosphere and mangrove soil (e.g., I_2_, IGT3, I_7_, I_21_, IGT25, I_50_, I_54_, I_60_, I_66_, I_135_, and I_154_), marine and water sediment (I_5_, I_13_, I_16_, and I_76_), and hot compost (I_20_). The landscape profiling of this family revealed that enzymes acting on a solid-state PET substrate exist on a variety of islands, with at least 90 houses discovered with PETase activity and 17 benchmarks across 43 islands. Of these templates, 82 houses from 33 islands have not, to the best of our knowledge, been previously identified as a homologous group in past studies, indicating that 90% of the templates were unrecognized. Our stratified and cluster sampling methods sought to trace the fitness landscape and may provide a bird’s-eye view of the natural distribution of these enzymes. We highlighted three islands, IGT3, IGT25, and IGT158, that contained houses with high PET decomposition potential, and two PETases, Mipa-P and kubu-P, with the characteristics of potent PETases were discovered on IGT25 and IGT158, respectively.

Both bacterial strains *M. pattaloongensis* (*28*) and *K. buriramensis* (*29*), containing genes coding for Mipa-P and Kubu-P, respectively, were isolated from tropical deciduous areas in Thailand, which is in line with the general belief that the efficient evolution of depolymerases occurs in regions with an abundance of ester bioresources in a rainforest climate. However, the genomes of these two microbes do not appear to have a 1,2-dihydroxy-3,5-cyclohexadiene-1,4-dicarboxylate dehydrogenase-coding gene, suggesting that they are not likely to assimilate terephthalate via a known aerobic metabolic pathway (*1*). These observations provide evidence that closely functioning genes that can be exploited to respond to artificial substances do exist. It is also worth noting that the optimal growth temperatures for most microorganisms containing enzymes with a high Tm_B_ are not high. Although gene mining that is limited to bacteria growing at high temperatures is effective in identifying enzymes with a high thermal stability, the strategy may overlook the potential enzymes that are more effective in harsh extracellular environments.

The increase in initial rate with higher temperatures before Tm (Fig. 4B) indicates that low temperature activity better represents catalytic efficiency than does high temperature activity, which is affected by the duration of catalyst activity. However, current research directions place greater weight on slow-and-steady thermophilic templates such as LCC and BhrPETase, because differences in durability are more prominent above the glass transition temperature of PET (*34*) than are differences in catalytic efficiency between enzymes. Indeed, as shown by the IsPETase (*15*) and CaPETase variants (*19*), the strategy of increasing the durability of templates with high activity at low temperatures has only been semi-successful, increasing the speed of PETase but still lacking durability under harsh conditions. Therefore, we believe that our discovery approach, which simultaneously evaluates low-temperature activity and thermal stability, is crucial to finding highly potent PETases such as Mipa-P and Kubu-P, which offer both high catalytic capability and thermal stability. In particular, Kubu-P appears to be a more optimal template for the engineering of a variant with a high tolerance to high temperatures, high substrate loads, and the use of water-depleted EG as a solvent.

In the PET supply chain and downstream recycling processes, enzymatic glycolysis offers an economic advantage because it does not require a prior esterification process, or titrants for pH adjustment and it has a negligible water content, which usually requires high latent heat exchange for purification. A previous study using the commercial enzymes *Candida antarctica* lipase B and HiC could not achieve a sufficient depolymerization rate or amount even under optimal conditions (*32*). In contrast, the apparent rate of glycolysis using Kubu-P^M12^ was approximately 109 h^-1^ at 40 °C (Fig. 7C), a speed comparable to that of hydrolysis. In addition, the observation that the BHET mol fraction of the released monomers was above 98% in a water-depleted EG solvent indicates that PET decomposition by Kubu-P^M12^ occurred solely via enzymatic glycolysis (fig. S22). This study demonstrates that enzymatic PET glycolysis with a significant degradation rate and amount can be achieved using an enzyme with a high tolerance to EG and high activity and durability for hydrolysis.

Nevertheless, there are still important limitations that need to be overcome for industrial applications. For example, the water added to the enzyme solution and water vapor appeared to produce a small amount of anionic chemicals such as mono(2-hydroxyethyl) terephthalate in our enzymatic glycolysis experiments (fig. S23). We observed that the addition of a buffer such as Tris and imidazole increased the enzymatic glycolysis activity (fig. S24), suggesting that these anions had a negative effect on enzymatic glycolysis. Thus, the design of the reactor system and the use of supplements need to be carefully considered. Another issue is that the glycolysis reaction involving Kubu-P^M12^ at 40 °C produced up to 30 mM BHET (fig. S25), which is close to its solubility in EG at this temperature (*35*), and other reactions also converged to a certain BHET concentration. This suggests that PET decomposition via enzyme-catalyzed glycolysis reactions cannot proceed beyond the saturation curve for BHET in EG. However, contrary to expectations based on the temperature-dependent solubility of BHET, the degree of depolymerization by Kubu-P^M12^ at 50 °C was lower than that at 40 °C (Fig. 7C), indicating that its tolerance to EG remains insufficient. Thus, further engineering of Kubu-P^M12^ for higher EG tolerance would allow glycolysis reactions to occur at higher temperatures, enhancing both its depolymerization rate and degree. This may also act as a starting point for the development of a BHET recovery process based on cooling crystallization (*36*), by producing BHET at concentrations sufficient to form crystalline nuclei at low temperatures (fig. S26). Overall, however, we believe that our landscape profiling of PET depolymerases provides a new perspective for understanding the natural distribution of PETases and for more efficient enzymatic PET depolymerization through hydrolysis, glycolysis, and potentially other methods.

