## Supplementary material for "Landscape profiling of PET depolymerases using a natural sequence cluster framework": fig. S1, fig. S26, table S1, table S2

**The PDF file includes:**

Materials and Methods
Text S1
Figs. S1 to S26
Tables S1 and S5
References

### Materials and Methods

#### Library construction, Clustering, and Visualization

The initial set of 25,418 non-redundant sequences were retrieved from Genbank Release 235 at Dec. 2019 with PSI-BLAST searches using seed sequences. The value of search threshold for each seed are summarized in Table S1. The subset library with 2,064 sequences was constructed, by reducing the seed sequences with a criterion of polyesterase-lipase-cutinase family and partial gene removal, and then clustered. The clustering algorithm uses a Wilbur and Lipman (*1*) pairwise distance matrix (values ranging from 0 to 1) equipped in clustalo (*2*), individually ($N^2$) arranged by distance, and cuts off the data at a distance of significant density to reduce meaningless relationship entanglement. The histogram data of all sequences are shown in Fig. S3. The cut-off strategy was based on the high probability of existence of cluster-able groups at a high-dense distance in histogram. Unlike the conventional k-Nearest Neighbor (kNN) clustering, we used flexible k numbers of neighbor utilizing the complexity of distance-dependent histogram pattern for each sequence, using a python code with eight hyperparameters (Fig. S4). It can make directional edges, but we considered them undirected edges. The network was visualized in Cytoscape (*3*) software via the Prefuse force-directed-cl layout, which provide forces to spread relative position of sequences along homology inside the clusters. The layout parameters and edge weight settings are summarized in Table S3. The procedures in this work can visualize the high dimensional sequence homology matrix data into easily recognizable 2-dimensional graph.

Considering the diversity of signal peptide sequence, artificial mutation, and dual prediction of start codon in non-redundant sequences, we made a concept of House merging the diverse sequences into their main representing templates. For the housing, we used accession codes to get entry of Identical Protein Group (IPG) using Entrez software (*4*) combined with utilization of idmapping_selected.tab file of UniProtKB database (24, Feb, 2022), and then merged some sequences if they have inclusion sequences or same accession mapping in the databases. The id value of the library, the numbers of Island and House, accession code, and other analytical and experimental data are summarized in Data S3. The landscape of the network data is also available in Data S7. The phyla of each bioresources were analyzed by taxonomy data in NCBI, genus and species names, with _search_scientific table of The Catalogue of Life (COL) database (*5*). For Biosample (*6*) analysis, we accessed to the database with accession codes of each nucleotide sequences in IPG database using Entrez.

#### Genetic source preparation

Genetic information is in Data S3. For cloning, we obtained double-stranded DNAs from synthetic (codon-optimized for *E. coli*, *Twist Bioscience*) or genomic (*Marinactinospora thermotolerans*, *Roseateles depolymerans*, and *C. aurantiacum*) gene sources. The genes were amplified by DH10B strain culture or polymerase chain reaction (PCR) with oligonucleotides listed in Table S4, and inserted them into commercial vectors, pET21b, pET22b, or pET30a, with restriction enzymes Nde1 and Xho1. Signal peptides of the genes were predicted by SignalP 5.0 (*7*), and the expressed protein construct with secondary structure prediction and the signal peptide information were designed and recorded at Data S3. The pET21b-based plasmids for LCC-ICCG and Hot-PETase were synthesized and subcloned with the enzymes (Table S5). Site-directed mutagenesis for variants were performed by PCR with oligonucleotides listed in Table S4 and all variants were verified by Sanger sequencing.

**Recombinant protein expression and purification**

The constructed plasmids were transformed into the *E. coli* strain BL21(DE3)-T1$^R$. To help formation of disulfide bonds, the variants of Mipa-P and Kubu-P, Mipa-P$^{M19}$ and Kubu-P$^{M12}$, LCC-ICCG, and Hot-PETase were produced with cultivation of Rosetta gami-B strain transformed by their plasmids. The transformed cells were cultured in 1 L of lysogeny broth (LB) medium containing 100 mg $L^{-1}$ Ampicillin or 50 mg $L^{-1}$ Kanamycin, depending on antibiotic resistance of each vector, at 37 °C. When the optical density of the culture broth reached $OD_{600}$ of 0.6, 0.5 mM isopropyl β-D-1-thiogalactopyranoside (IPTG) was added to induce protein expression in T7-LacO promoter system. After further culture for 20 h at 18 °C with shaking at 120 rpm, the cells were harvested via centrifugation at relative centrifugal force (RCF) of 4,000 xg for 15 min at 20 °C.

The cell pellet was resuspended in lysis buffer (40 mM Tris-HCl, pH 8.0) and cell lysis was conducted via ultrasonication. The cell lysate was centrifuged at RCF of 13,000 xg for 30 min, and the supernatant was obtained. For affinity chromatography, the supernatant was applied to a Ni-NTA agarose column (Qiagen) that was equilibrated with the lysis buffer. After washing with the lysis buffer containing 30 mM imidazole, the bound proteins were eluted with the 300 mM imidazole in lysis buffer. All protein purification steps were conducted at 4 °C. The purities of the eluted proteins were analyzed by sodium dodecyl sulphate-polyacrylamide gel electrophoresis (SDS-PAGE) and Coomassie blue staining, and shown in Fig. S. Protein production titer of each protein was quantified using the solution by a BioTek™ Epoch microplate spectrophotometer and Gen5™ microplate data analysis software. For the primary, secondary, and final contest, the produced enzyme solutions were concentrated by Amicon Ultra Centrifugal filter unit (10,000 Da, *Millipore*, *Billerica*, *MA*, *USA*) to final concentration of 1 mg $mL^{-1}$. For the hydrolysis reactions in bioreactor, the solutions were concentrated to 5 mg $mL^{-1}$, and then applied to the reaction. To avoid precipitation of Mipa-P$^{M19}$ and LCC-ICCG during the concentration, sodium chloride is added to the eluted solution to reach 150 mM. For the feasibility tests and bioreactor experiments for enzyme-catalyzed PET glycolysis, we prepared enzyme solutions of 10.2, and 20 mg $mL^{-1}$, respectively.

For crystallization, further purification through size exclusion chromatography was performed using a HiPrep 26/60 Sephacryl S-100 HR column (320 mL, *GE Healthcare Life Sciences*) equilibrated with the lysis buffer. The eluted proteins were similarly concentrated to 10 mg $mL^{-1}$ and prepared for next crystallization steps.

**PET powder preparation**

Molecular weights, such as number-average (Mn), weight-average (Mw), highest peak (Mp), and average diameter of the used PET powders were analyzed using gel permeation chromatography and laser diffraction particle size analyzer, respectively, certified by Korea Polymer Testing & Research Institute (KOPTRI). The microcrystalline PET powder pulverized from a commercial bottle, prepared as in Sagong *et al*. (*8*) (average diameter of 182.4 μm, 8,800 Mn, 23,800 Mw, 19,900 Mp, and 17.4% crystallinity), was used for the primary and secondary contests of PETase. The amorphized post-consumer powder (average diameter of 247.8 μm, 7,060 Mn, 16,700 Mw, 13,400 Mp, and 9.9% crystallinity) from transparent flake (Yoochang R&C, Republic of Korea), prepared as in Hong *et al*., (*9*) with 200 μm sieving was used for final contest, further comparisons, bioreactor, and enzyme-catalyzed PET glycolysis experiments. The crystallinity values were estimated from differential scanning calorimetry data of the previous articles.

**PETase activity assays**

To measure PETase activity for the contests, 15 mg of PET powder was soaked in 1 mL reaction solution with 50 mM glycine-NaOH (pH 9.0) and 500 nanomolar enzyme (approximately 13~16 μL of 1 mg $mL^{-1}$ enzyme solution). The primary and secondary contests were performed at 30 °C for 3 days, but final contest was held at 30~70 °C for 12 h. For the comparison of time-course activity, the reactions were conducted for 3, 6, 12, 24, 48, 72, 96, or 168 h at 30~70 °C. All the reaction was terminated by eliminating substrate using a PVDF syringe filter (0.22 μm) and the main products, terephthalic acid, mono(2-hydroxyethyl) terephthalic acid, and bis(2-hydroxylethyl) terephthalate, were quantified via analytical procedure of high-performance liquid chromatography (HPLC). The pH-dependent activities of Mipa-P, Kubu-P, and LCC were measured in the same mixture containing 100 mM Britton-Robinson universal pH buffer, instead of 50 mM glycine buffer, by relative absorbance at 270 nm.

**HPLC Analytical procedure**

Reaction samples were analyzed using a CBM-20A (*Shimadzu*, *Japan*) connected to a UV/Vis detector (SPD-20A) and a C18 column (*Shimadzu Shim-pack GIST C18, 3 μm*, *4.6 × 50 mm*) guarded with a cartridge (*YMC ODS-A*, *3 μm*). Mobile phases A (0.1% formic acid) and B (100% methanol) were used with isocratic flow of 33% concentration of B at a flow rate of 1 mL $min^{-1}$ and a chamber temperature of 40 °C. The chromatograms were detected at 260 nm, and subtracted with fitted baseline using arPLS algorithm (*10*) of Rampy (*11*), and then the peak areas were integrated. The amounts of main products, terephthalic acid (*Sigma-Aldrich*, *MO*, *USA*, *Cat. No.: 185361*), mono(2-hydroxyethyl) terephthalic acid (*Ambeed*, *IL*, *USA*, *Cat. No.: A875019*), and bis(2-hydroxylethyl) terephthalate (*Sigma-Aldrich*, *MO*, *USA*, *Cat. No.: 465151*), were calculated using standard curves of the integrated area.

**Differential scanning fluorimetry**

Fluorescent intensity for enzyme melting curve was measured using the protein thermal shift™ dye kit (*Applied Biosystems*) in a Quantstudio™ 5 Real-time PCR instrument (*Thermo Fisher Scientific*). Each reaction mixture contained 5 µL of 1 mg $mL^{-1}$ enzyme solution, 100mM $Na_2HPO_4$-HCl (pH 7.0), and 2.5 µL of 1× dye, and distilled water up to the final volume of 20 µL. Signal changes were monitored by increasing the temperature from 25 °C to 99 °C with 0.1 $°C·s^{-1}$ rate. The Boltzmann or derivative melting temperatures were calculated from the intensity change based on Boltzmann equation fitting or peak of the first derivative curve, and all data is shown in Fig. S7, S15, and S19. We referred Boltzmann melting temperature as Tm throughout the manuscript.

**Crystallization**

Crystallization conditions of Mipa-P, Mipa-$P^{M19}$, Kubu-P, Kubu-$P^{P185V}$, and Kubu-$P^{M12+P185V}$ were screened using the sitting-drop vapor diffusion method (drop: 1.0 µL of the protein solution + 1.0 µL of reservoir solution; reservoir: 50 µL) with reagents of PEG/Ion HT™ and Index HT™ (*Hampton Research*), Wizard™ Classic I&II HT96 (*Rigaku*), Structure Screen 1&2 HT96 (*Molecular Dimensions*) and Wizard Cryo I&II HT96 (*Rigaku*) at 20 °C. The crystals of Mipa-P and Mipa-$P^{M19}$ appeared under conditions of 20% polyethylene glycol (PEG) 8,000 and 100 mM N-cyclohexyl-3-aminopropanesulfonic acid (CAPS)-NaOH (pH 10.5), and 20% PET3350 and 40 mM citric acid + 60mM BIS-Tris propane (pH 6.4). The crystals of Kubu-P and Kubu-$P^{P185V}$ appeared in a condition of 40% PEG 300, 100 mM imidazole-HCl (pH 8.0), and 200 mM zinc acetate. The crystals of Kubu-$P^{M12+P185V}$ appeared in a condition of 16% PEG 3,350, 100 mM sodium acetate tribasic (pH 4.6), and 2% (v/v) Tacsimate (pH 4.0). The best quality crystals were soaked in reservoir solution with 20% (v/v) glycerol for cryoprotection, fished out with a loop, and mounted on a goniometer in beamline.

**Structure determination**

Data were collected using a 7 A beamline of the Pohang Accelerator Laboratory (Republic of Korea) (*12*) with ADSC Q270 CCD or EIGER 9M detector under 100 K. The diffraction data were indexed, integrated, scaled, and processed using XDSGUI (*13*) and determination of space group with screw axis and resolution cutoff were performed using Aimless (*14*). The Mipa-P crystals belonged to a space group $P2_12_12_1$ with a unit cell parameter of 49.07 a, 103.41 b, 131.88 c, 90.0 α, 90.0 β, and 90.0 γ, with highest resolution of 1.34 Å, and there are three molecules in the asymmetric unit. The Mipa-P$^{M19}$ crystals belonged to a space group $P12_11$ with a unit cell parameter of 42.62 a, 51.36 b, 107.65 c, 90.0 α, 94.3 β, and 120.0 γ, with highest resolution of 1.89 Å, and there is two molecules in the asymmetric unit. The Kubu-P crystals belonged to a space group $I4_132$ with a unit cell parameter of 159.1 a, 159.10 b, 159.10 c, 90.0 α, 90.0 β, and 90.0 γ, with highest resolution of 2.65 Å, and there are is molecule in the asymmetric unit. The Kubu-P$^{P185V}$ crystals belonged to a space group $I4_132$ with a unit cell parameter of 157.22 a, 157.22 b, 157.22 c, 90.0 α, 90.0 β, and 90.0 γ, with highest resolution of 1.7 Å, and there is one molecule in the asymmetric unit. The Kubu-P$^{P185V}$ crystals belonged to a space group *I*222 with a unit cell parameter of 64.96 a, 81.65 b, 83.73 c, 90.0 α, 90.0 β, and 90.0 γ, with highest resolution of 1.15 Å, and there is one molecule in the asymmetric unit.

The phases of the diffraction patterns of the Mipa-P and Kubu-P crystals were inferred by molecular replacement using MOLREP (*15*) and PHASER (*16*) in CCP4 (*17*) and Phenix (*18*) using the structure of CaPETase (PDB code: 7YM9) as a search model, and those of the variants were inferred by same method using their wild type structures. Structural models were built using Wincoot (*19*), and map refinements were performed using REFMAC5 (*20*). The statistics are summarized in Table S2. Kubu-P$^{M12}$ structure was prepared using a homology model of colabfold (*21*) from template structures of Kubu-P, Kubu-P$^{P185V}$, and Kubu-P$^{M12+P185V}$. The structures of Mipa-P, Mipa-P$^{M19}$, Kubu-P, Kubu-P$^{P185V}$, and Kubu-P$^{M12+P185V}$ were deposited in the Protein Data Bank, with PDB codes 8YTU, 8YTV, 8YTW, 8YTZ and 8YTY, respectively.

**Depolymerization via hydrolysis in bioreactor**

The customized bioreactor (*MARADO-PDA*, *BIOCNS Co. Ltd*, *South Korea*) and thermostat bath circulator (*RW3-1025*, *JEIO TECH*, *Daejeon*, *South Korea*) (*9*) were operated to test large-scale performance of enzymes in pH-stat hydrolytic PET depolymerization. The real-time values of pH during reaction were detected using a pH electrode (pH Sensor InPro3030/120, METTLER TOLEDO, Switzerland) and titrated by NaOH solution using PID controller to maintain pH 8.0. To consider stirring rate, diffusion of alkalinity and disodium terephthalate solubility, the titrant concentrations were used differently depending on the substrate load (1 M for 2% load, 7.15 M for 20% load, and 4 M for 30% load). For the reactions in a condition of 2% (w/w) PET and 3 $mg_{enzyme} \cdot g_{PET}^{-1}$ enzyme load, 3.06 g PET powder was mixed with a 150 mL solution containing 100 mM $Na_2HPO_4$ (pH 8.0) with a stirring rate of 300 rpm. Once the temperature reached the correct conditions, 1.836 mL of enzyme solution was added to the solution to initiate the reaction. For the reactions in a harsh condition of 20% (w/w) PET and 1 $mg_{enzyme} \cdot g_{PET}^{-1}$ enzyme load, 37.50 g PET powder and 7.5 mL of enzyme solution were added to the 150 mL solution with a stirring rate of 600 rpm. For the reactions in a harsher condition of 30% (w/w) PET and 0.58 $mg_{enzyme} \cdot g_{PET}^{-1}$ enzyme load, 64.29 g PET powder and 7.5 mL of enzyme solution were added to the 150 mL solution with a stirring rate of 600 rpm. Experimental data including temperature, pH change, titrant consumption, and revolutions per minute of impeller were recorded in real time on a personal computer. During the reactions, 500 μL aliquots were sampled using syringe, filtered through a PVDF syringe filter (0.22 μm, *SFPVDF013022CB*, *Futecs*, *South Korea*), diluted to optimal concentration, and analyzed using HPLC for quantifying the released monomers. The degree of depolymerization was calculated by two methods: ester bond cleavage ratio (NaOH consumption) and available monomer release ratio (NaOH consumption with terephthalic acid/mono(2-hydroxyethyl) terephthalic acid ratio). The weight loss values were determined by a weight change of 0.22 μm filter holder before and after filtration with complete drying.

**Depolymerization via enzyme-catalyzed PET glycolysis**

The all reactions for enzyme-catalyzed PET glycolysis were performed with 99.5% ethylene glycol (EG) solution (Guaranteed Reagents grade, *Daejung*, South Korea). For 1-day feasibility test, enzyme-EG mixture containing 5 μL of the 10.2 mg $mL^{-1}$ enzyme solution, 45 μL of the lysis buffer, and 950 μL of EG solution was added to the 50 mg of PET powder in a 1.5 mL microtube, and after mixing for 10 s with vortex mixer and centrifuging for 30 s with RCF of 20,000, the reaction was initiated. The reaction of the PET suspension in ~95% EG was conducted at 40 °C without shaking. To evaluate the tolerances of the three enzymes to EG, the mixture was incubated for 0, 2, 4, 6, 8, and 10 days at 40~70 °C. Then, the solution was loaded on 50 mg of the PET powder, and the 40 °C reaction was initiated by the same manner. For the buffer reagent test, we performed the enzyme-catalyzed PET glycolysis for 17 h with a suspension of 50 mg PET powder, 5 μL enzyme, 950 μL EG, and 45 μL buffer solution (100 mM Tris or imidazole, dissolved in EG). In this ~99.5% condition, the relative activities of LCC-ICCG in enzyme-catalyzed PET glycolysis were severely decreased compared to ~95% condition experiments (Fig. S24). For the reactions of enzyme-catalyzed PET glycolysis in the bioreactor at 40 and 50 °C, 7.5 g PET powder, 0.81 g Tris, and 0.75 mL of the enzyme solution were added to the 150 mL solution with a stirring rate of 300 rpm, the condition corresponds to 0.45% (w/w) PET 2 $mg_{enzyme} \cdot g_{PET}^{-1}$ enzyme load. It is note to worthy that EG is hygroscopic but water vapor level was not tightly controlled during reactions, especially for bioreactor experiments. The degree of depolymerization was inferred from produced BHET concentration with the mass per repeating unit of PET and EG molecular weight. The weight loss of the 100 h reaction by Kubu-$P^{M12}$ at 40 °C was 17%, determined by a weight change of 0.22 μm filter holder before and after filtration with complete drying.

### Supplementary Text

#### Text S1. Structural features of Mipa-P and Kubu-P

Overall shape of the two enzymes is quite similar to that of the known superficial hydrolases, which have six superficial loops constituting the active site: L1 to L6 (Fig. S12). Based on the docking models and the crystal structures of this enzyme class in complex with terephthalic acid or 2-hydroxymonoethylene terephthalate (*22*, *23*, *24*, *25*), the scissile ester bond between the PET monomers should be located on the catalytic serine of L3, and the linked terephthalate moiety is thought to extend toward a highly conserved tryptophan residue of L4. The tryptophan residues of the Mipa-P and Kubu-P enzymes, W193 and W189, whose positions are fixed by histidine residues on L5, H222 and H218 respectively, do not exhibit the wobbling mechanism in IsPETase (*22*) (Fig. S13). Combined with the case of CaPETase (*9*), the observations that the substrate binding tryptophan does not largely fluctuate even in the two more highly active enzymes suggests that the mechanism of IsPETase may not be an essential mechanism for high activity against glassy PET at moderate temperature. Unfixing the tryptophan by the H222S or H218S variants of Mipa-P or Kubu-P severely decreased their Tm values by 8.6 or 12°C, respectively, and consequently their activities (Fig. S14-15). To our interest, Mipa-P and Kubu-P were found to share most of the local sub-network structures in CaPETase (*9*). The unique phenylalanine anchor of the A-G-F local network in CaPETase (*9*) is also displayed on the L6 loops of Mipa-P and Kubu-P: F247 and F242, respectively (Fig. S13 and S16). Kubu-P has the same local network, but in Mipa-P it is A-A-F (Fig. S16). The Mipa-$P^{F247A}$ and Kubu-$P^{F242A}$ variants showed the decreased Tm values by 11.8 or 10.5°C, respectively (Fig. S14-15). Due to the stability difference between the two templates by ~20 °C in Tm, the activity of the Mipa-$P^{F247A}$ variant decreased by more than 80% on day 1 compared to wild type, while the Kubu-$P^{F242A}$ variant maintained its activity (Fig. S14-15). However, after 3 days, the Kubu-$P^{F242A}$ variant showed the decreased activity by 34% (Fig. S14-15). The observation that the three enzymes or IsPETase that can attack glassy PET at low temperatures happen to have the phenylalanine or the disulfide bond (*26*) at this region implies the importance of this region upholding the histidine residue of the catalytic triad. Mipa-P and Kubu-P also have the unique tryptophan residue of the W-L-G local network in CaPETase (*9*); they have W-I-G and W-F-G, respectively (Fig. S16). The substitutions of the tryptophan in Mipa-P and Kubu-P with alanine and glutamine residues of IsPETase and LCC showed the opposite results (Fig. S14-15), which appear to be due to the difference in the local network, I or F. The Mipa-$P^{W102A}$ and Mipa-$P^{W102Q}$ variants had the increased Tm values by approximately 3°C and the increased activities by ~10%, whereas the Kubu-$P^{W96A}$ and Kubu-$P^{W96Q}$ variants had the decreased Tm and activity values by ~7°C and 80%, respectively (Fig. S14-15). In addition, Mipa-P has H167 but Kubu-P has W161 at the first position of L3, mostly conserved as histidine or tryptophan in other cutinases (Fig.

S12-13). Uniquely, Mipa-P has E100 on L1 that interacts by a 2.7 Å distance hydrogen bond with a delta nitrogen atom of the H167 (Fig. S13). The fact that the Mipa-P$^{E100S}$ and Mipa-P$^{E100A}$ variants showed similar activity but ~5.8 °C higher Tm values (Fig. S14-15) suggests that the hydrogen bond seems to affect the activity but not critical, and rather be negative for stability. The Mipa-P$^{H167W}$ variant showed a large decrease in activity that suggests a physical backlash between E100 and H167W, and the Kubu-P$^{W161H}$ variants was also worse than its wild type (Fig. S14-15), indicating the optimum of the residues in their complementary structure.

L4 loop was reported to be important region for activity in many PETases and their variants (*8*, *27*, *28*). L4 contacts to the core via three helices, which shows diverse and complex force network including salt bridges, polar interactions, and van der Waals forces (Fig. S12). A conserved non-polar residue, mostly phenylalanine or tryptophan, in the L4's hydrophobic patch underlies the loop conformation weakening its structural disorder (Fig. S12-13). Unlike the similar orientations of the phenylalanine residues, such as F195, F191, and F196, in three enzymes, Kubu-P, IsPETase, and LCC, the indole rings of the tryptophan residues, W199 and W200, in Mipa-P and CaPETase showed a significant difference by 70° rotation; that in CaPETase is connected toward the inner core of the protein, while that in Mipa-P is partially exposed to the surface (Fig. S12-13 and S16). If the tryptophan residue in Mipa-P was replaced with phenylalanine, its activity and Tm values increased, but the reverse case in Kubu-P showed the decreased values of both (Fig. S14-15). Another key residue corresponds to H194 and D190 of Mipa-P and Kubu-P, respectively (Fig. S12-S13). Considering that many efficient variants of IsPETase (*27*, *29*, *30*, *31*) includes the D-to-H replacement at the site, which is originated from the IsPETase-EHA variant (*27*), it was predicted that variation at this site would result in significant differences in activity for the two enzymes. As expected, the Kubu-P$^{D190H}$ variants showed activity increase of 35% in 1d and 22% in 3d, respectively (Fig. S14-15), showing that the site is a strong engineering target for many PETase. Some examples of the key residue variation have convinced us that they still have not evolved to an optimal structure for *in vitro* depolymerization and that there is large room for introducing artificial mutations from these templates.

**Figs. S1 to S26**

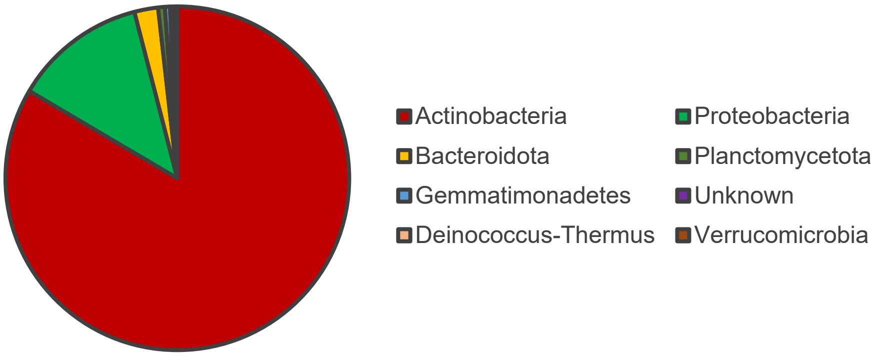


**Fig. S1. Pie chart of phyla of the 2,064 sequences in the library.** Percentages from major phylum data for each sequence. See Data S3 for the details. Each of the 2,064 amino acid sequences are originated from nucleotide sequences of natural species, and the major genetic resource is analyzed for the major phylum data.

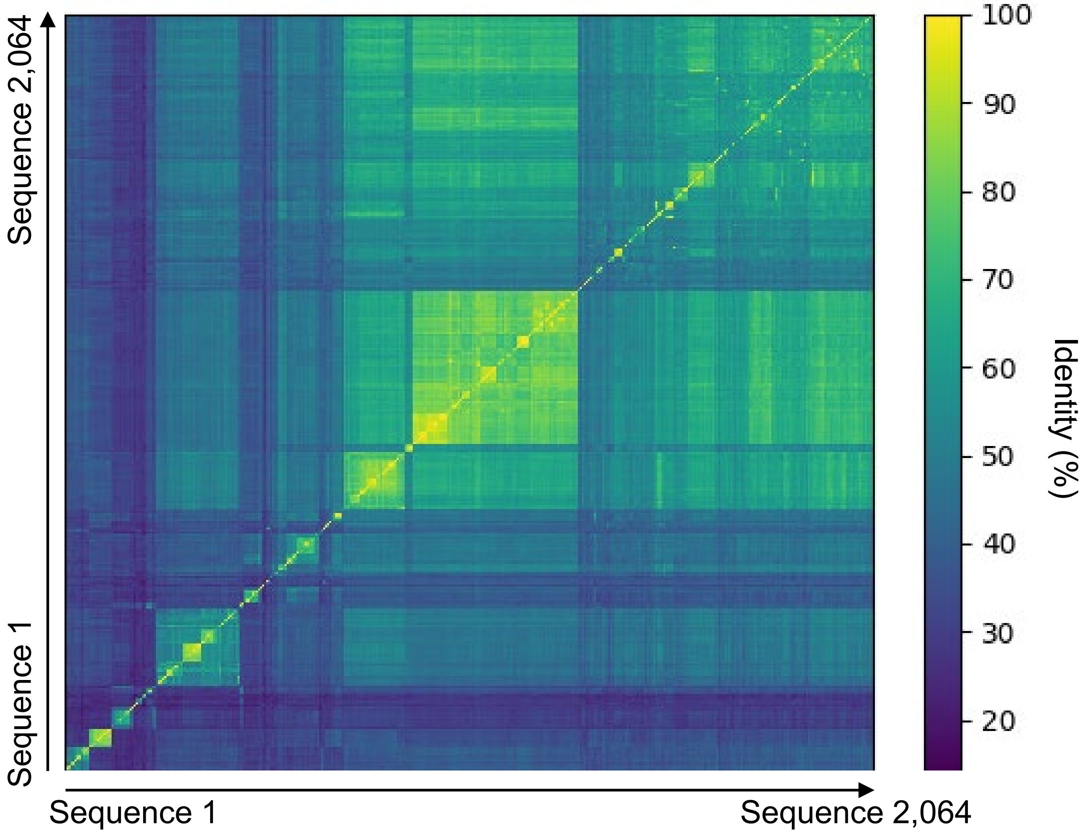


**Fig. S2. Heatmap of percent identity matrix of the 2,064 sequences in the library.** See Data S4 for the sequence (accession code) order and details.

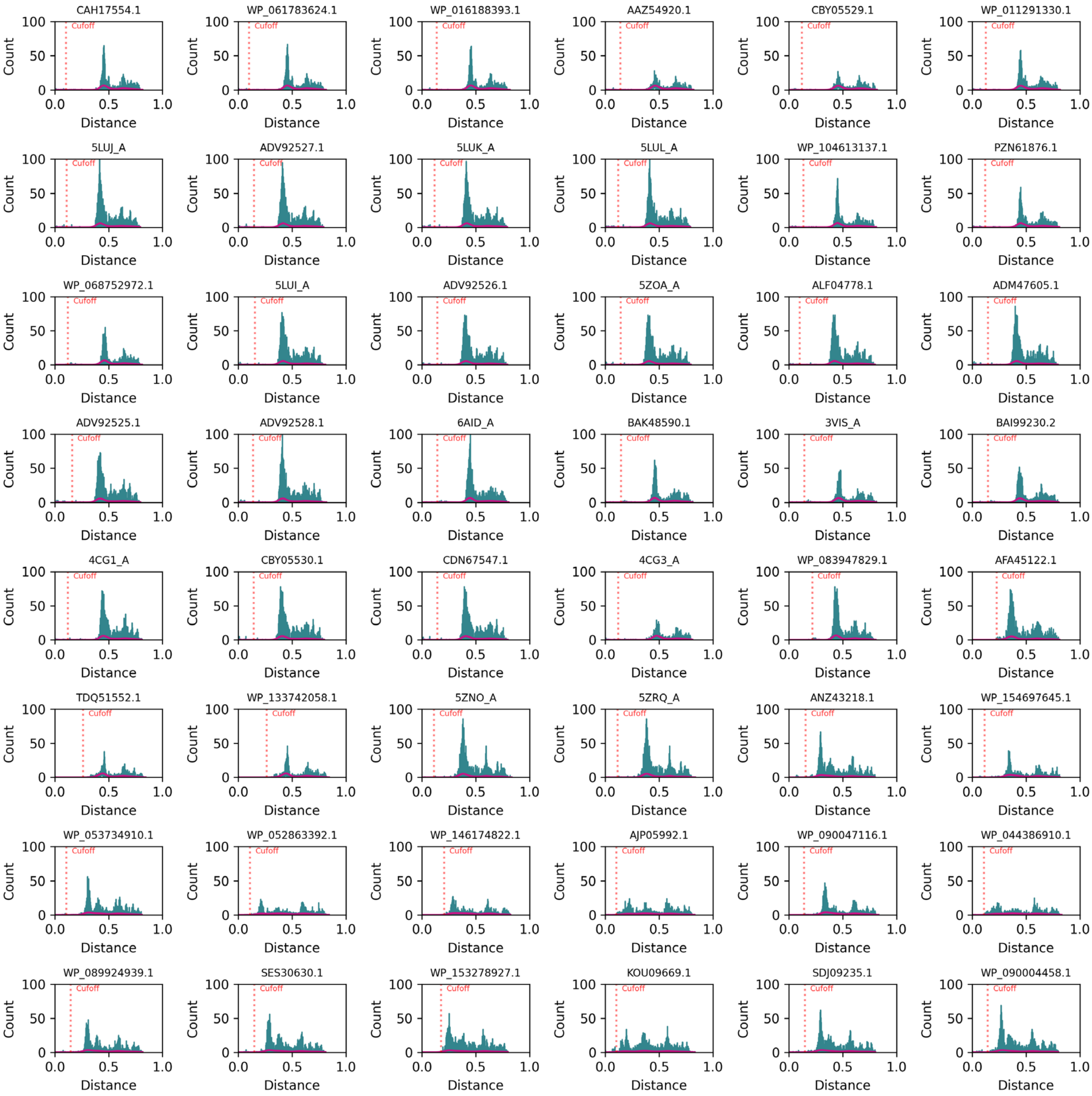


**Fig. S3. Histograms for the 2,064 sequences (continued, 1/43).** Cut-off distance (the first neighbor that is not linked to this sequence with a distance edge) for each sequence is labeled and indicated with a red vertical dashed line. The histograms are shown in order of library id value.

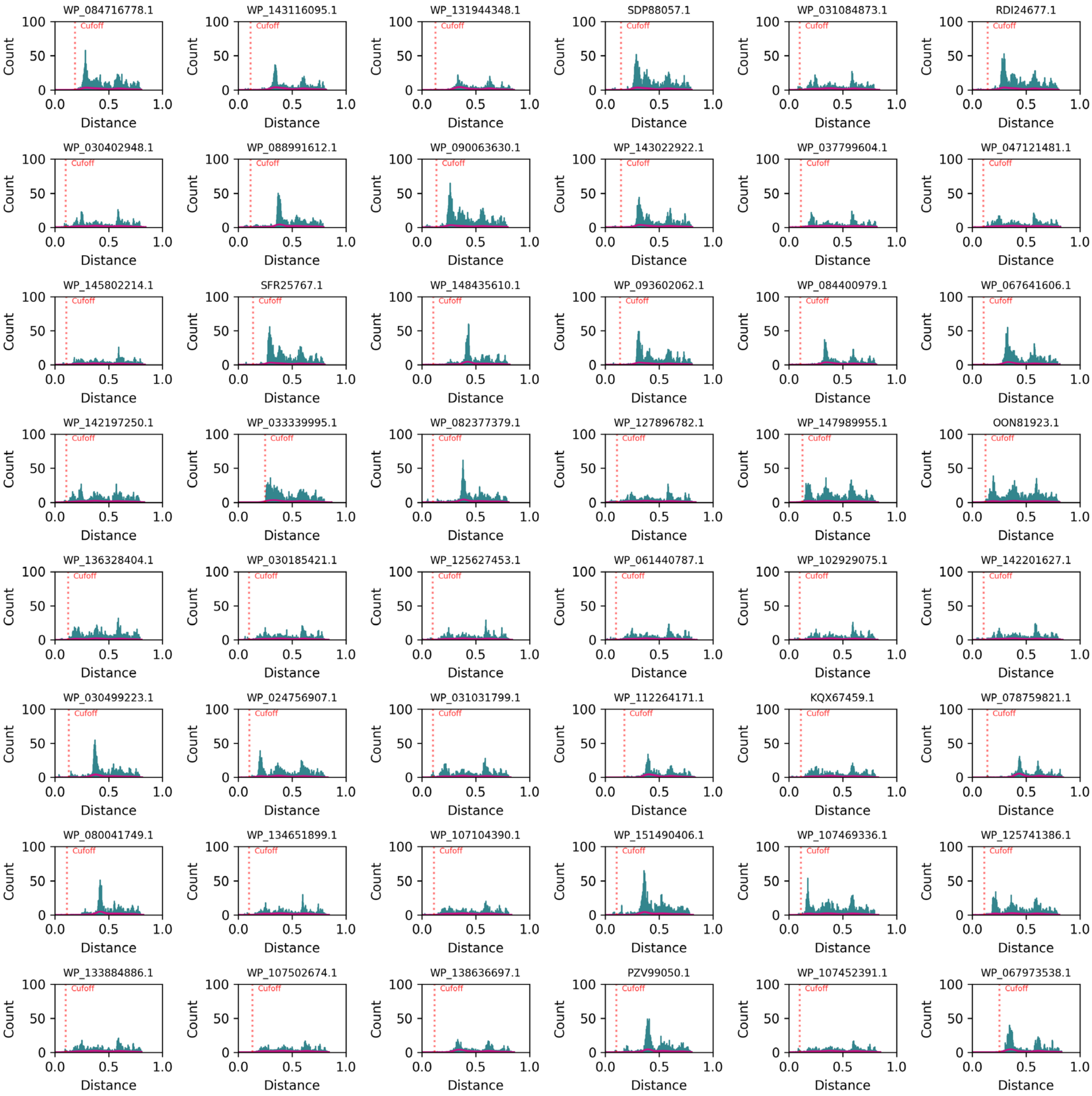


**Fig. S3. Histograms for the 2,064 sequences (continued, 2/43).** Cut-off distance (the first neighbor that is not linked to this sequence with a distance edge) for each sequence is labeled and indicated with a red vertical dashed line. The histograms are shown in order of library id value.

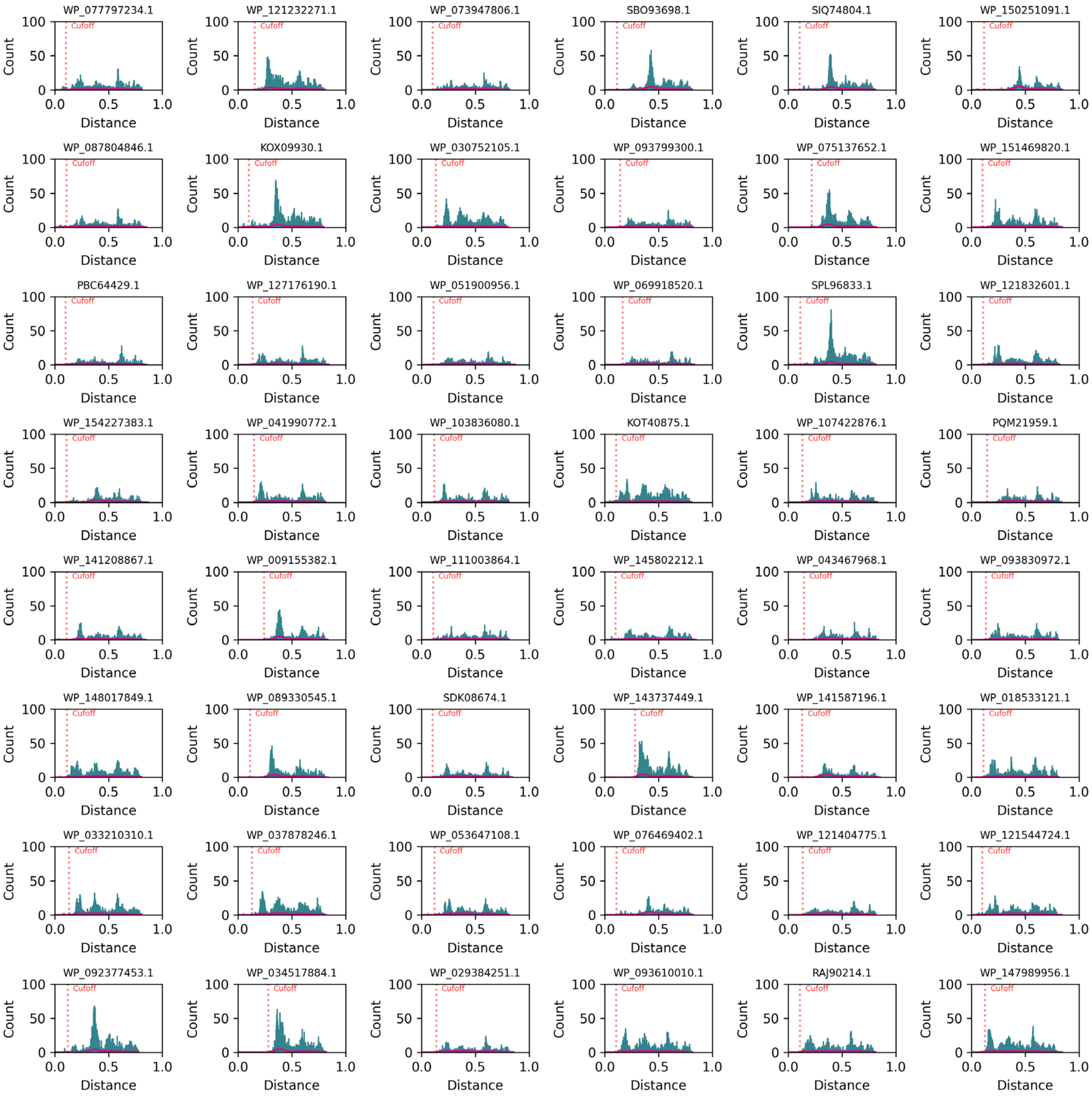


**Fig. S3. Histograms for the 2,064 sequences (continued, 3/43).** Cut-off distance (the first neighbor that is not linked to this sequence with a distance edge) for each sequence is labeled and indicated with a red vertical dashed line. The histograms are shown in order of library id value.

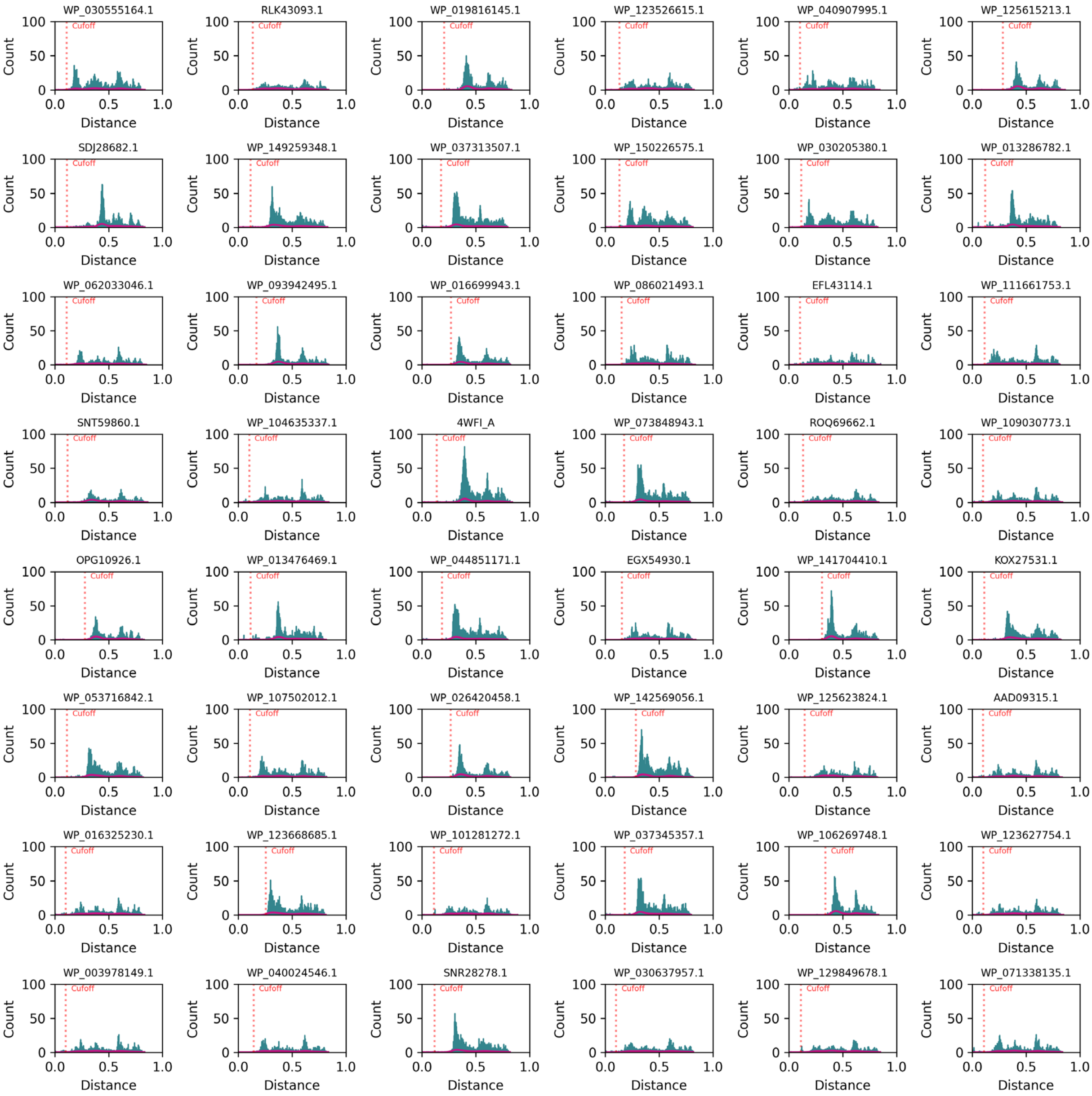


**Fig. S3. Histograms for the 2,064 sequences (continued, 4/43).** Cut-off distance (the first neighbor that is not linked to this sequence with a distance edge) for each sequence is labeled and indicated with a red vertical dashed line. The histograms are shown in order of library id value.

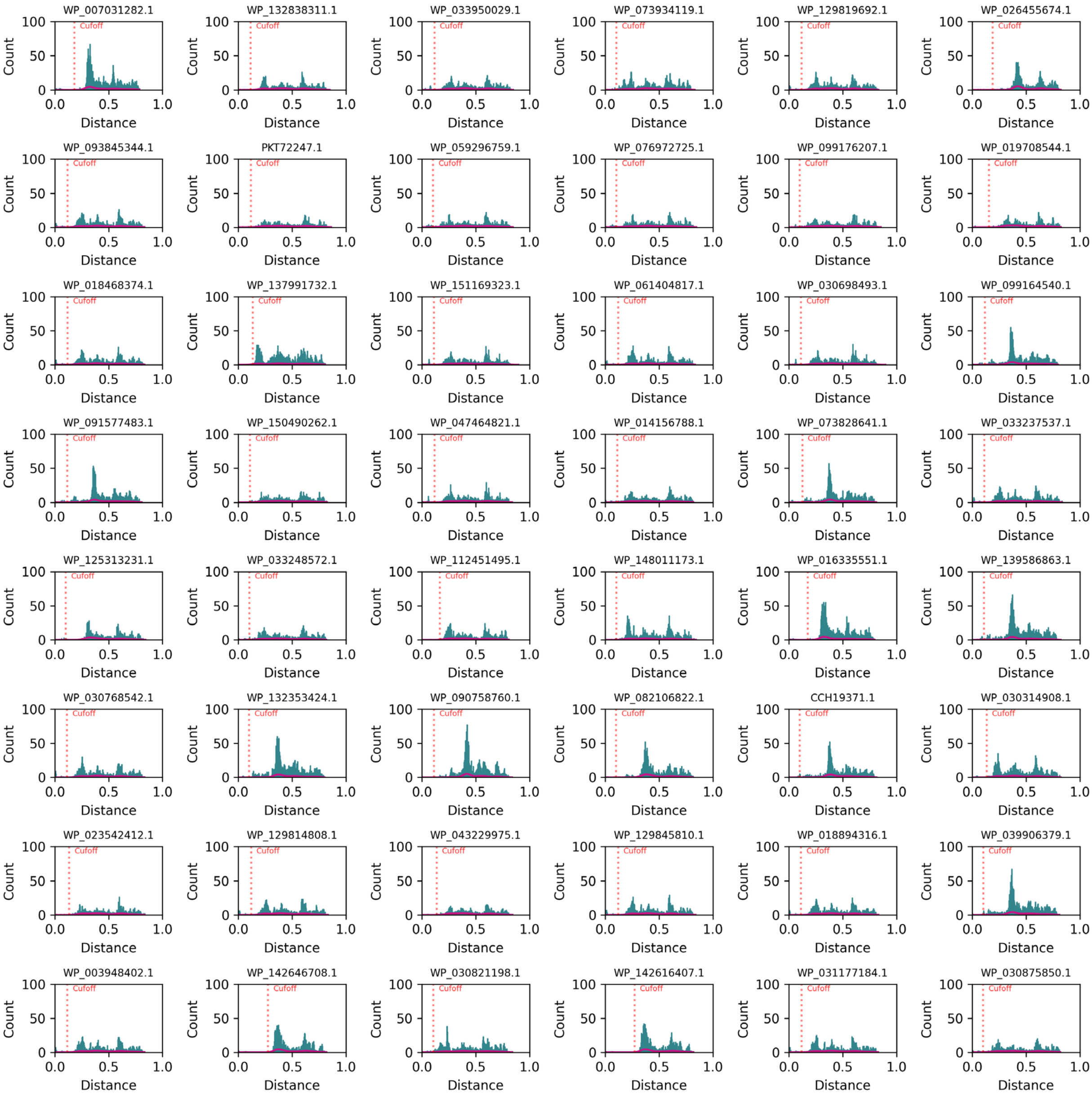


**Fig. S3. Histograms for the 2,064 sequences (continued, 5/43).** Cut-off distance (the first neighbor that is not linked to this sequence with a distance edge) for each sequence is labeled and indicated with a red vertical dashed line. The histograms are shown in order of library id value.

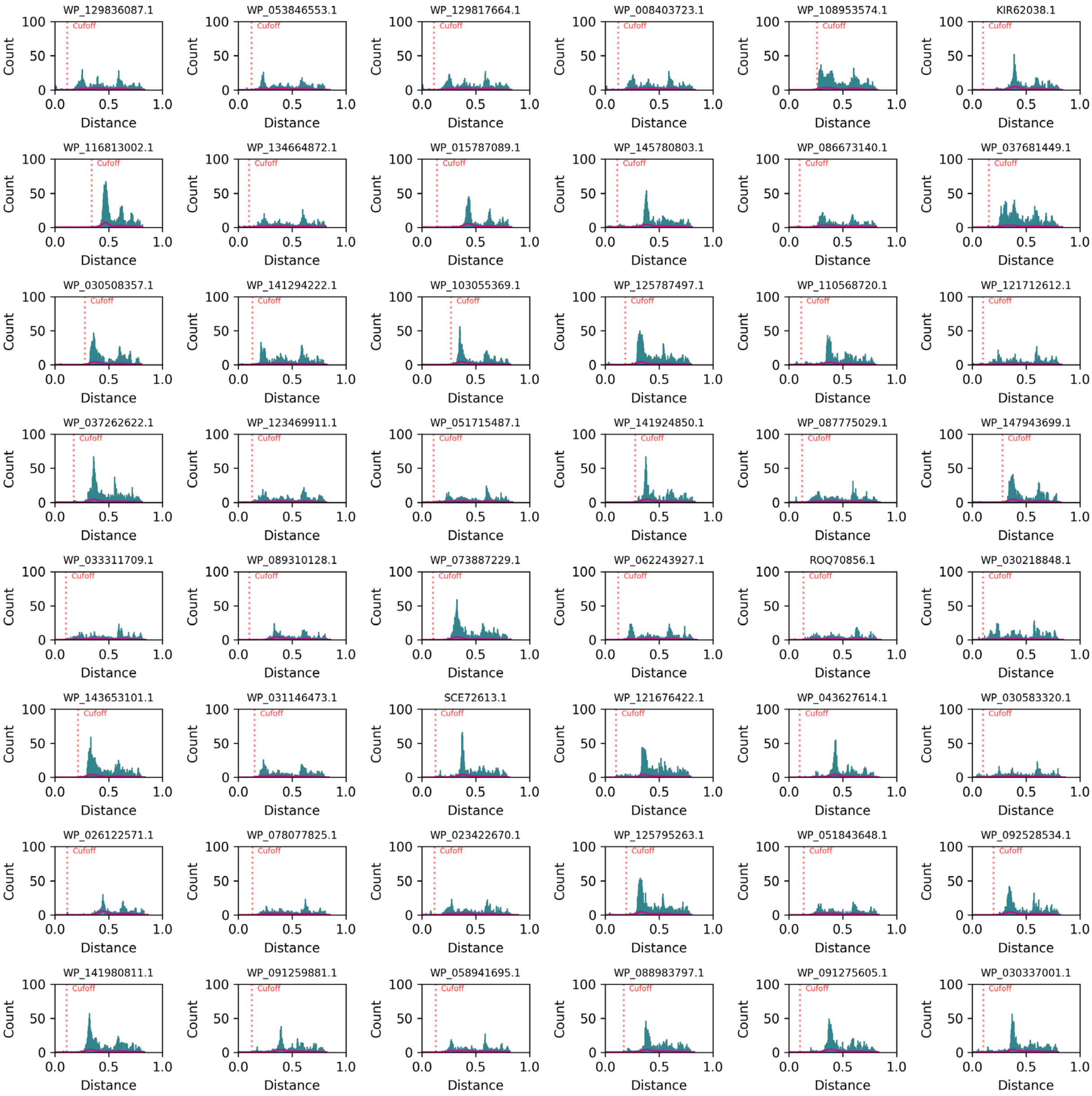


**Fig. S3. Histograms for the 2,064 sequences (continued, 6/43).** Cut-off distance (the first neighbor that is not linked to this sequence with a distance edge) for each sequence is labeled and indicated with a red vertical dashed line. The histograms are shown in order of library id value.

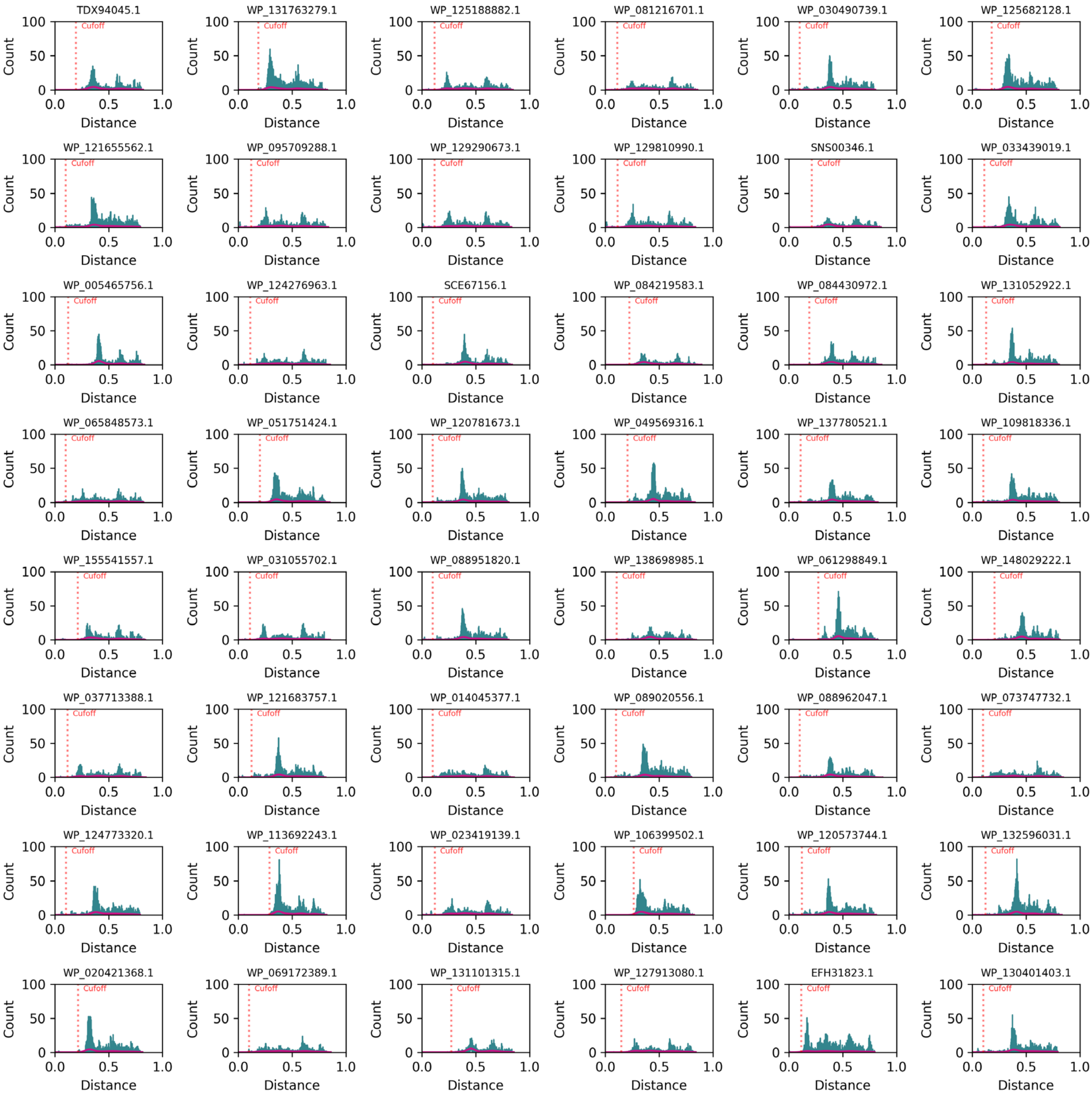


**Fig. S3. Histograms for the 2,064 sequences (continued, 7/43).** Cut-off distance (the first neighbor that is not linked to this sequence with a distance edge) for each sequence is labeled and indicated with a red vertical dashed line. The histograms are shown in order of library id value.

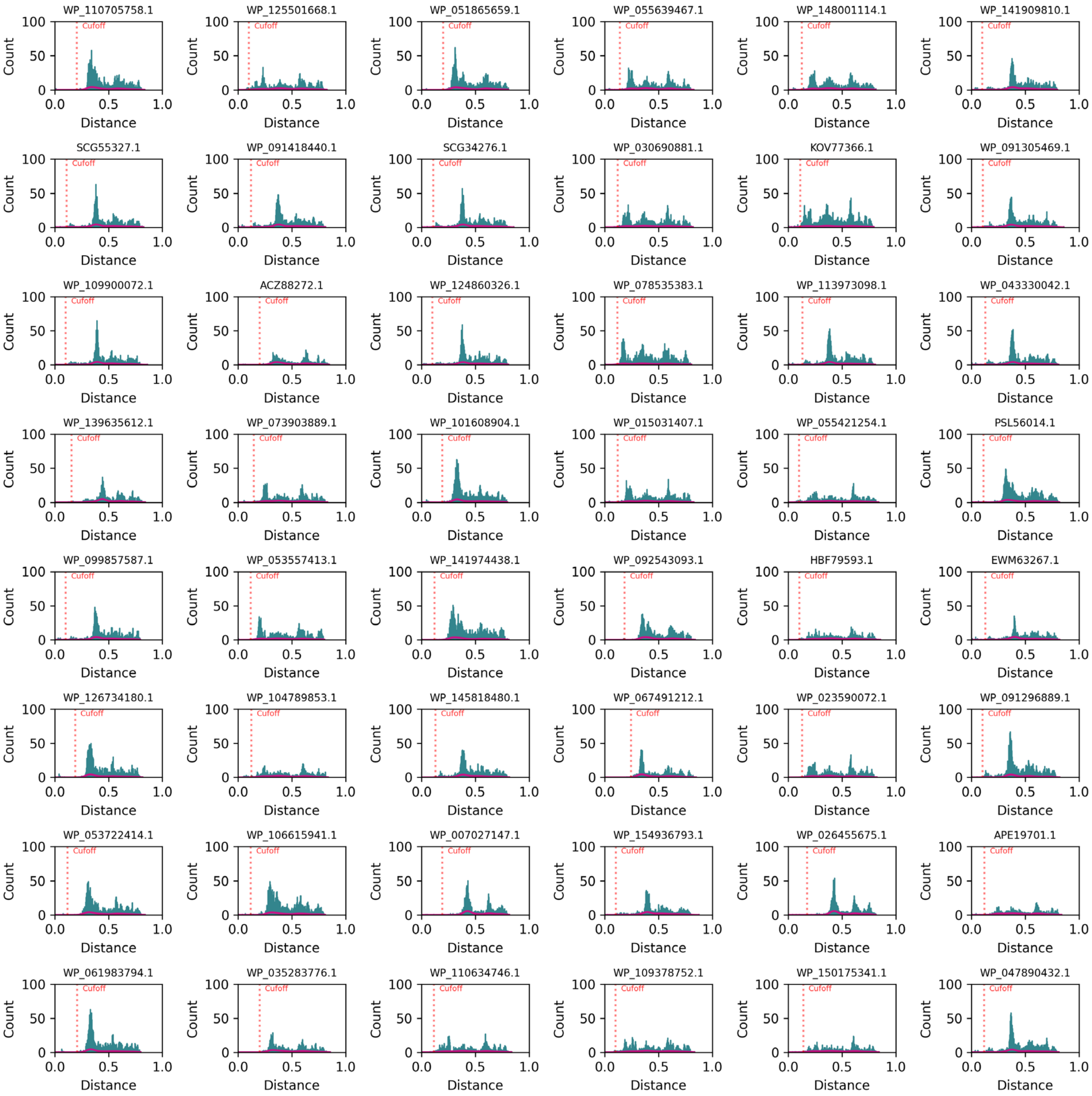


**Fig. S3. Histograms for the 2,064 sequences (continued, 8/43).** Cut-off distance (the first neighbor that is not linked to this sequence with a distance edge) for each sequence is labeled and indicated with a red vertical dashed line. The histograms are shown in order of library id value.

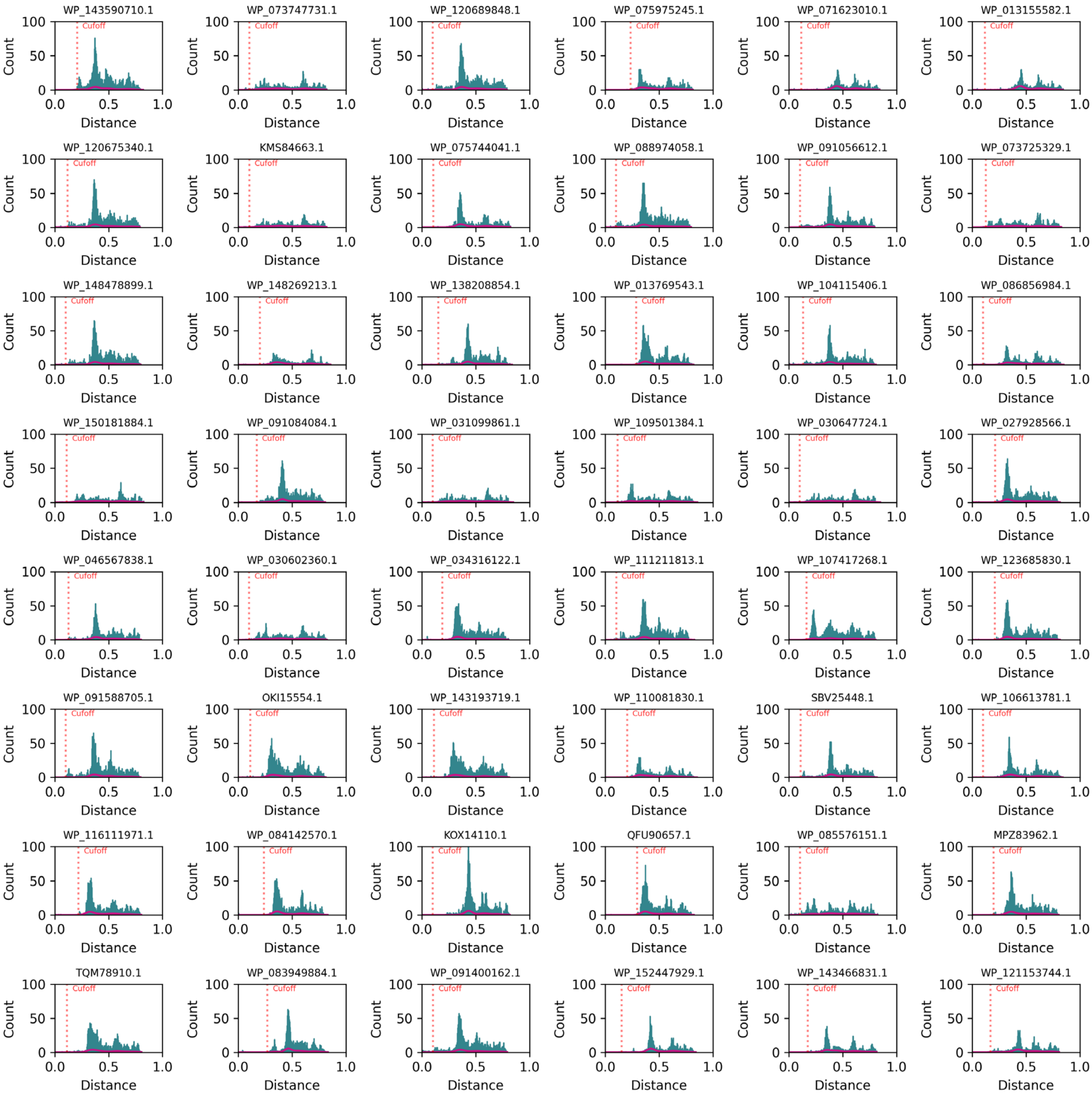


**Fig. S3. Histograms for the 2,064 sequences (continued, 9/43).** Cut-off distance (the first neighbor that is not linked to this sequence with a distance edge) for each sequence is labeled and indicated with a red vertical dashed line. The histograms are shown in order of library id value.

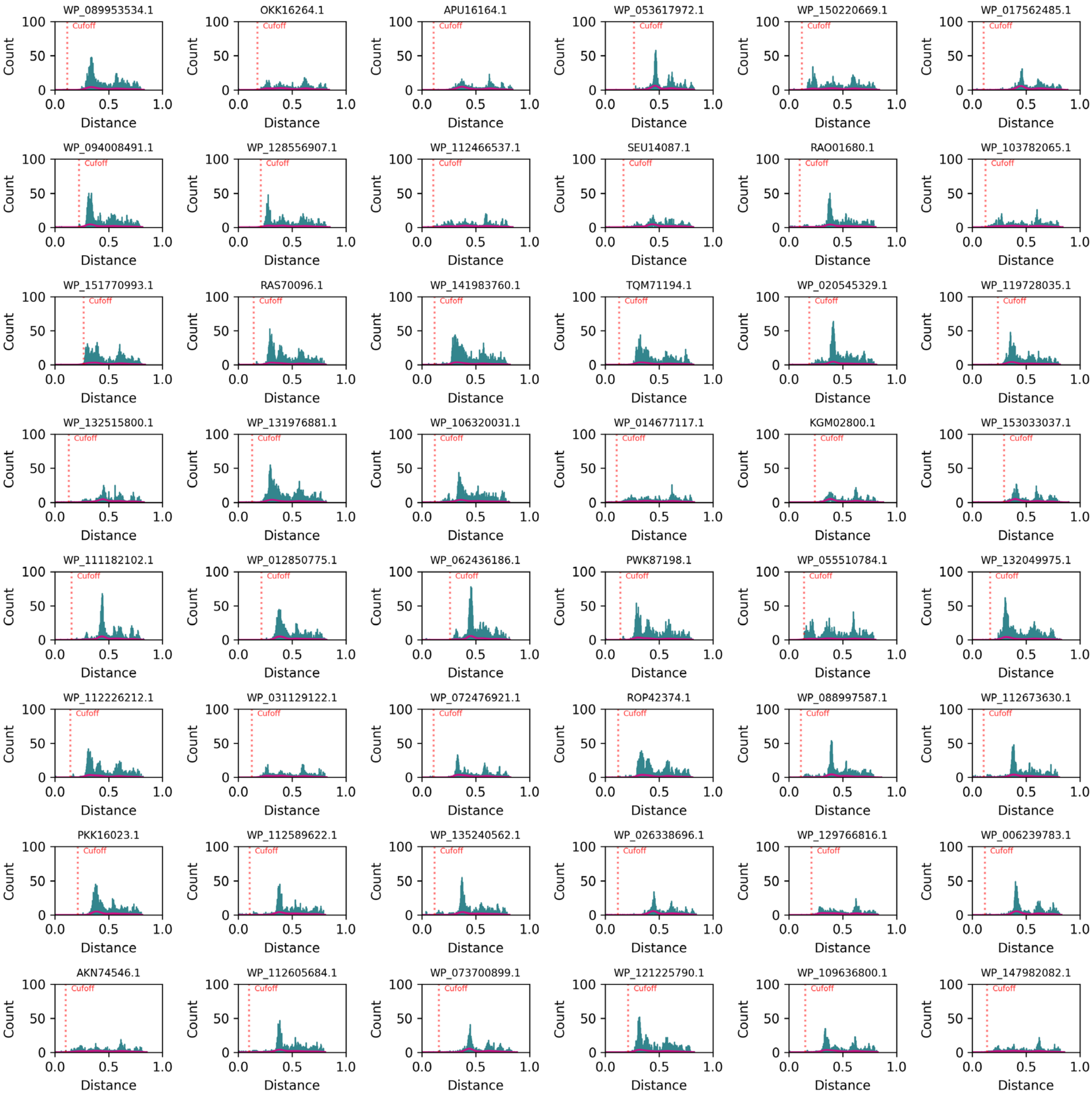


**Fig. S3. Histograms for the 2,064 sequences (continued, 10/43).** Cut-off distance (the first neighbor that is not linked to this sequence with a distance edge) for each sequence is labeled and indicated with a red vertical dashed line. The histograms are shown in order of library id value.

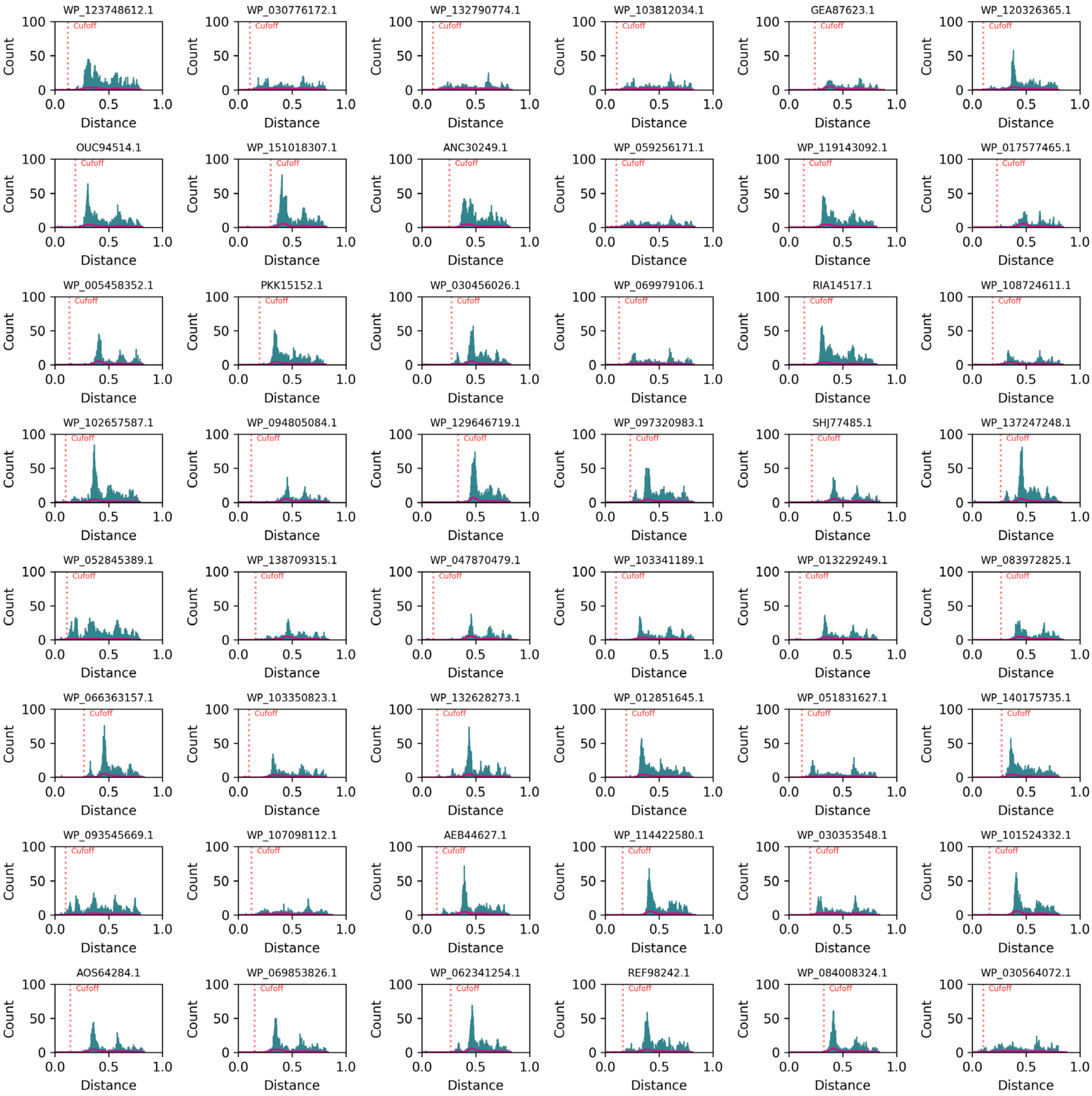


**Fig. S3. Histograms for the 2,064 sequences (continued, 11/43).** Cut-off distance (the first neighbor that is not linked to this sequence with a distance edge) for each sequence is labeled and indicated with a red vertical dashed line. The histograms are shown in order of library id value.

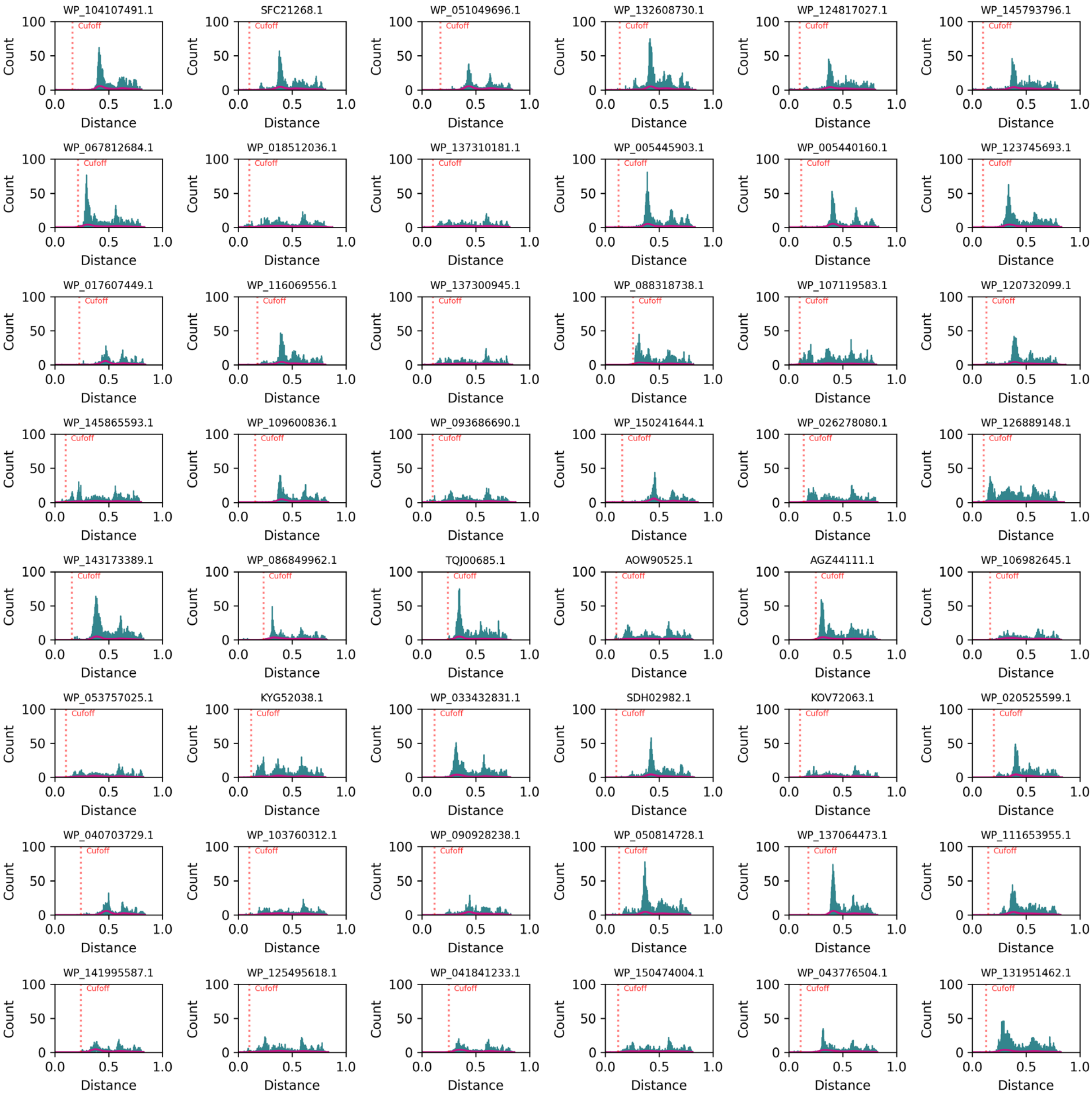


**Fig. S3. Histograms for the 2,064 sequences (continued, 12/43).** Cut-off distance (the first neighbor that is not linked to this sequence with a distance edge) for each sequence is labeled and indicated with a red vertical dashed line. The histograms are shown in order of library id value.

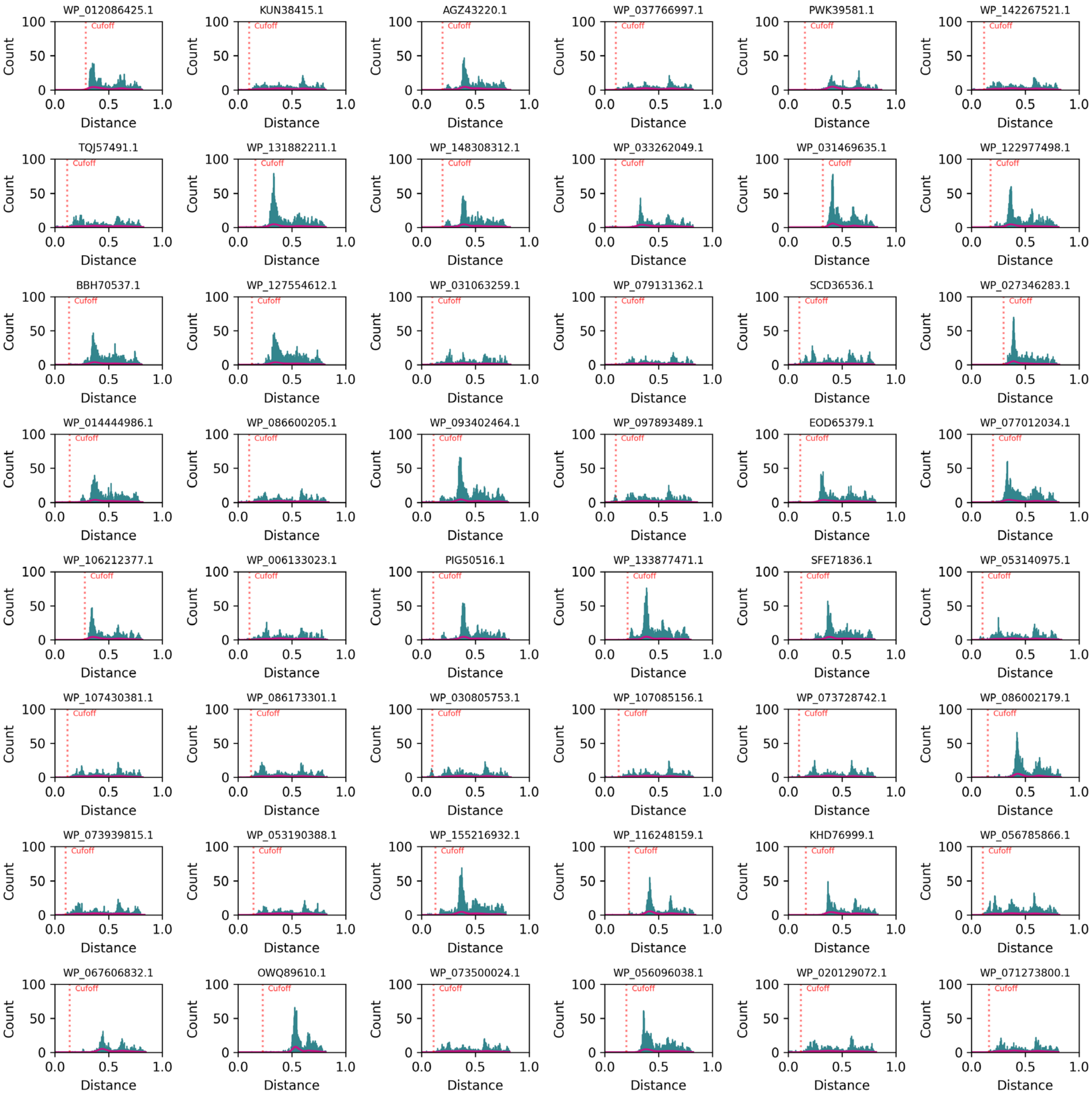


**Fig. S3. Histograms for the 2,064 sequences (continued, 13/43).** Cut-off distance (the first neighbor that is not linked to this sequence with a distance edge) for each sequence is labeled and indicated with a red vertical dashed line. The histograms are shown in order of library id value.

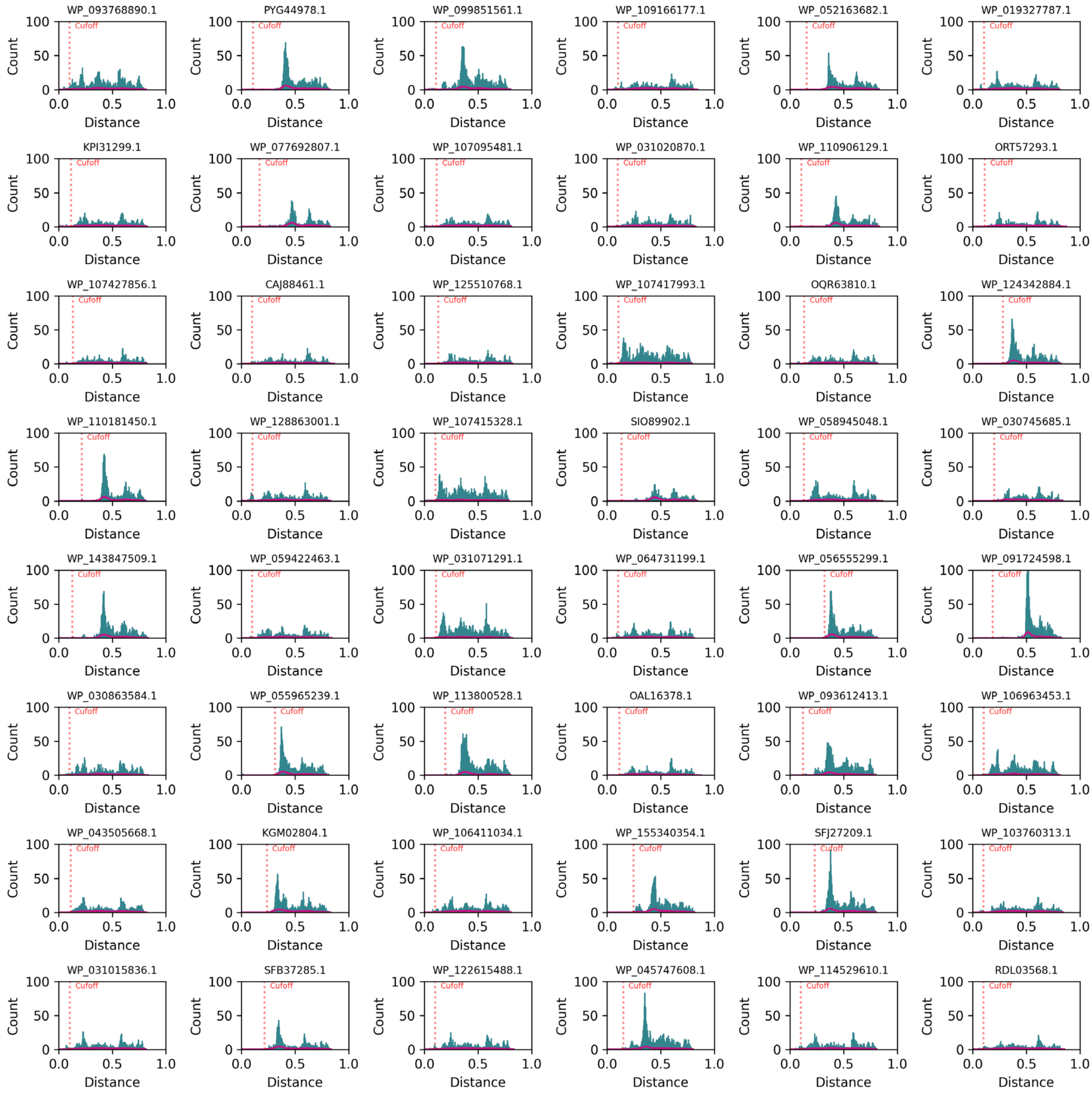


**Fig. S3. Histograms for the 2,064 sequences (continued, 14/43).** Cut-off distance (the first neighbor that is not linked to this sequence with a distance edge) for each sequence is labeled and indicated with a red vertical dashed line. The histograms are shown in order of library id value.

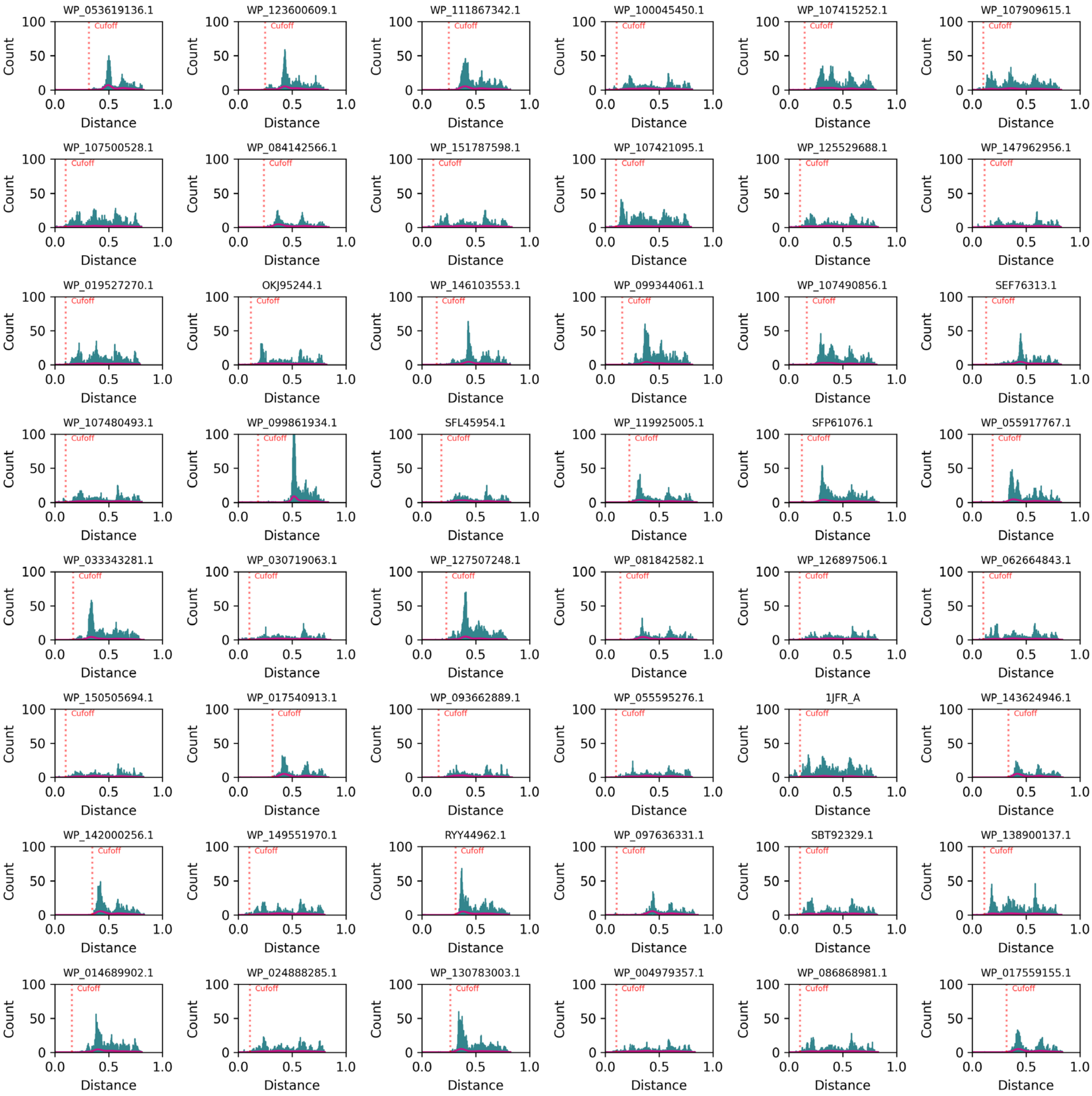


**Fig. S3. Histograms for the 2,064 sequences (continued, 15/43).** Cut-off distance (the first neighbor that is not linked to this sequence with a distance edge) for each sequence is labeled and indicated with a red vertical dashed line. The histograms are shown in order of library id value.

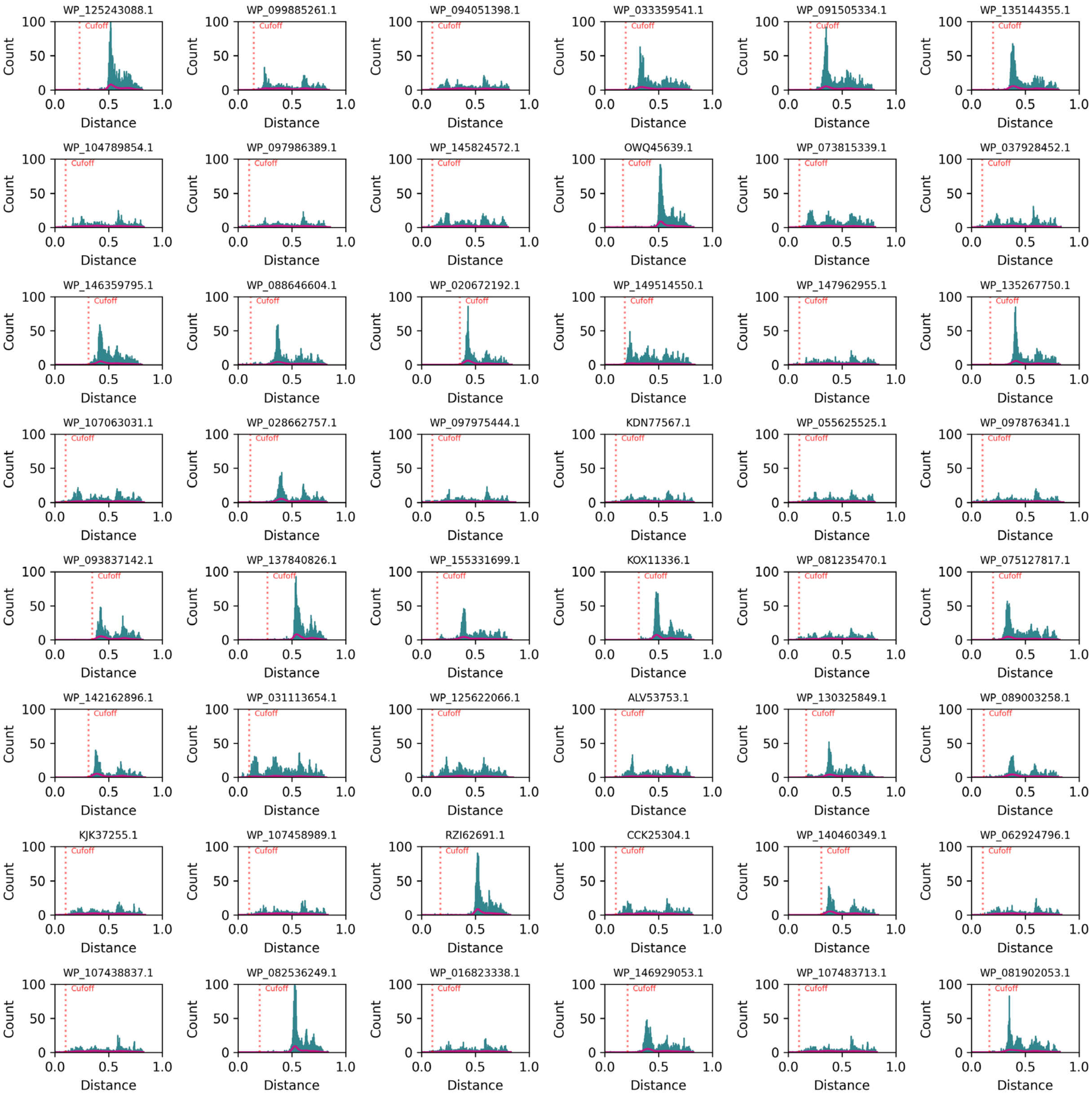


**Fig. S3. Histograms for the 2,064 sequences (continued, 16/43).** Cut-off distance (the first neighbor that is not linked to this sequence with a distance edge) for each sequence is labeled and indicated with a red vertical dashed line. The histograms are shown in order of library id value.

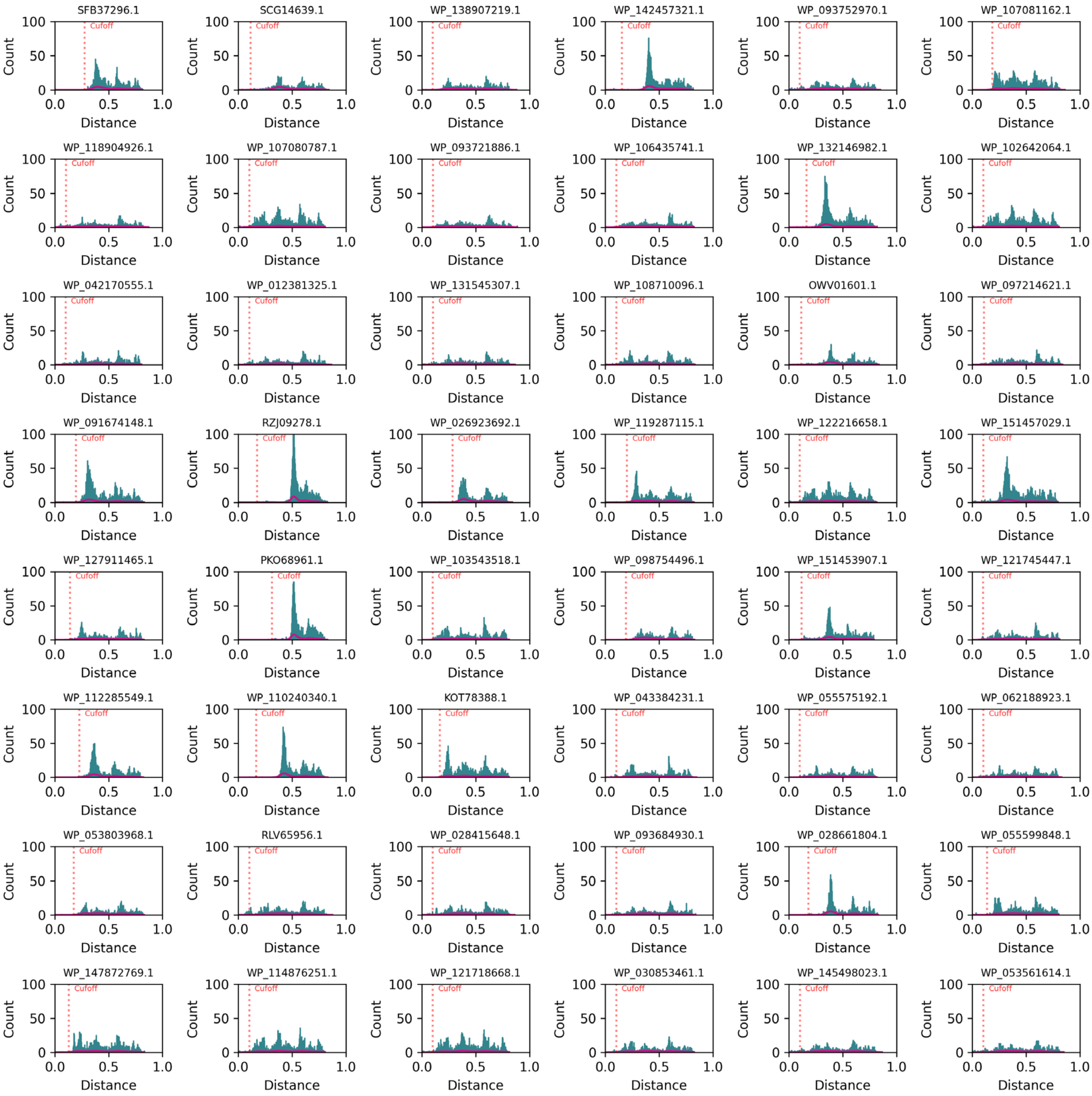


**Fig. S3. Histograms for the 2,064 sequences (continued, 17/43).** Cut-off distance (the first neighbor that is not linked to this sequence with a distance edge) for each sequence is labeled and indicated with a red vertical dashed line. The histograms are shown in order of library id value.

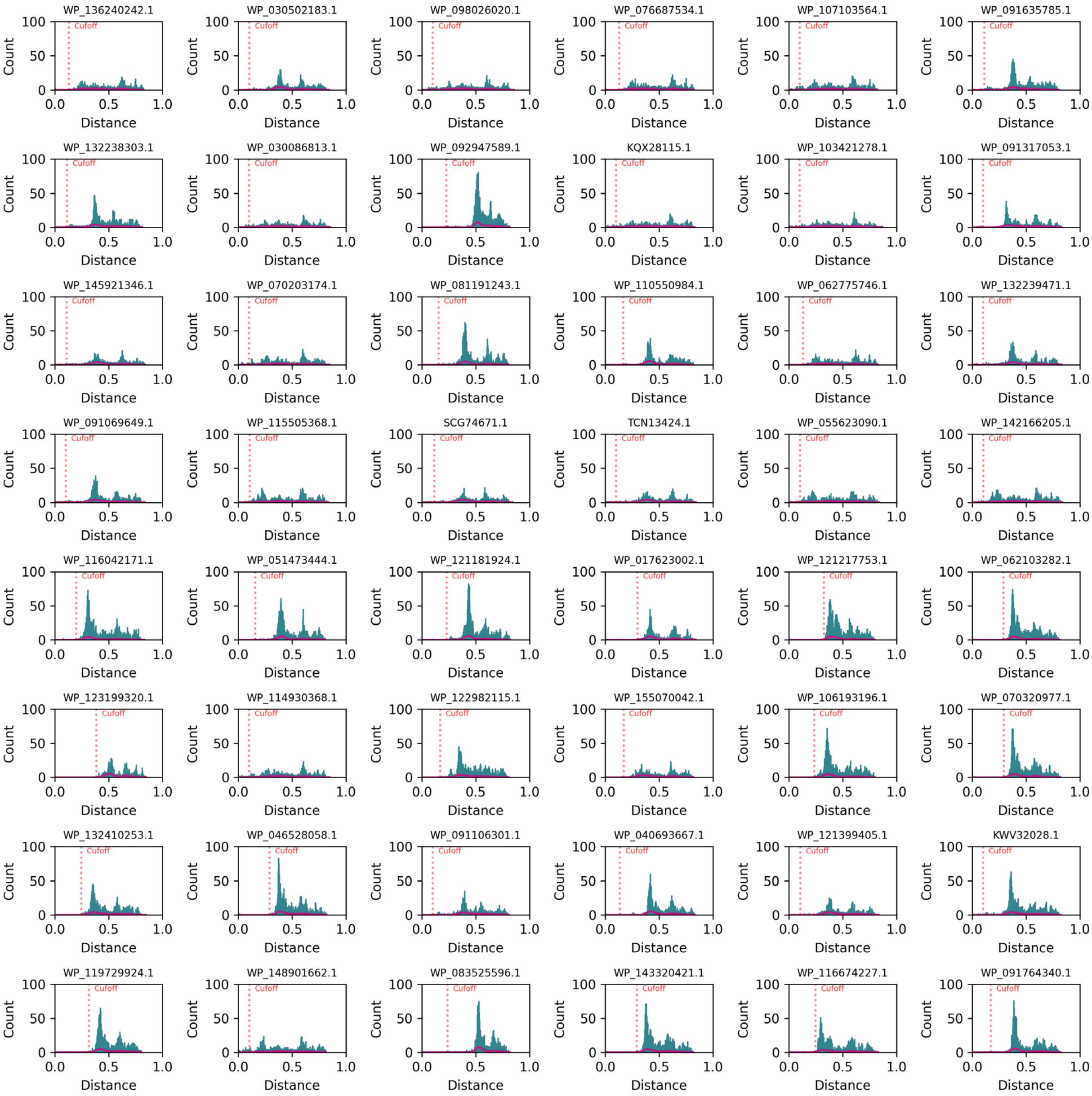


**Fig. S3. Histograms for the 2,064 sequences (continued, 18/43).** Cut-off distance (the first neighbor that is not linked to this sequence with a distance edge) for each sequence is labeled and indicated with a red vertical dashed line. The histograms are shown in order of library id value.

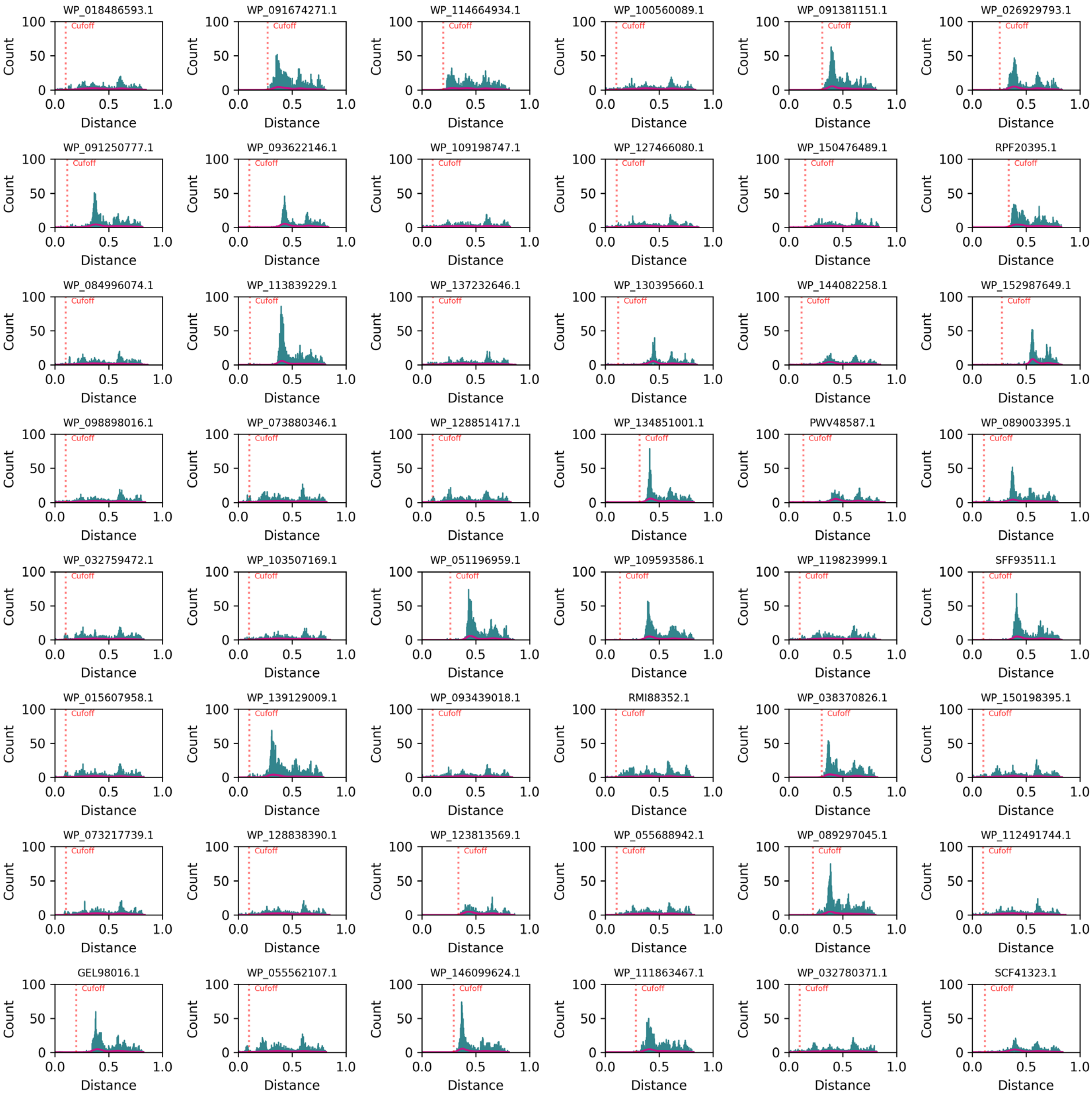


**Fig. S3. Histograms for the 2,064 sequences (continued, 19/43).** Cut-off distance (the first neighbor that is not linked to this sequence with a distance edge) for each sequence is labeled and indicated with a red vertical dashed line. The histograms are shown in order of library id value.

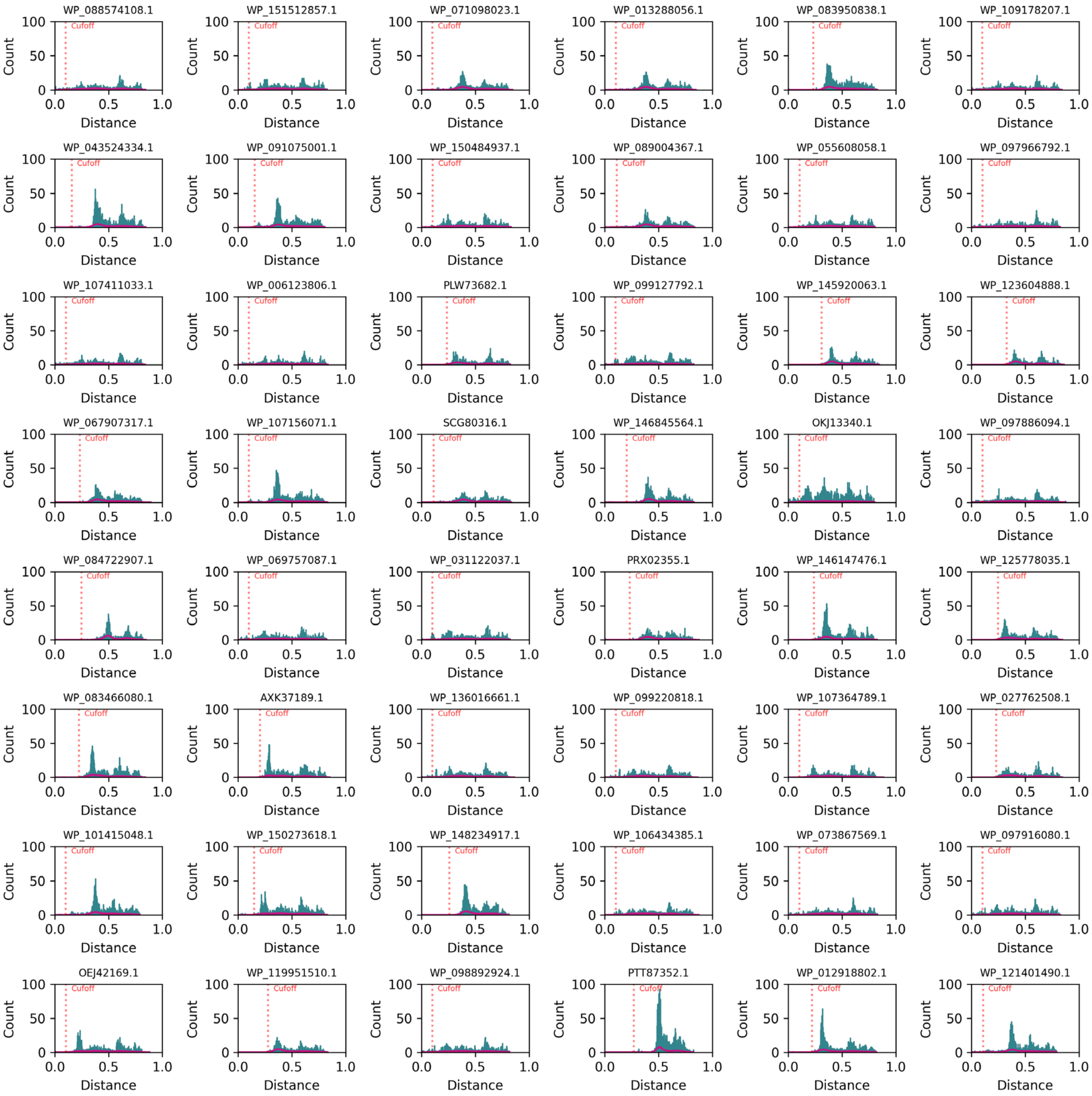


**Fig. S3. Histograms for the 2,064 sequences (continued, 20/43).** Cut-off distance (the first neighbor that is not linked to this sequence with a distance edge) for each sequence is labeled and indicated with a red vertical dashed line. The histograms are shown in order of library id value.

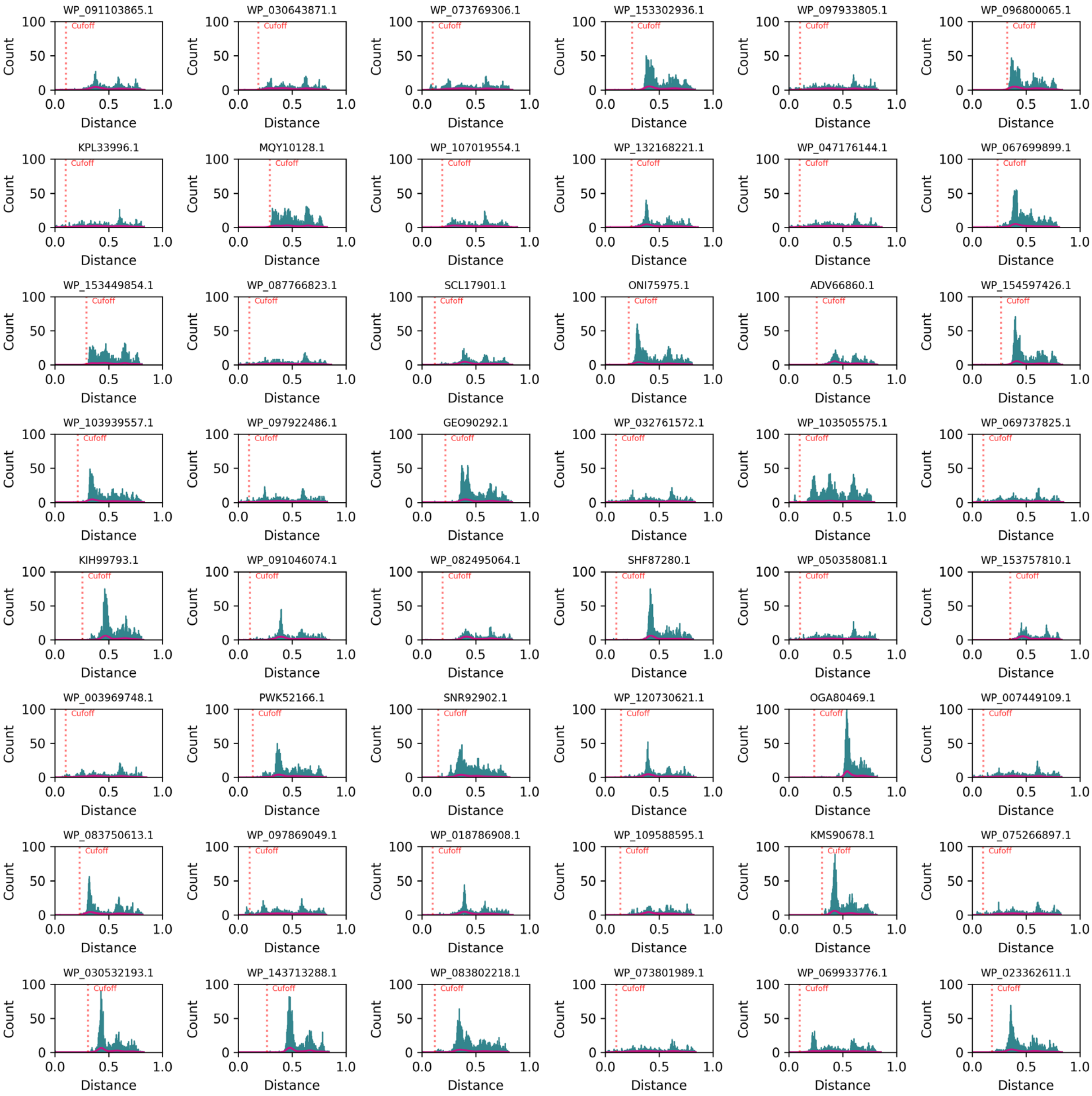


**Fig. S3. Histograms for the 2,064 sequences (continued, 21/43).** Cut-off distance (the first neighbor that is not linked to this sequence with a distance edge) for each sequence is labeled and indicated with a red vertical dashed line. The histograms are shown in order of library id value.

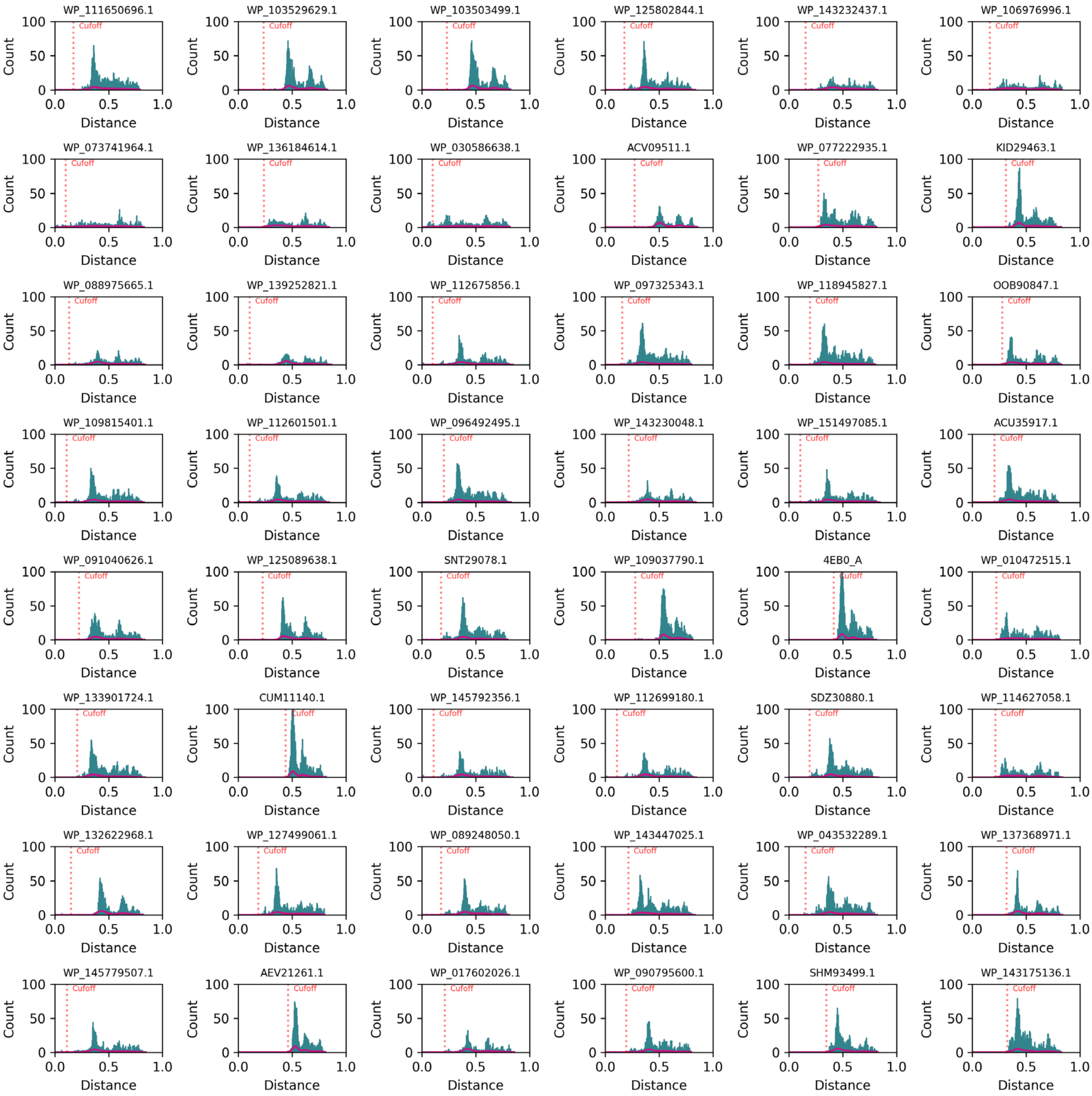


**Fig. S3. Histograms for the 2,064 sequences (continued, 22/43).** Cut-off distance (the first neighbor that is not linked to this sequence with a distance edge) for each sequence is labeled and indicated with a red vertical dashed line. The histograms are shown in order of library id value.

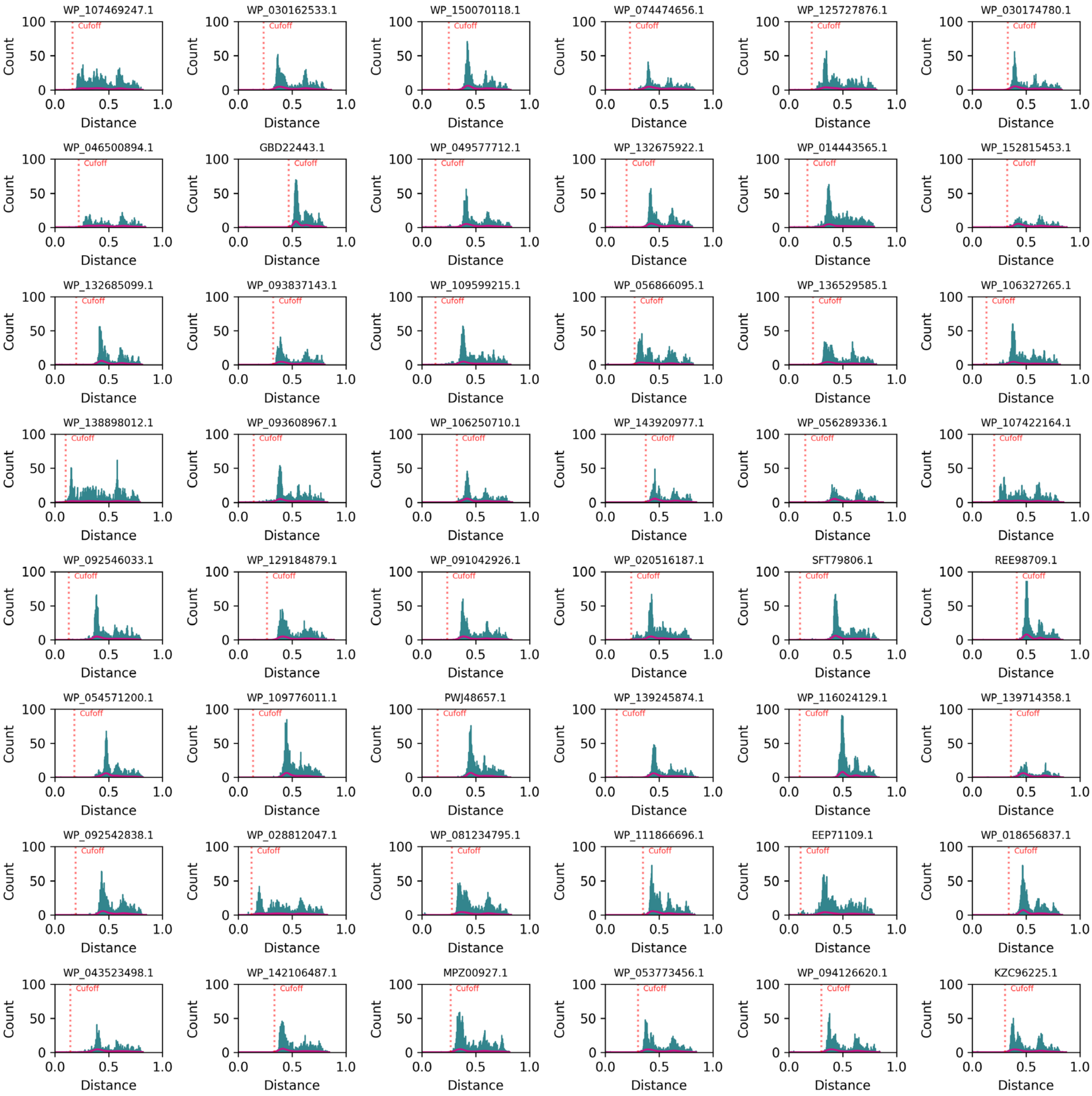


**Fig. S3. Histograms for the 2,064 sequences (continued, 23/43).** Cut-off distance (the first neighbor that is not linked to this sequence with a distance edge) for each sequence is labeled and indicated with a red vertical dashed line. The histograms are shown in order of library id value.

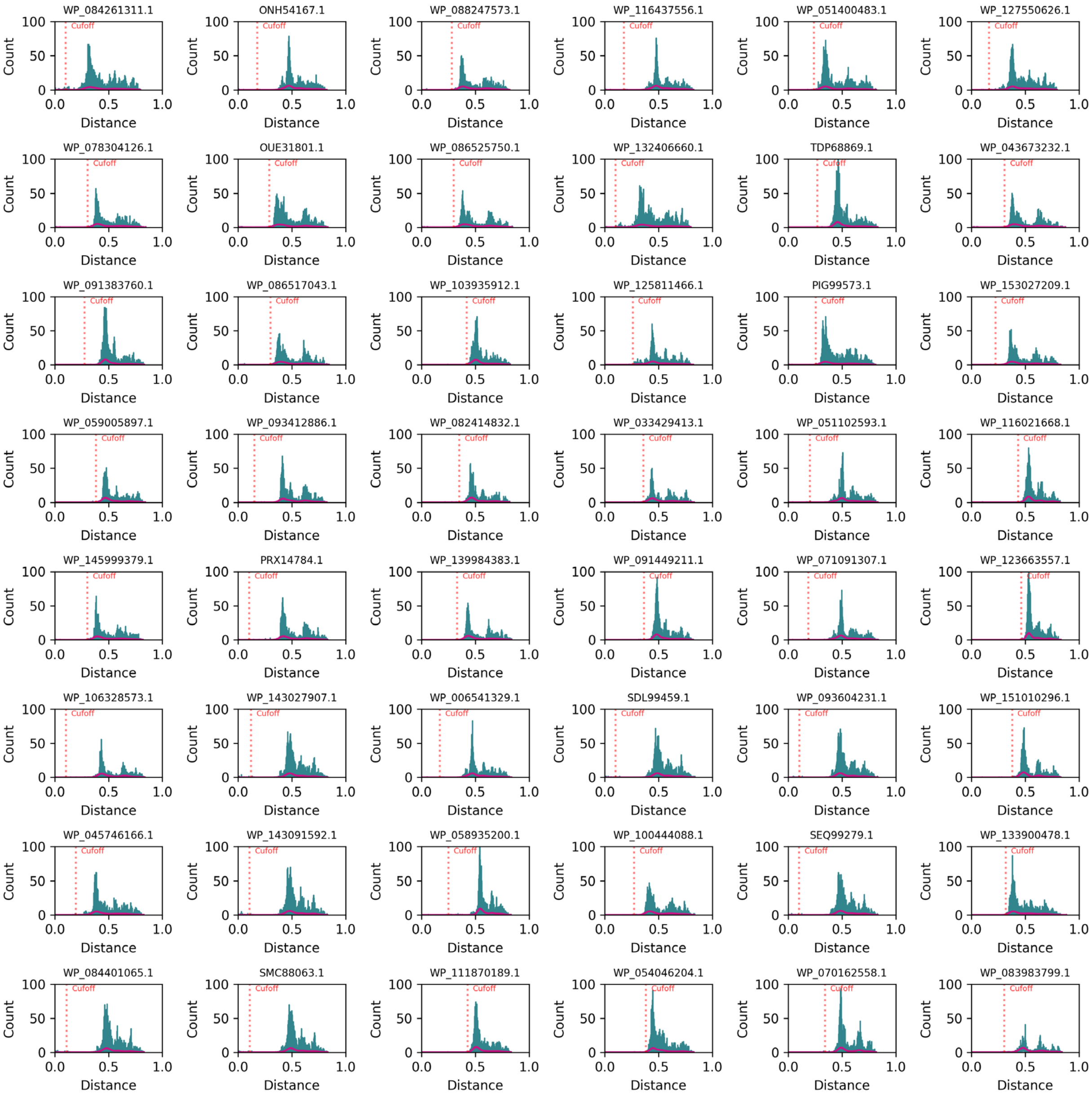


**Fig. S3. Histograms for the 2,064 sequences (continued, 24/43).** Cut-off distance (the first neighbor that is not linked to this sequence with a distance edge) for each sequence is labeled and indicated with a red vertical dashed line. The histograms are shown in order of library id value.

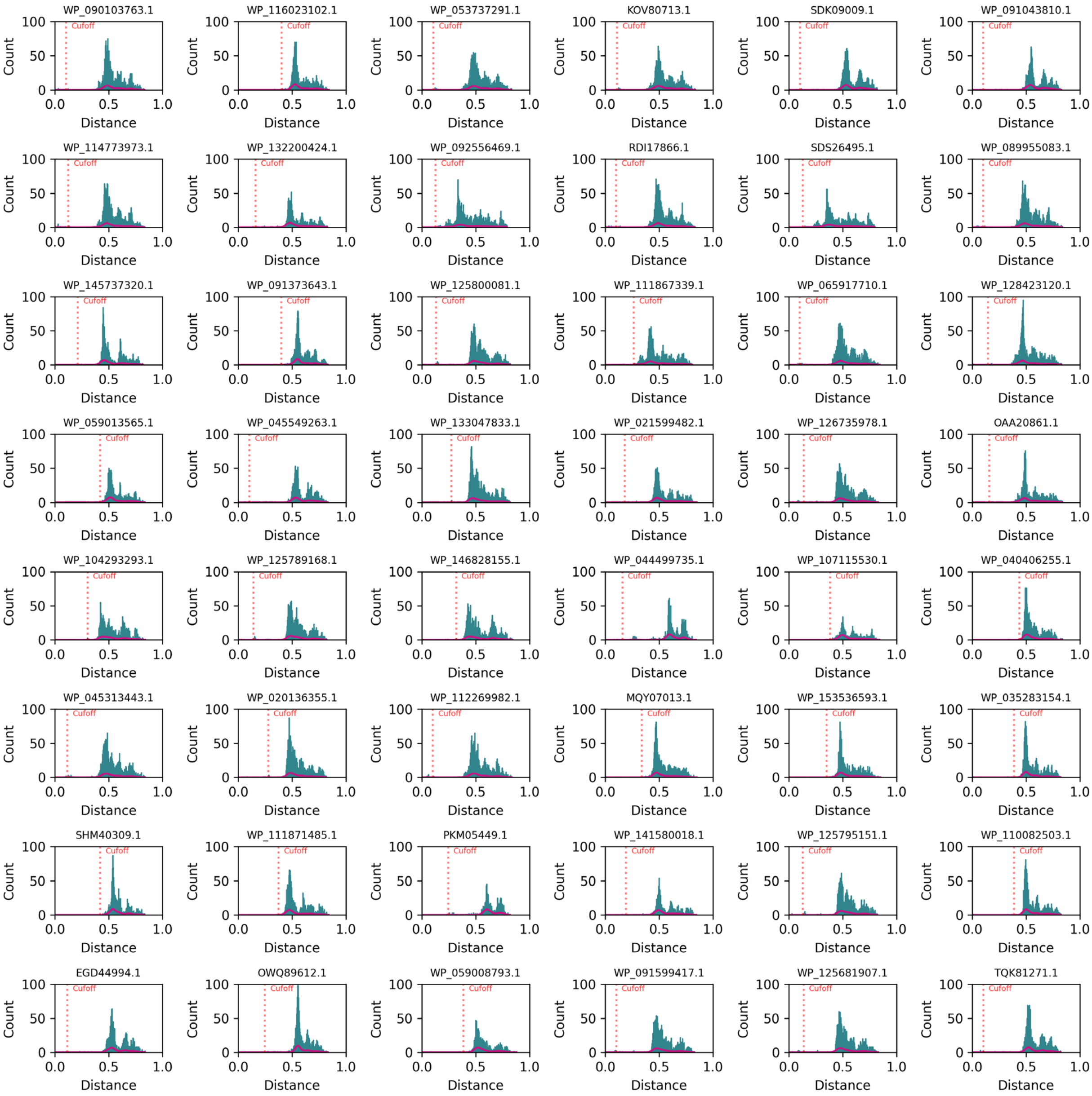


**Fig. S3. Histograms for the 2,064 sequences (continued, 25/43).** Cut-off distance (the first neighbor that is not linked to this sequence with a distance edge) for each sequence is labeled and indicated with a red vertical dashed line. The histograms are shown in order of library id value.

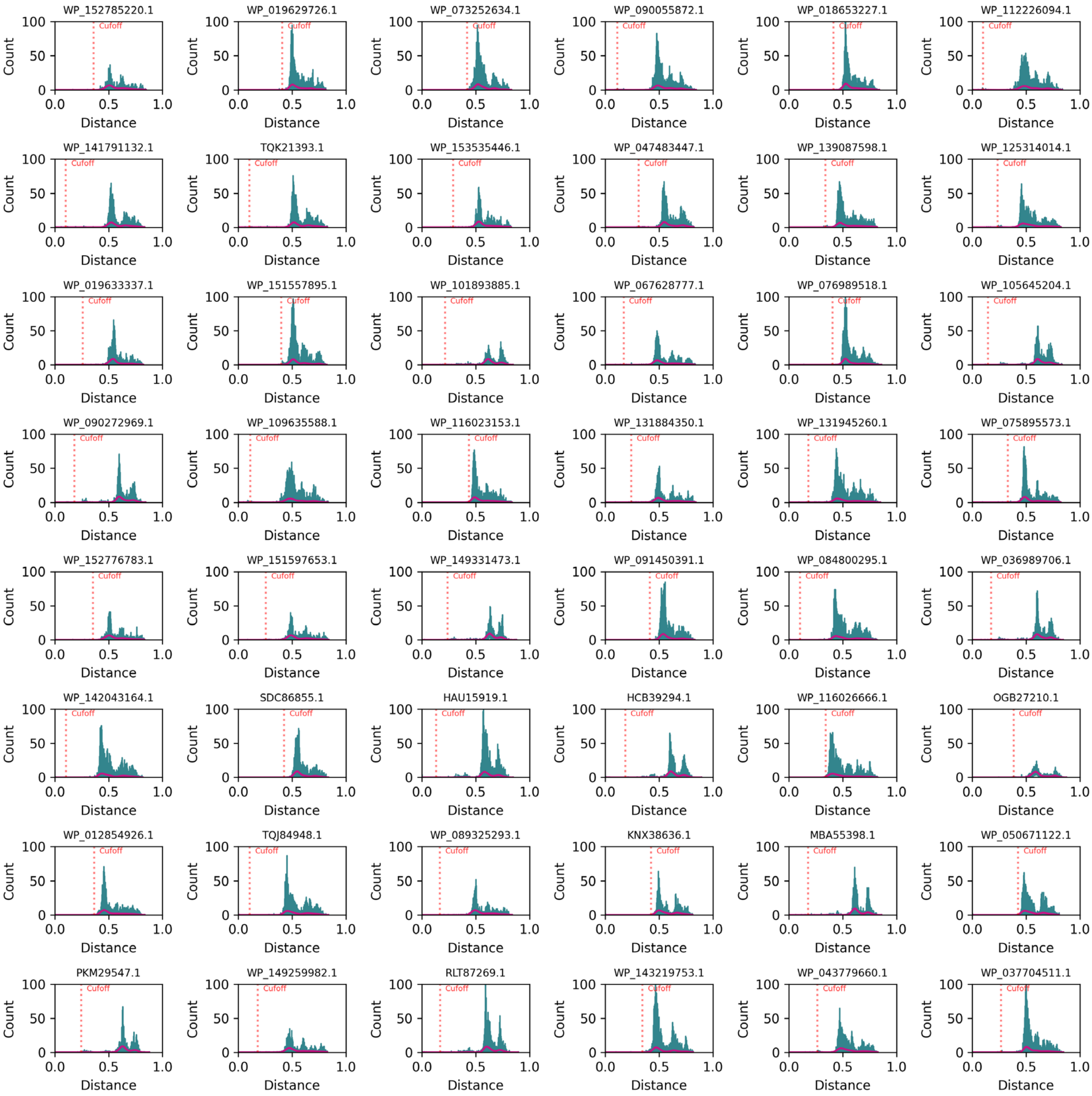


**Fig. S3. Histograms for the 2,064 sequences (continued, 26/43).** Cut-off distance (the first neighbor that is not linked to this sequence with a distance edge) for each sequence is labeled and indicated with a red vertical dashed line. The histograms are shown in order of library id value.

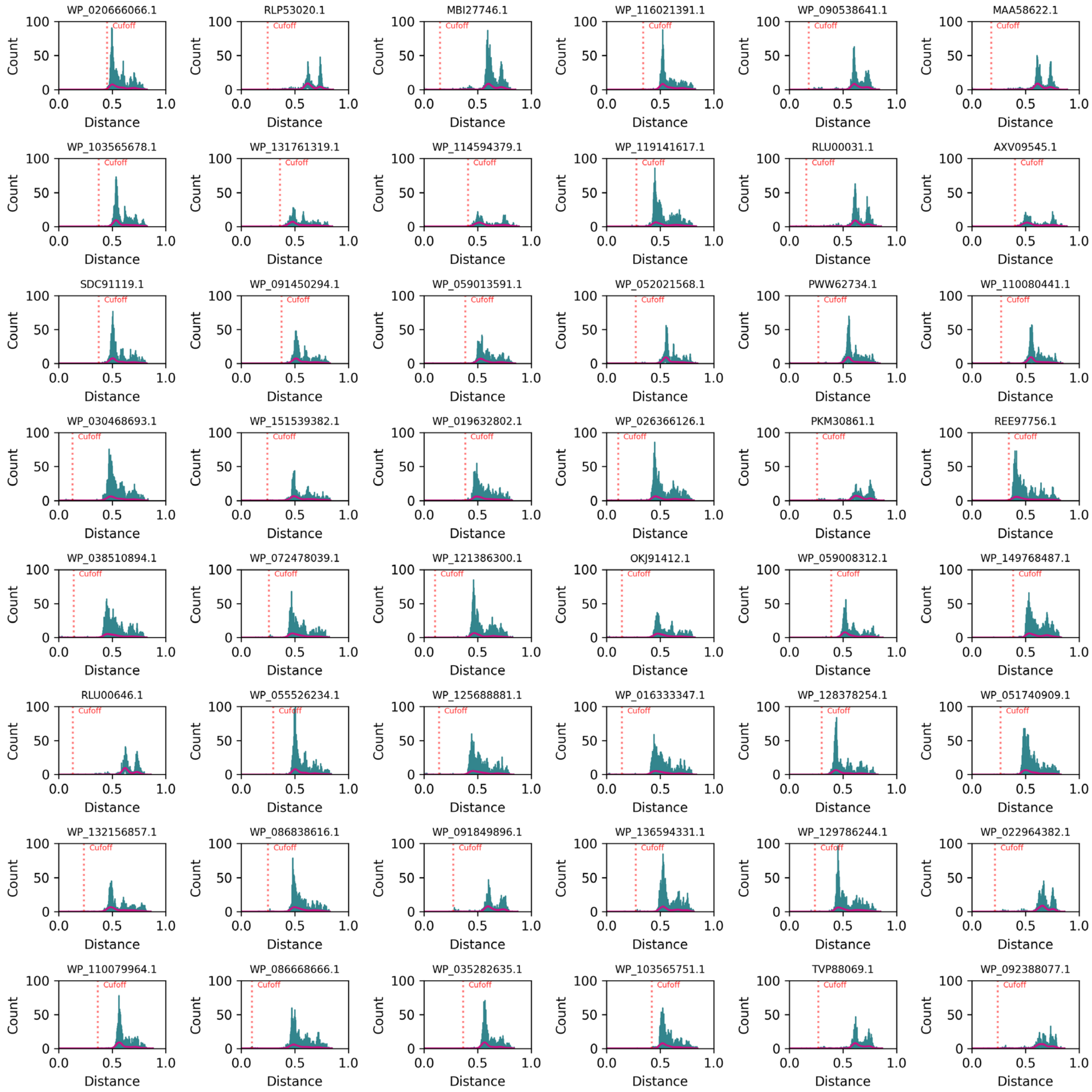


**Fig. S3. Histograms for the 2,064 sequences (continued, 27/43).** Cut-off distance (the first neighbor that is not linked to this sequence with a distance edge) for each sequence is labeled and indicated with a red vertical dashed line. The histograms are shown in order of library id value.

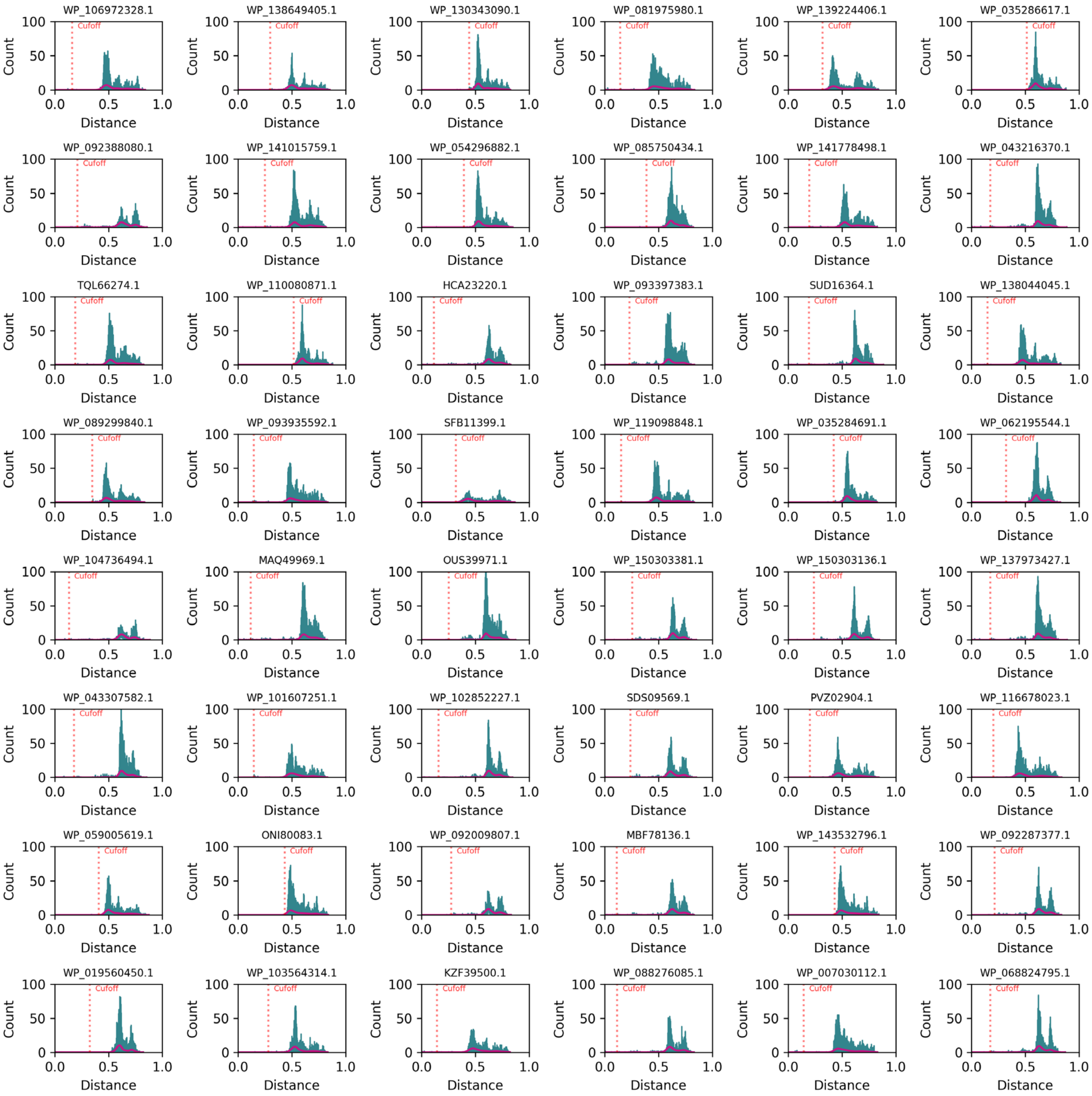


**Fig. S3. Histograms for the 2,064 sequences (continued, 28/43).** Cut-off distance (the first neighbor that is not linked to this sequence with a distance edge) for each sequence is labeled and indicated with a red vertical dashed line. The histograms are shown in order of library id value.

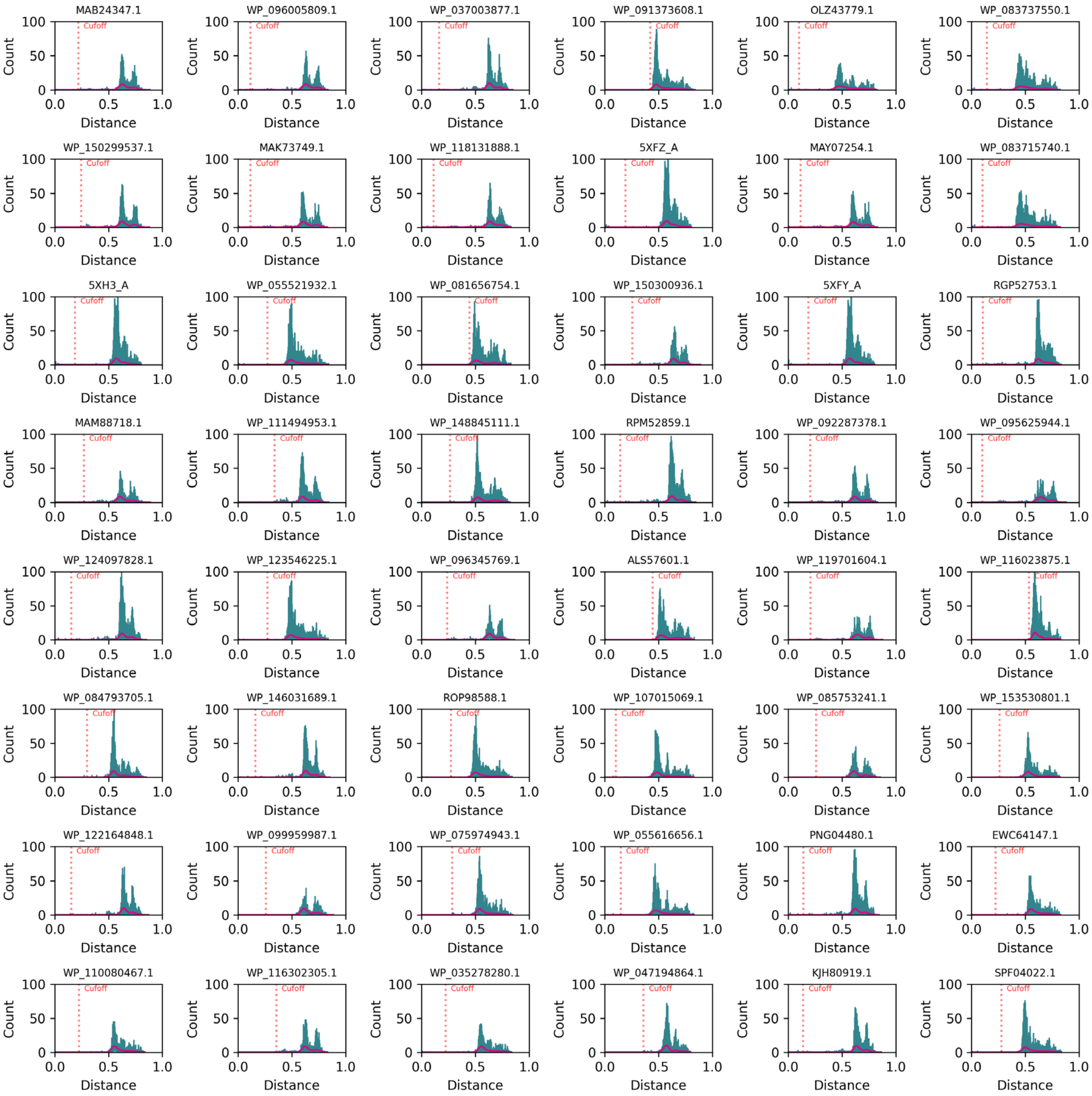


**Fig. S3. Histograms for the 2,064 sequences (continued, 29/43).** Cut-off distance (the first neighbor that is not linked to this sequence with a distance edge) for each sequence is labeled and indicated with a red vertical dashed line. The histograms are shown in order of library id value.

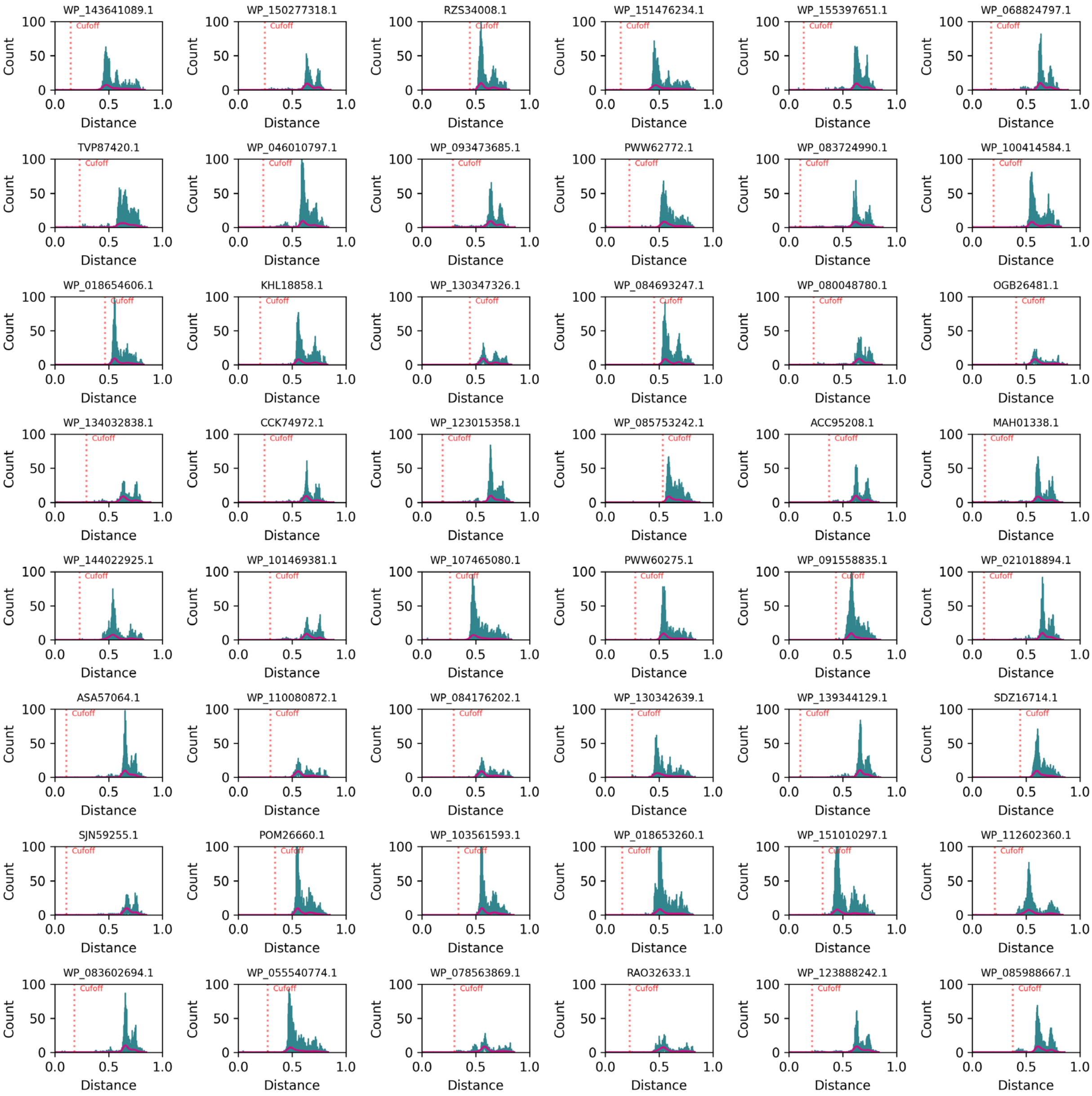


**Fig. S3. Histograms for the 2,064 sequences (continued, 30/43).** Cut-off distance (the first neighbor that is not linked to this sequence with a distance edge) for each sequence is labeled and indicated with a red vertical dashed line. The histograms are shown in order of library id value.

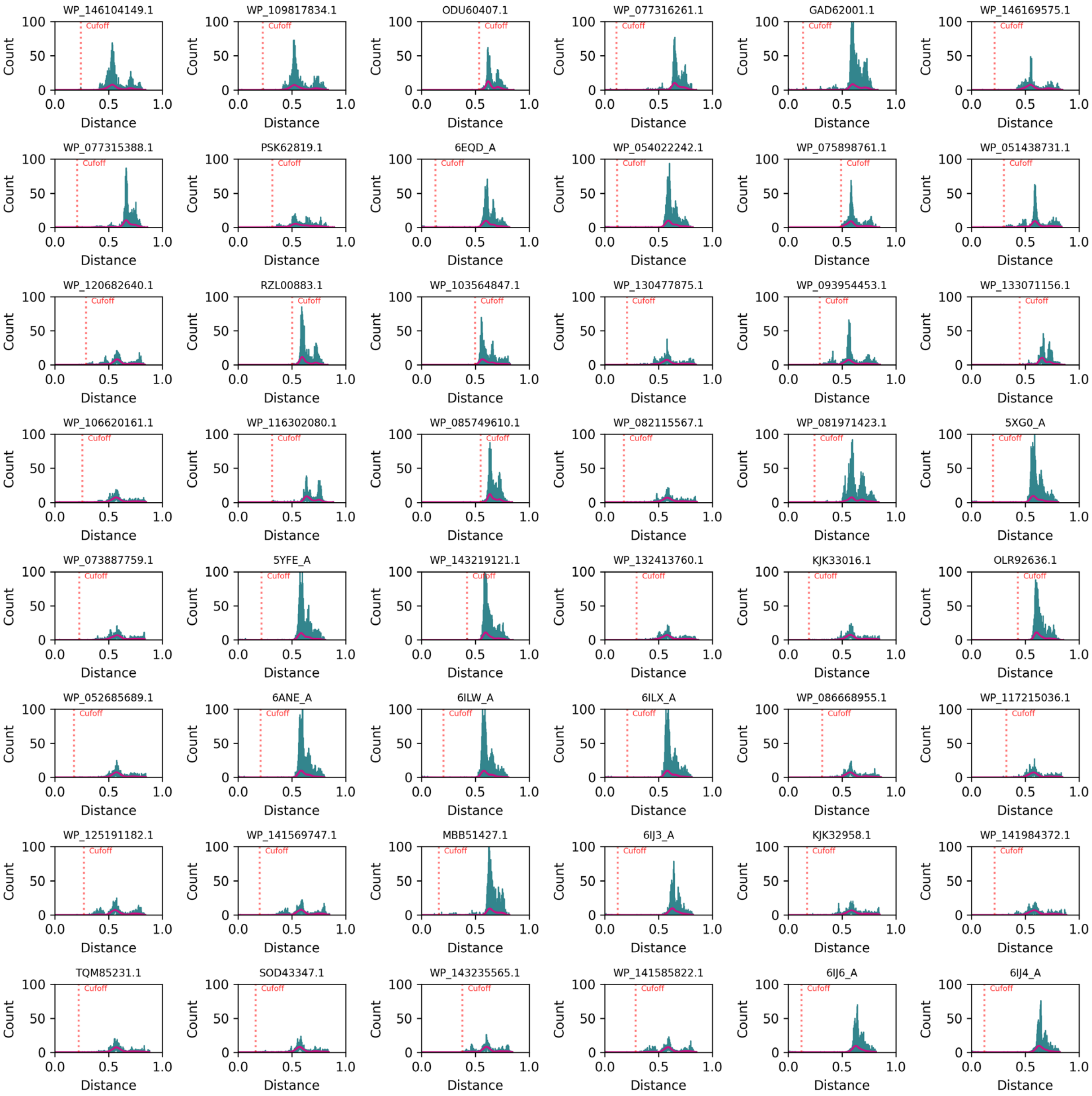


**Fig. S3. Histograms for the 2,064 sequences (continued, 31/43).** Cut-off distance (the first neighbor that is not linked to this sequence with a distance edge) for each sequence is labeled and indicated with a red vertical dashed line. The histograms are shown in order of library id value.

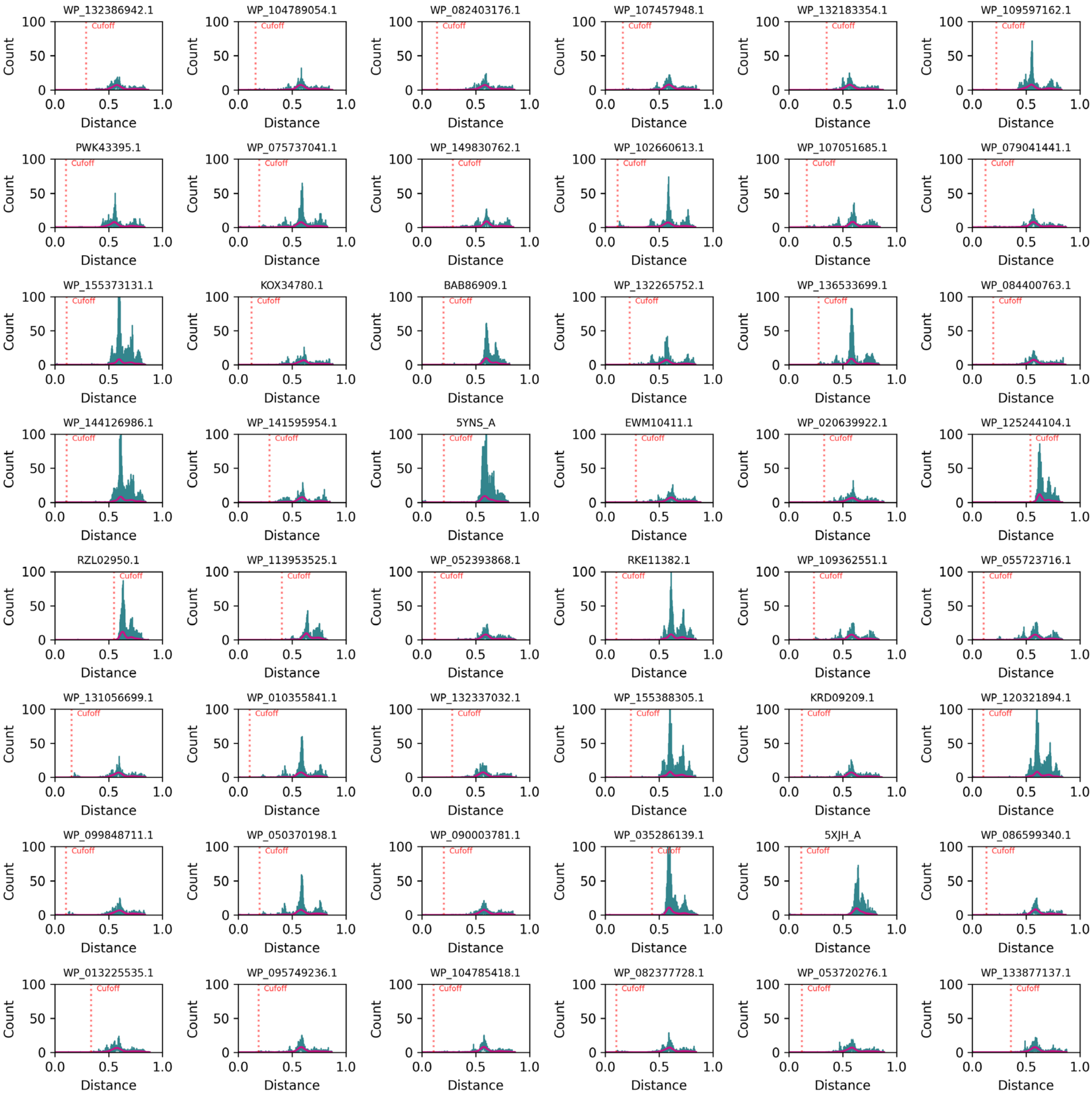


**Fig. S3. Histograms for the 2,064 sequences (continued, 32/43).** Cut-off distance (the first neighbor that is not linked to this sequence with a distance edge) for each sequence is labeled and indicated with a red vertical dashed line. The histograms are shown in order of library id value.

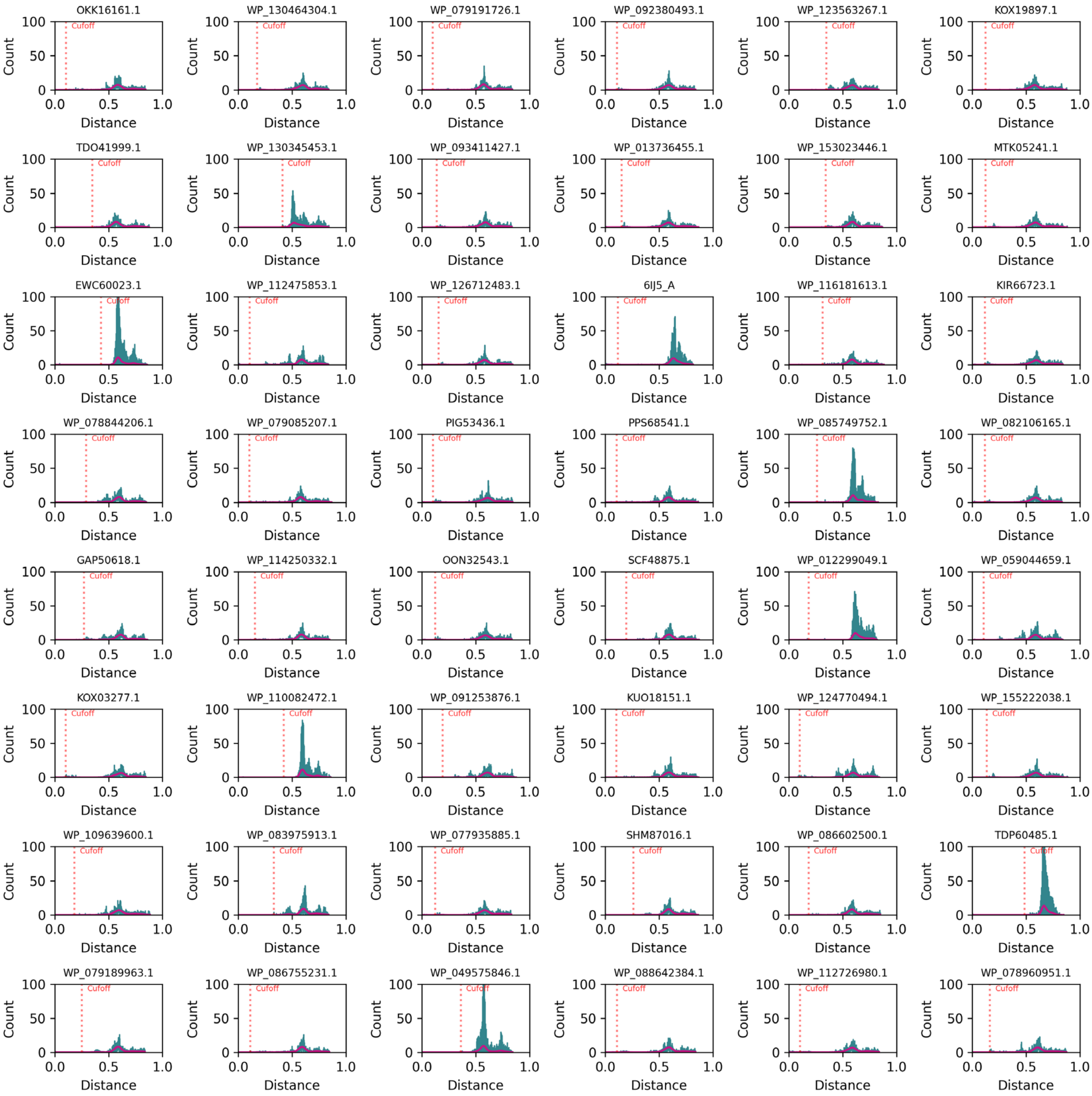


**Fig. S3. Histograms for the 2,064 sequences (continued, 33/43).** Cut-off distance (the first neighbor that is not linked to this sequence with a distance edge) for each sequence is labeled and indicated with a red vertical dashed line. The histograms are shown in order of library id value.

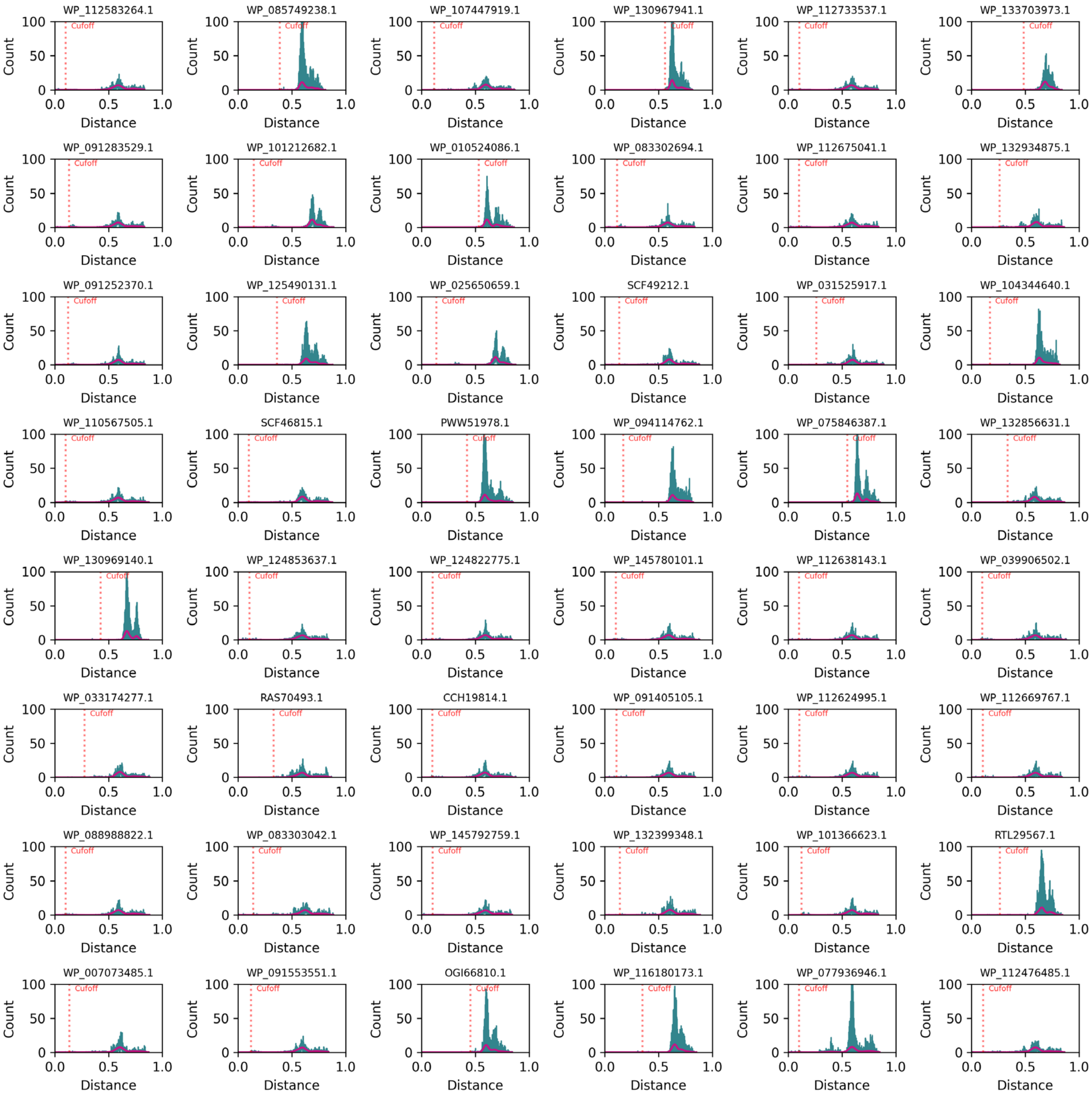


**Fig. S3. Histograms for the 2,064 sequences (continued, 34/43).** Cut-off distance (the first neighbor that is not linked to this sequence with a distance edge) for each sequence is labeled and indicated with a red vertical dashed line. The histograms are shown in order of library id value.

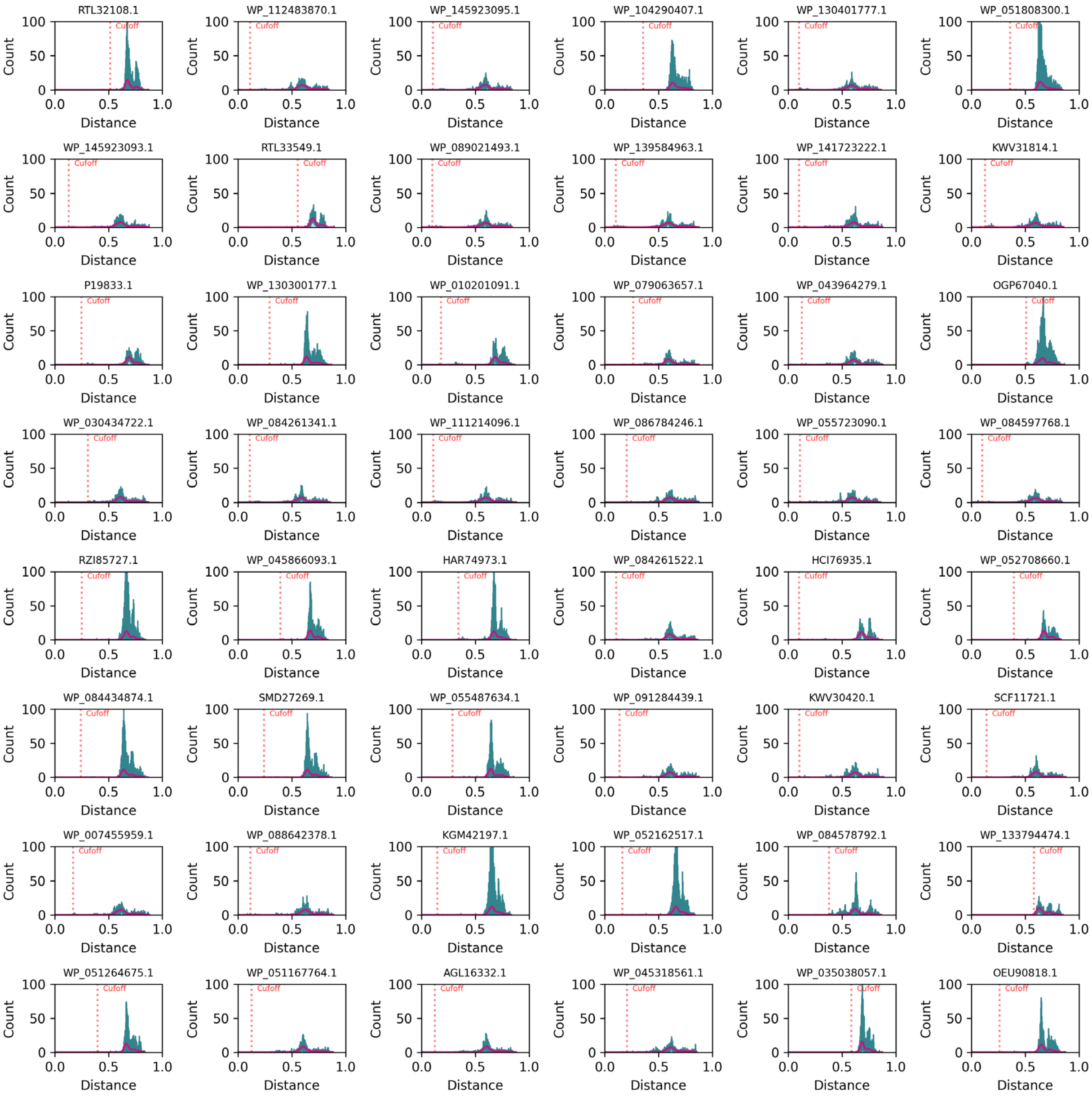


**Fig. S3. Histograms for the 2,064 sequences (continued, 35/43).** Cut-off distance (the first neighbor that is not linked to this sequence with a distance edge) for each sequence is labeled and indicated with a red vertical dashed line. The histograms are shown in order of library id value.

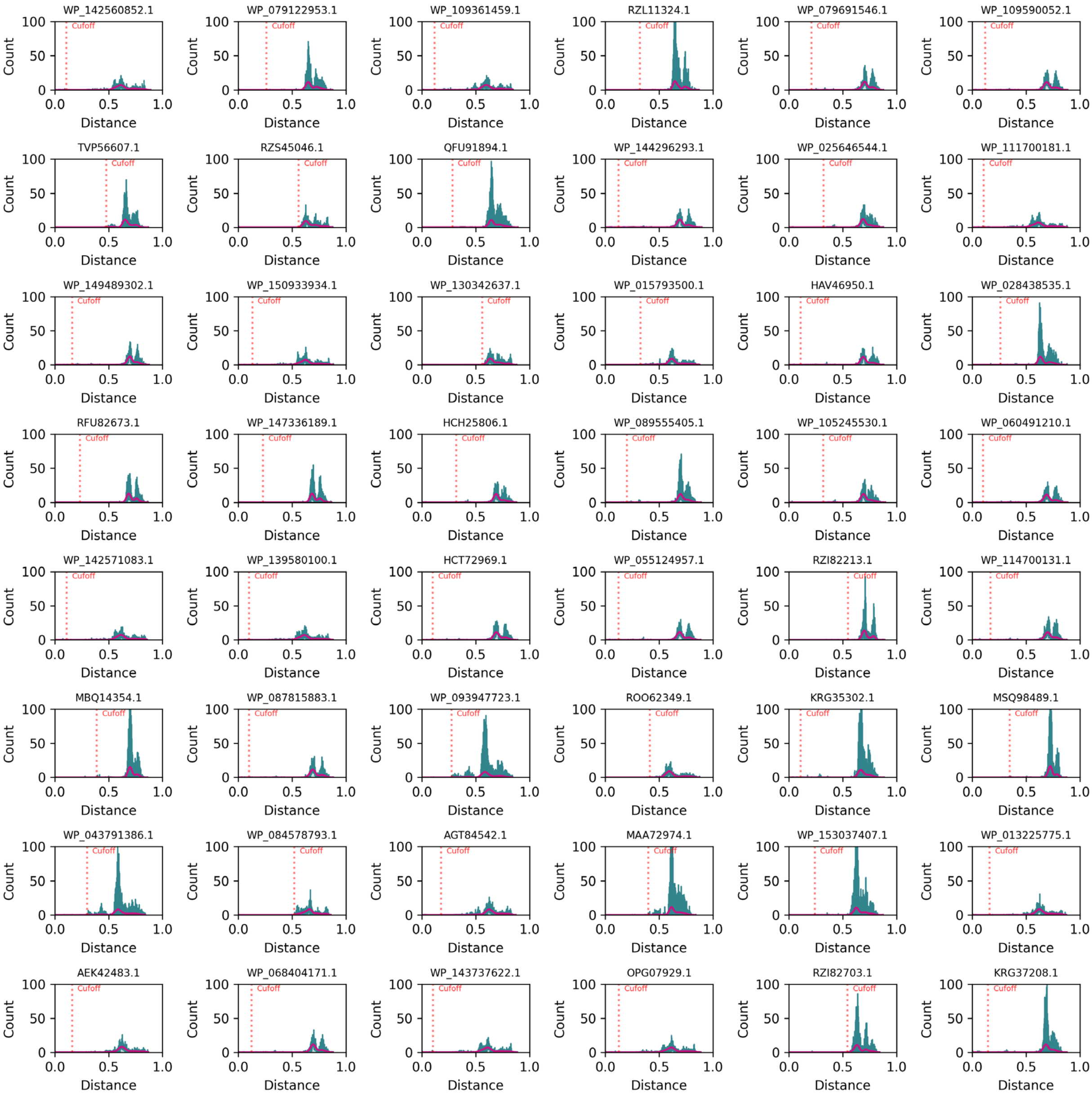


**Fig. S3. Histograms for the 2,064 sequences (continued, 36/43).** Cut-off distance (the first neighbor that is not linked to this sequence with a distance edge) for each sequence is labeled and indicated with a red vertical dashed line. The histograms are shown in order of library id value.

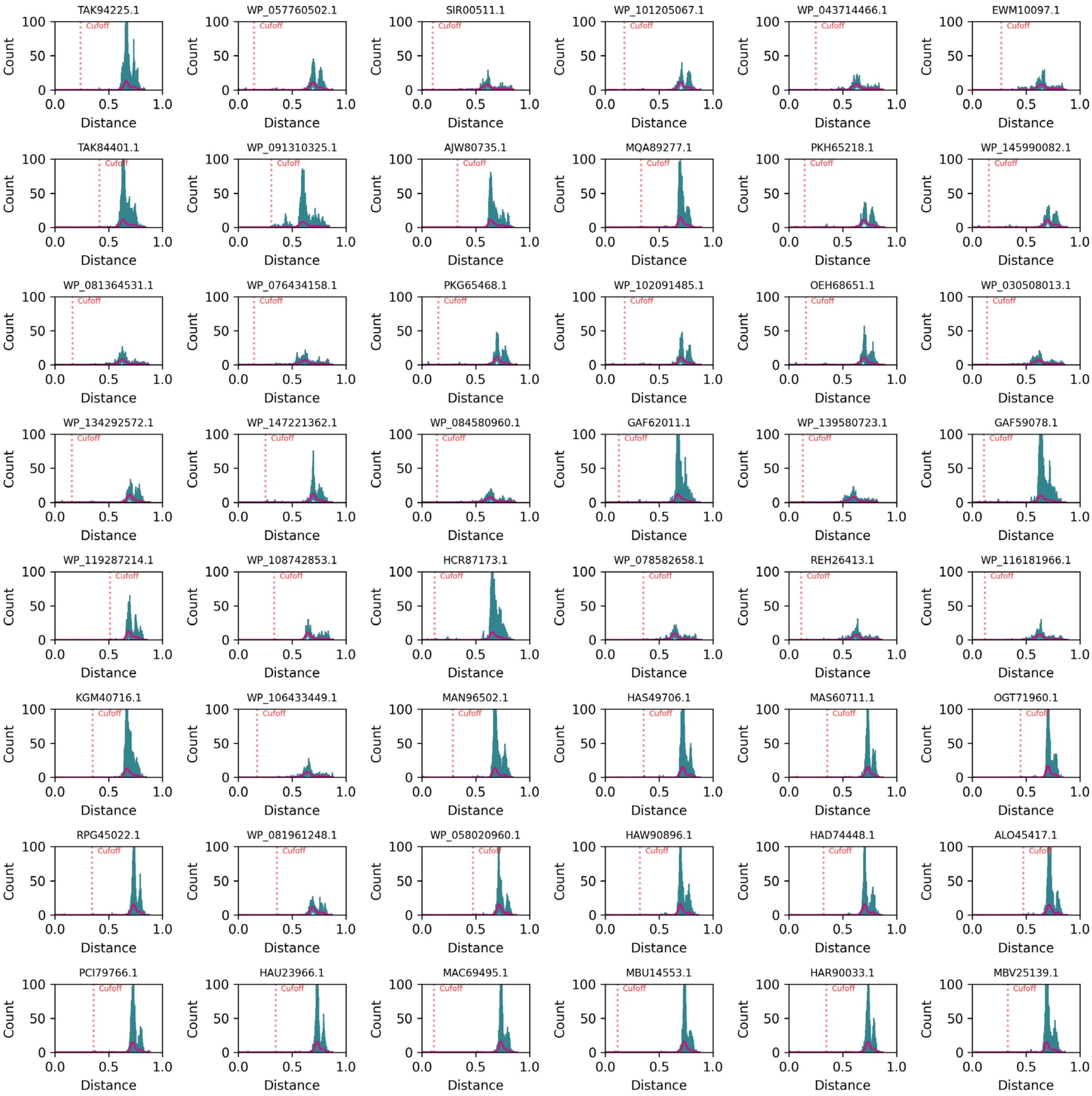


**Fig. S3. Histograms for the 2,064 sequences (continued, 37/43).** Cut-off distance (the first neighbor that is not linked to this sequence with a distance edge) for each sequence is labeled and indicated with a red vertical dashed line. The histograms are shown in order of library id value.

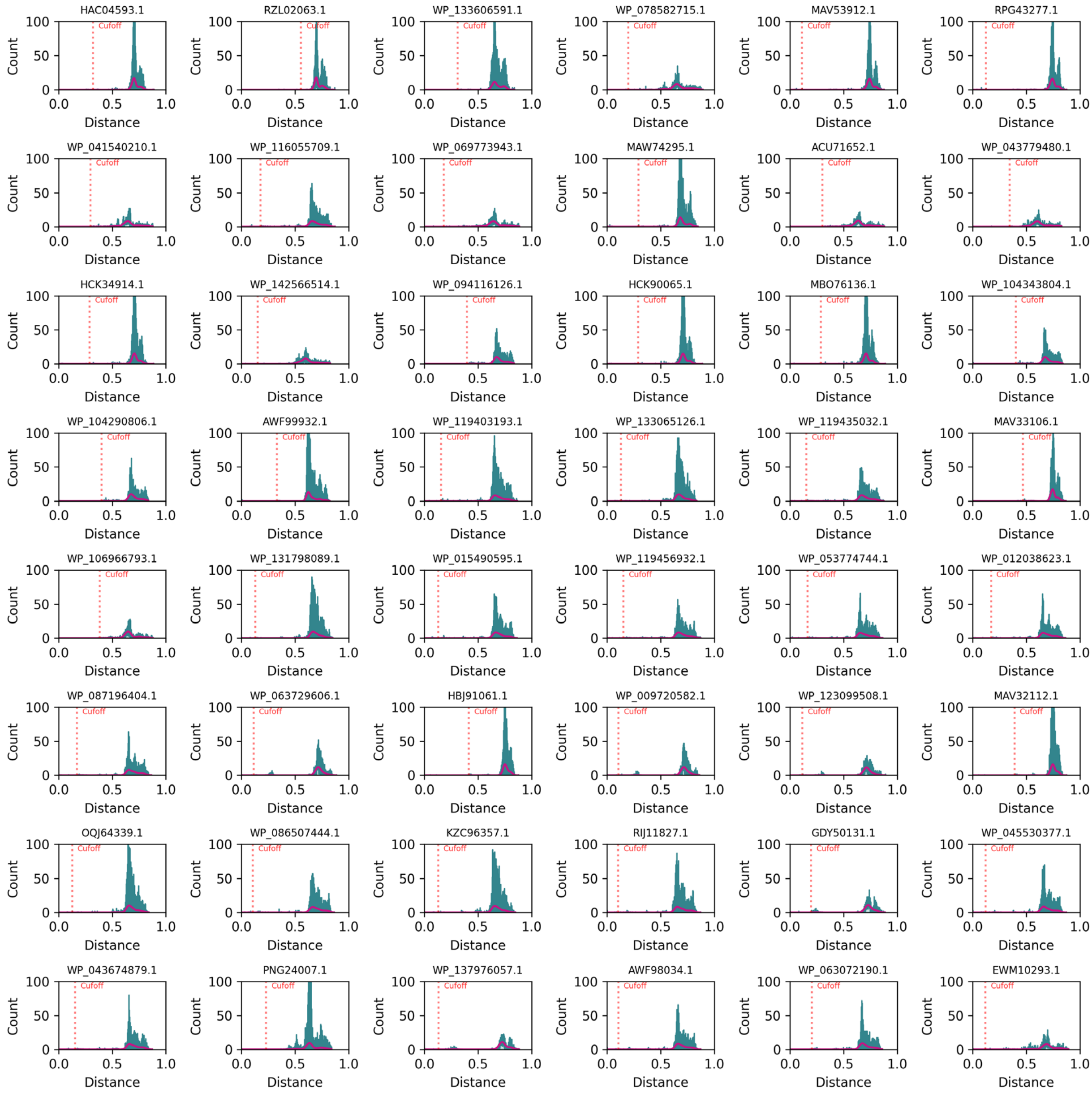


**Fig. S3. Histograms for the 2,064 sequences (continued, 38/43).** Cut-off distance (the first neighbor that is not linked to this sequence with a distance edge) for each sequence is labeled and indicated with a red vertical dashed line. The histograms are shown in order of library id value.

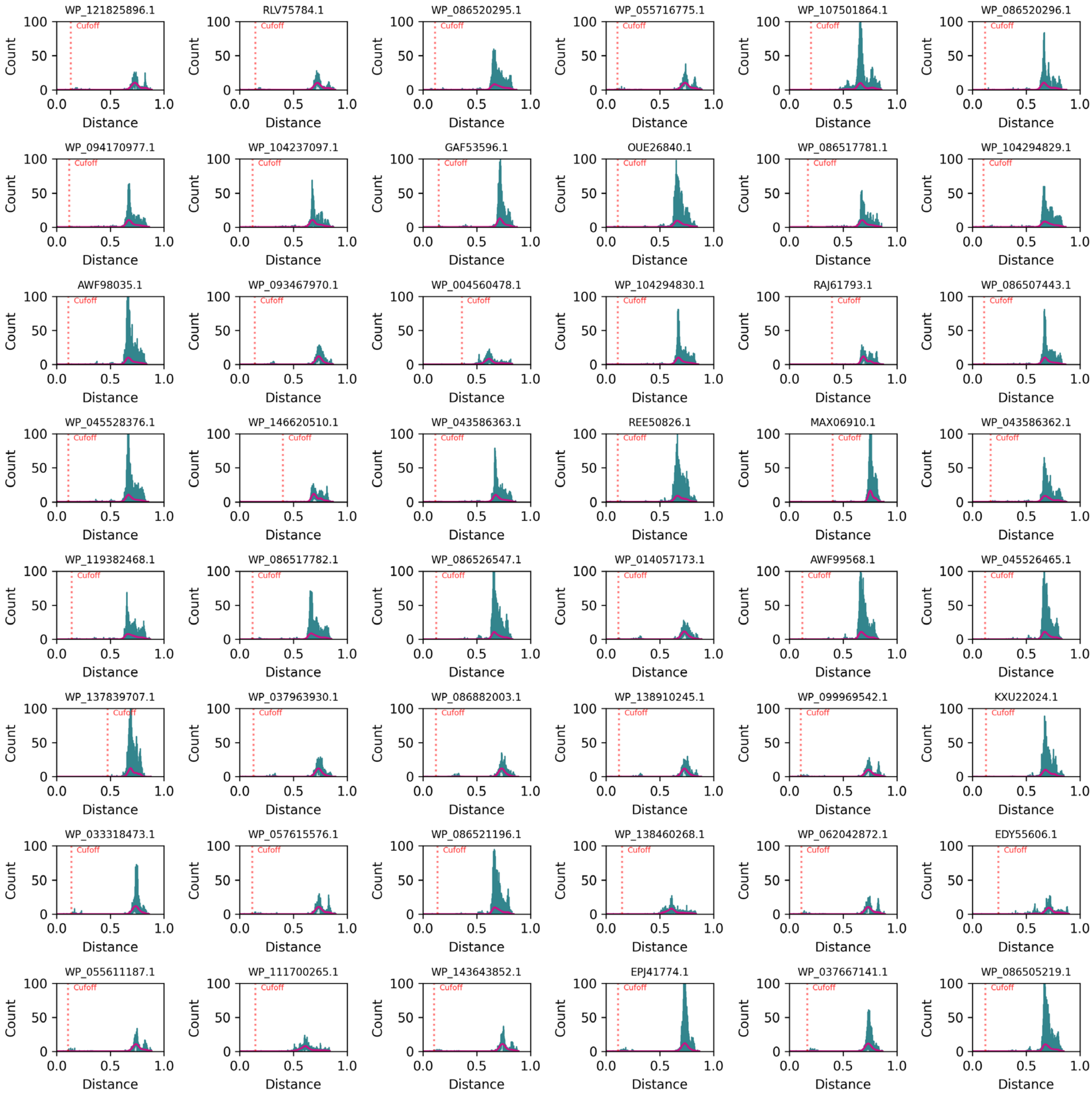


**Fig. S3. Histograms for the 2,064 sequences (continued, 39/43).** Cut-off distance (the first neighbor that is not linked to this sequence with a distance edge) for each sequence is labeled and indicated with a red vertical dashed line. The histograms are shown in order of library id value.

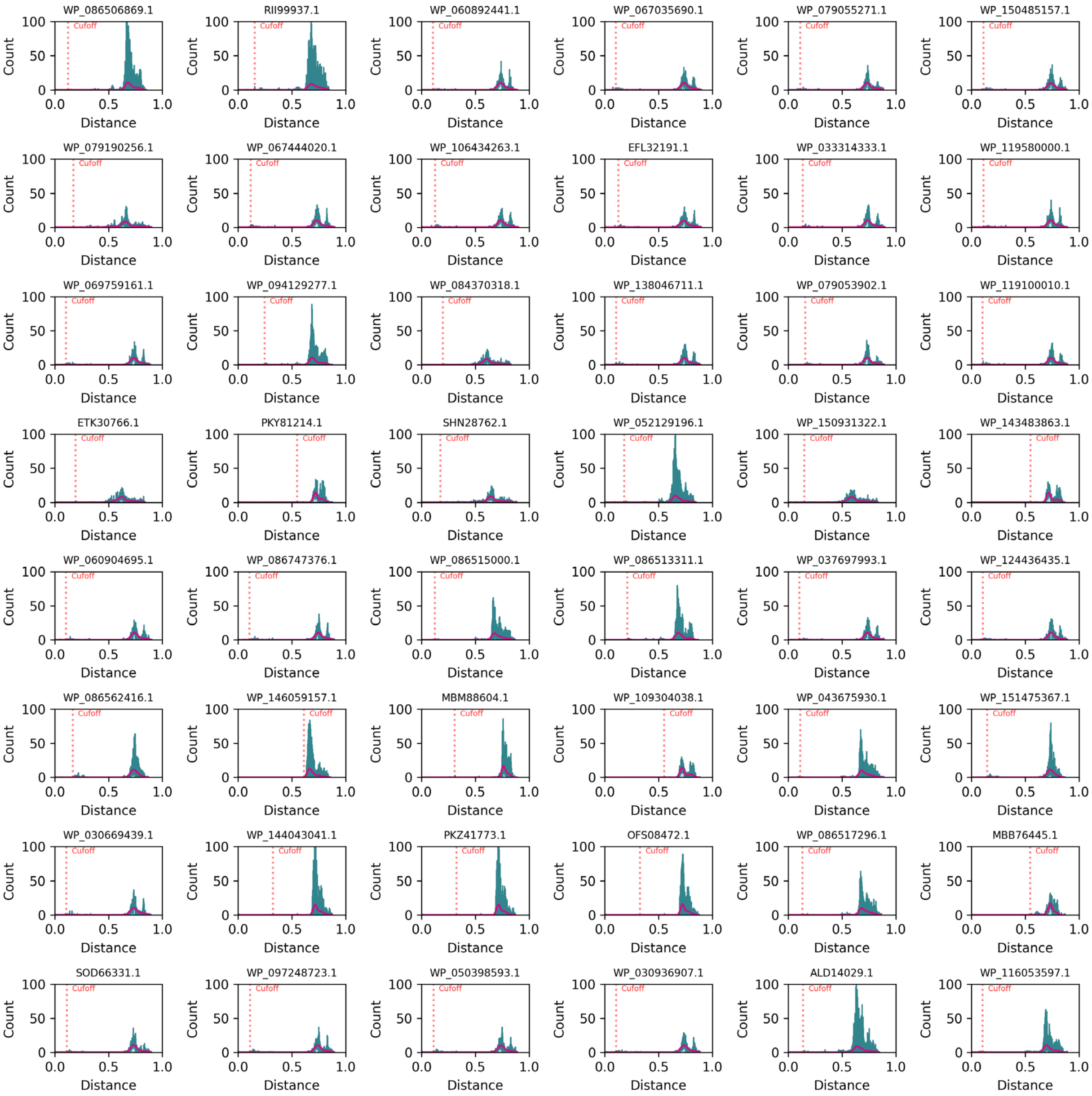


**Fig. S3. Histograms for the 2,064 sequences (continued, 40/43).** Cut-off distance (the first neighbor that is not linked to this sequence with a distance edge) for each sequence is labeled and indicated with a red vertical dashed line. The histograms are shown in order of library id value.

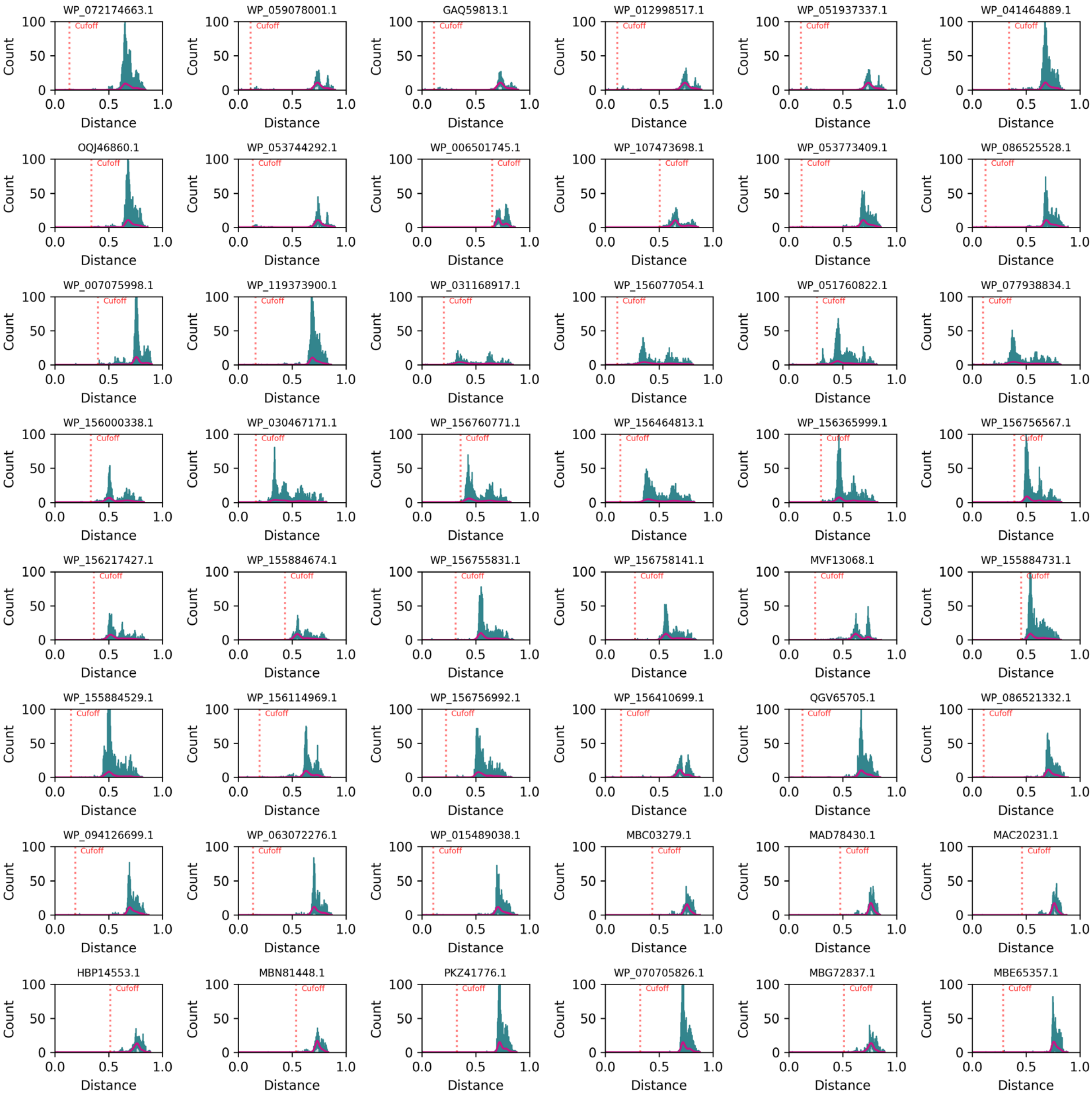


**Fig. S3. Histograms for the 2,064 sequences (continued, 41/43).** Cut-off distance (the first neighbor that is not linked to this sequence with a distance edge) for each sequence is labeled and indicated with a red vertical dashed line. The histograms are shown in order of library id value.

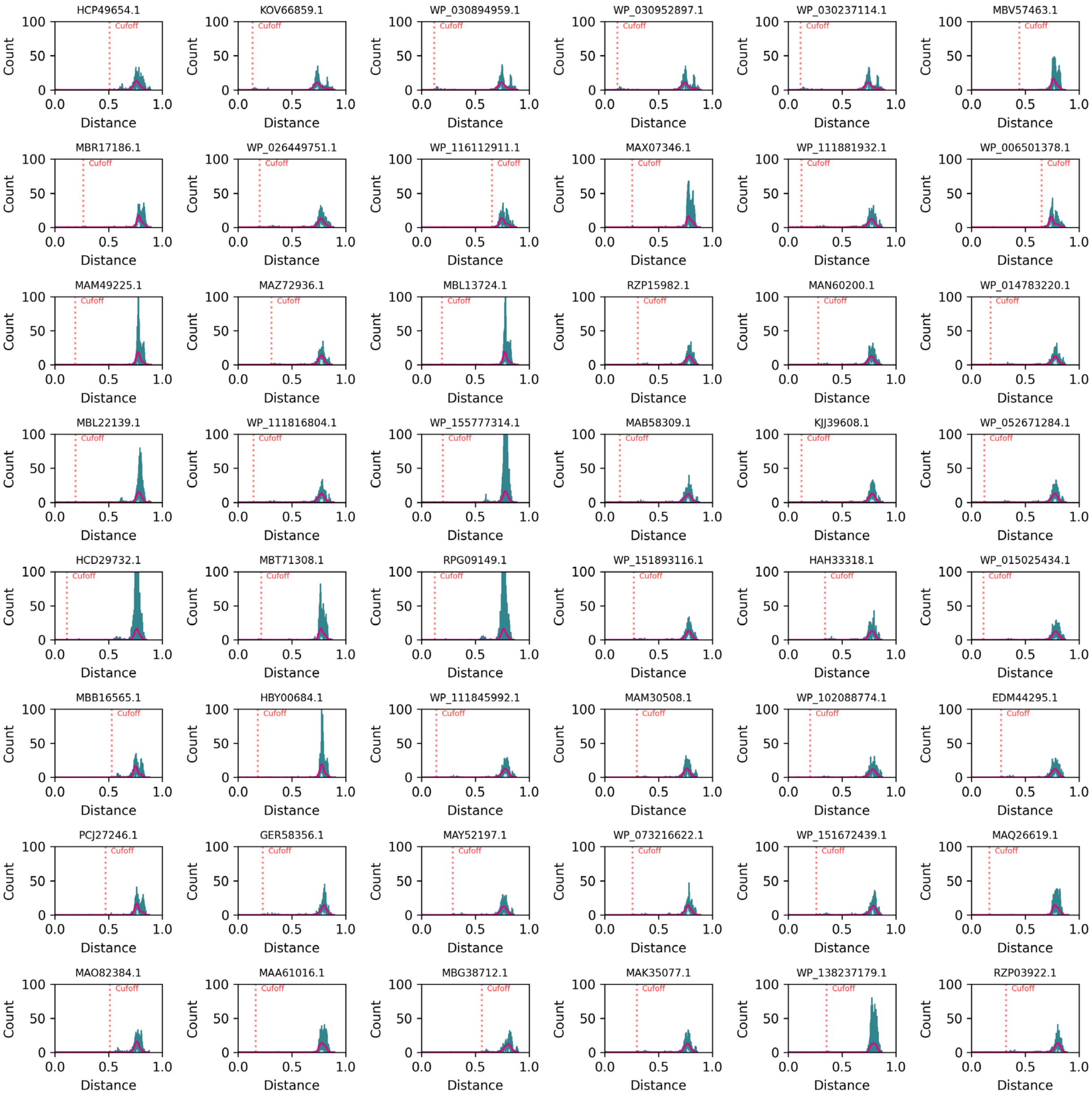


**Fig. S3. Histograms for the 2,064 sequences (continued, 42/43).** Cut-off distance (the first neighbor that is not linked to this sequence with a distance edge) for each sequence is labeled and indicated with a red vertical dashed line. The histograms are shown in order of library id value.

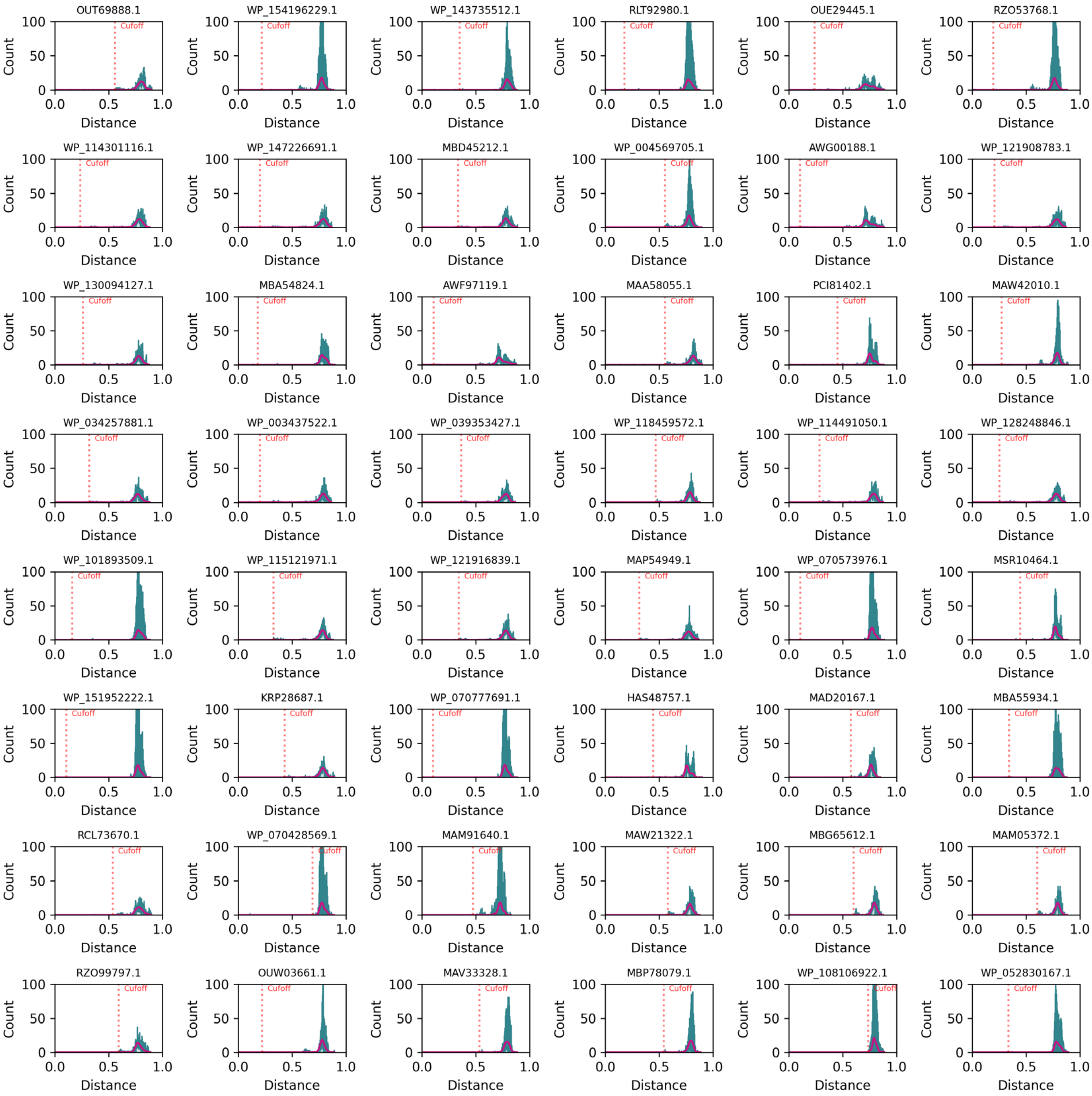


**Fig. S3. Histograms for the 2,064 sequences (43/43).** Cut-off distance (the first neighbor that is not linked to this sequence with a distance edge) for each sequence is labeled and indicated with a red vertical dashed line. The histograms are shown in order of library id value.

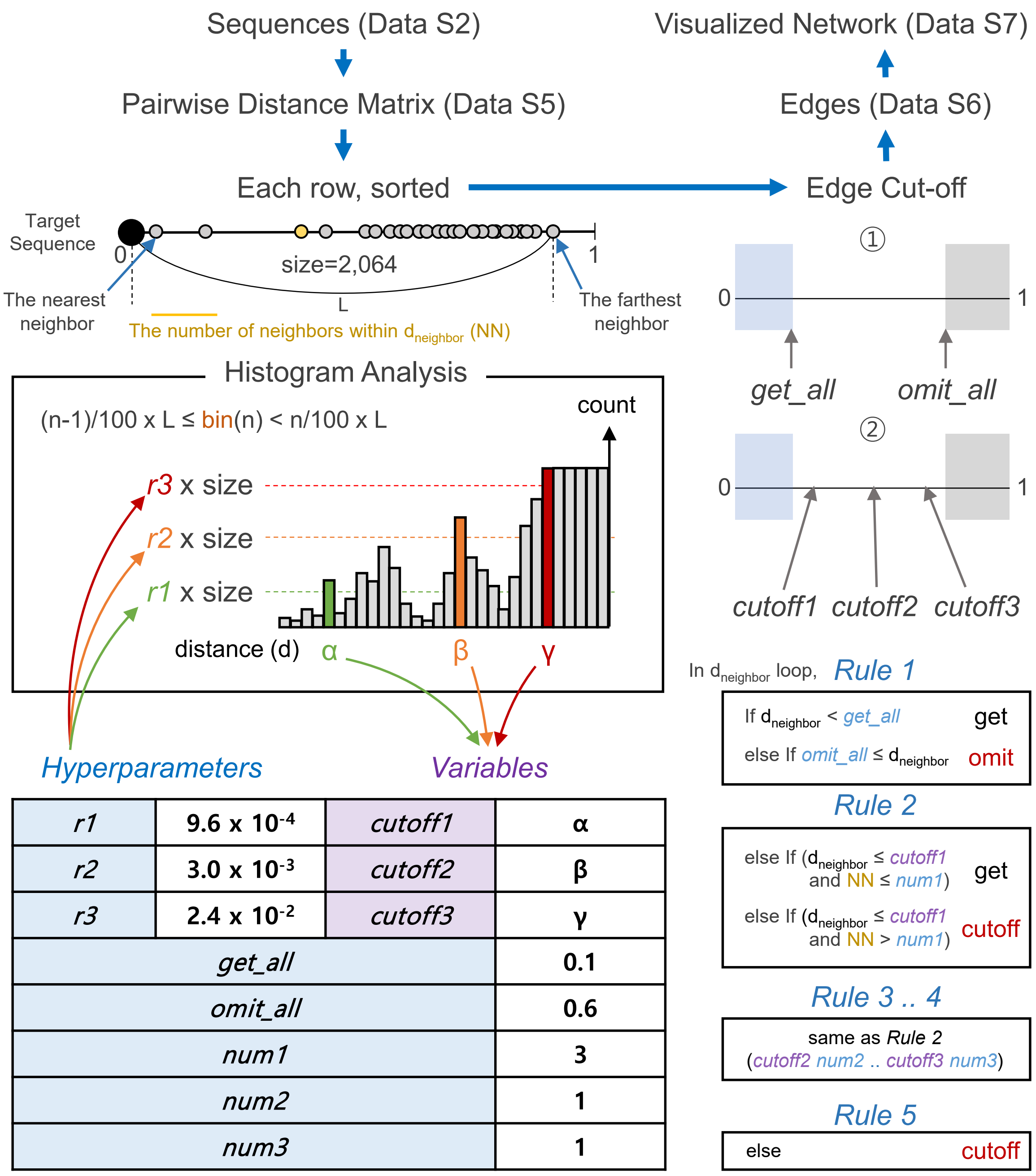


| | | | |
|---|---|---|---|
| *r1* | $9.6 \times 10^{-4}$ | *cutoff1* | α |
| *r2* | $3.0 \times 10^{-3}$ | *cutoff2* | β |
| *r3* | $2.4 \times 10^{-2}$ | *cutoff3* | γ |
| *get_all* | | | 0.1 |
| *omit_all* | | | 0.6 |
| *num1* | | | 3 |
| *num2* | | | 1 |
| *num3* | | | 1 |

**Fig. S4. A heuristic algorithm of density analysis with eight hyperparameters.** The algorithm flow is indicated by blue arrows. The eight hyperparameters, *r1*, *r2*, *r3*, *get_all*, *omit_all*, *num1*, *num2*, and *num3* and their relationship with variables, *cutoff1*, *cutoff2*, and *cutoff3*, determined from the histogram analysis are shown.

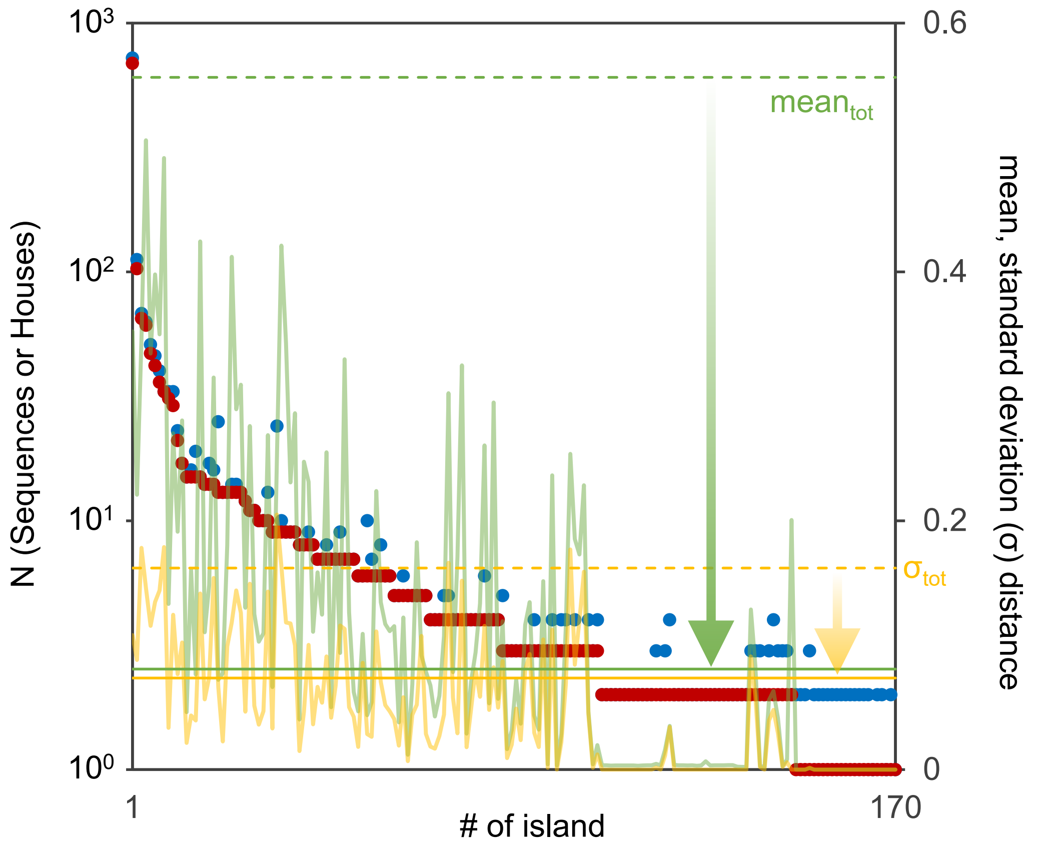


**Fig. S5. The number of sequences/Houses and the mean and standard deviation values of edge distance in each Islands.** Horizontal axis indicates Island number ranging from 1 to 170. The number of sequences/Houses (N) is shown with blue and red circles, respectively. The line graphs indicates the mean and standard deviation values of each islands. The dashed and solid horizontal lines indicate those of total edge distances before clustering and after clustering, respectively. Arrows represent clustering effect.

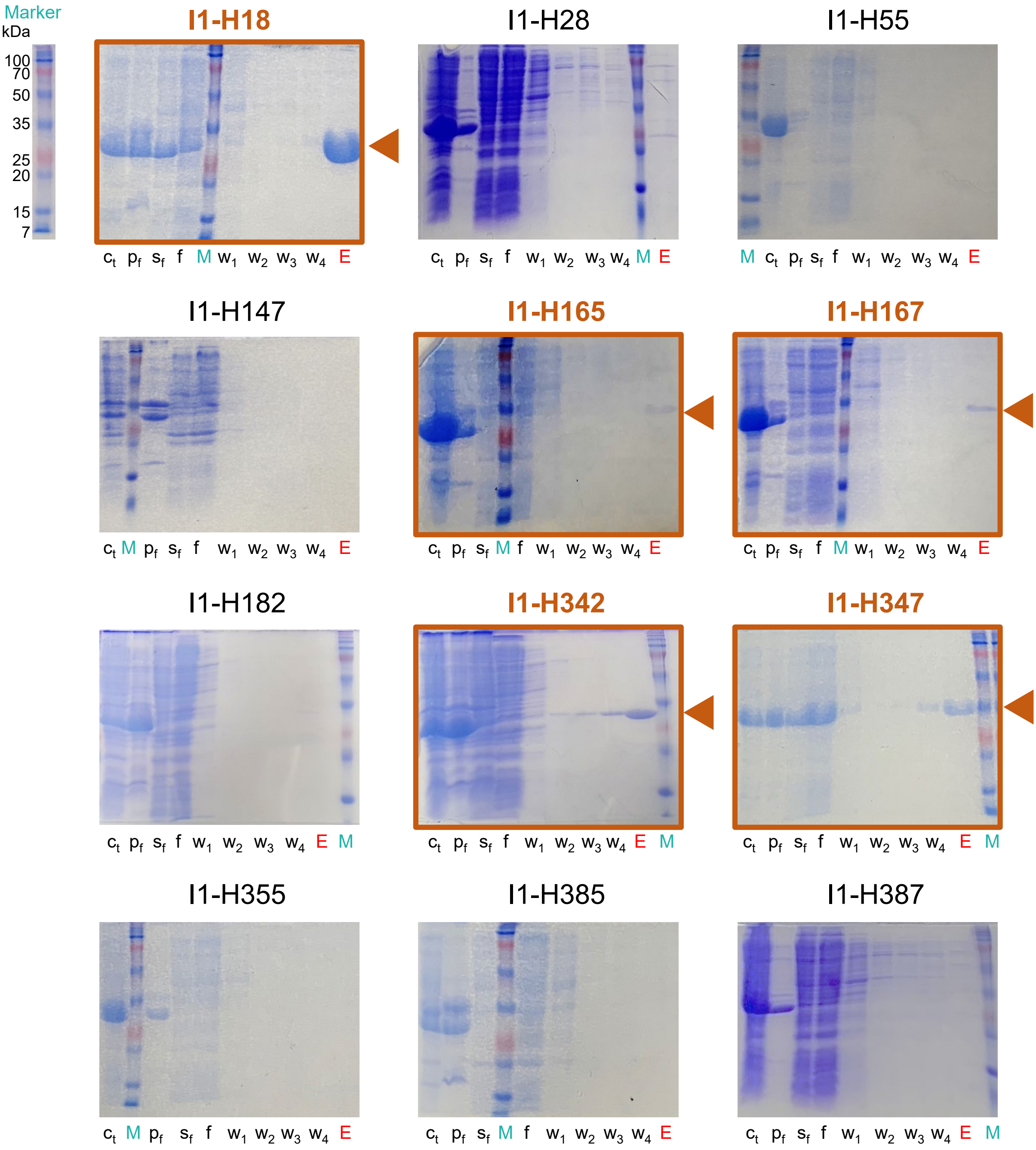


**Fig. S6. Expression and purification of 289 Houses (continued, 1/25).** Abbreviations: Ct, cell total lysate; Pf, pellet fraction; Sf, soluble fraction; W1-4, wash; E, elution; M, marker.

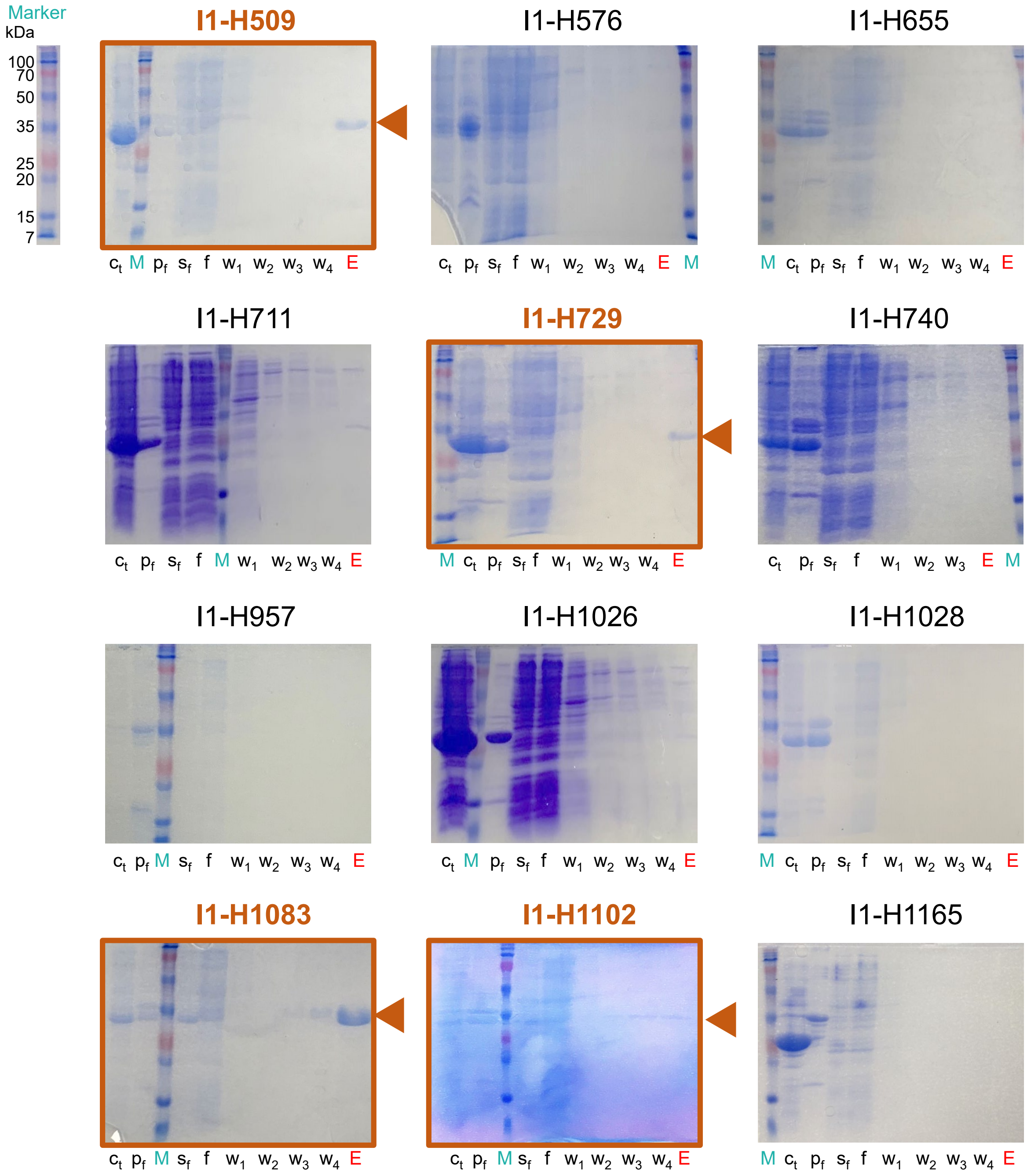


**Fig. S6. Expression and purification of 289 Houses (continued, 2/25).** Abbreviations: Ct, cell total lysate; Pf, pellet fraction; Sf, soluble fraction; W1-4, wash; E, elution; M, marker.

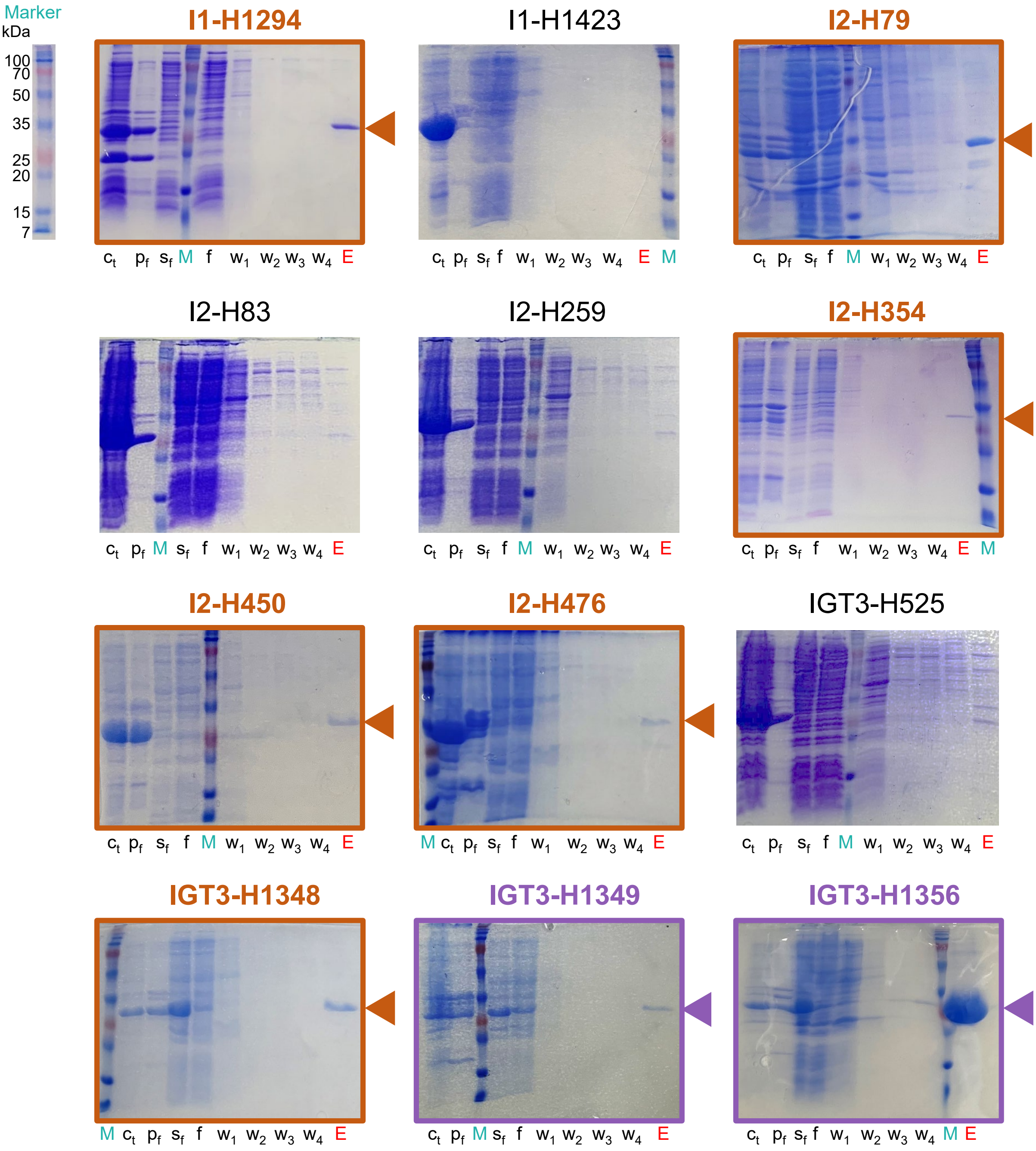


**Fig. S6. Expression and purification of 289 Houses (continued, 3/25).** Abbreviations: Ct, cell total lysate; Pf, pellet fraction; Sf, soluble fraction; W1-4, wash; E, elution; M, marker.

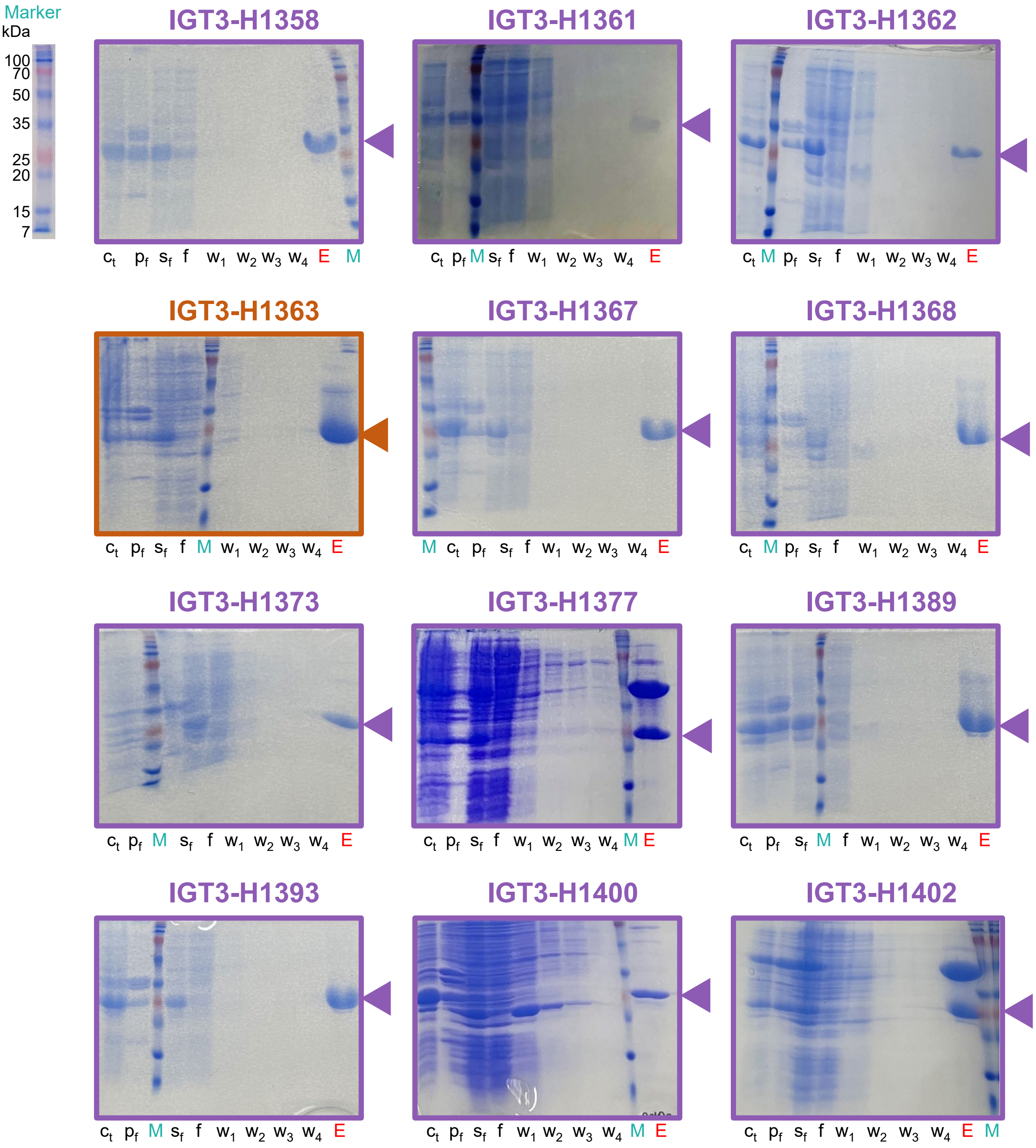


**Fig. S6. Expression and purification of 289 Houses (continued, 4/25).** Abbreviations: Ct, cell total lysate; Pf, pellet fraction; Sf, soluble fraction; W1-4, wash; E, elution; M, marker.

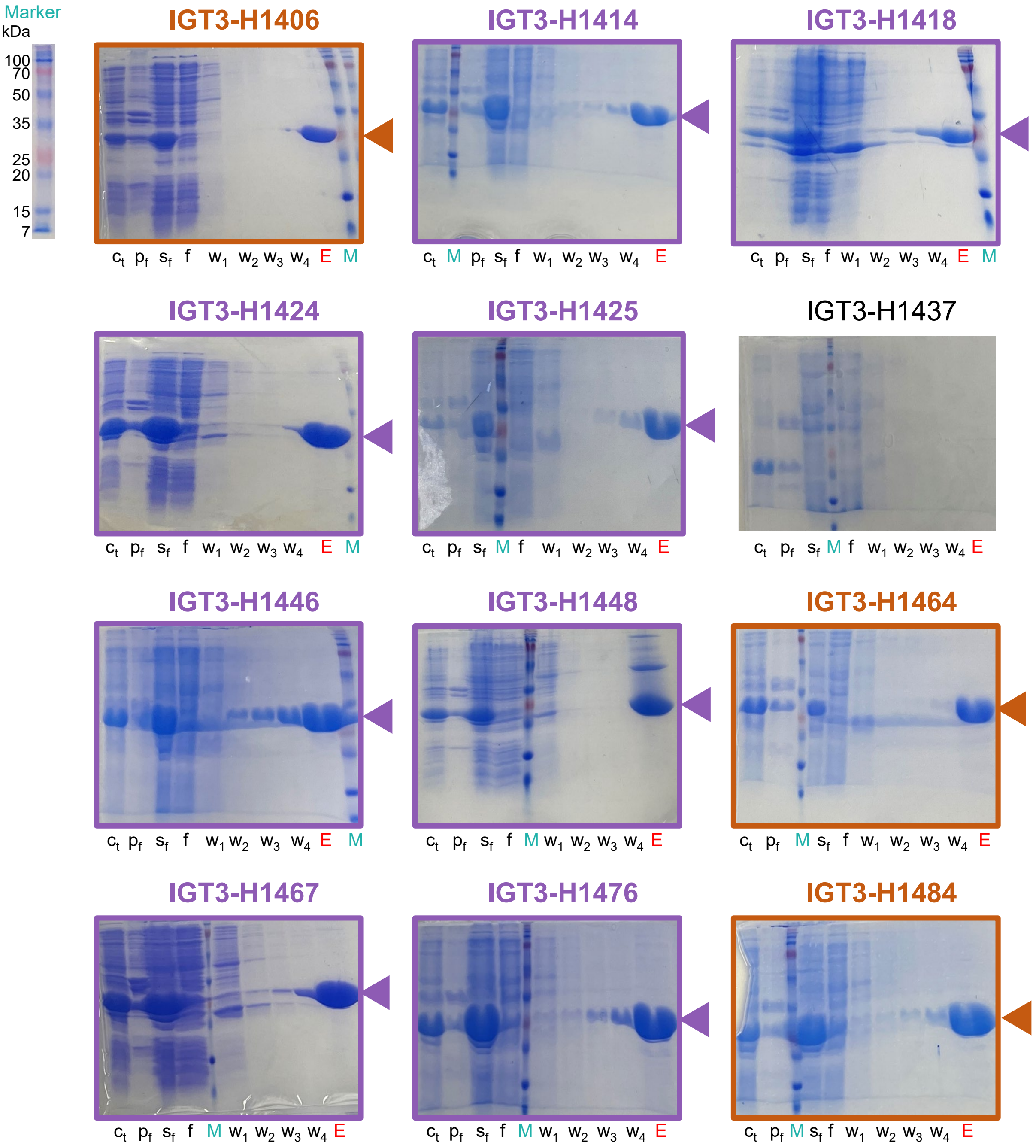


**Fig. S6. Expression and purification of 289 Houses (continued, 5/25).** Abbreviations: Ct, cell total lysate; Pf, pellet fraction; Sf, soluble fraction; W1-4, wash; E, elution; M, marker.

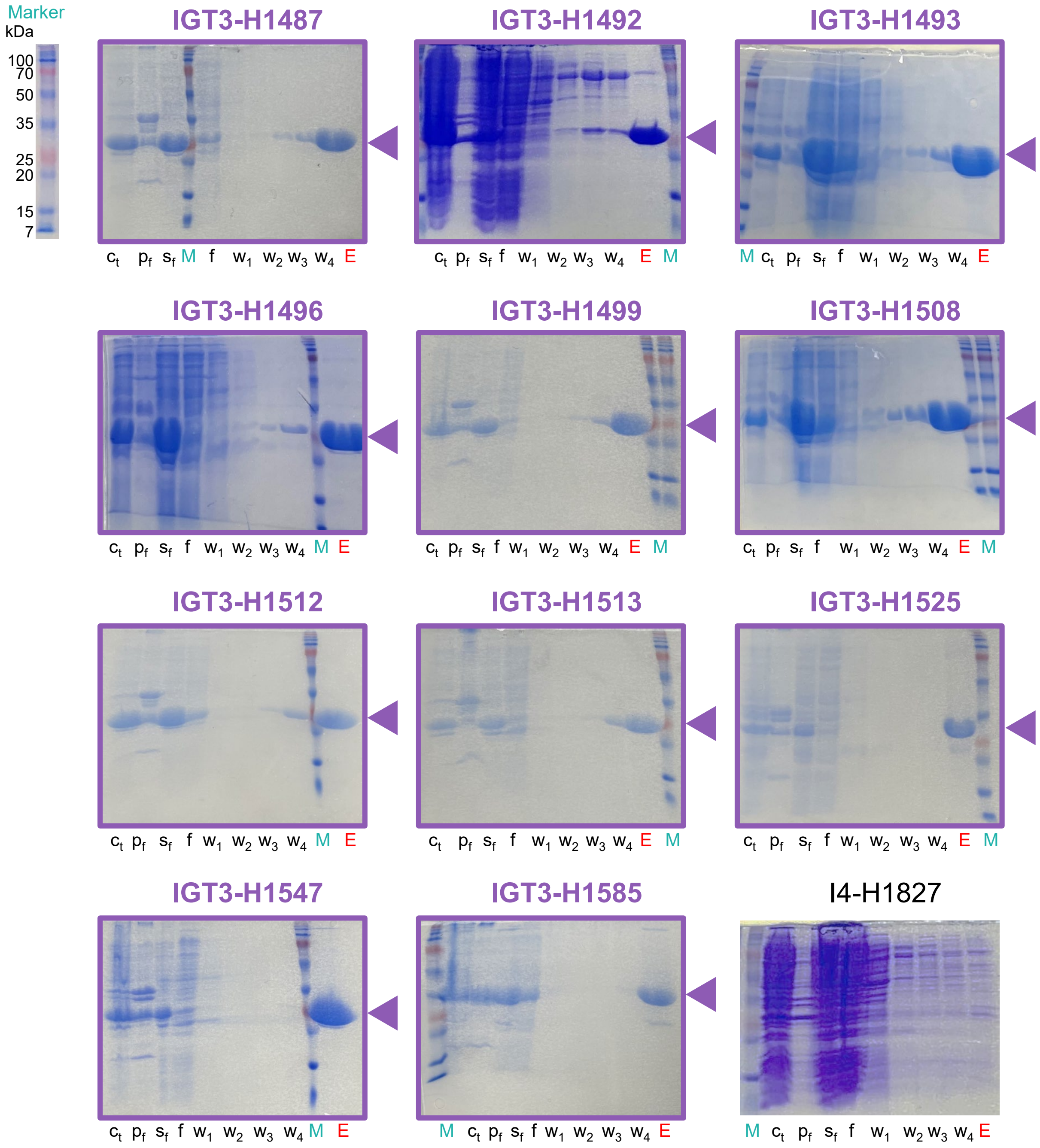


**Fig. S6. Expression and purification of 289 Houses (continued, 6/25).** Abbreviations: Ct, cell total lysate; Pf, pellet fraction; Sf, soluble fraction; W1-4, wash; E, elution; M, marker.

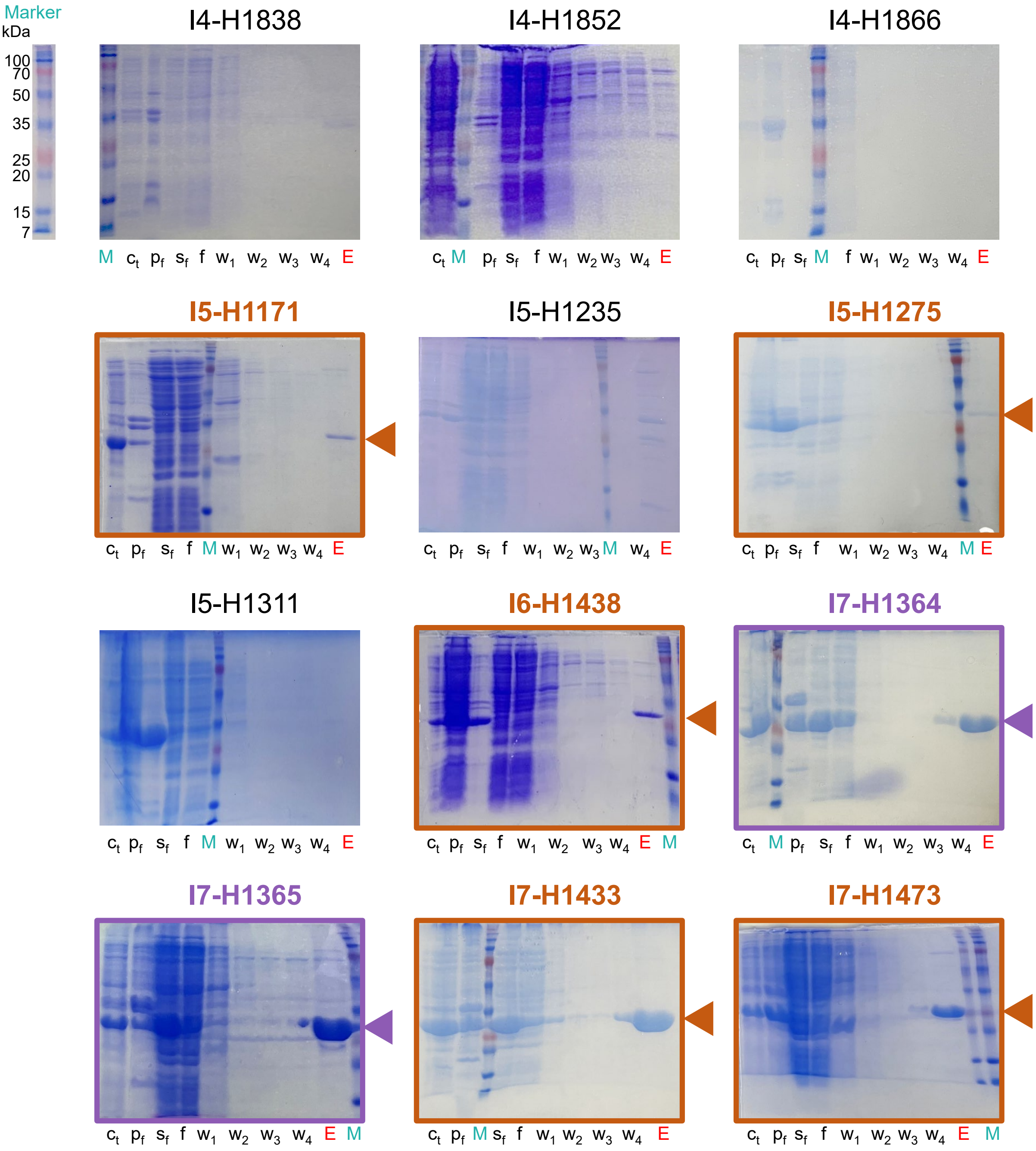


**Fig. S6. Expression and purification of 289 Houses (continued, 7/25).** Abbreviations: Ct, cell total lysate; Pf, pellet fraction; Sf, soluble fraction; W1-4, wash; E, elution; M, marker.

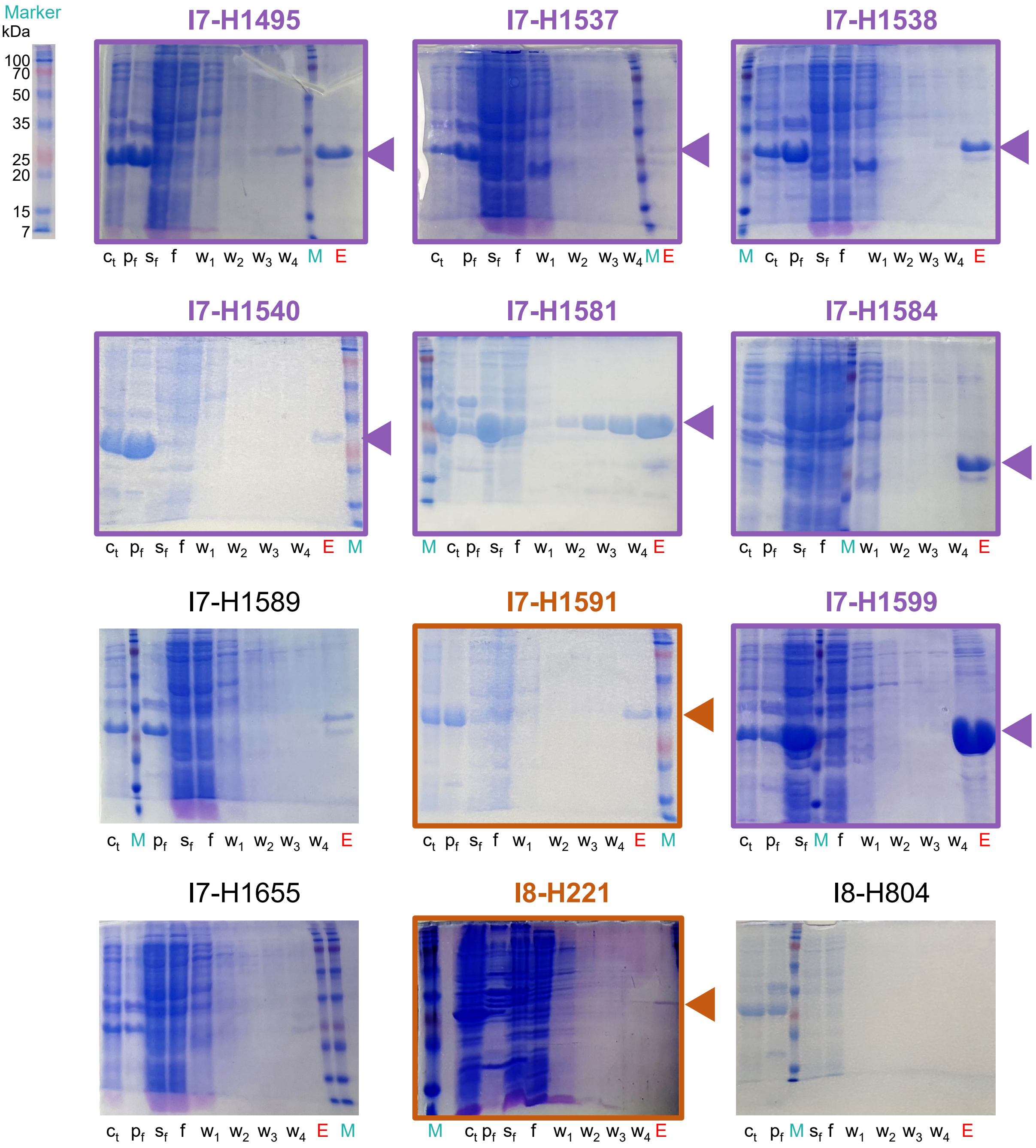


**Fig. S6. Expression and purification of 289 Houses (continued, 8/25).** Abbreviations: Ct, cell total lysate; Pf, pellet fraction; Sf, soluble fraction; W1-4, wash; E, elution; M, marker.

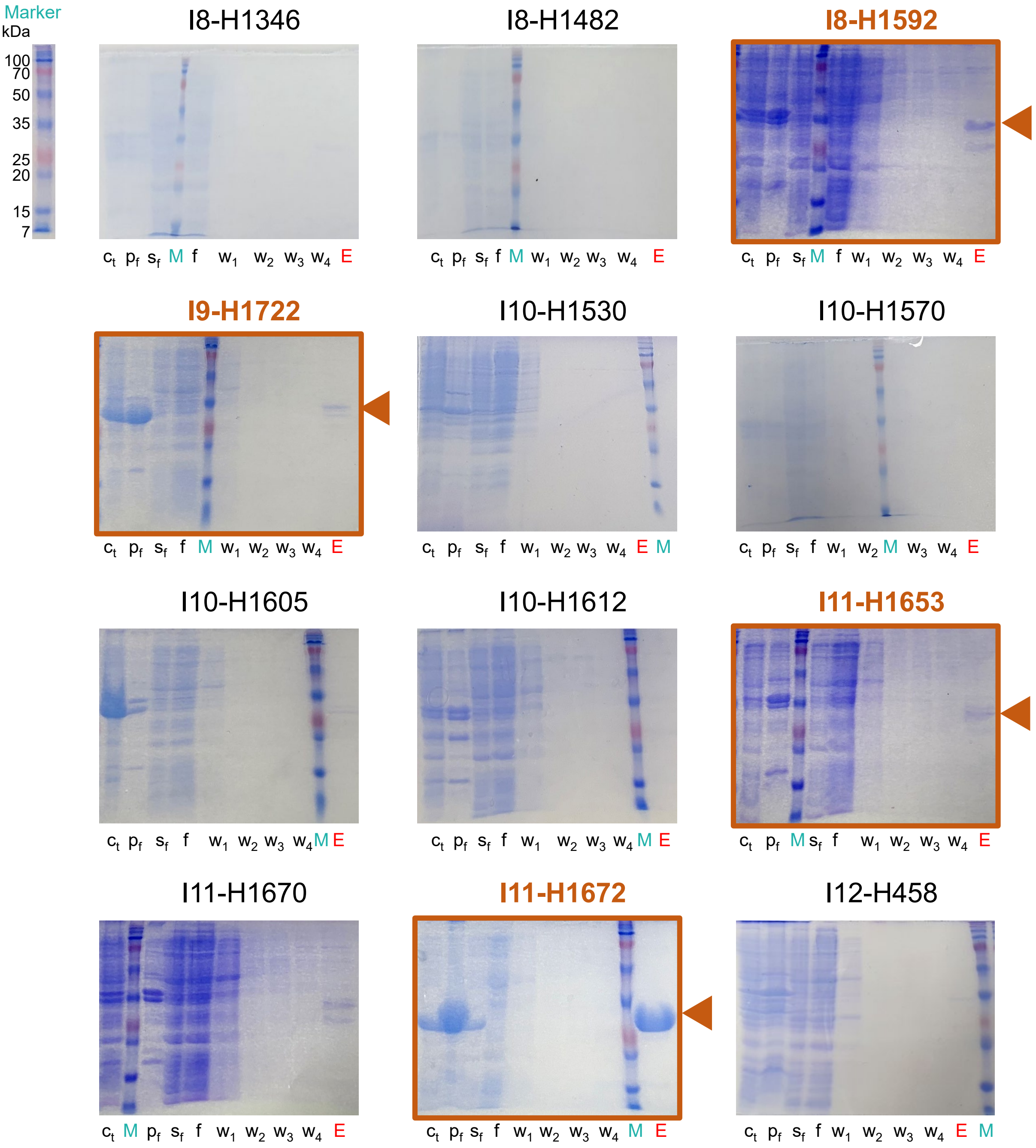


**Fig. S6. Expression and purification of 289 Houses (continued, 9/25).** Abbreviations: Ct, cell total lysate; Pf, pellet fraction; Sf, soluble fraction; W1-4, wash; E, elution; M, marker.

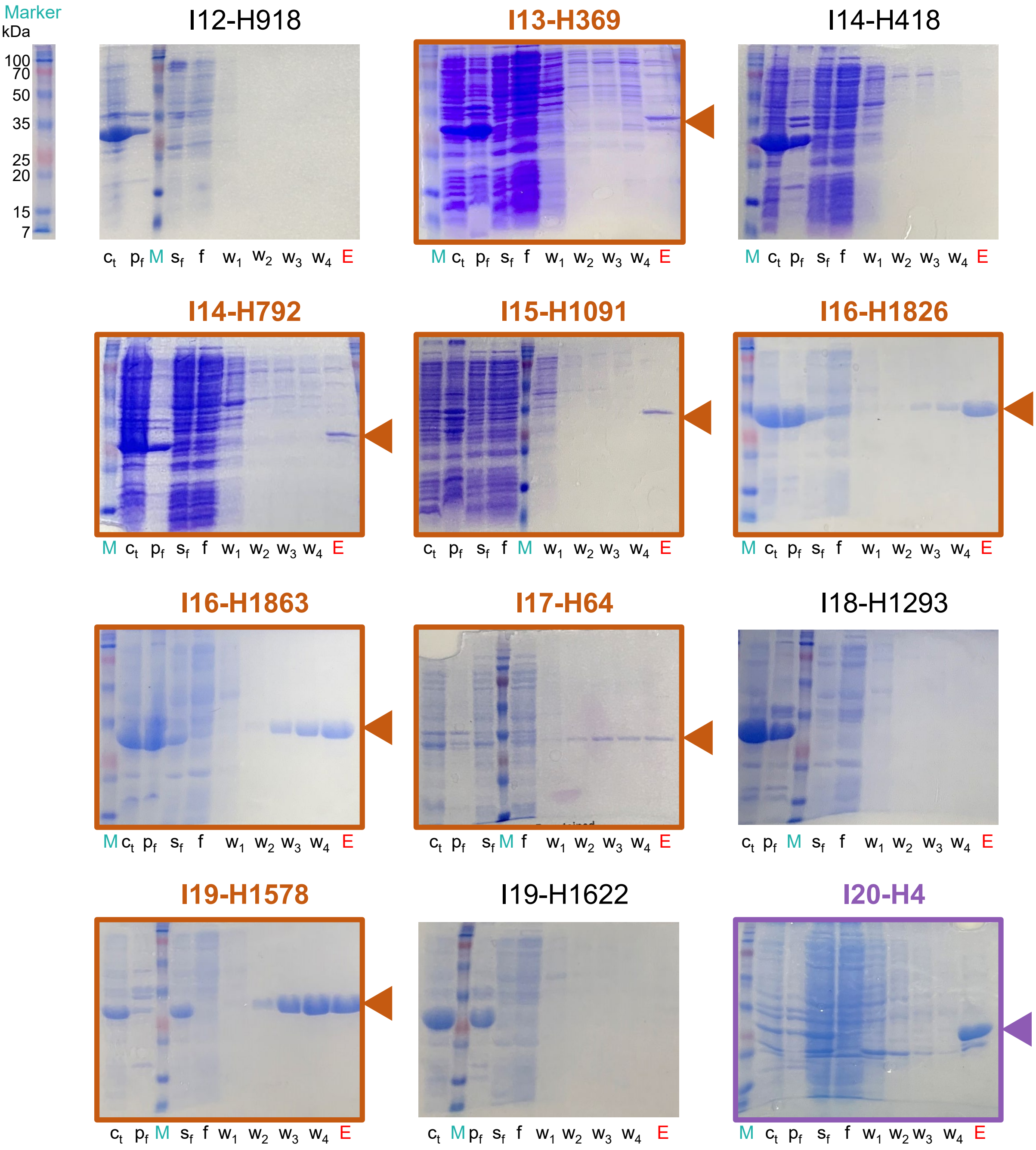


**Fig. S6. Expression and purification of 289 Houses (continued, 10/25).** Abbreviations: Ct, cell total lysate; Pf, pellet fraction; Sf, soluble fraction; W1-4, wash; E, elution; M, marker.

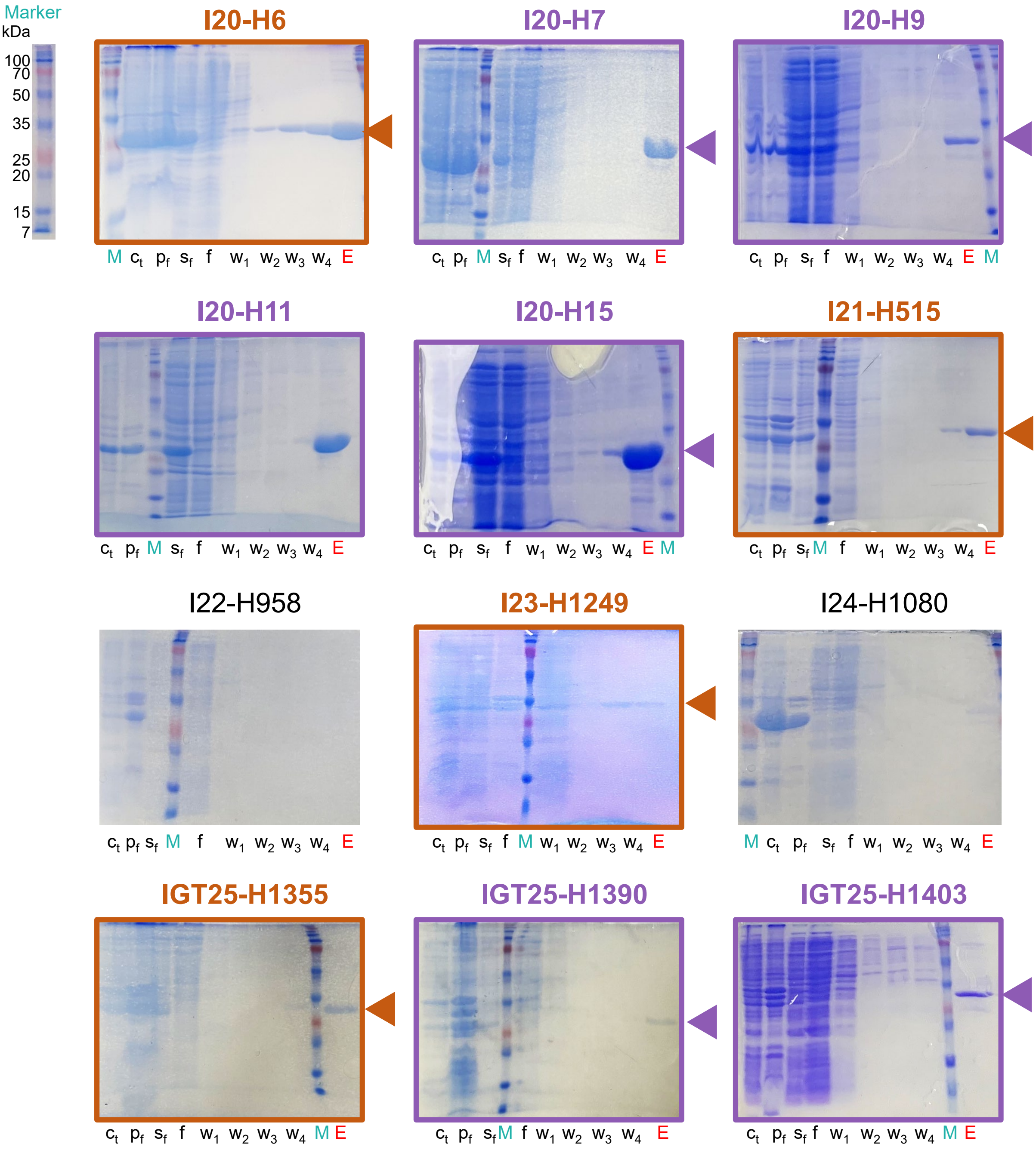


**Fig. S6. Expression and purification of 289 Houses (continued, 11/25).** Abbreviations: Ct, cell total lysate; Pf, pellet fraction; Sf, soluble fraction; W1-4, wash; E, elution; M, marker.

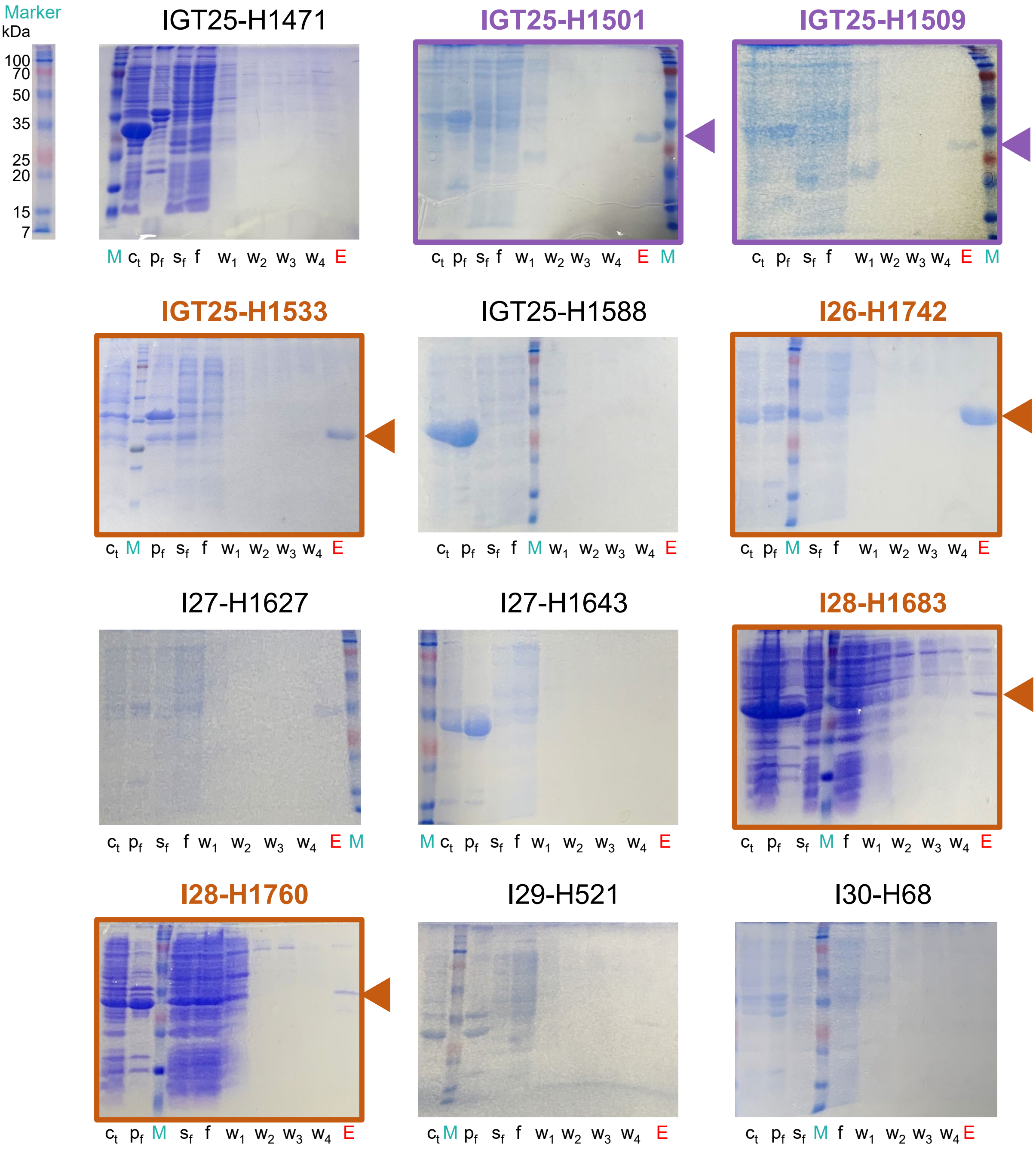


**Fig. S6. Expression and purification of 289 Houses (continued, 12/25).** Abbreviations: Ct, cell total lysate; Pf, pellet fraction; Sf, soluble fraction; W1-4, wash; E, elution; M, marker.

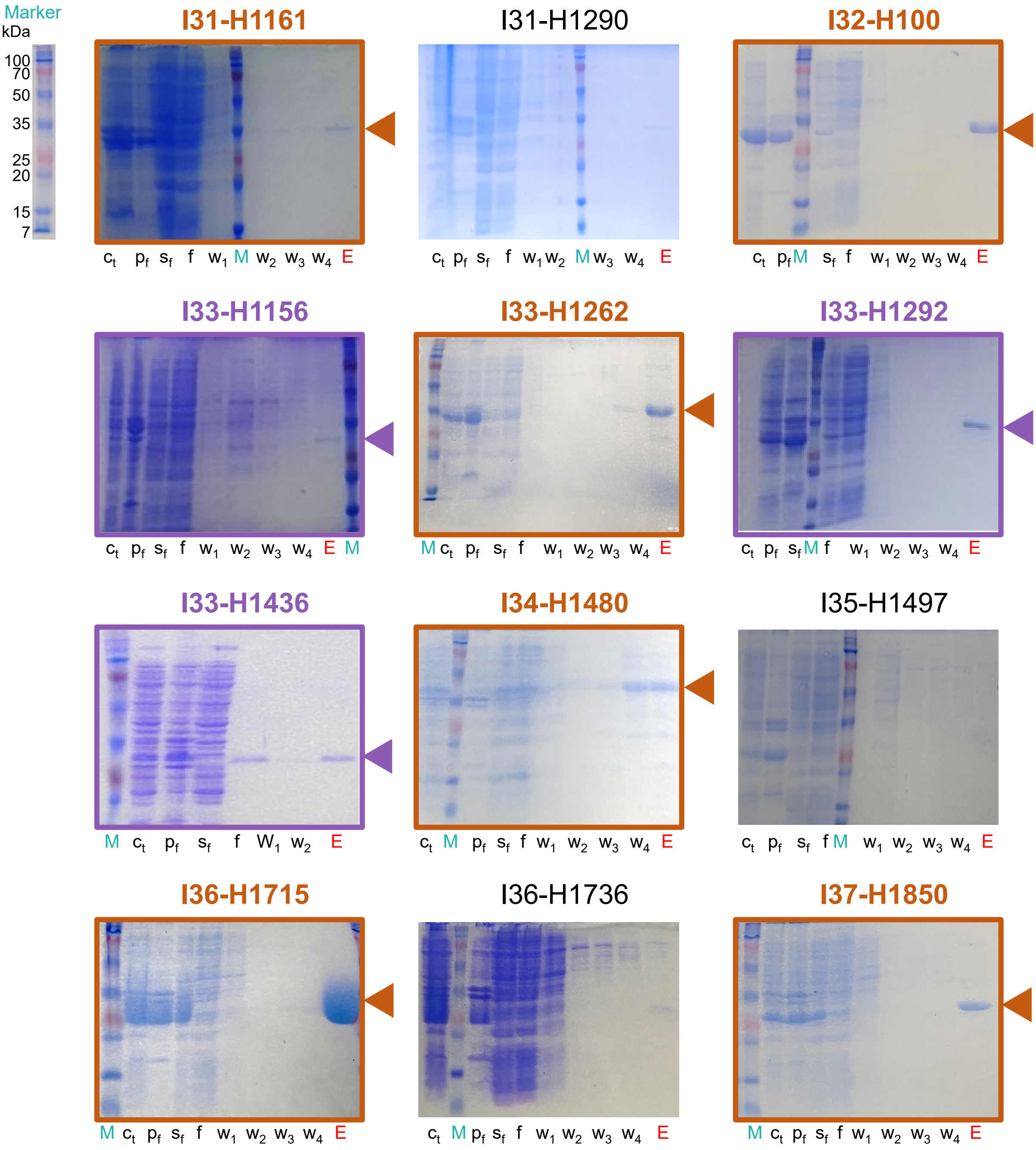


**Fig. S6. Expression and purification of 289 Houses (continued, 13/25).** Abbreviations: Ct, cell total lysate; Pf, pellet fraction; Sf, soluble fraction; W1-4, wash; E, elution; M, marker.

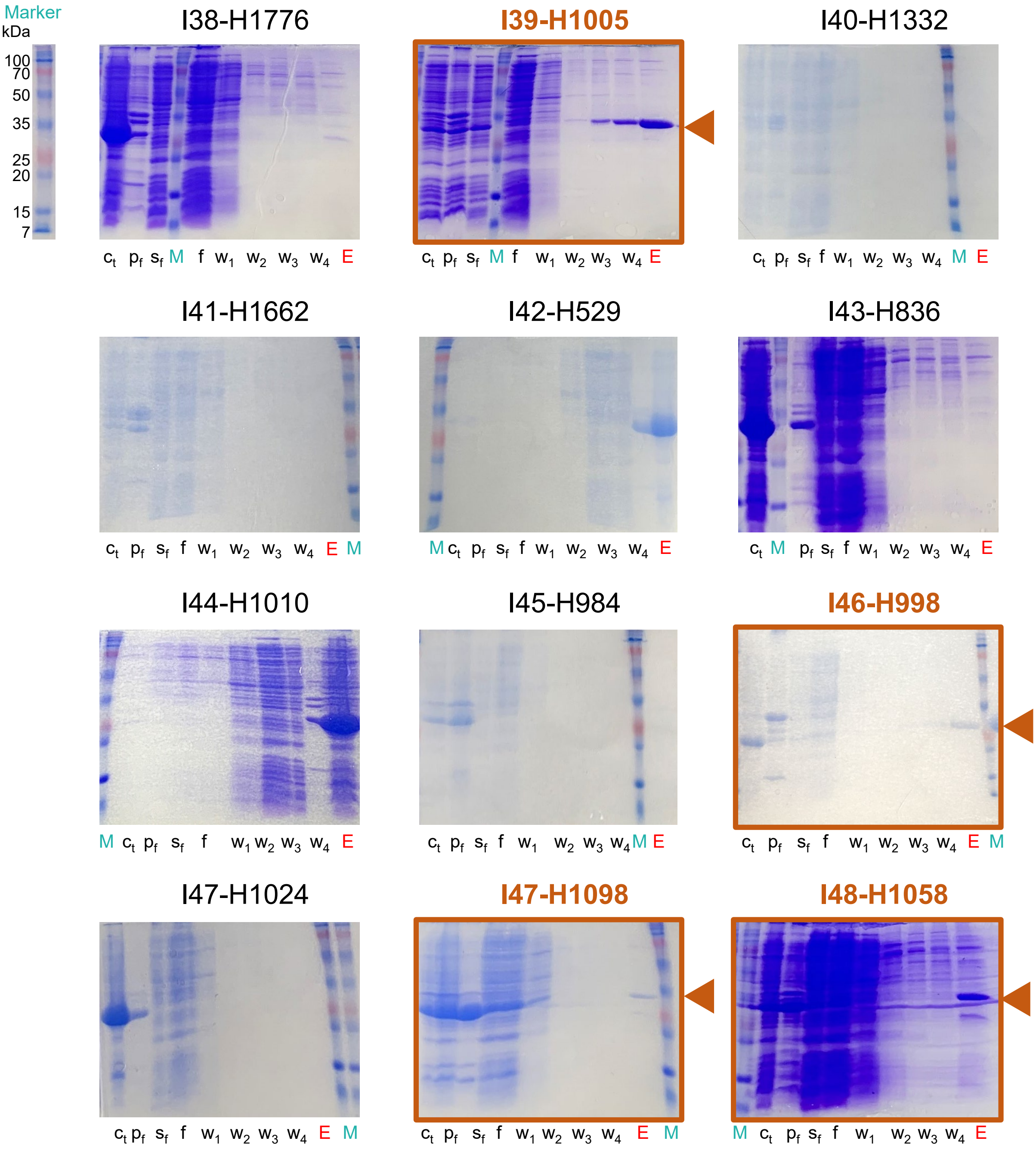


**Fig. S6. Expression and purification of 289 Houses (continued, 14/25).** Abbreviations: Ct, cell total lysate; Pf, pellet fraction; Sf, soluble fraction; W1-4, wash; E, elution; M, marker.

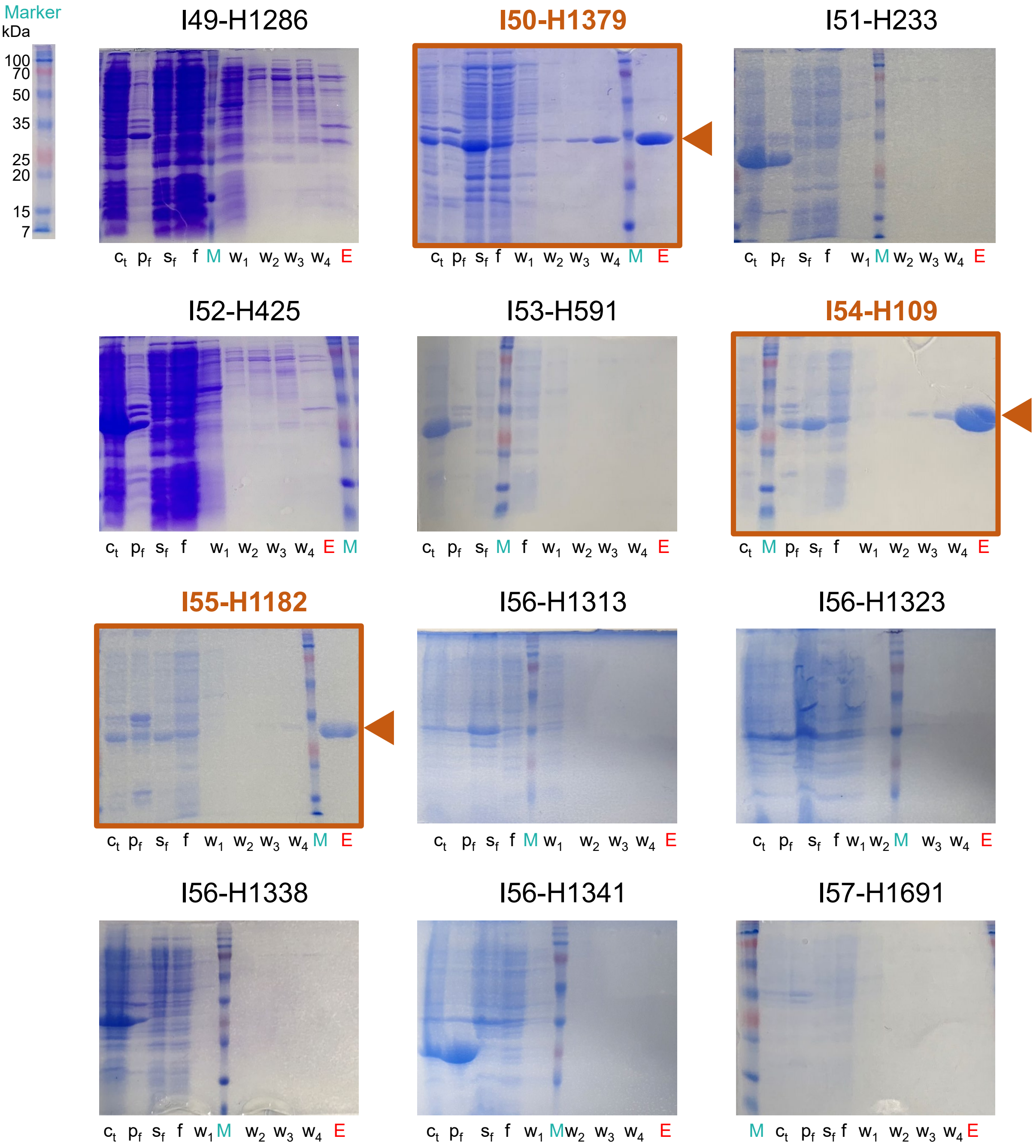


**Fig. S6. Expression and purification of 289 Houses (continued, 15/25).** Abbreviations: Ct, cell total lysate; Pf, pellet fraction; Sf, soluble fraction; W1-4, wash; E, elution; M, marker.

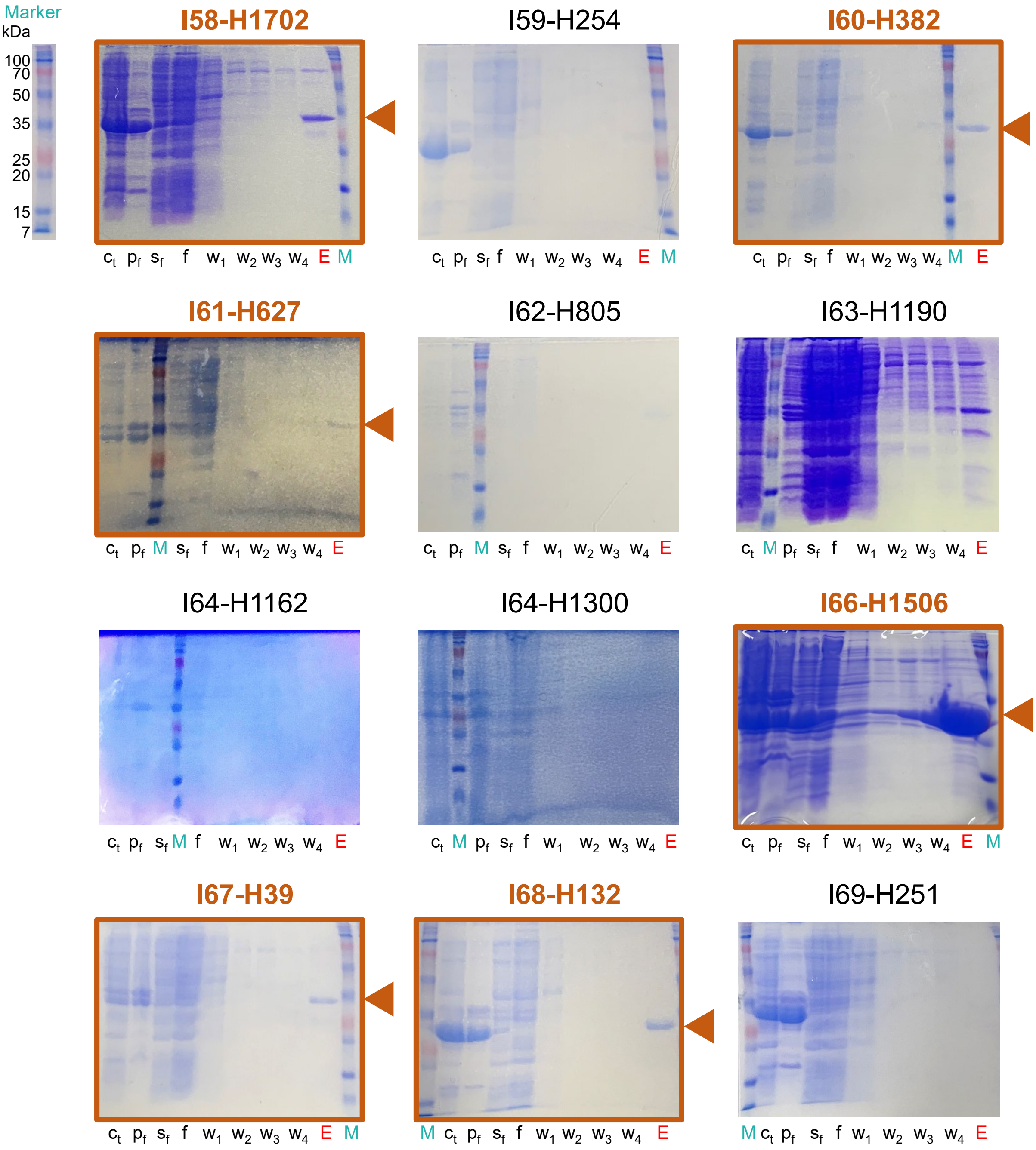


**Fig. S6. Expression and purification of 289 Houses (continued, 16/25).** Abbreviations: Ct, cell total lysate; Pf, pellet fraction; Sf, soluble fraction; W1-4, wash; E, elution; M, marker.

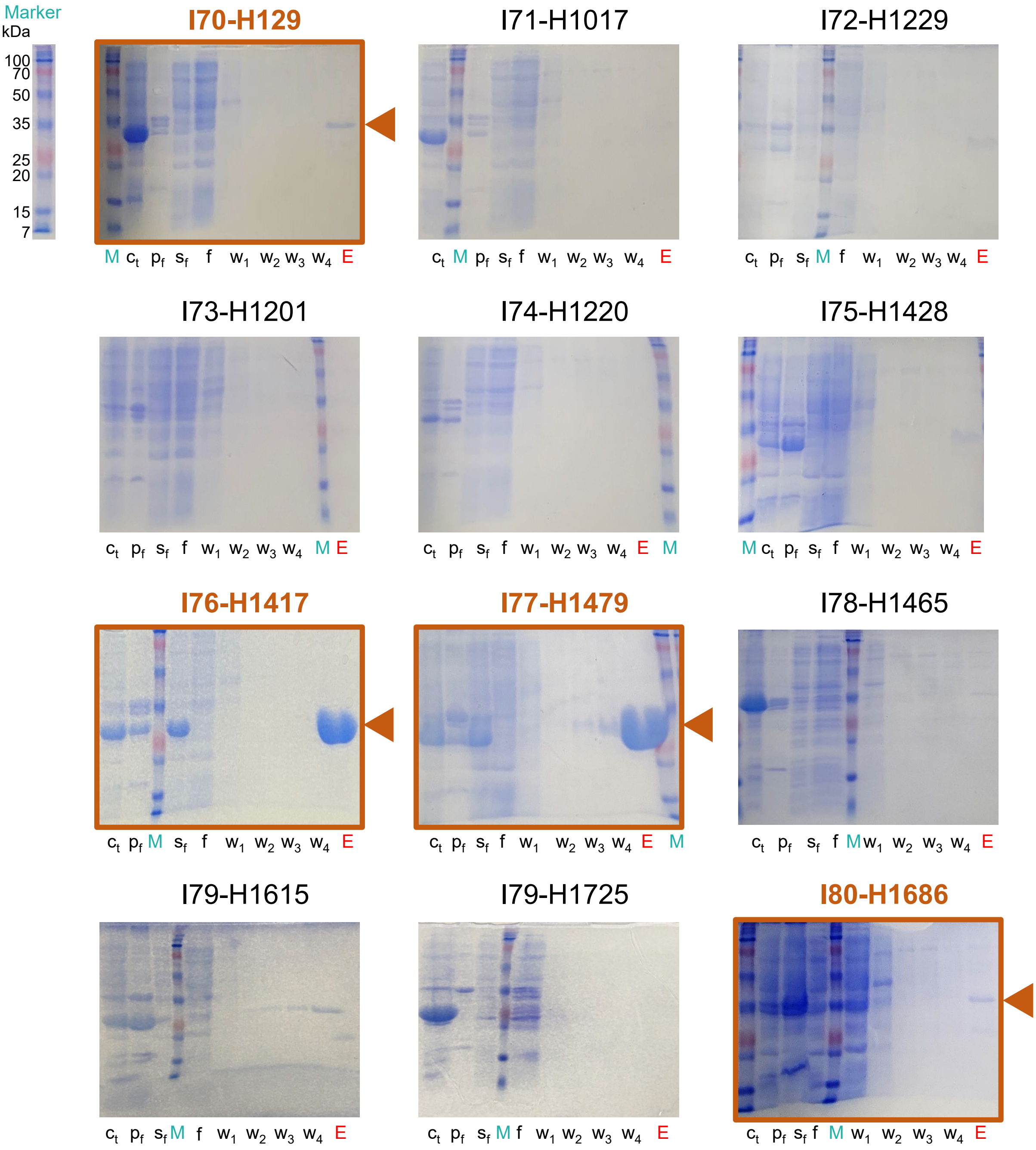


**Fig. S6. Expression and purification of 289 Houses (continued, 17/25).** Abbreviations: Ct, cell total lysate; Pf, pellet fraction; Sf, soluble fraction; W1-4, wash; E, elution; M, marker.

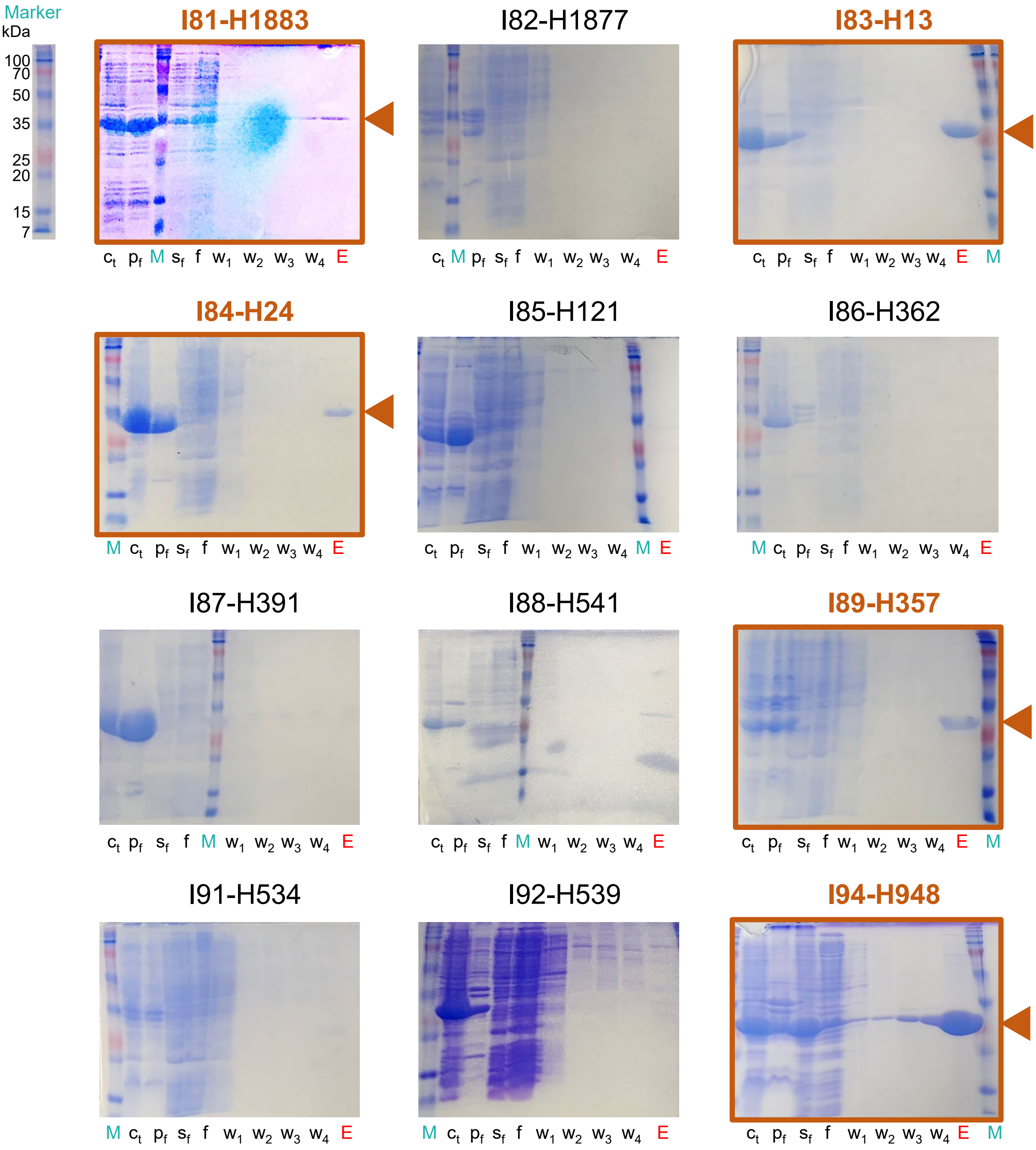


**Fig. S6. Expression and purification of 289 Houses (continued, 18/25).** Abbreviations: Ct, cell total lysate; Pf, pellet fraction; Sf, soluble fraction; W1-4, wash; E, elution; M, marker.

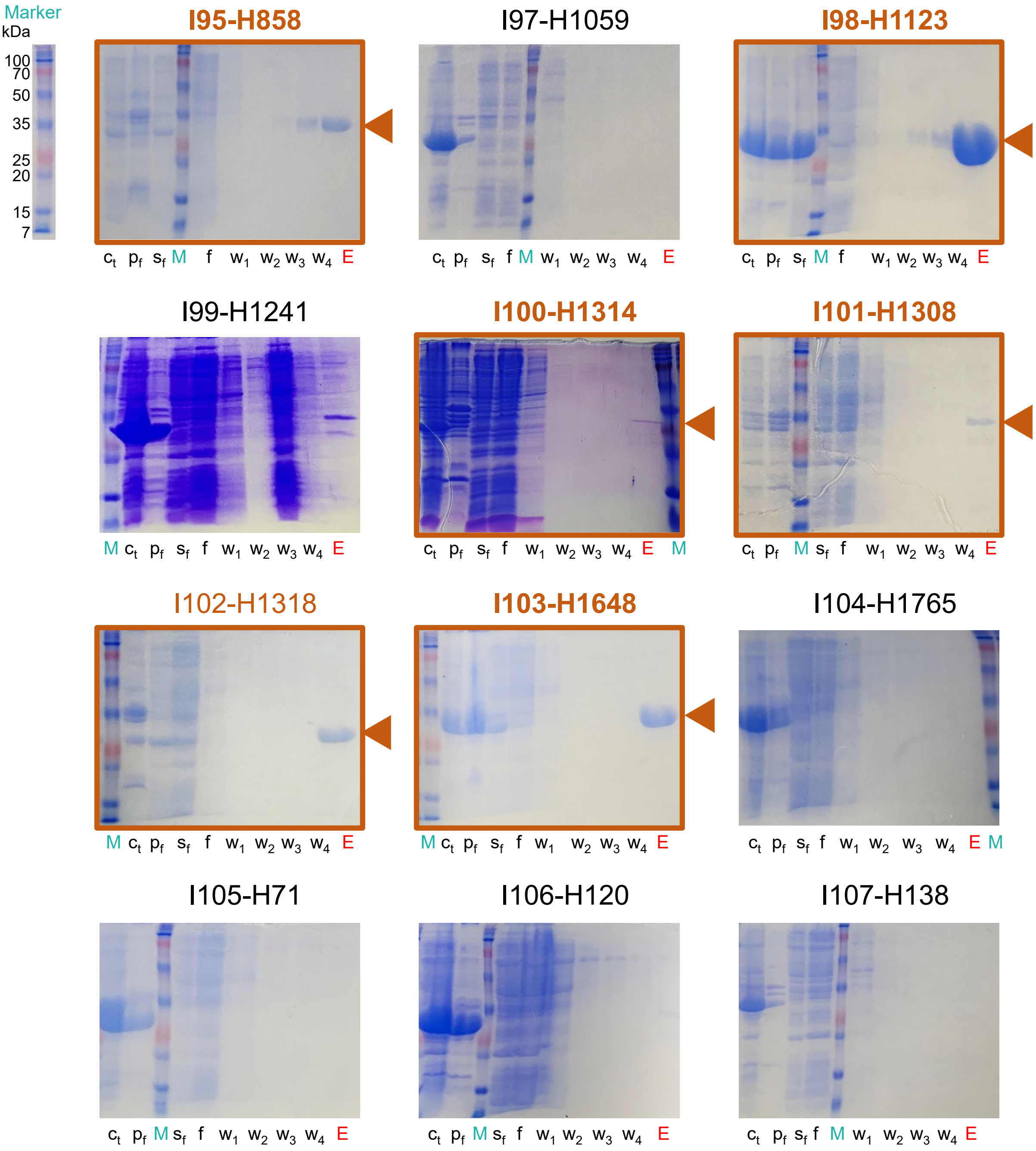


**Fig. S6. Expression and purification of 289 Houses (continued, 19/25).** Abbreviations: Ct, cell total lysate; Pf, pellet fraction; Sf, soluble fraction; W1-4, wash; E, elution; M, marker.

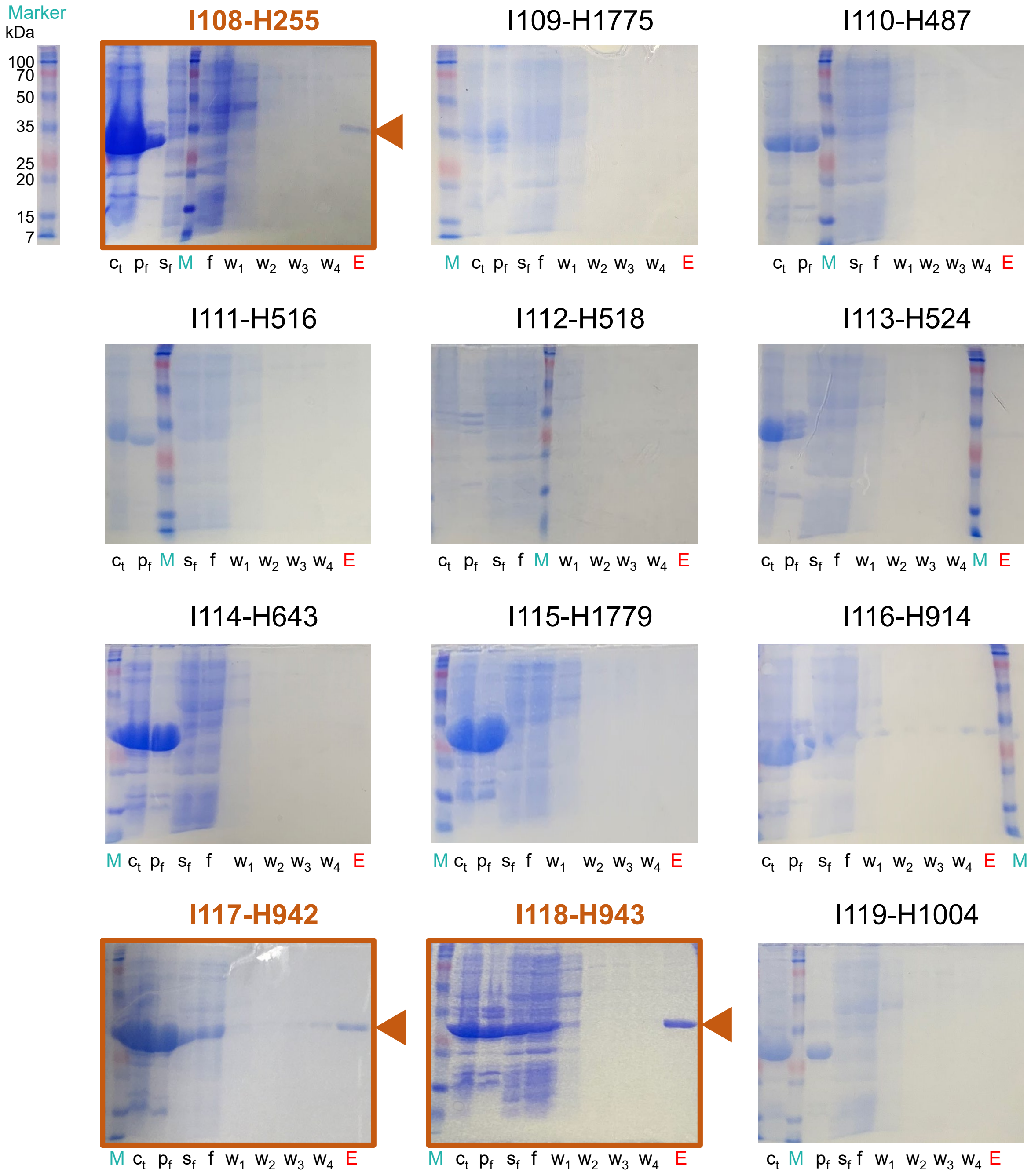


**Fig. S6. Expression and purification of 289 Houses (continued, 20/25).** Abbreviations: Ct, cell total lysate; Pf, pellet fraction; Sf, soluble fraction; W1-4, wash; E, elution; M, marker.

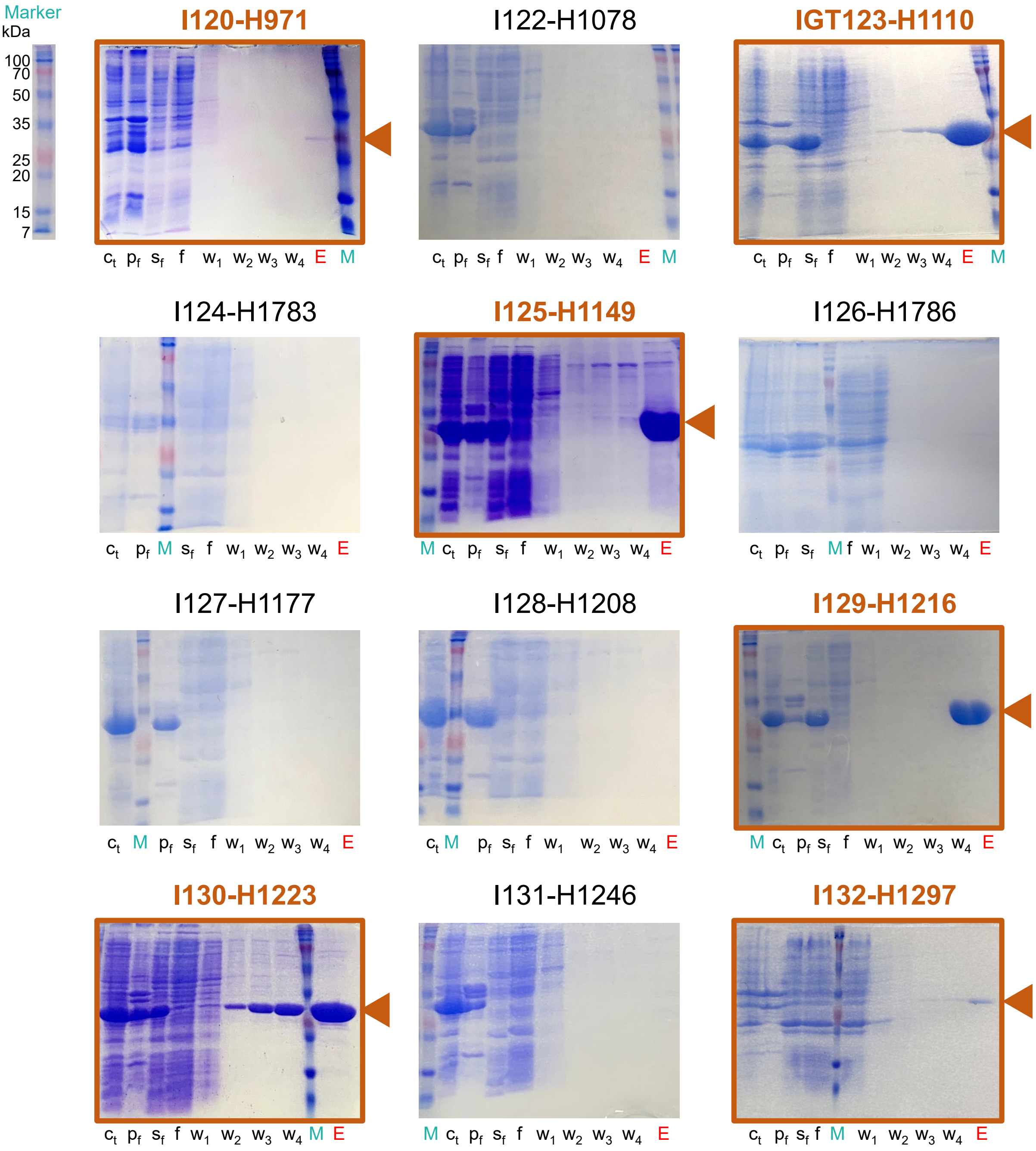


**Fig. S6. Expression and purification of 289 Houses (continued, 21/25).** Abbreviations: Ct, cell total lysate; Pf, pellet fraction; Sf, soluble fraction; W1-4, wash; E, elution; M, marker.

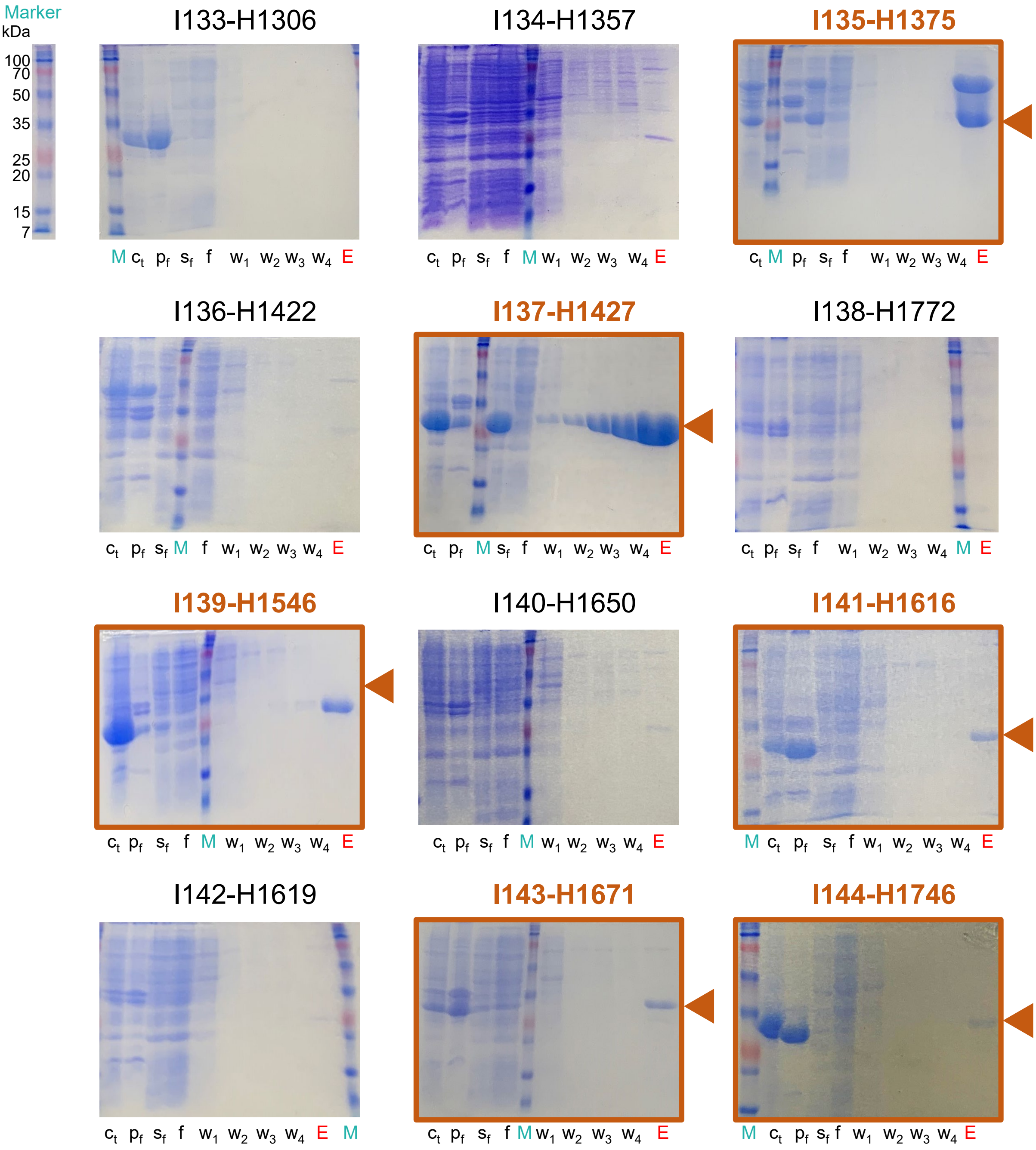


**Fig. S6. Expression and purification of 289 Houses (continued, 22/25).** Abbreviations: Ct, cell total lysate; Pf, pellet fraction; Sf, soluble fraction; W1-4, wash; E, elution; M, marker.

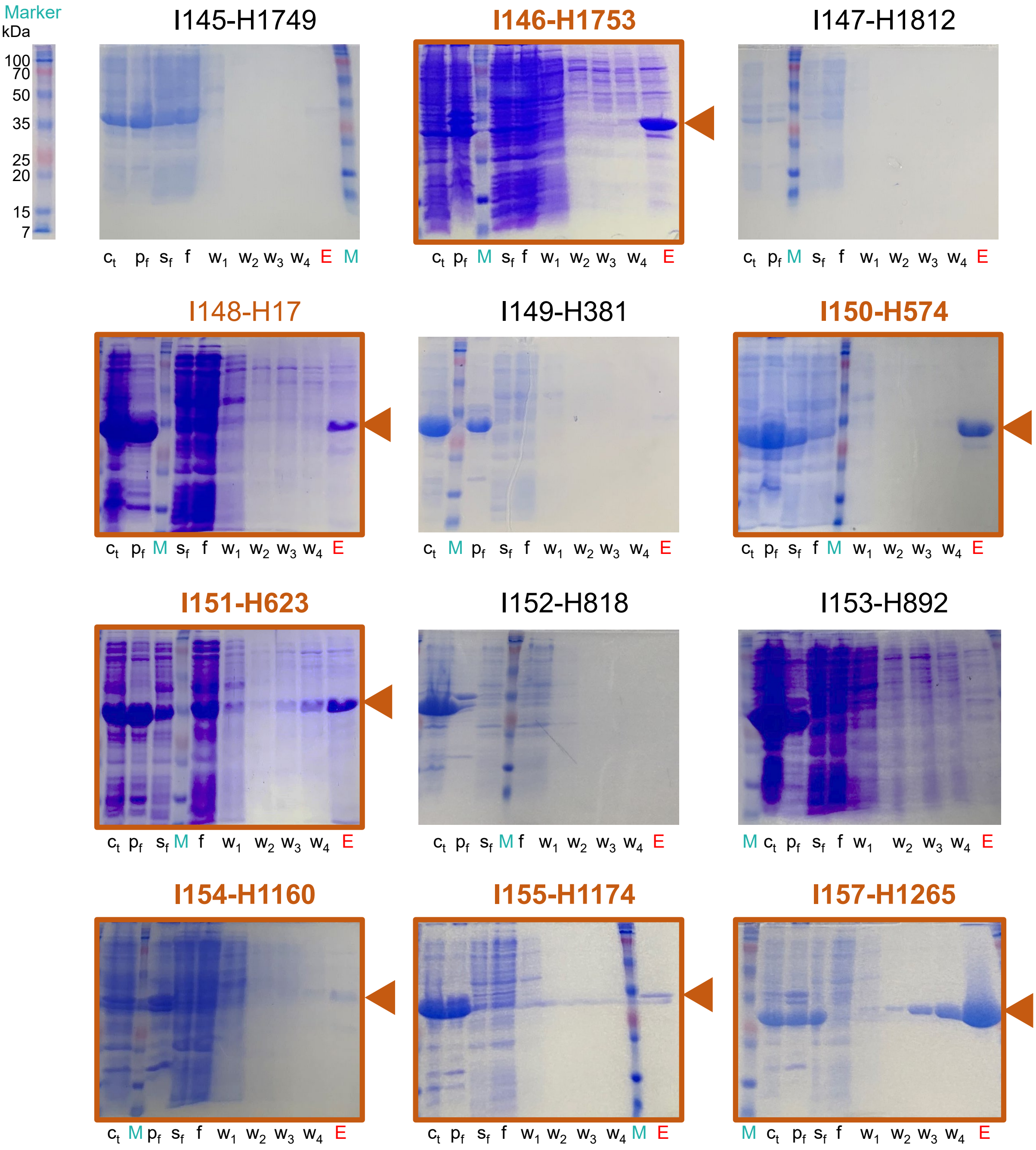


**Fig. S6. Expression and purification of 289 Houses (continued, 23/25).** Abbreviations: Ct, cell total lysate; Pf, pellet fraction; Sf, soluble fraction; W1-4, wash; E, elution; M, marker.

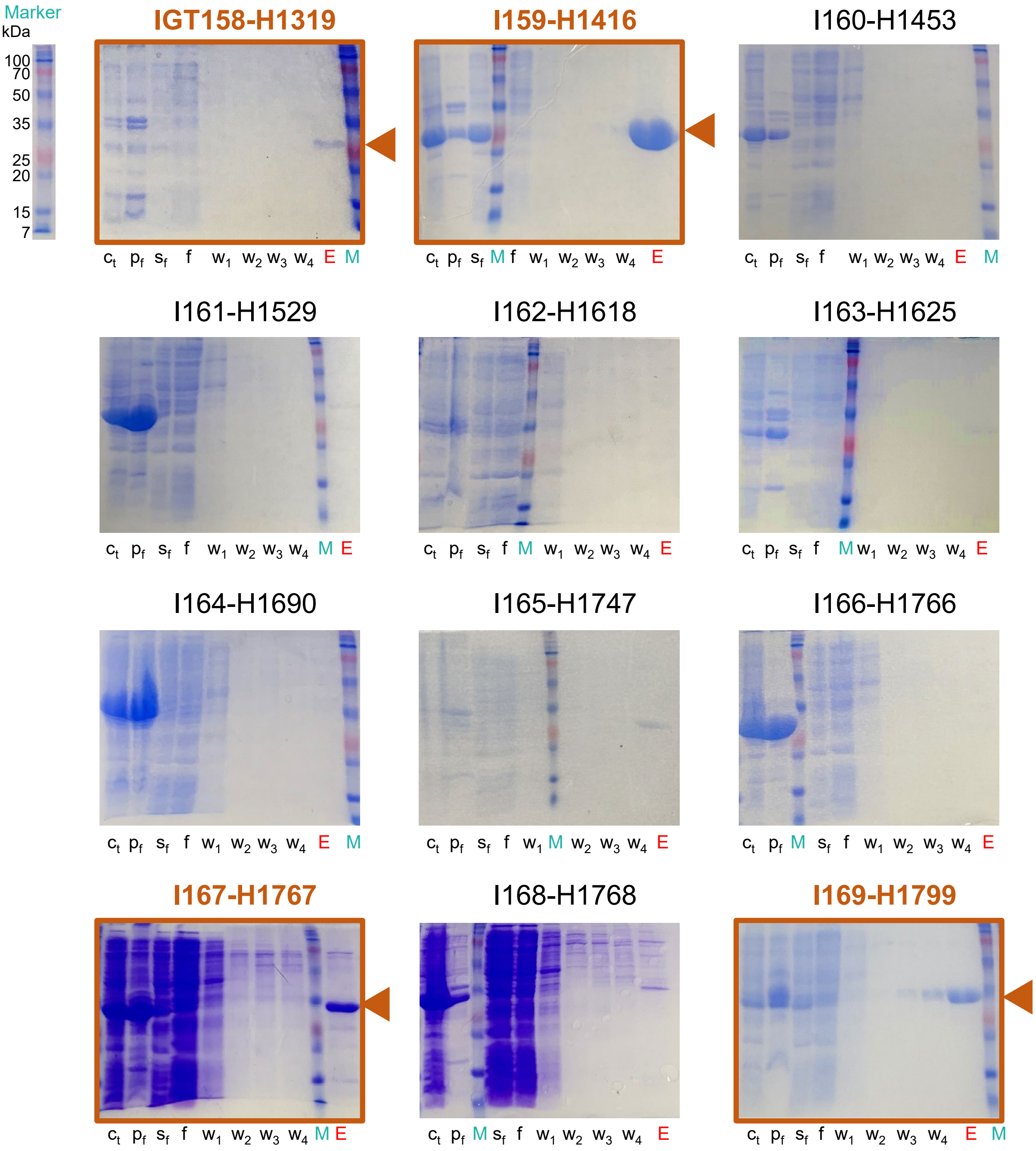


**Fig. S6. Expression and purification of 289 Houses (continued, 24/25).** Abbreviations: Ct, cell total lysate; Pf, pellet fraction; Sf, soluble fraction; W1-4, wash; E, elution; M, marker.

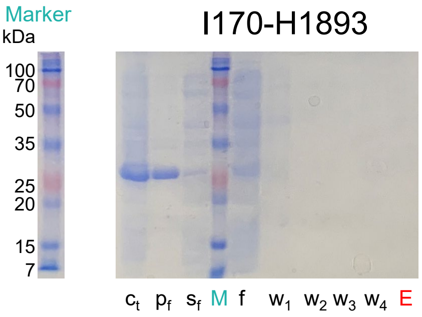


**Fig. S6. Expression and purification of 289 Houses (25/25).** Abbreviations: Ct, cell total lysate; Pf, pellet fraction; Sf, soluble fraction; W1-4, wash; E, elution; M, marker.

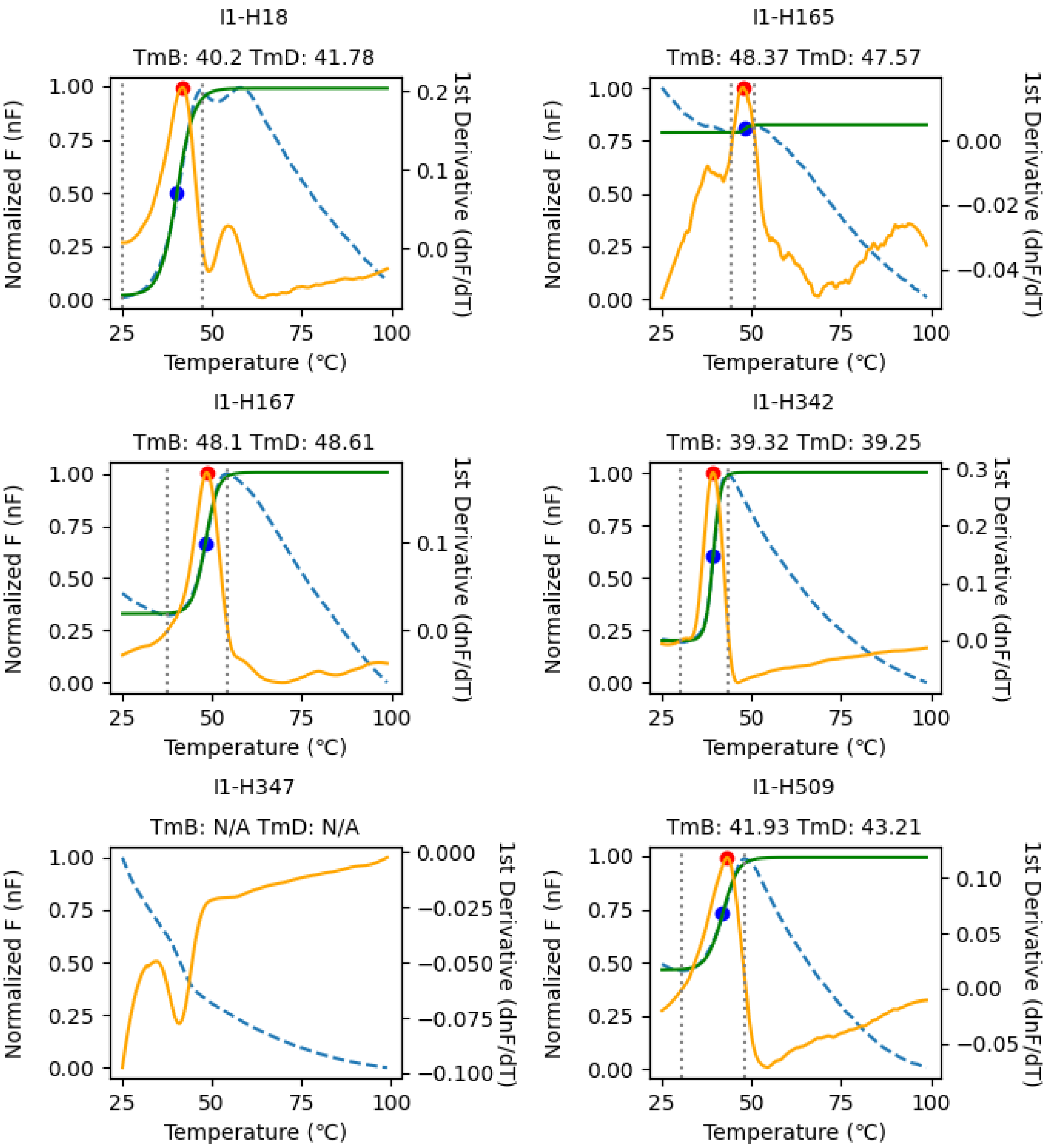


**Fig. S7. Melting temperature (Tm) determination of 158 Houses through differential scanning fluorimetry (continued, 1/27).** The normalized fluorescence (nF) and its 1st derivative is shown with a blue dashed and yellow solid lines, respectively. The Boltzmann curve fitted to the nF is shown with a green solid line. The determined $Tm_B$ and $Tm_D$ is shown with blue and red circles, respectively, and labeled.

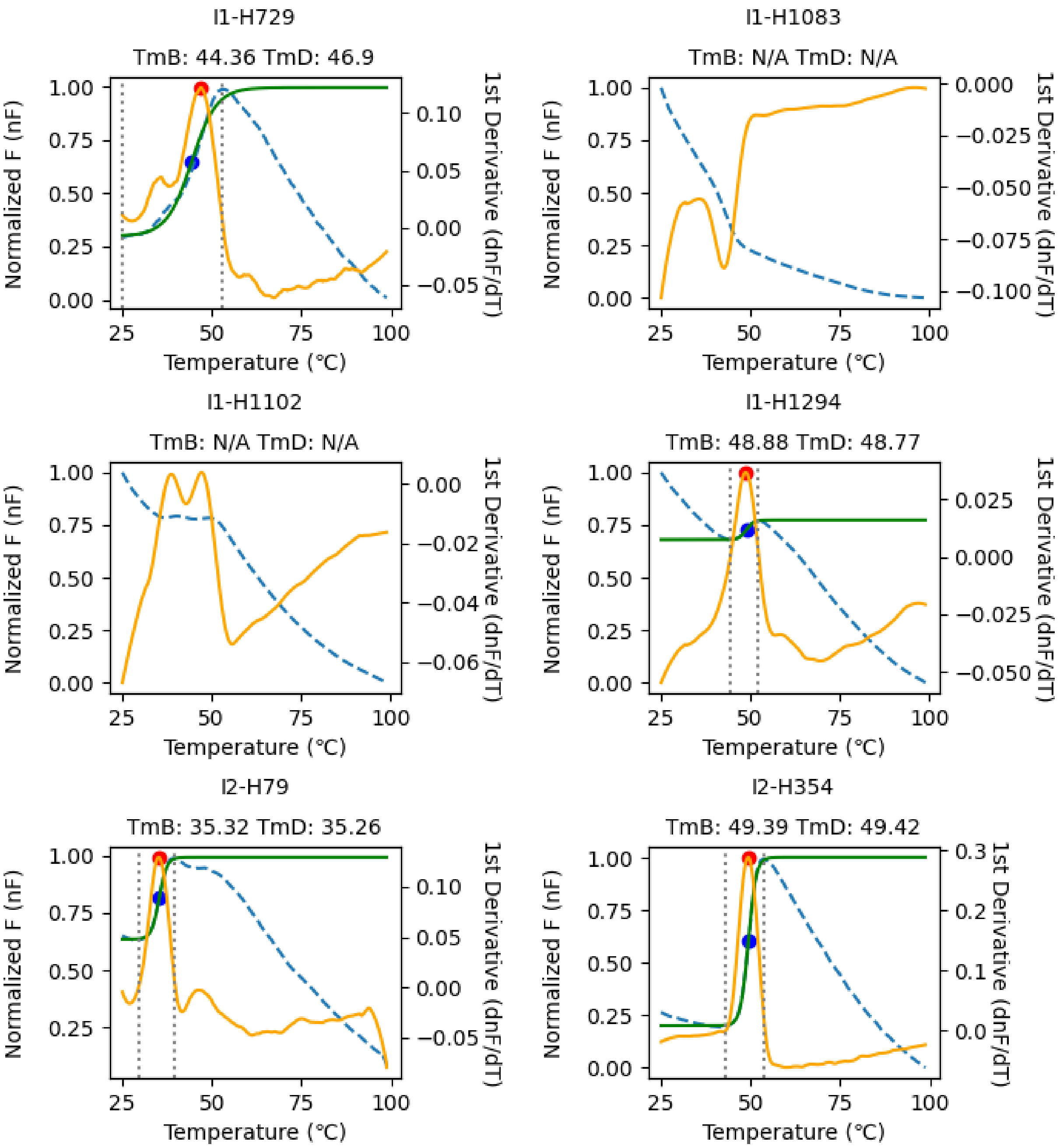


**Fig. S7. Melting temperature (Tm) determination of 158 Houses through differential scanning fluorimetry (continued, 2/27).** The normalized fluorescence (nF) and its 1st derivative is shown with a blue dashed and yellow solid lines, respectively. The Boltzmann curve fitted to the nF is shown with a green solid line. The determined $Tm_B$ and $Tm_D$ is shown with blue and red circles, respectively, and labeled.

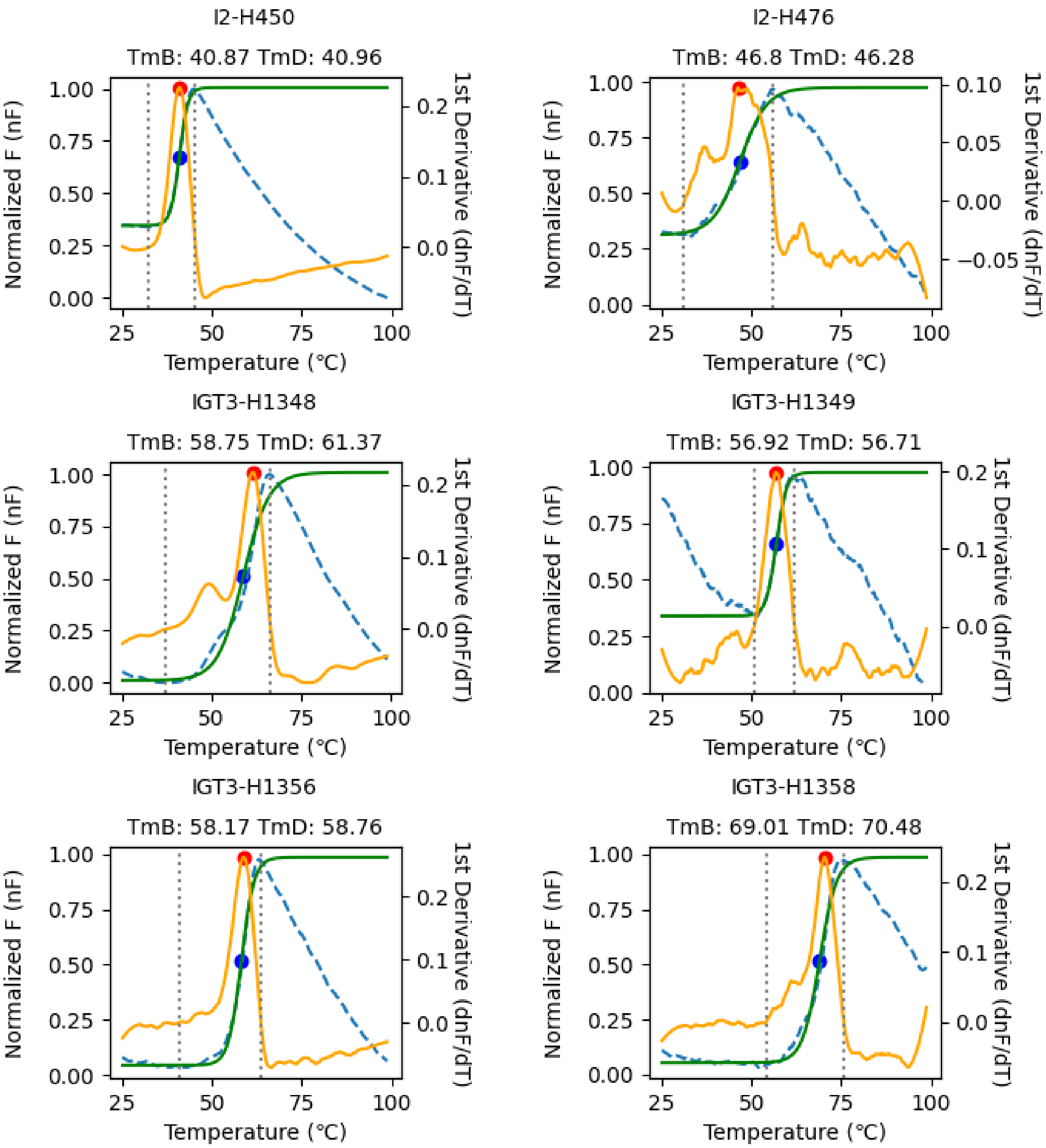


**Fig. S7. Melting temperature (Tm) determination of 158 Houses through differential scanning fluorimetry (continued, 3/27).** The normalized fluorescence (nF) and its 1$^{st}$ derivative is shown with a blue dashed and yellow solid lines, respectively. The Boltzmann curve fitted to the nF is shown with a green solid line. The determined $Tm_B$ and $Tm_D$ is shown with blue and red circles, respectively, and labeled.

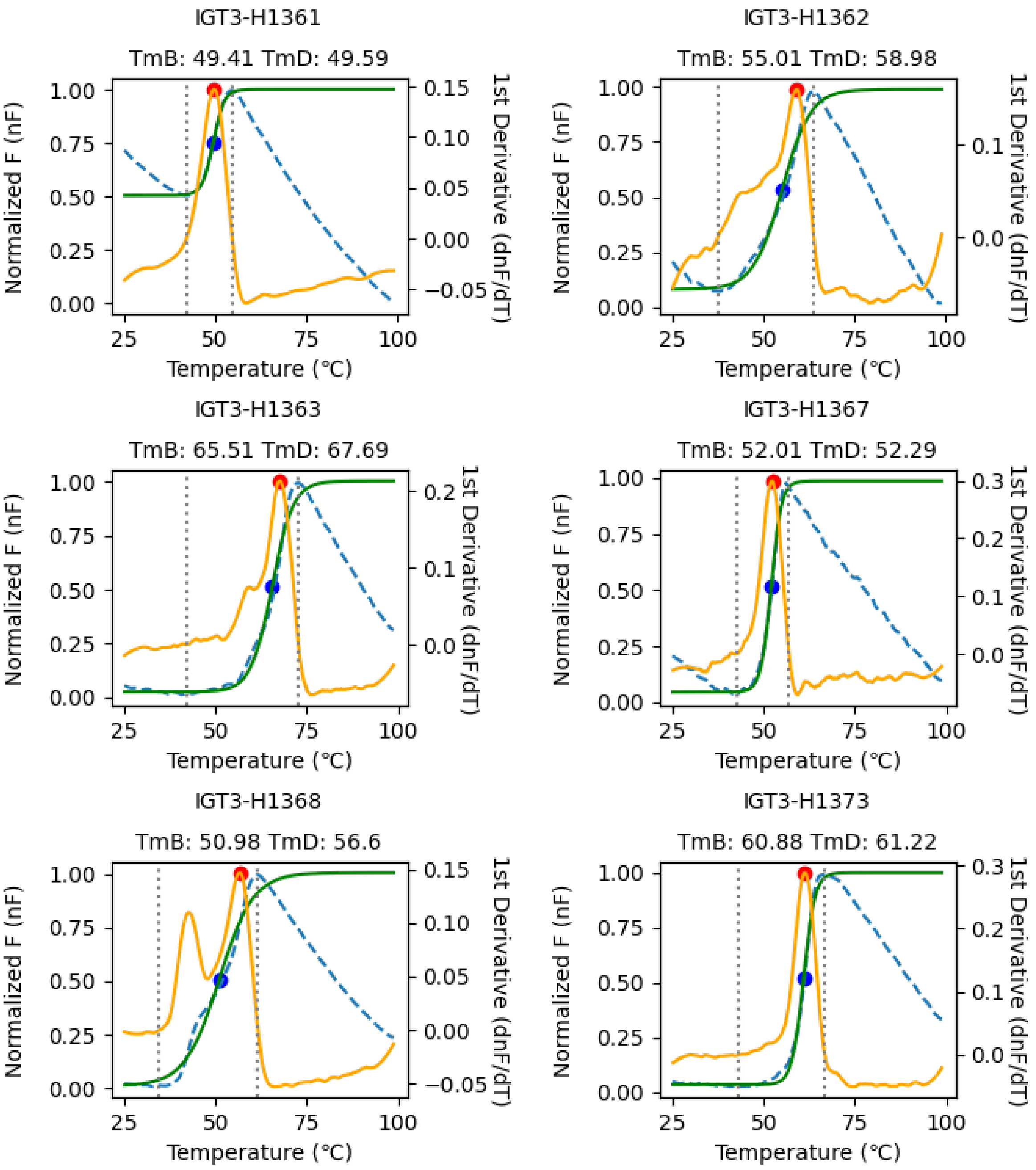


**Fig. S7. Melting temperature (Tm) determination of 158 Houses through differential scanning fluorimetry (continued, 4/27).** The normalized fluorescence (nF) and its 1st derivative is shown with a blue dashed and yellow solid lines, respectively. The Boltzmann curve fitted to the nF is shown with a green solid line. The determined $Tm_B$ and $Tm_D$ is shown with blue and red circles, respectively, and labeled.

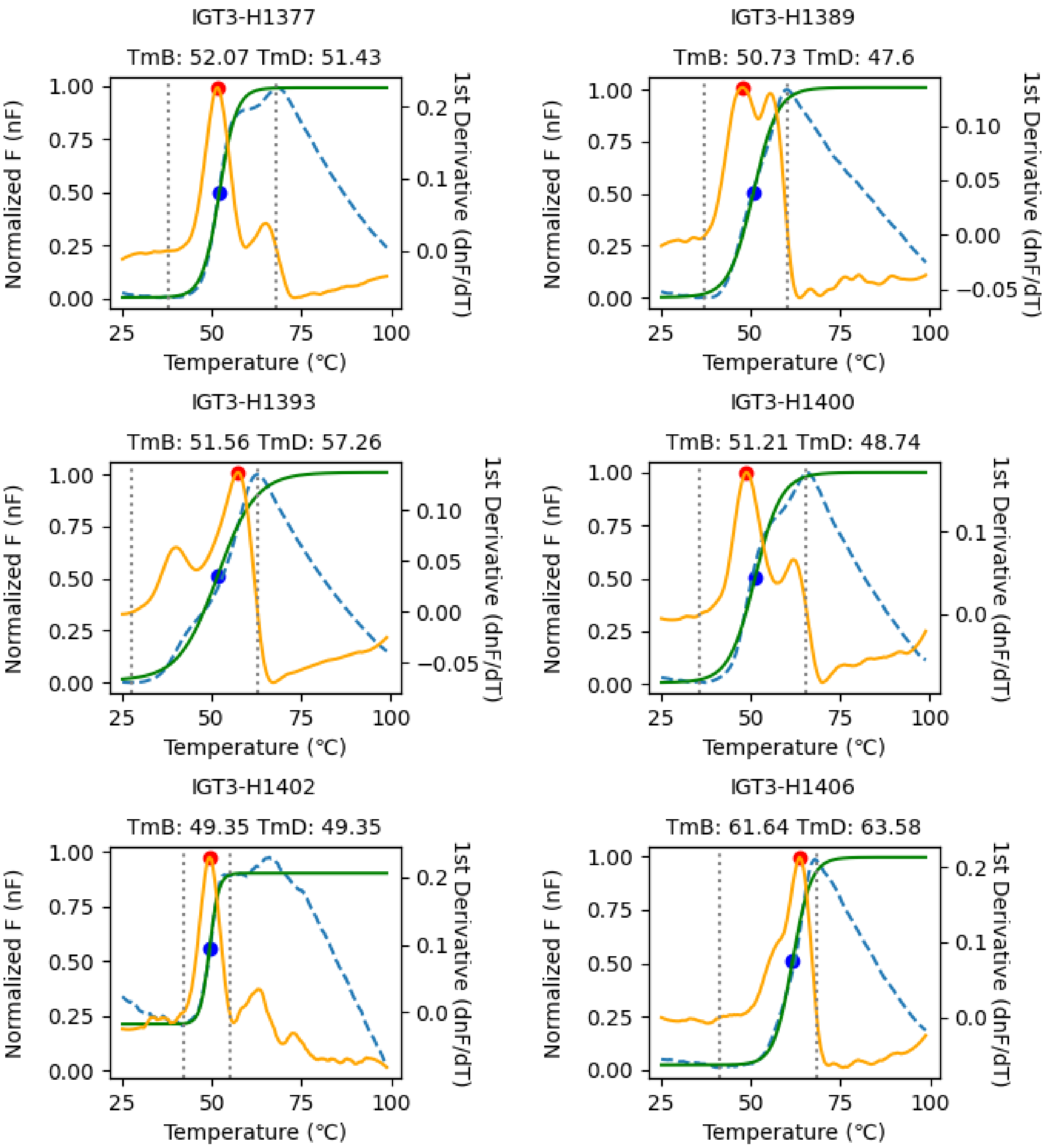


**Fig. S7. Melting temperature (Tm) determination of 158 Houses through differential scanning fluorimetry (continued, 5/27).** The normalized fluorescence (nF) and its 1st derivative is shown with a blue dashed and yellow solid lines, respectively. The Boltzmann curve fitted to the nF is shown with a green solid line. The determined $Tm_B$ and $Tm_D$ is shown with blue and red circles, respectively, and labeled.

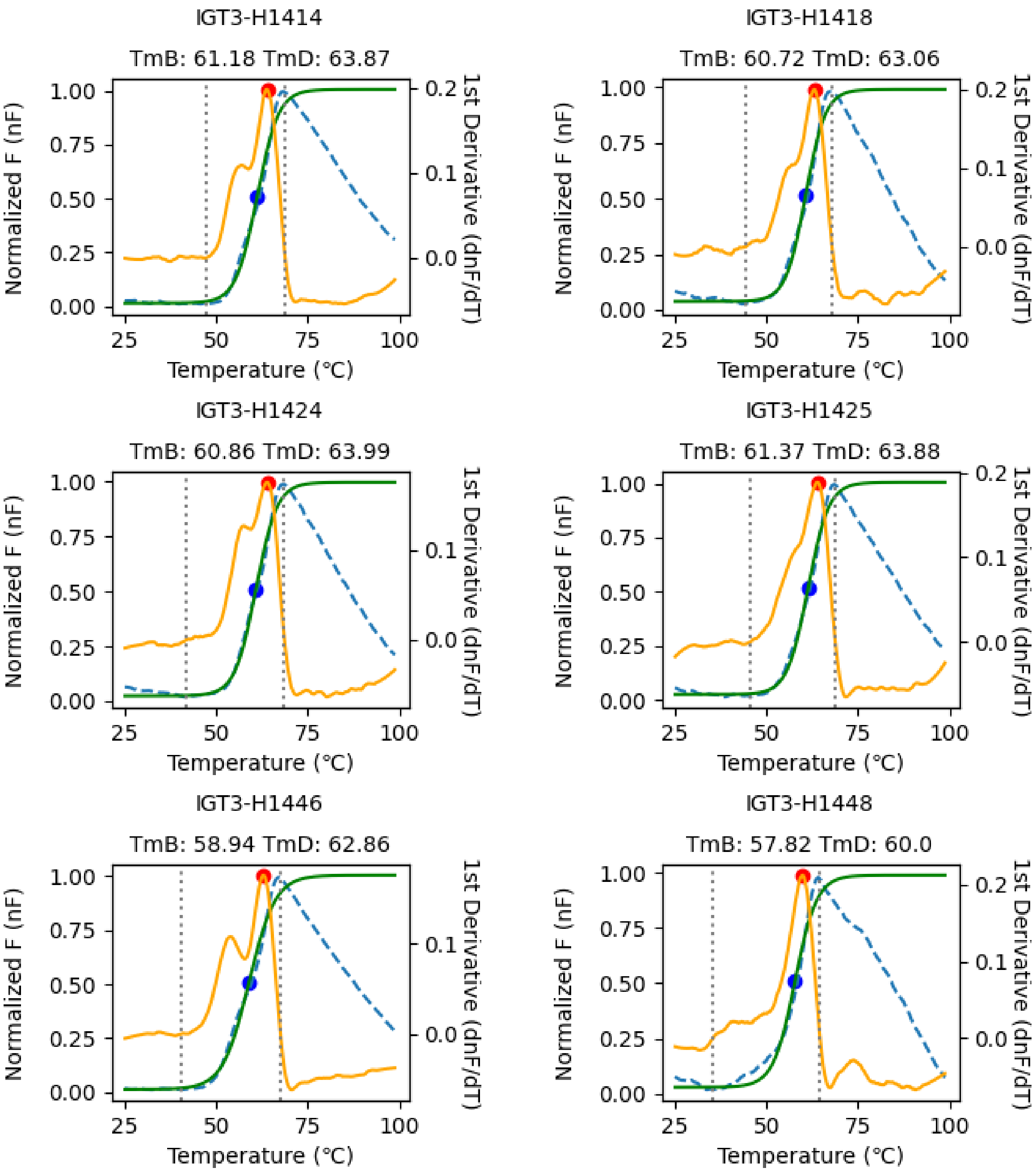


**Fig. S7. Melting temperature (Tm) determination of 158 Houses through differential scanning fluorimetry (continued, 6/27).** The normalized fluorescence (nF) and its $1^{st}$ derivative is shown with a blue dashed and yellow solid lines, respectively. The Boltzmann curve fitted to the nF is shown with a green solid line. The determined $Tm_B$ and $Tm_D$ is shown with blue and red circles, respectively, and labeled.

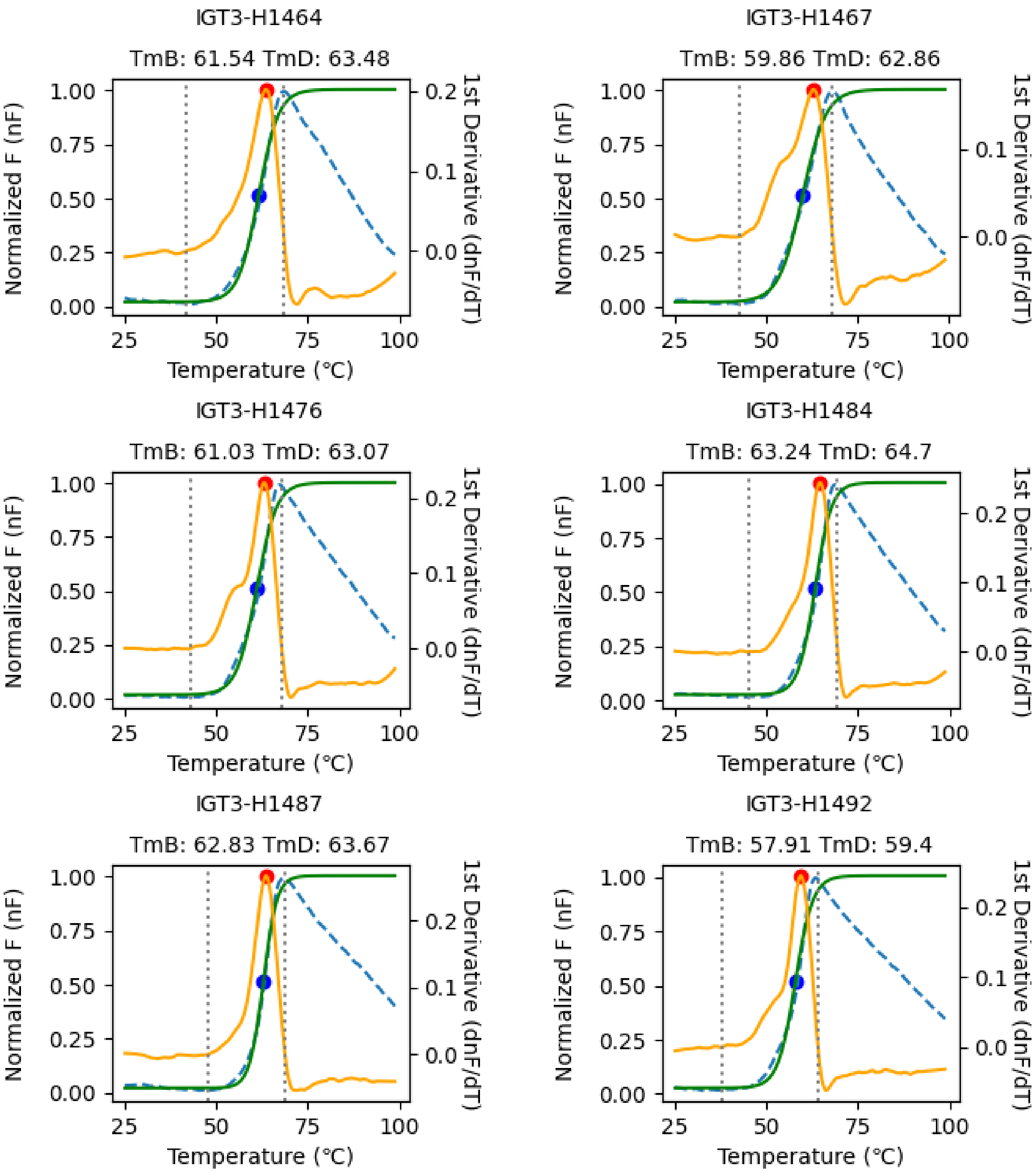


**Fig. S7. Melting temperature (Tm) determination of 158 Houses through differential scanning fluorimetry (continued, 7/27).** The normalized fluorescence (nF) and its 1st derivative is shown with a blue dashed and yellow solid lines, respectively. The Boltzmann curve fitted to the nF is shown with a green solid line. The determined $Tm_B$ and $Tm_D$ is shown with blue and red circles, respectively, and labeled.

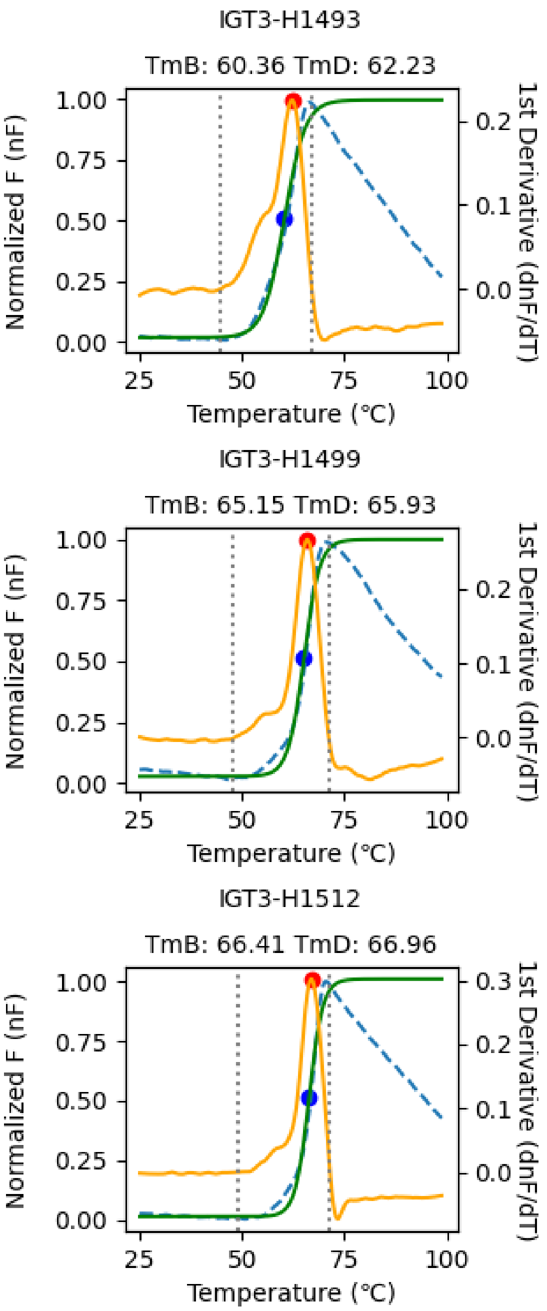


**Fig. S7. Melting temperature (Tm) determination of 158 Houses through differential scanning fluorimetry (continued, 8/27).** The normalized fluorescence (nF) and its 1st derivative is shown with a blue dashed and yellow solid lines, respectively. The Boltzmann curve fitted to the nF is shown with a green solid line. The determined $Tm_B$ and $Tm_D$ is shown with blue and red circles, respectively, and labeled.

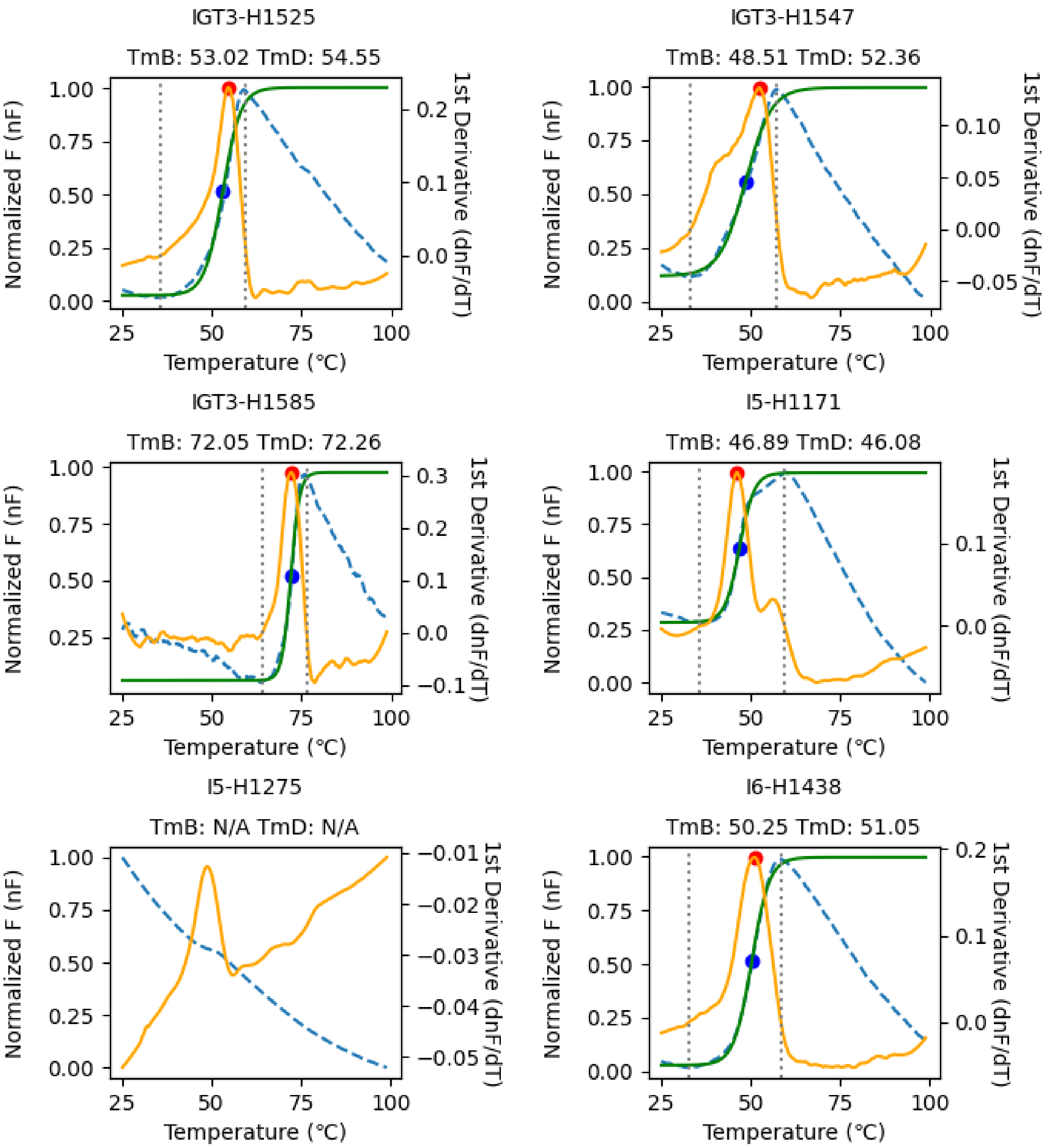


**Fig. S7. Melting temperature (Tm) determination of 158 Houses through differential scanning fluorimetry (continued, 9/27).** The normalized fluorescence (nF) and its 1st derivative is shown with a blue dashed and yellow solid lines, respectively. The Boltzmann curve fitted to the nF is shown with a green solid line. The determined $Tm_B$ and $Tm_D$ is shown with blue and red circles, respectively, and labeled.

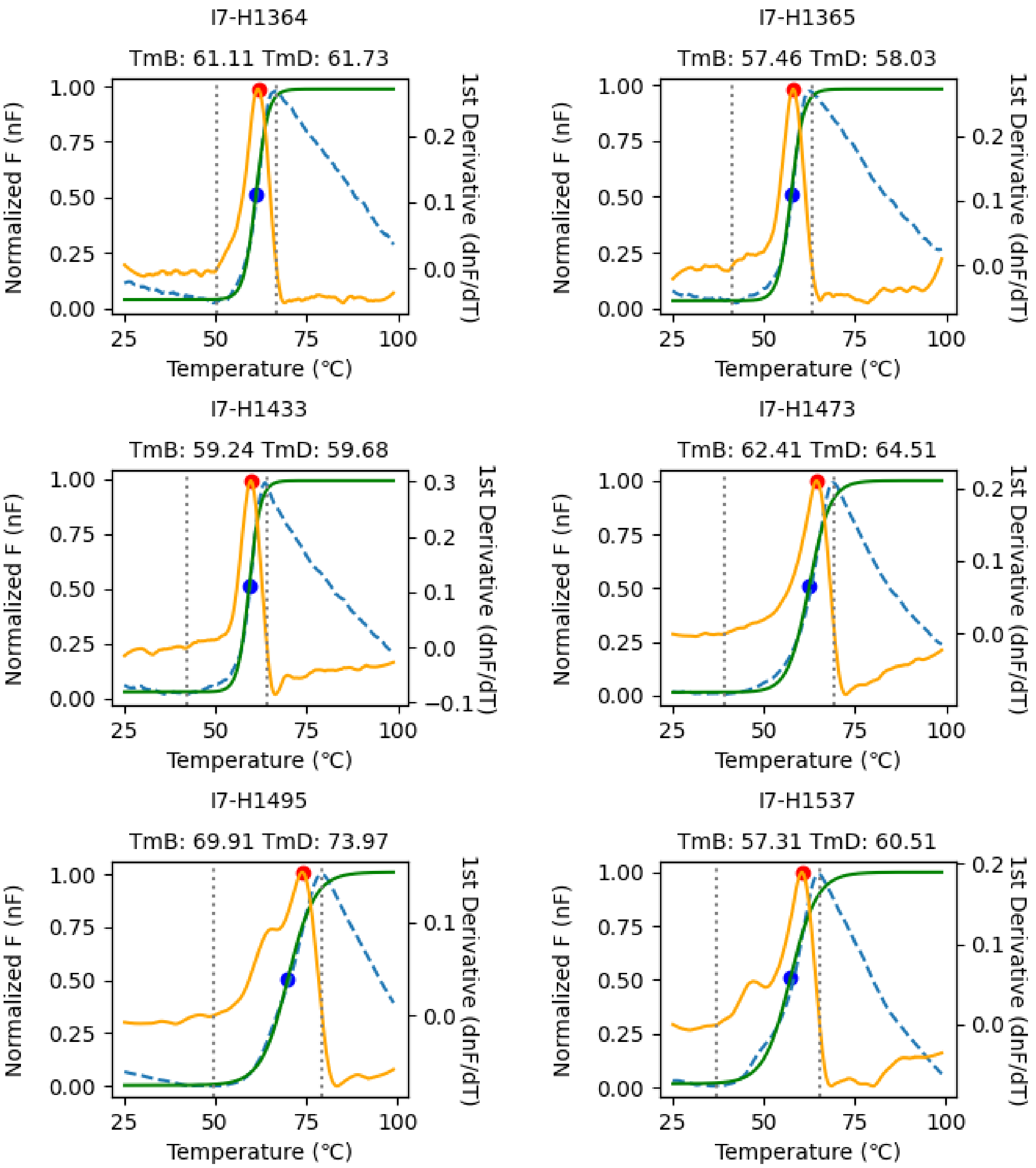


**Fig. S7. Melting temperature (Tm) determination of 158 Houses through differential scanning fluorimetry (continued, 10/27).** The normalized fluorescence (nF) and its 1st derivative is shown with a blue dashed and yellow solid lines, respectively. The Boltzmann curve fitted to the nF is shown with a green solid line. The determined $Tm_B$ and $Tm_D$ is shown with blue and red circles, respectively, and labeled.

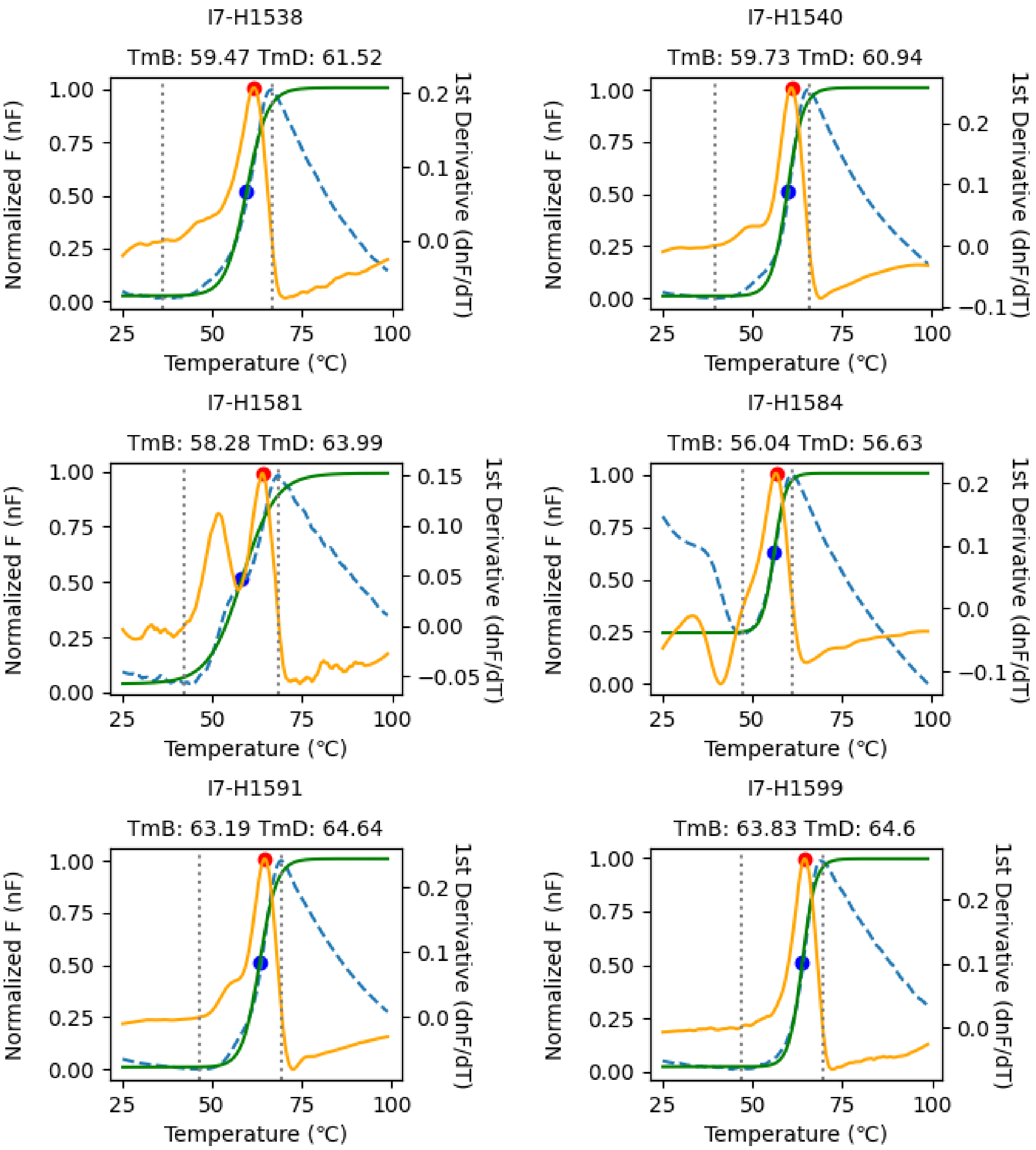


**Fig. S7. Melting temperature (Tm) determination of 158 Houses through differential scanning fluorimetry (continued, 11/27).** The normalized fluorescence (nF) and its 1st derivative is shown with a blue dashed and yellow solid lines, respectively. The Boltzmann curve fitted to the nF is shown with a green solid line. The determined $Tm_B$ and $Tm_D$ is shown with blue and red circles, respectively, and labeled.

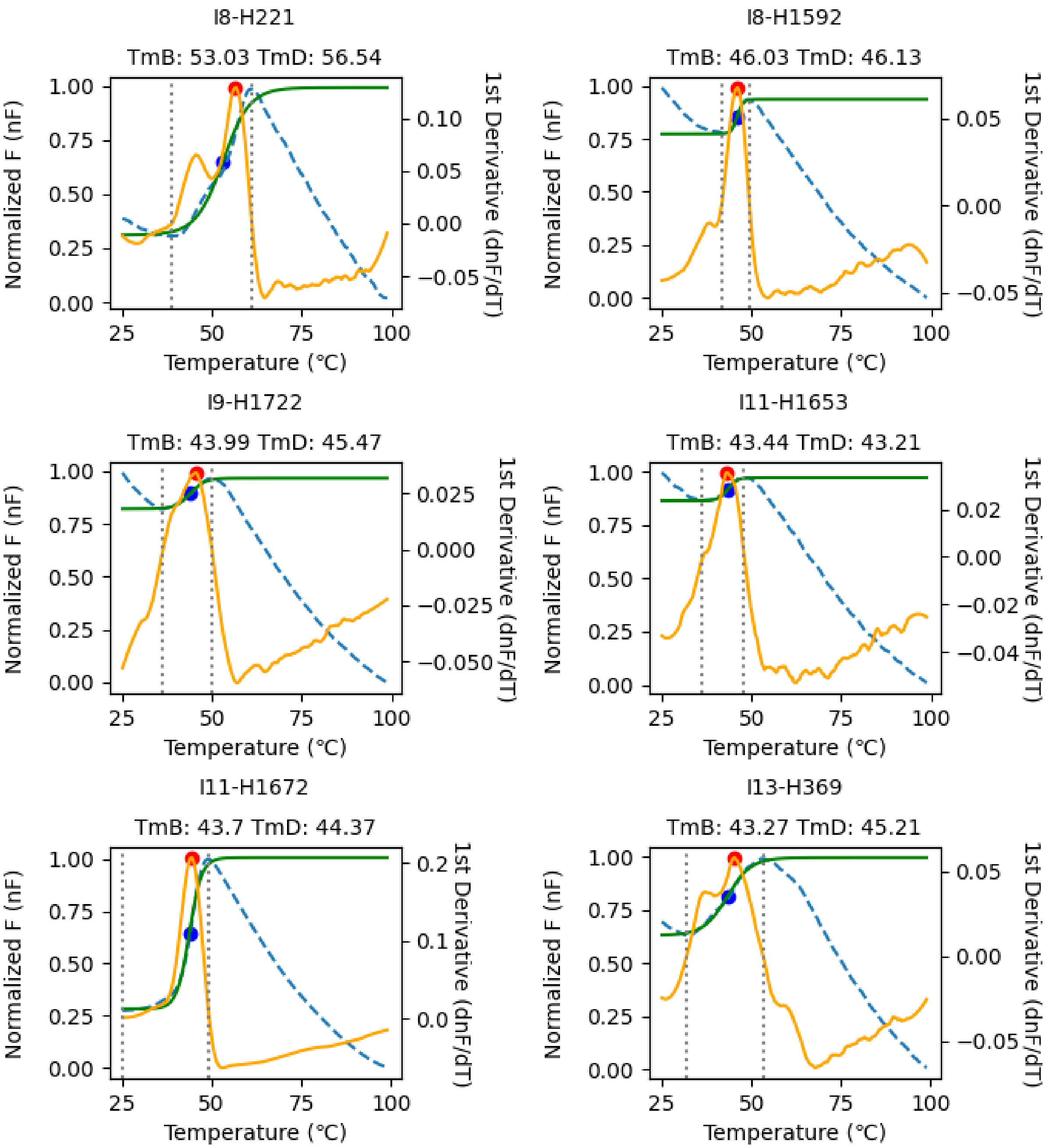


**Fig. S7. Melting temperature (Tm) determination of 158 Houses through differential scanning fluorimetry (continued, 12/27).** The normalized fluorescence (nF) and its 1st derivative is shown with a blue dashed and yellow solid lines, respectively. The Boltzmann curve fitted to the nF is shown with a green solid line. The determined $Tm_B$ and $Tm_D$ is shown with blue and red circles, respectively, and labeled.

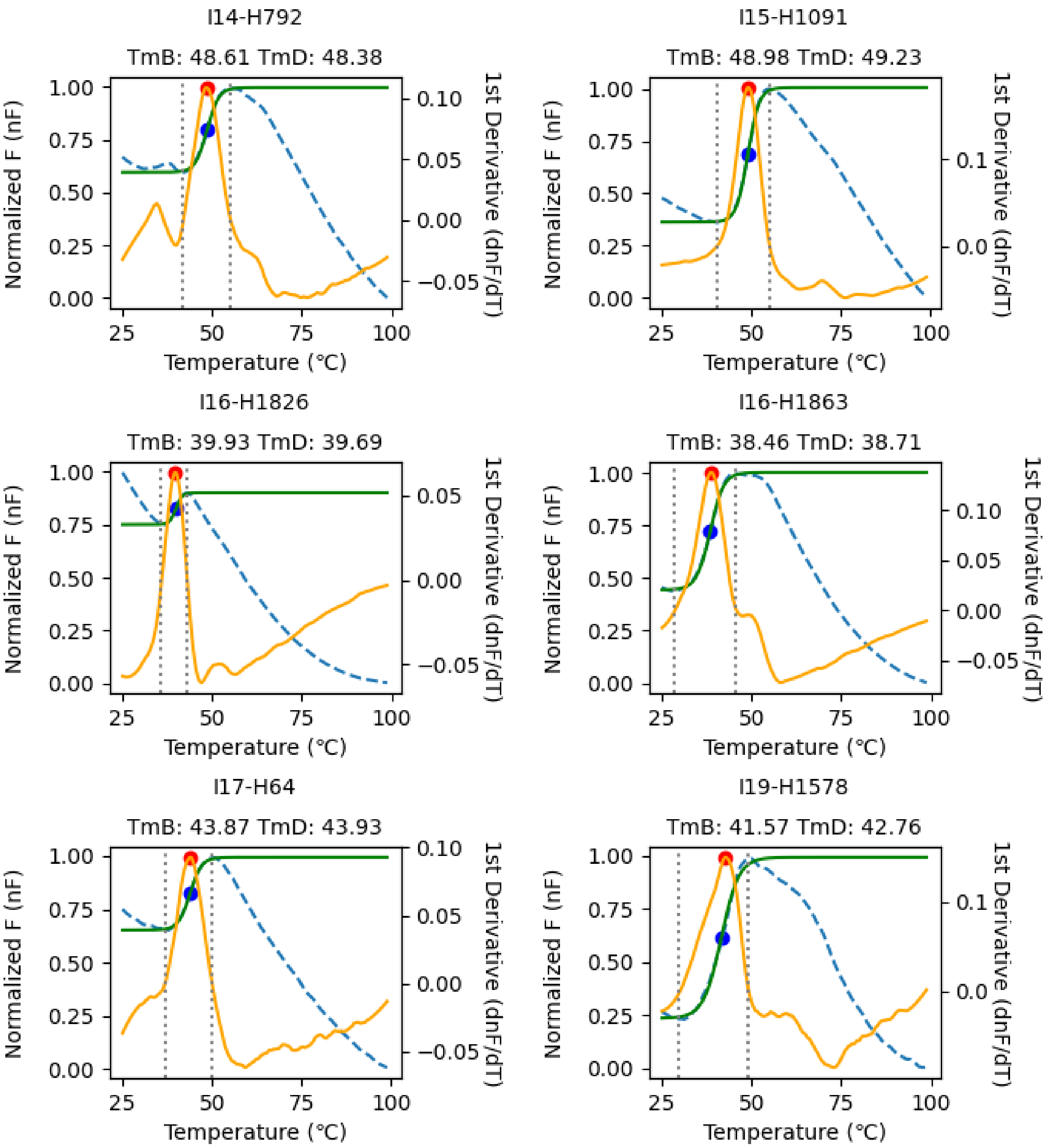


**Fig. S7. Melting temperature (Tm) determination of 158 Houses through differential scanning fluorimetry (continued, 13/27).** The normalized fluorescence (nF) and its 1st derivative is shown with a blue dashed and yellow solid lines, respectively. The Boltzmann curve fitted to the nF is shown with a green solid line. The determined $Tm_B$ and $Tm_D$ is shown with blue and red circles, respectively, and labeled.

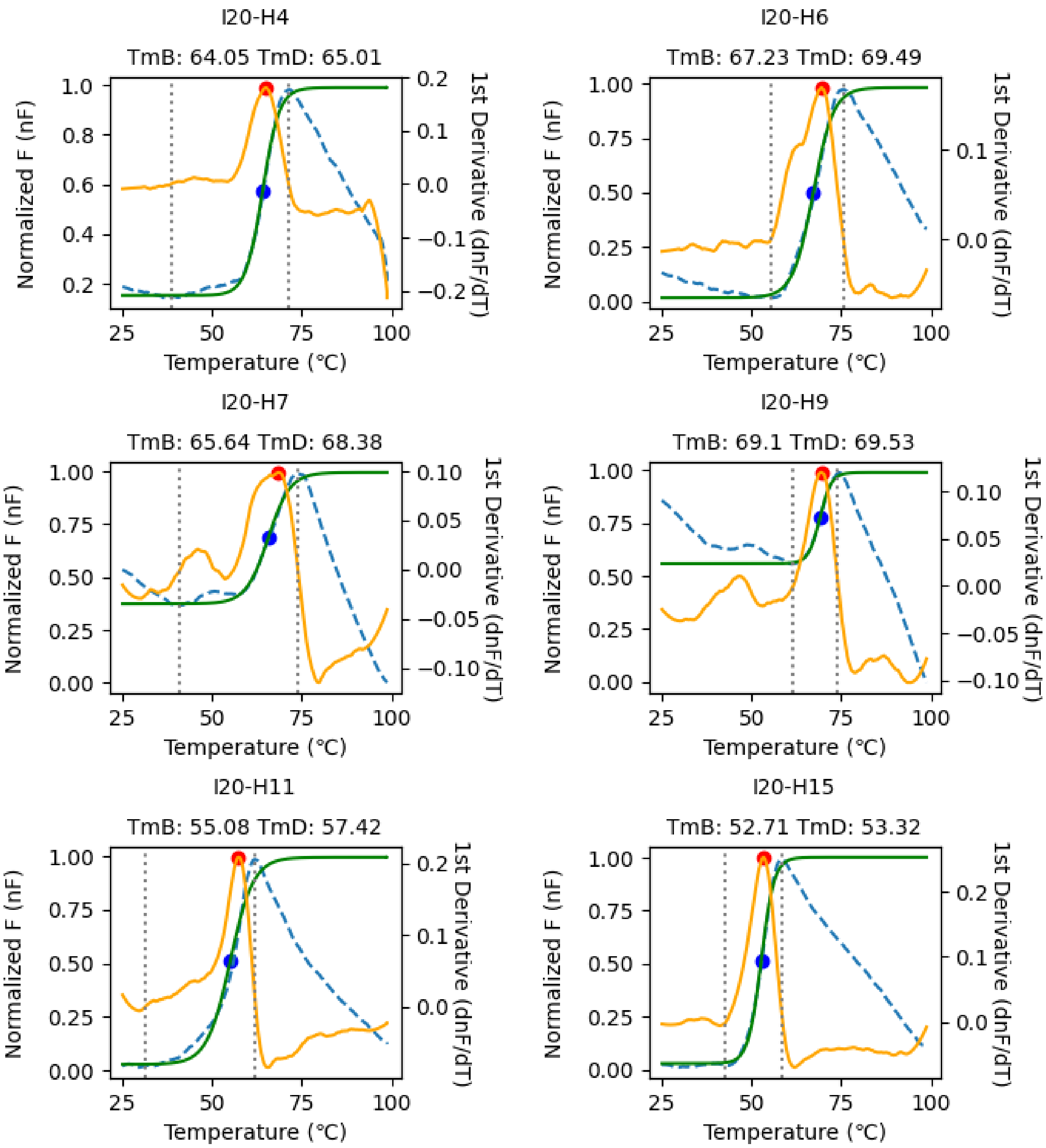


**Fig. S7. Melting temperature (Tm) determination of 158 Houses through differential scanning fluorimetry (continued, 14/27).** The normalized fluorescence (nF) and its 1st derivative is shown with a blue dashed and yellow solid lines, respectively. The Boltzmann curve fitted to the nF is shown with a green solid line. The determined $Tm_B$ and $Tm_D$ is shown with blue and red circles, respectively, and labeled.

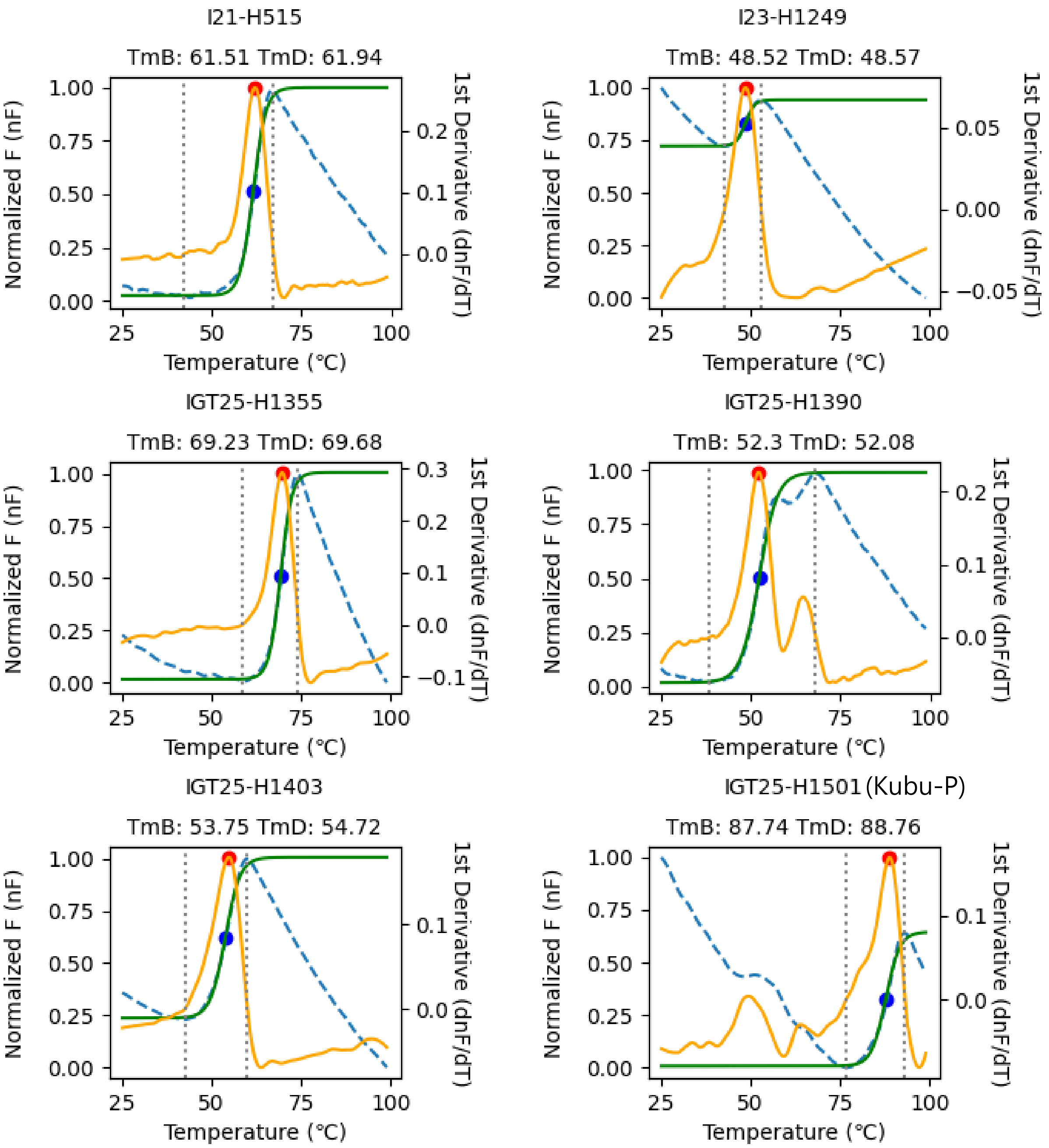


**Fig. S7. Melting temperature (Tm) determination of 158 Houses through differential scanning fluorimetry (continued, 15/27).** The normalized fluorescence (nF) and its 1st derivative is shown with a blue dashed and yellow solid lines, respectively. The Boltzmann curve fitted to the nF is shown with a green solid line. The determined $Tm_B$ and $Tm_D$ is shown with blue and red circles, respectively, and labeled.

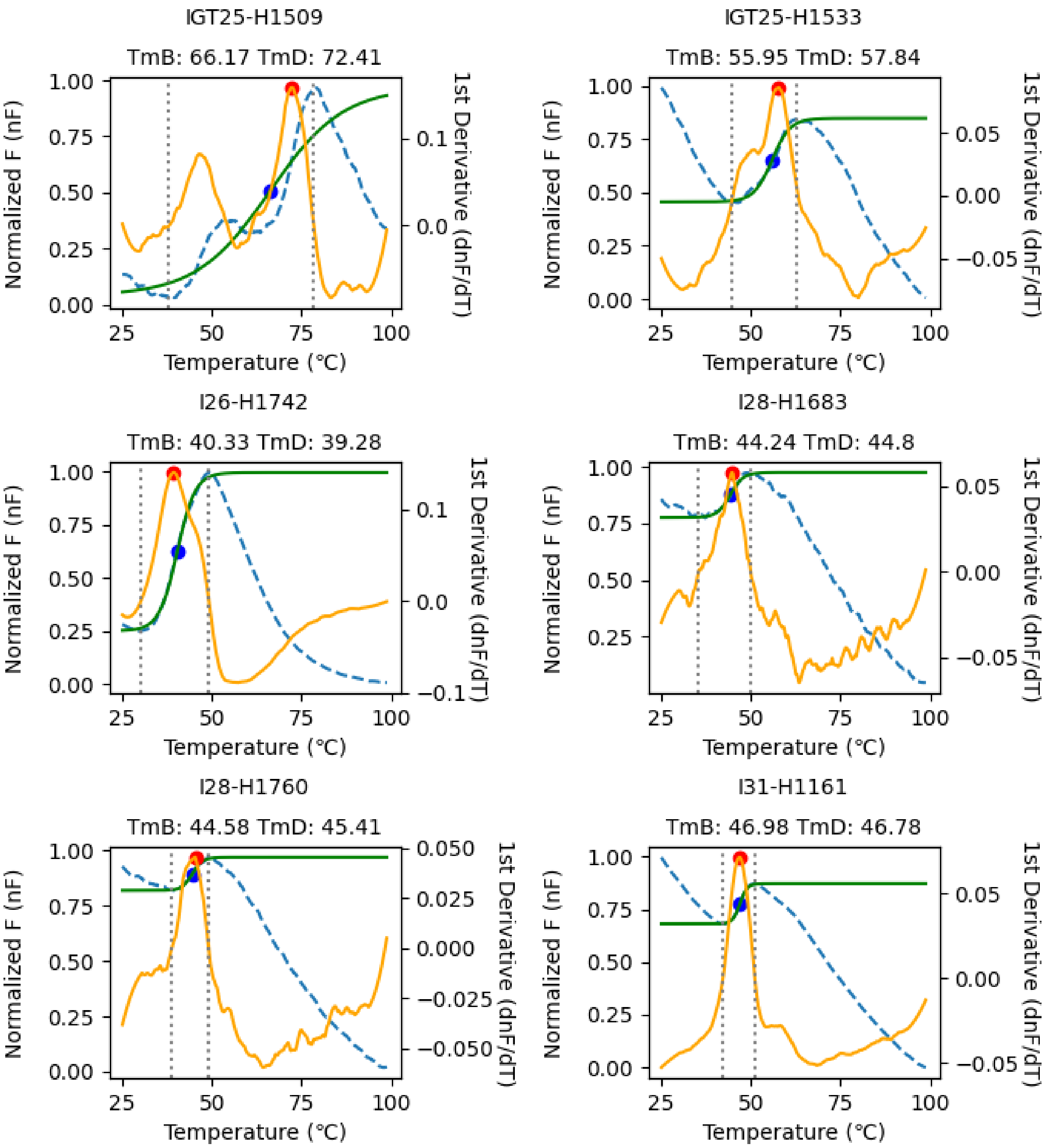


**Fig. S7. Melting temperature (Tm) determination of 158 Houses through differential scanning fluorimetry (continued, 16/27).** The normalized fluorescence (nF) and its 1st derivative is shown with a blue dashed and yellow solid lines, respectively. The Boltzmann curve fitted to the nF is shown with a green solid line. The determined $Tm_B$ and $Tm_D$ is shown with blue and red circles, respectively, and labeled.

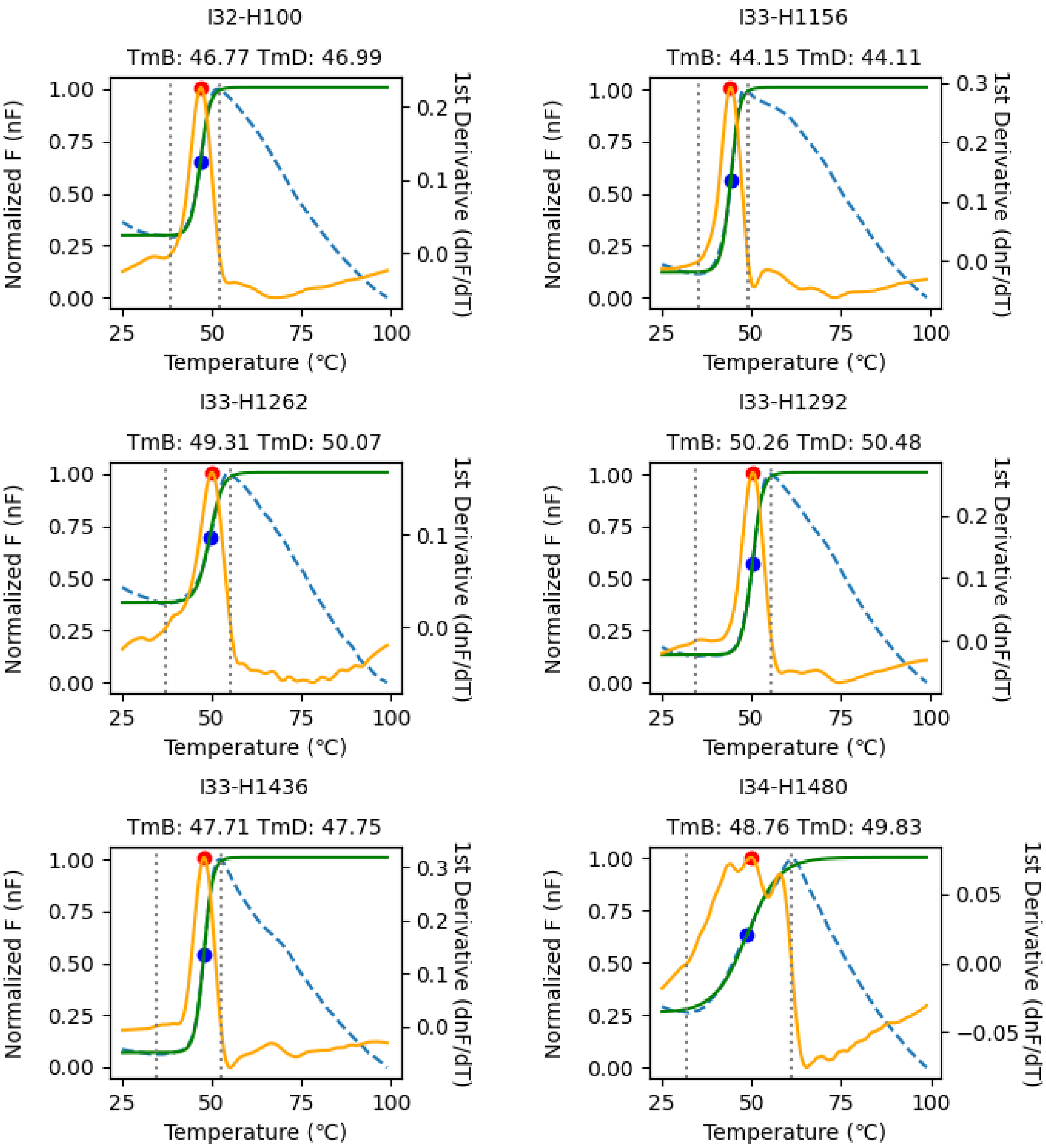


**Fig. S7. Melting temperature (Tm) determination of 158 Houses through differential scanning fluorimetry (continued, 17/27).** The normalized fluorescence (nF) and its 1st derivative is shown with a blue dashed and yellow solid lines, respectively. The Boltzmann curve fitted to the nF is shown with a green solid line. The determined $Tm_B$ and $Tm_D$ is shown with blue and red circles, respectively, and labeled.

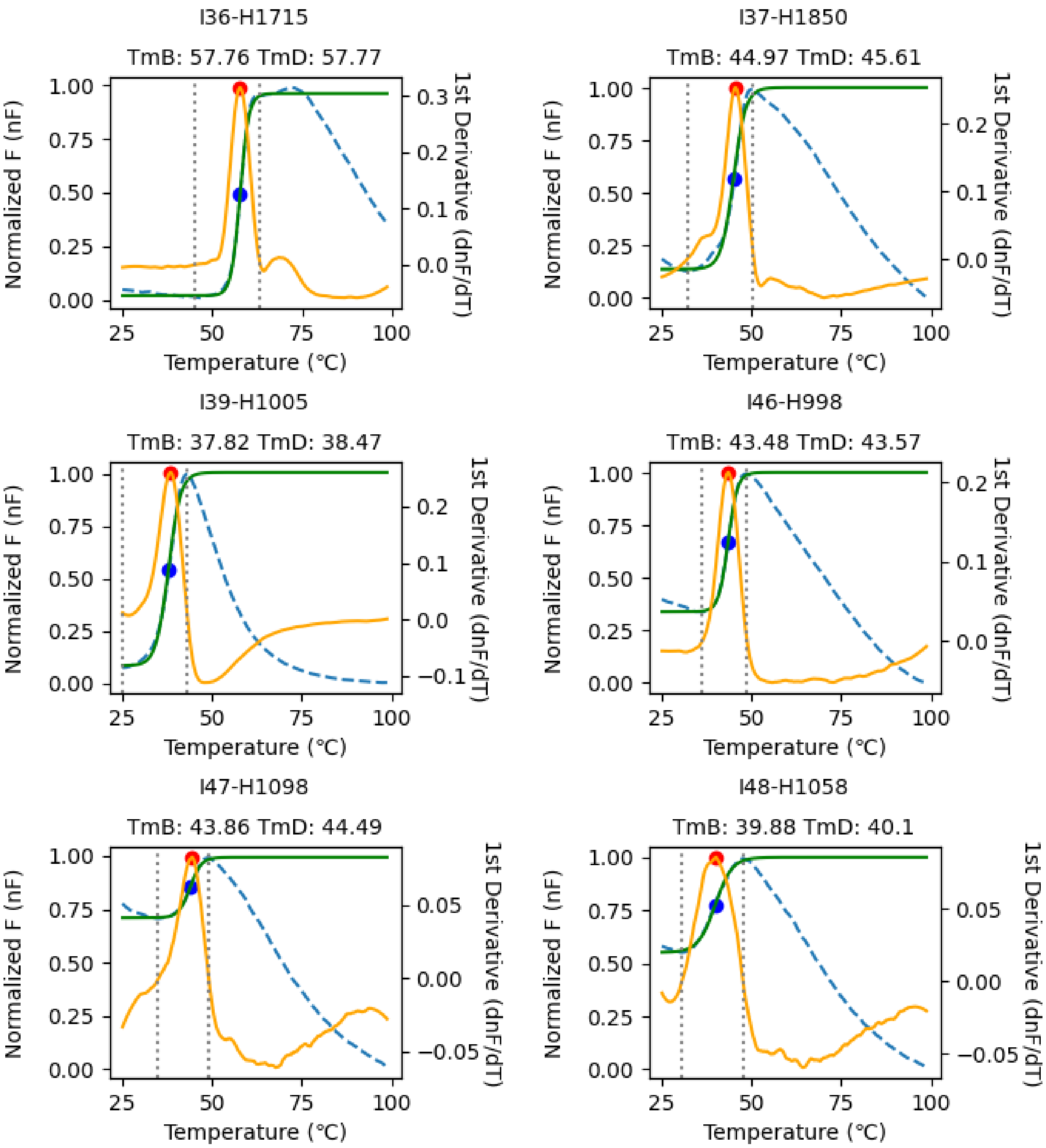


**Fig. S7. Melting temperature (Tm) determination of 158 Houses through differential scanning fluorimetry (continued, 18/27).** The normalized fluorescence (nF) and its 1st derivative is shown with a blue dashed and yellow solid lines, respectively. The Boltzmann curve fitted to the nF is shown with a green solid line. The determined $Tm_B$ and $Tm_D$ is shown with blue and red circles, respectively, and labeled.

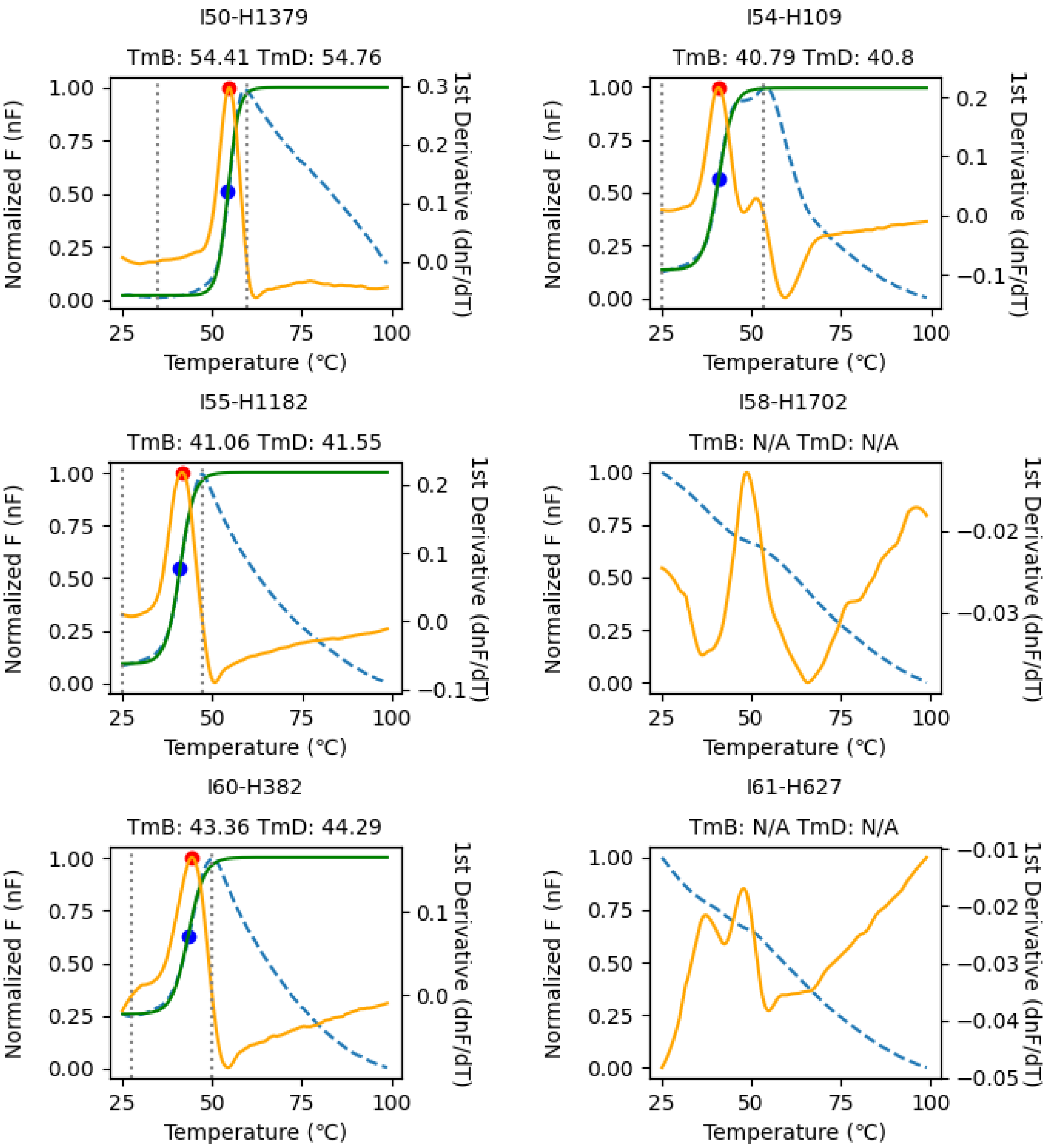


**Fig. S7. Melting temperature (Tm) determination of 158 Houses through differential scanning fluorimetry (continued, 19/27).** The normalized fluorescence (nF) and its 1st derivative is shown with a blue dashed and yellow solid lines, respectively. The Boltzmann curve fitted to the nF is shown with a green solid line. The determined $Tm_B$ and $Tm_D$ is shown with blue and red circles, respectively, and labeled.

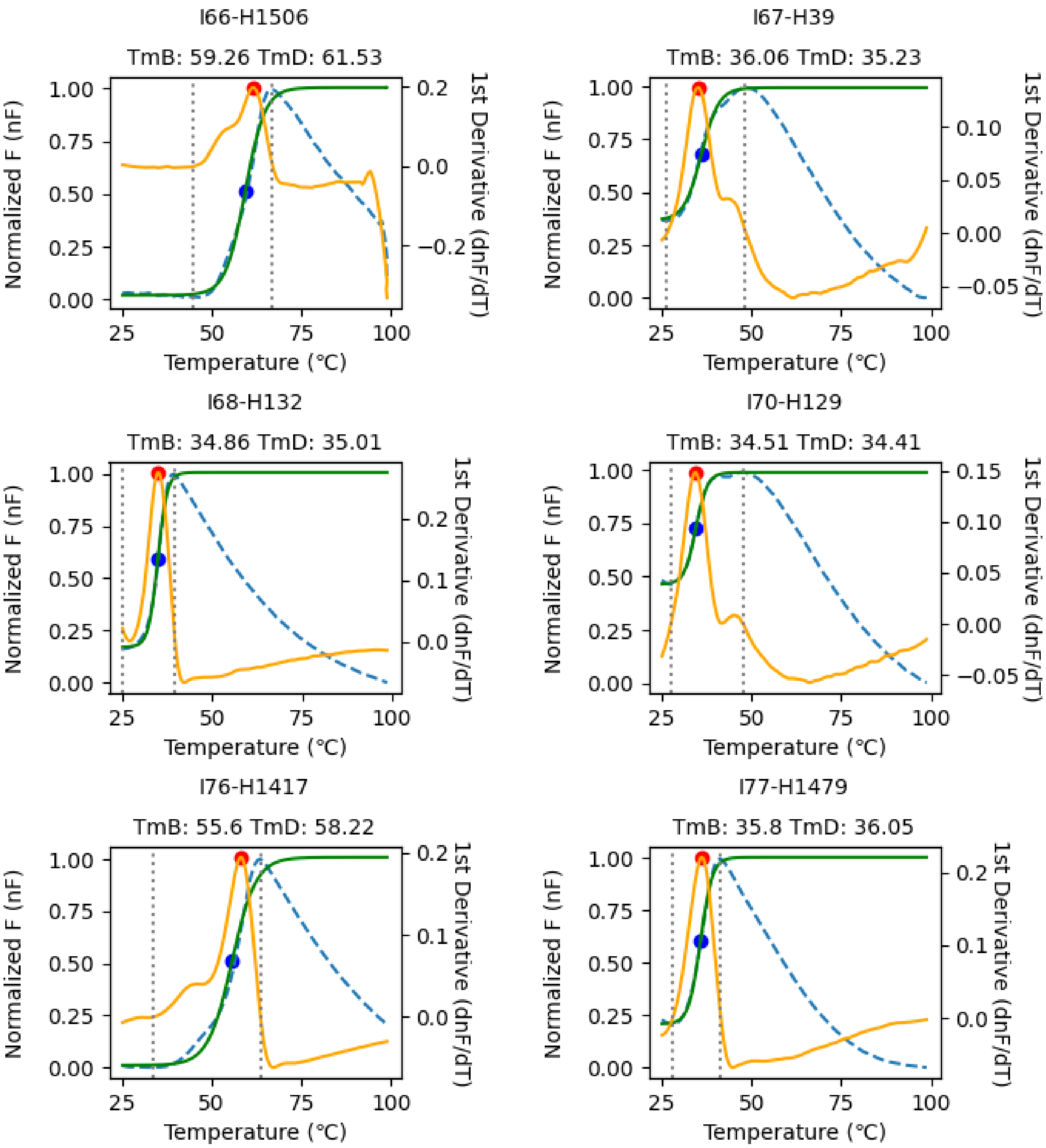


**Fig. S7. Melting temperature (Tm) determination of 158 Houses through differential scanning fluorimetry (continued, 20/27).** The normalized fluorescence (nF) and its 1st derivative is shown with a blue dashed and yellow solid lines, respectively. The Boltzmann curve fitted to the nF is shown with a green solid line. The determined $Tm_B$ and $Tm_D$ is shown with blue and red circles, respectively, and labeled.

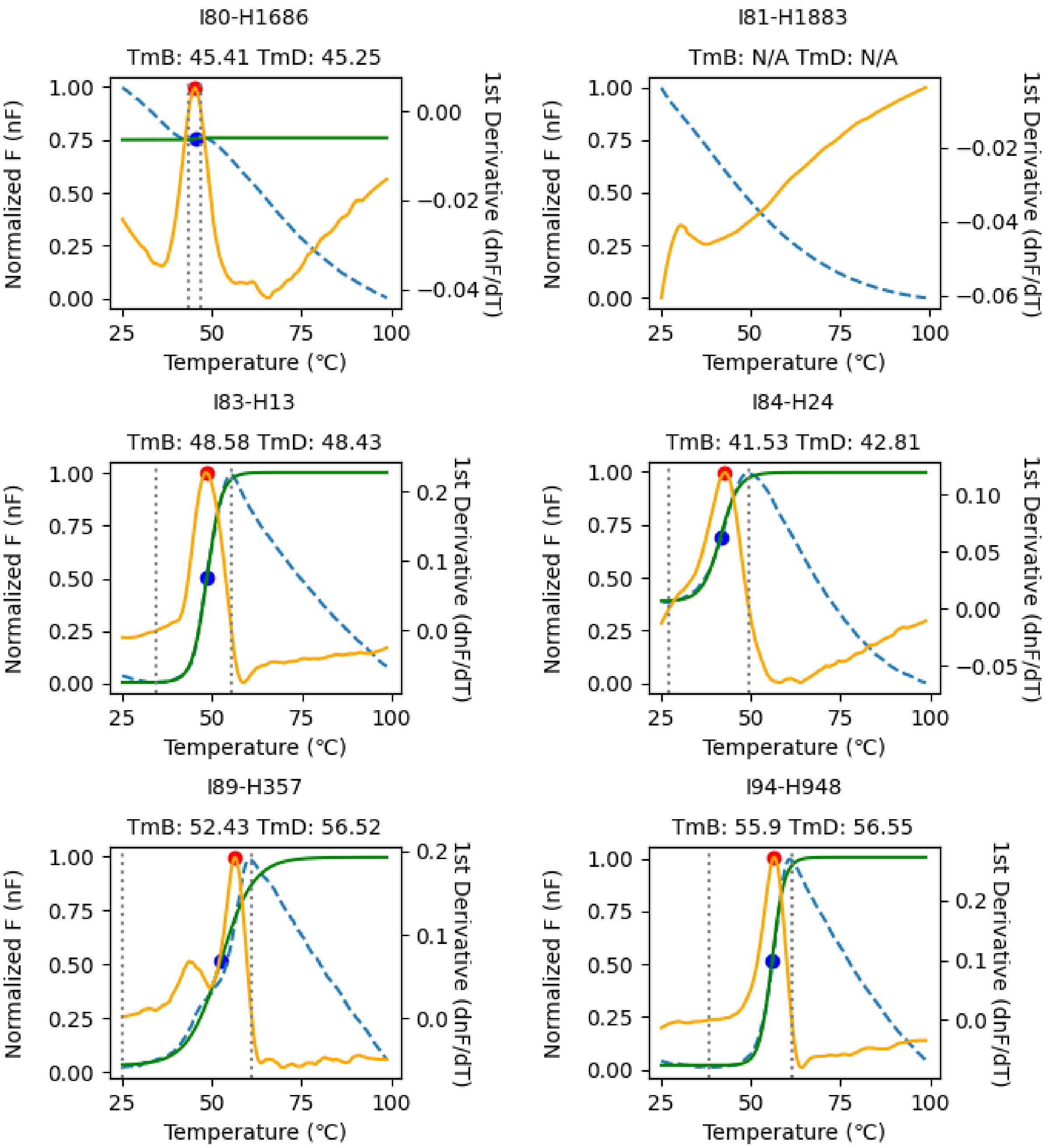


**Fig. S7. Melting temperature (Tm) determination of 158 Houses through differential scanning fluorimetry (continued, 21/27).** The normalized fluorescence (nF) and its 1st derivative is shown with a blue dashed and yellow solid lines, respectively. The Boltzmann curve fitted to the nF is shown with a green solid line. The determined $Tm_B$ and $Tm_D$ is shown with blue and red circles, respectively, and labeled.

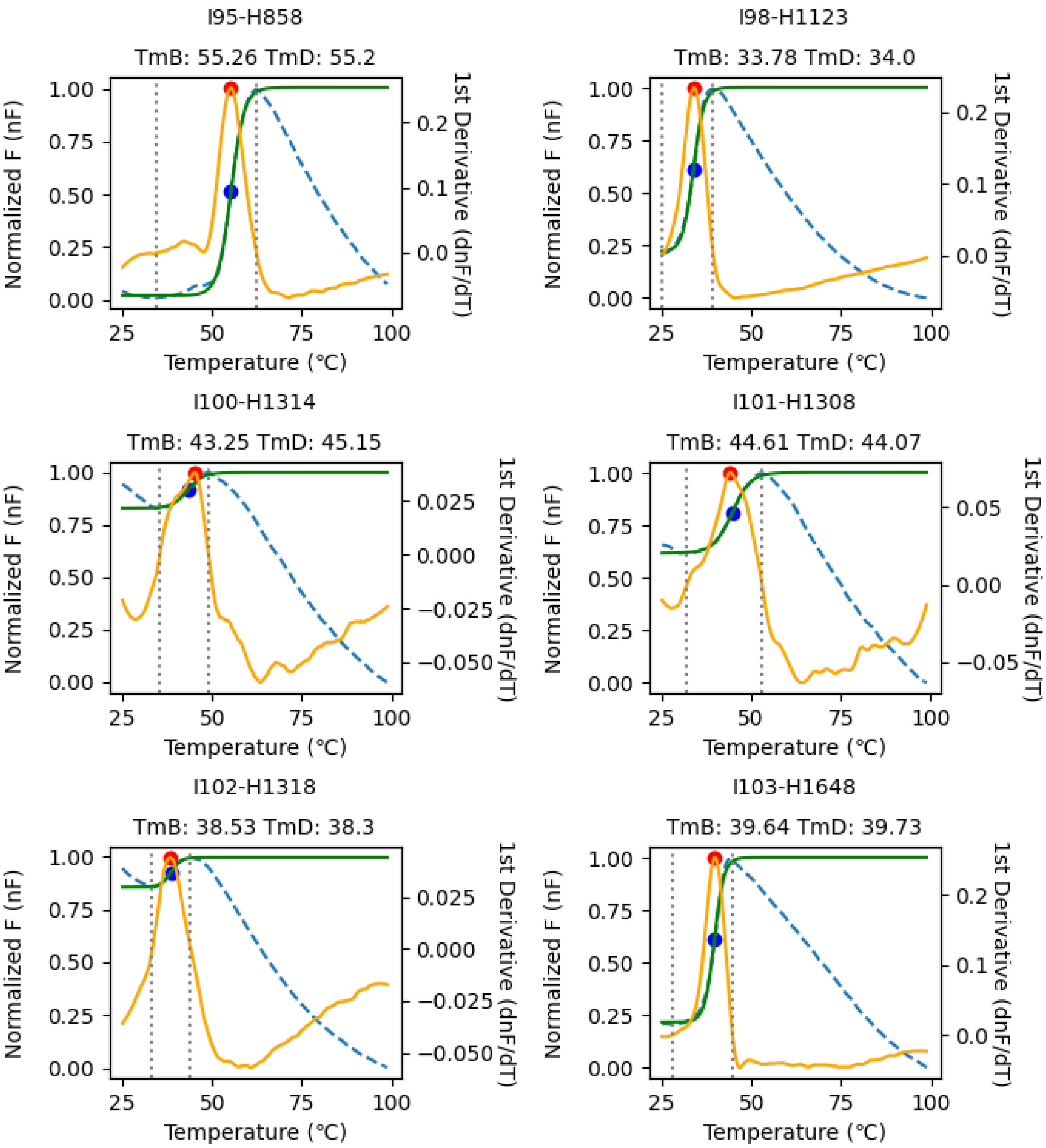


**Fig. S7. Melting temperature (Tm) determination of 158 Houses through differential scanning fluorimetry (continued, 22/27).** The normalized fluorescence (nF) and its 1st derivative is shown with a blue dashed and yellow solid lines, respectively. The Boltzmann curve fitted to the nF is shown with a green solid line. The determined $Tm_B$ and $Tm_D$ is shown with blue and red circles, respectively, and labeled.

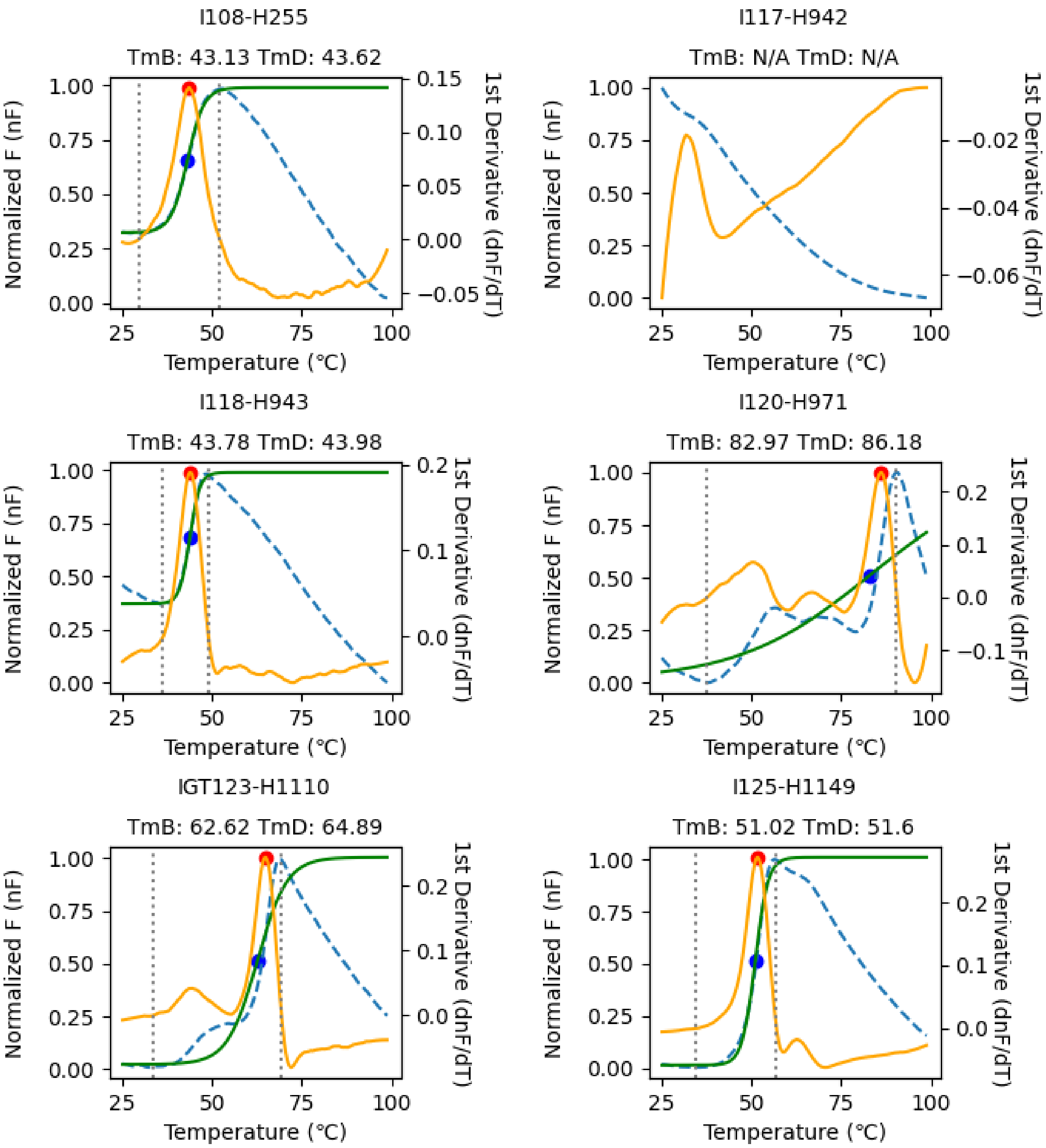


**Fig. S7. Melting temperature (Tm) determination of 158 Houses through differential scanning fluorimetry (continued, 23/27).** The normalized fluorescence (nF) and its 1st derivative is shown with a blue dashed and yellow solid lines, respectively. The Boltzmann curve fitted to the nF is shown with a green solid line. The determined $Tm_B$ and $Tm_D$ is shown with blue and red circles, respectively, and labeled.

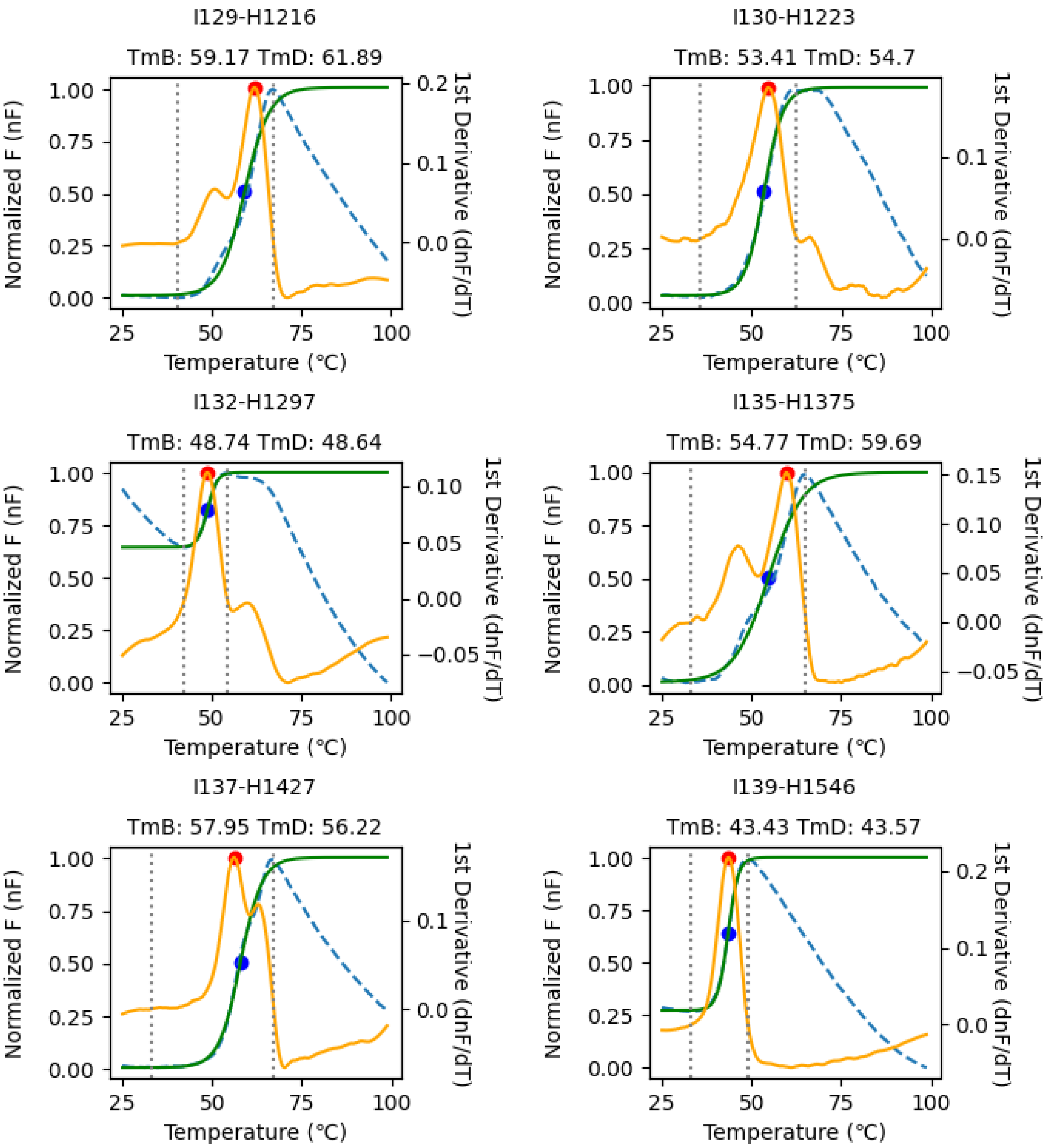


**Fig. S7. Melting temperature (Tm) determination of 158 Houses through differential scanning fluorimetry (continued, 24/27).** The normalized fluorescence (nF) and its 1st derivative is shown with a blue dashed and yellow solid lines, respectively. The Boltzmann curve fitted to the nF is shown with a green solid line. The determined $Tm_B$ and $Tm_D$ is shown with blue and red circles, respectively, and labeled.

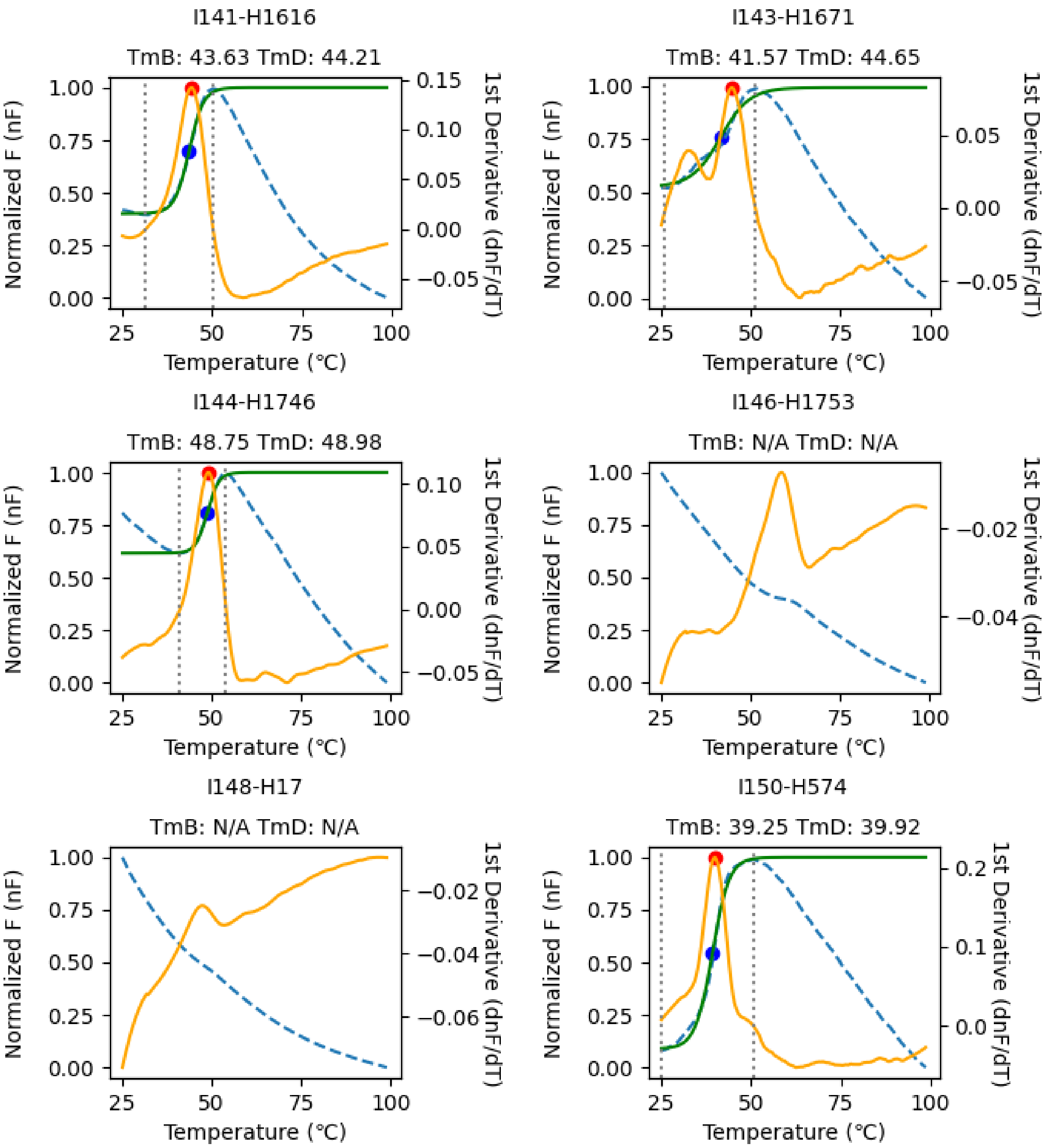


**Fig. S7. Melting temperature (Tm) determination of 158 Houses through differential scanning fluorimetry (continued, 25/27).** The normalized fluorescence (nF) and its 1st derivative is shown with a blue dashed and yellow solid lines, respectively. The Boltzmann curve fitted to the nF is shown with a green solid line. The determined $Tm_B$ and $Tm_D$ is shown with blue and red circles, respectively, and labeled.

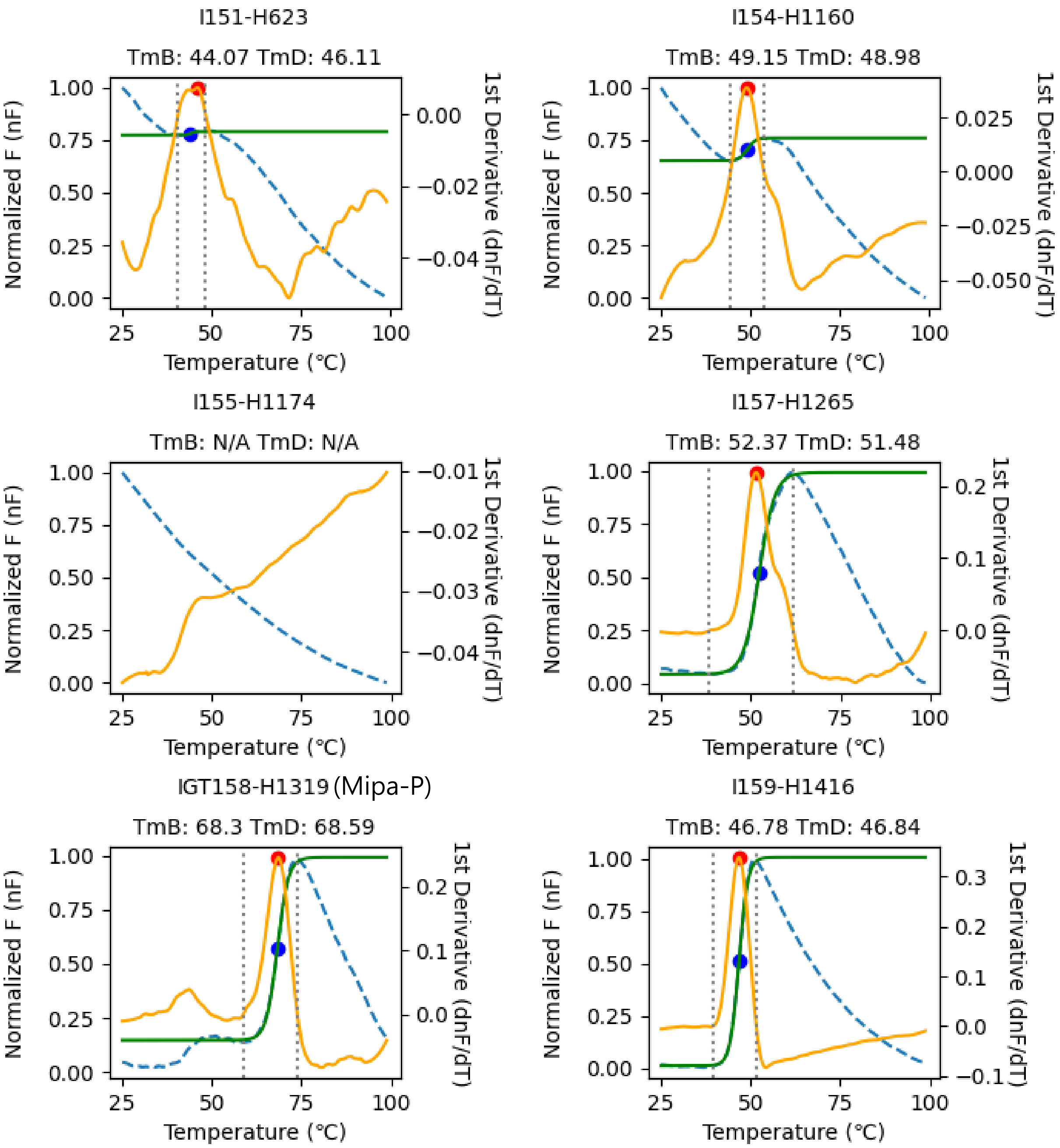


**Fig. S7. Melting temperature (Tm) determination of 158 Houses through differential scanning fluorimetry (continued, 26/27).** The normalized fluorescence (nF) and its 1st derivative is shown with a blue dashed and yellow solid lines, respectively. The Boltzmann curve fitted to the nF is shown with a green solid line. The determined $Tm_B$ and $Tm_D$ is shown with blue and red circles, respectively, and labeled.

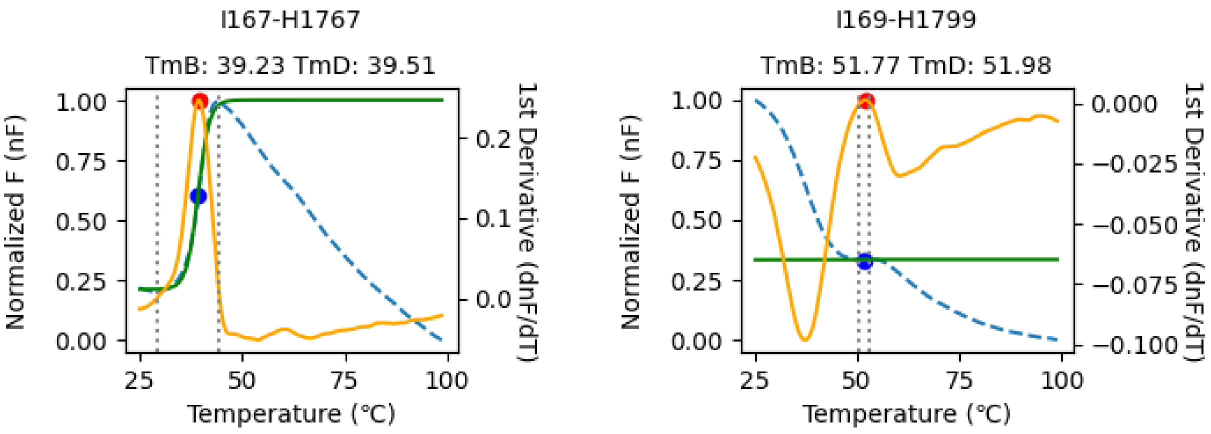


**Fig. S7. Melting temperature (Tm) determination of 158 Houses through differential scanning fluorimetry (27/27).** The normalized fluorescence (nF) and its 1st derivative is shown with a blue dashed and yellow solid lines, respectively. The Boltzmann curve fitted to the nF is shown with a green solid line. The determined $Tm_B$ and $Tm_D$ is shown with blue and red circles, respectively, and labeled.

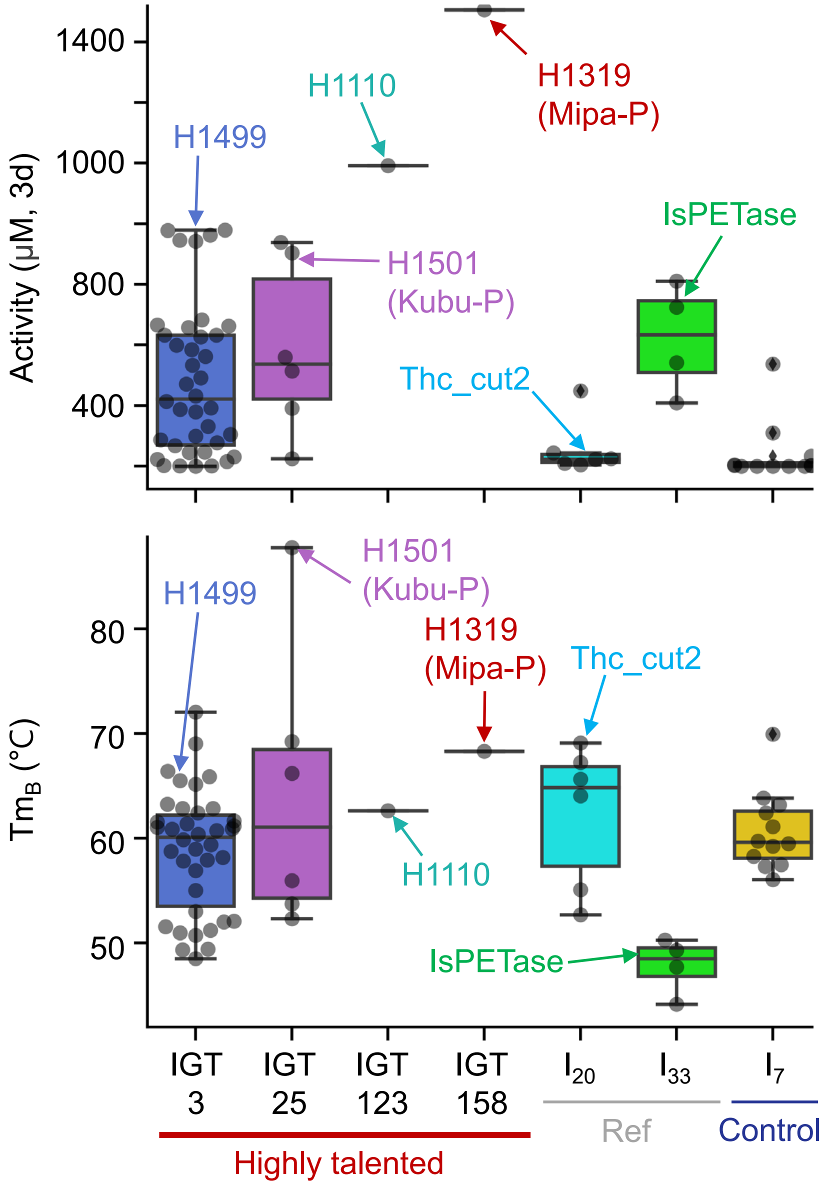


**Fig. S8. Box plots of activity and Tm values by Islands in the secondary contest.** H1499, H1501, H1110 (CaPETase), and H1319, and the benchmarks, IsPETase and Thc_Cut2, of the reference (well-known) islands are indicated by arrows and labeled. Control means the Island with low activity and medium number of Houses, showing representativeness. Houses tested in these seven Islands are individually shown as black dots.

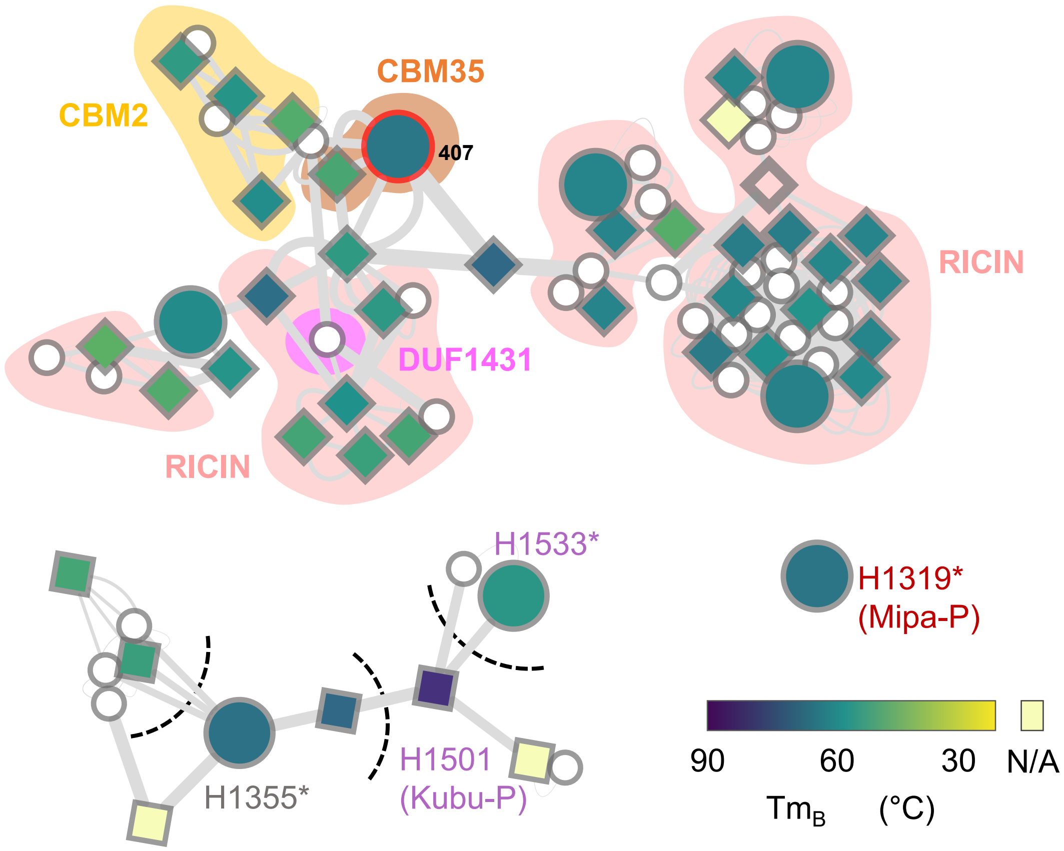


**Fig. S9. Network of unexplored IGTs and their Tm distributions.** The figure is drawn as in Fig. 2C, except for the color scheme of the node fill (Tm). The shadows in IGT3 with yellow, orange, salmon, and pink colors indicate additional C-terminal domains.

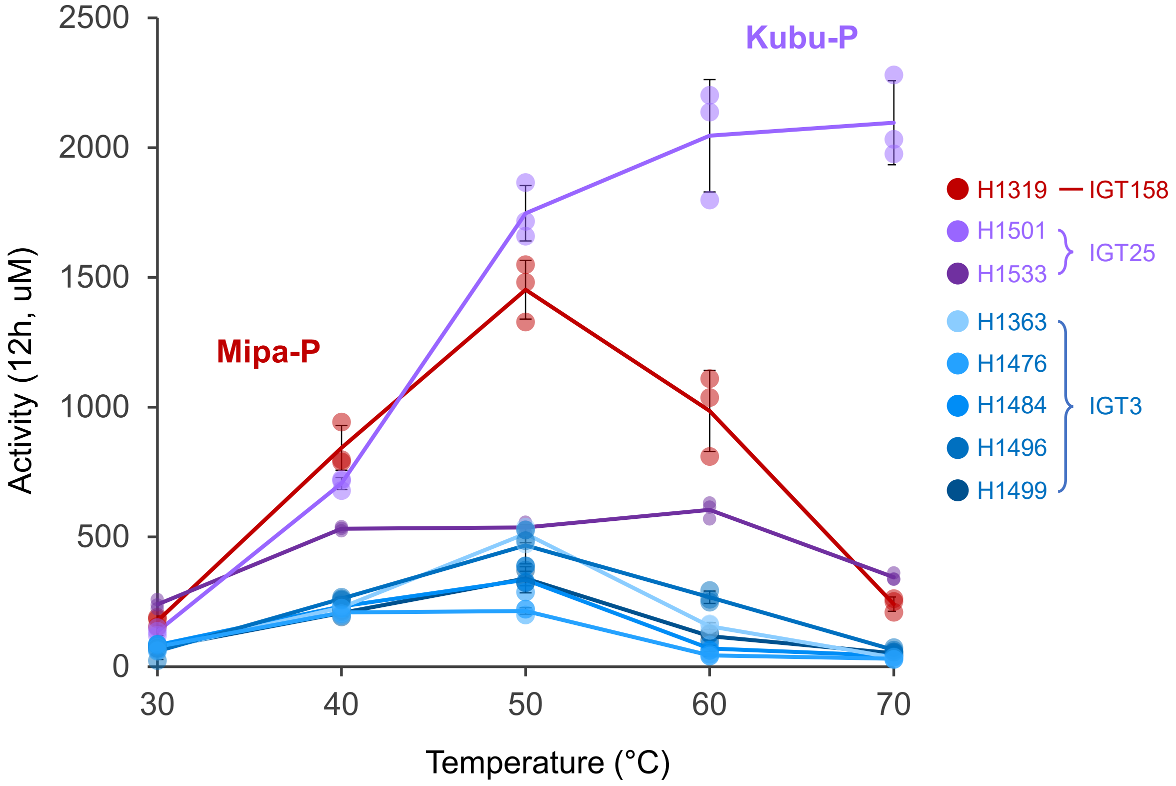


**Fig. S10. The final evaluation contest at the various temperatures.** 12 h monomer release activities along temperatures are shown. Figure legend with House and Island number is shown. The two outstanding Houses are labeled with their names.

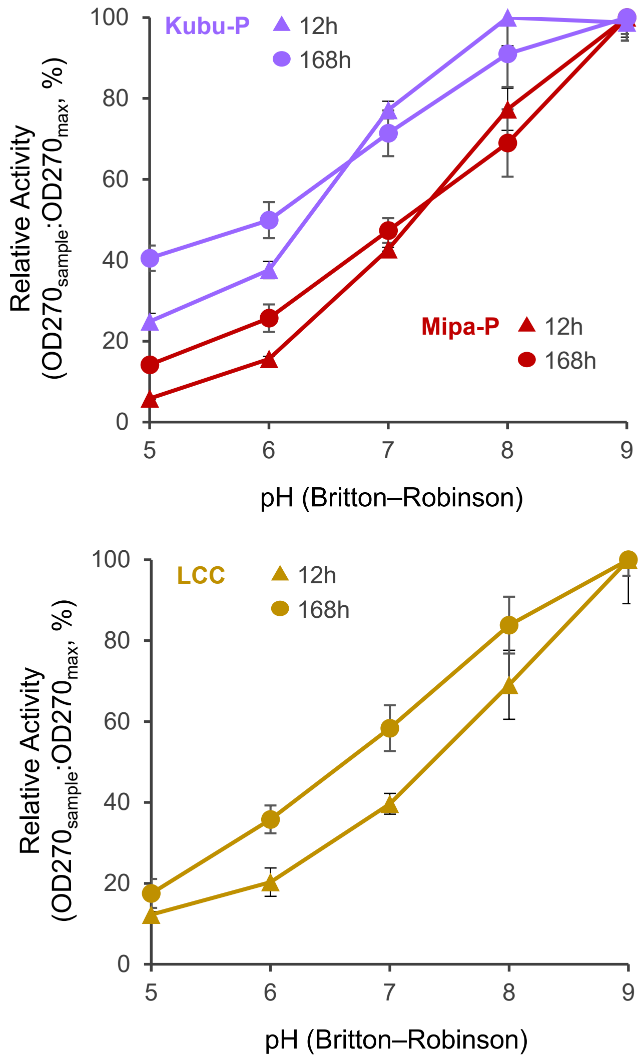


**Fig. S11. pH dependent activity of Mipa-P, Kubu-P, and LCC.** 12 and 168 h relative activities are shown along pH values. All three enzymes prefer alkaline conditions.

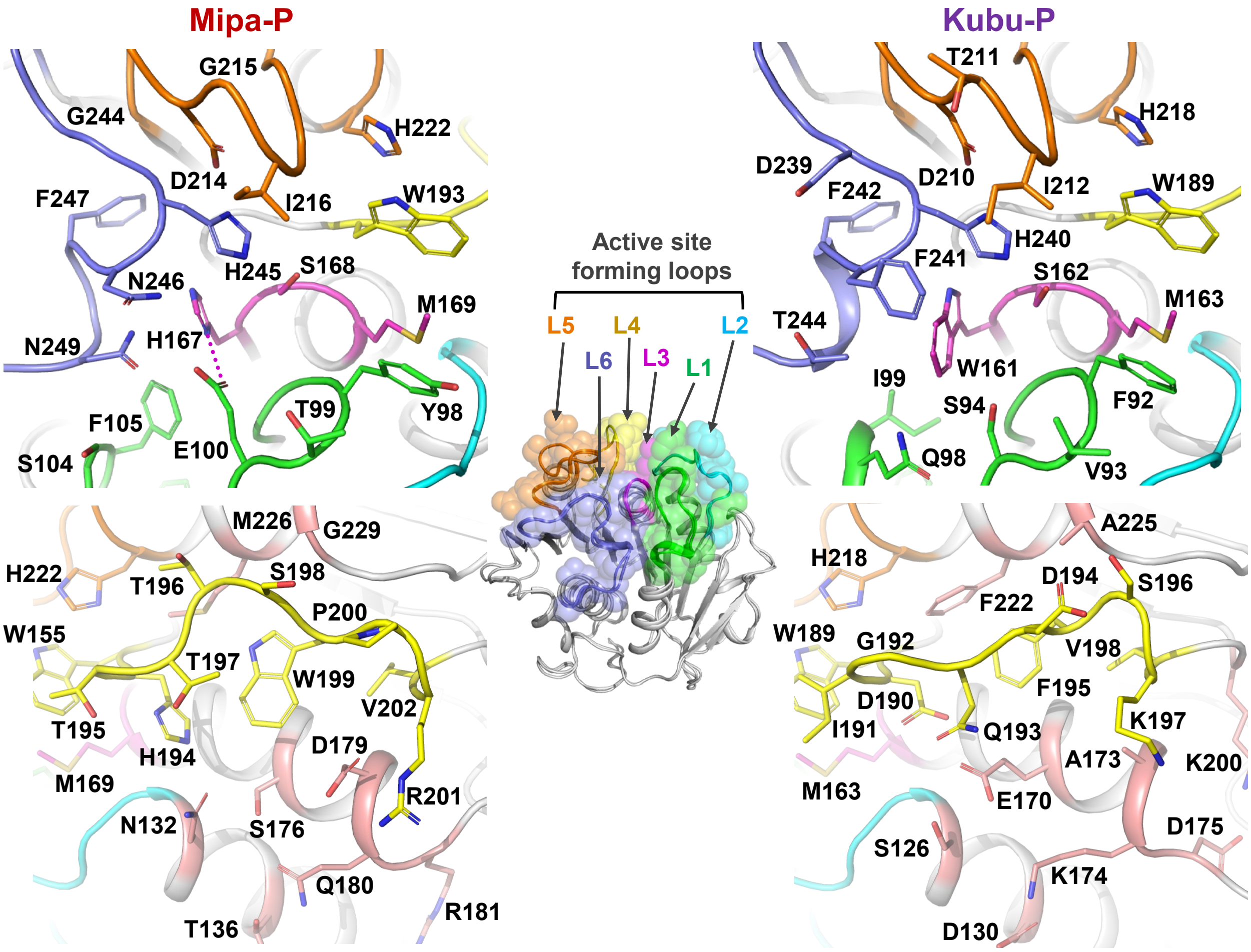


**Fig. S12. Structures of Mipa-P and Kubu-P.** The structures are shown as cartoon models. Each loop regions are distinguished with different color scheme. Upper and lower figures are in views of catalytic triad and L4, respectively. The salmon color in the lowers figures indicates additional positions that are not on the six loops but form the active site network. All residues involved in forming the active site are labeled.

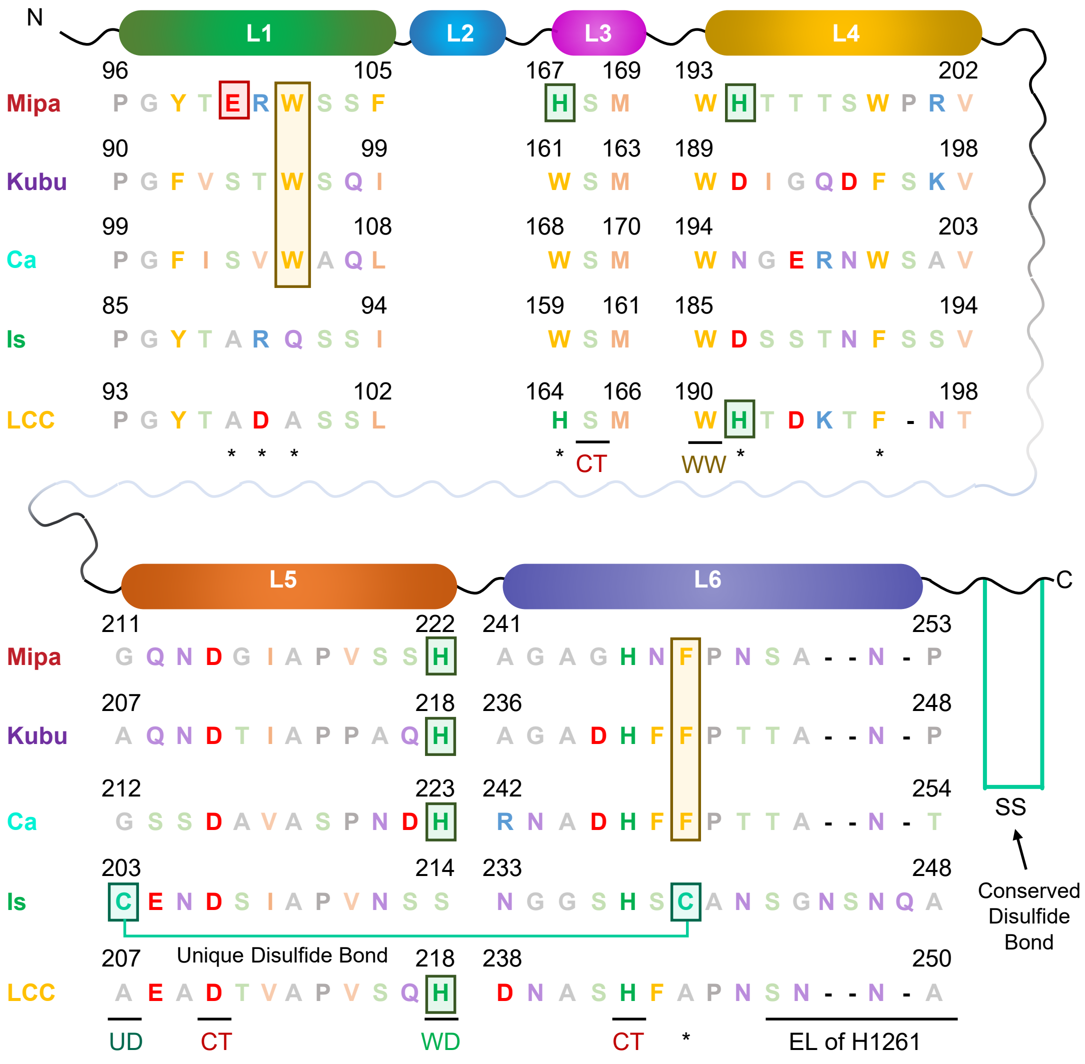


**Fig. S13. Alignments of key residues in the loops forming the active site.** Asterisk indicate residue position referred in the Supplementary Text. Residual features of Mipa-P and Kubu-P are distinguished with boxes. Abbreviations: WW, wobbling tryptophan; UD, unique disulfide bond in IsPETase; CT, catalytic triad; WD, wobbling tryptophan determinant; EL, extended loop.

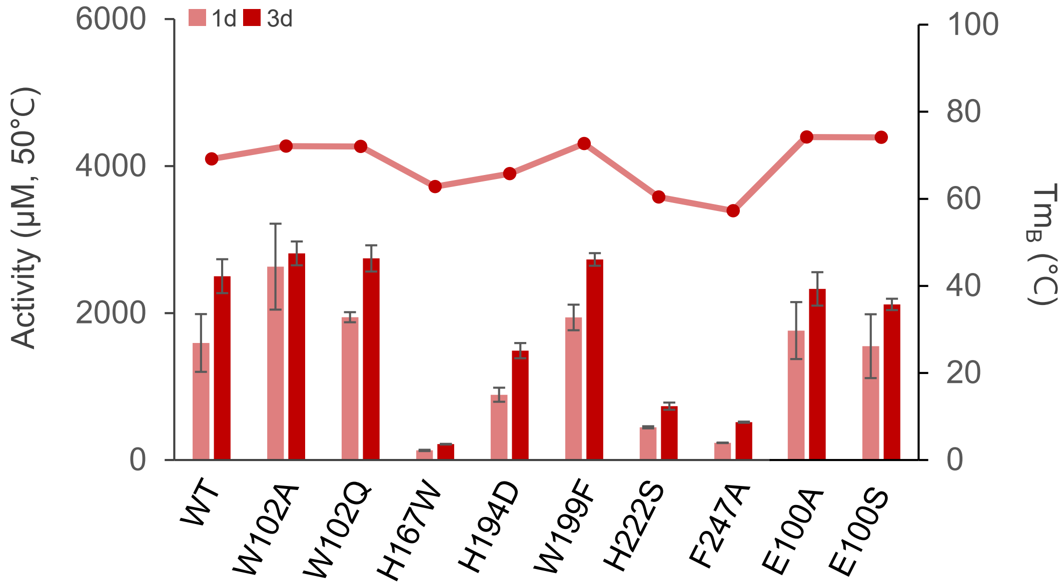


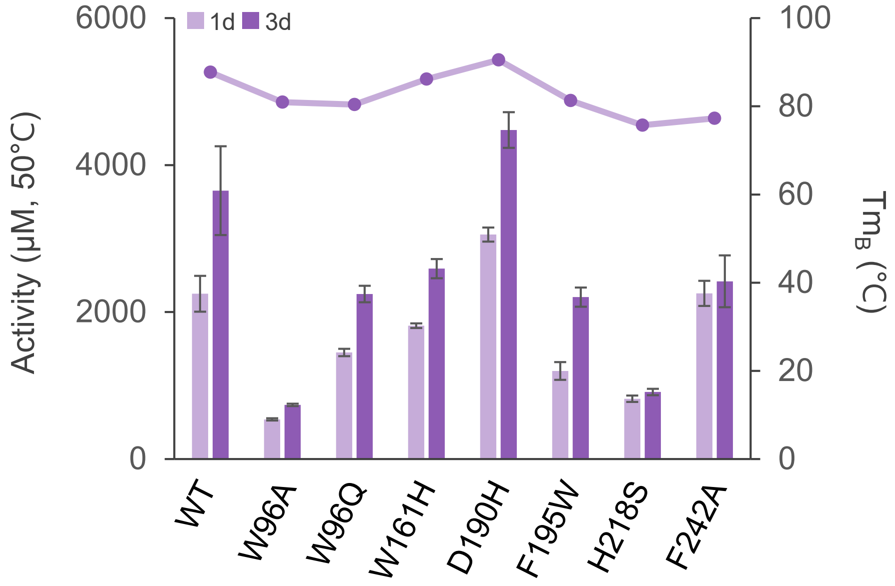


**Fig. S14. Activity and Tm of variants with structure-based replacement.** The upper (Mipa-P) and lower (Kubu-P) figures shows activity (bar graph) and Tm (line graph) values of the enzyme variants. The activities of 1 and 3 days at 50 °C were shown and labeled.

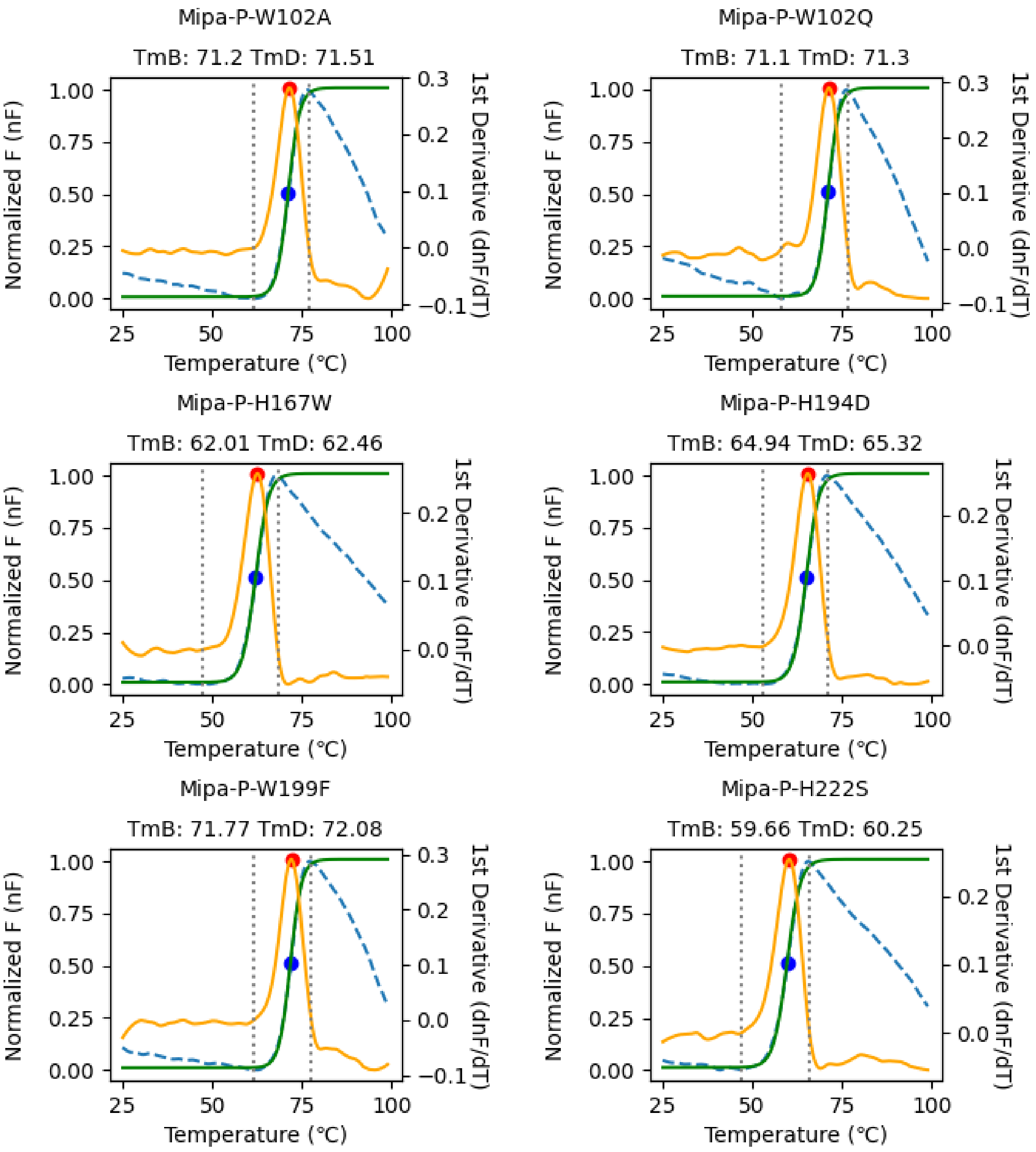


**Fig. S15. Melting temperature (Tm) determination of the structure-based replacement variants through differential scanning fluorimetry (continued, 1/3).** The normalized fluorescence (nF) and its 1st derivative is shown with a blue dashed and yellow solid lines, respectively. The Boltzmann curve fitted to the nF is shown with a green solid line. The determined $Tm_B$ and $Tm_D$ is shown with blue and red circles, respectively, and labeled.

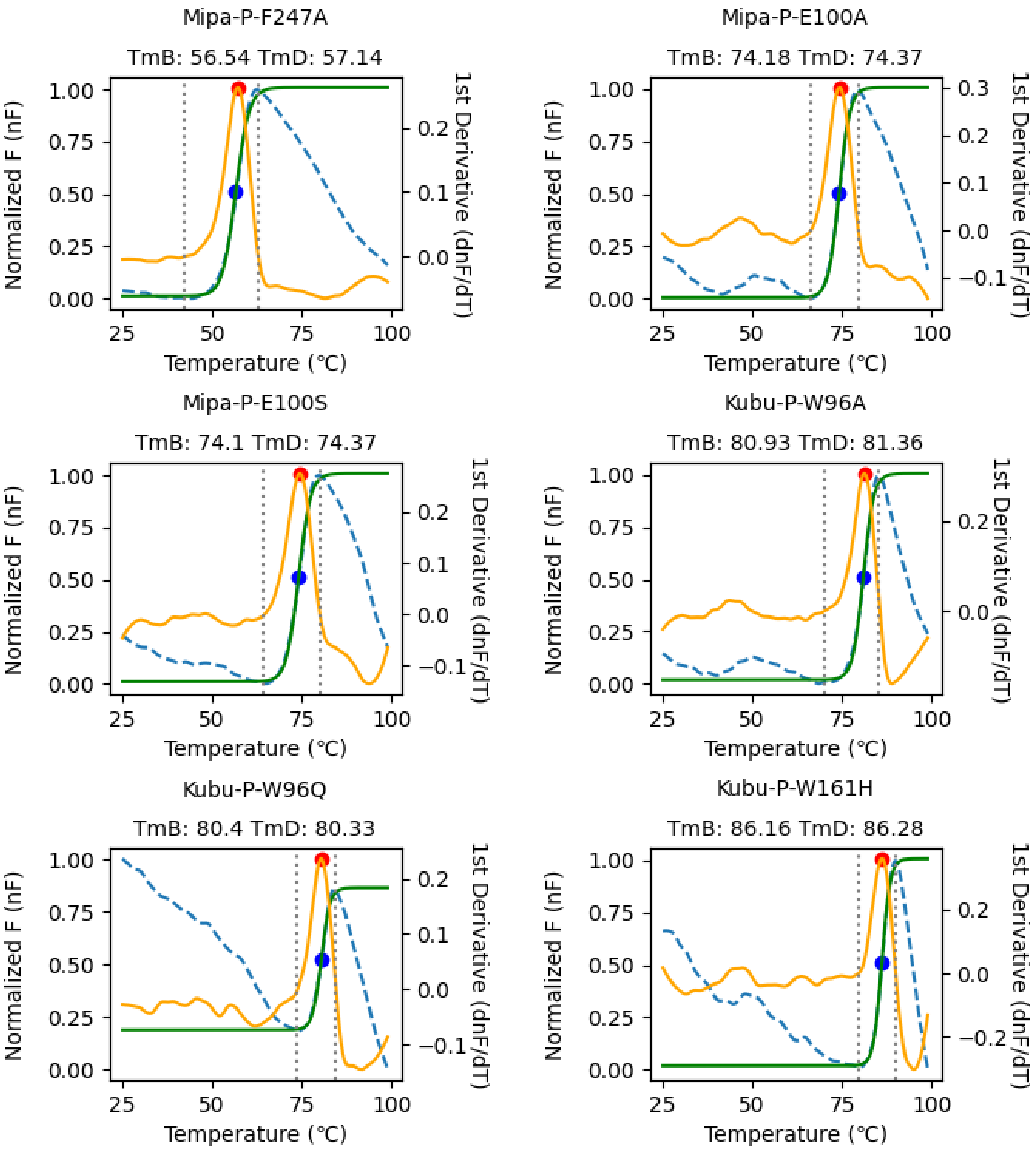


**Fig. S15. Melting temperature (Tm) determination of the structure-based replacement variants through differential scanning fluorimetry (continued, 2/3).** The normalized fluorescence (nF) and its 1st derivative is shown with a blue dashed and yellow solid lines, respectively. The Boltzmann curve fitted to the nF is shown with a green solid line. The determined $Tm_B$ and $Tm_D$ is shown with blue and red circles, respectively, and labeled.

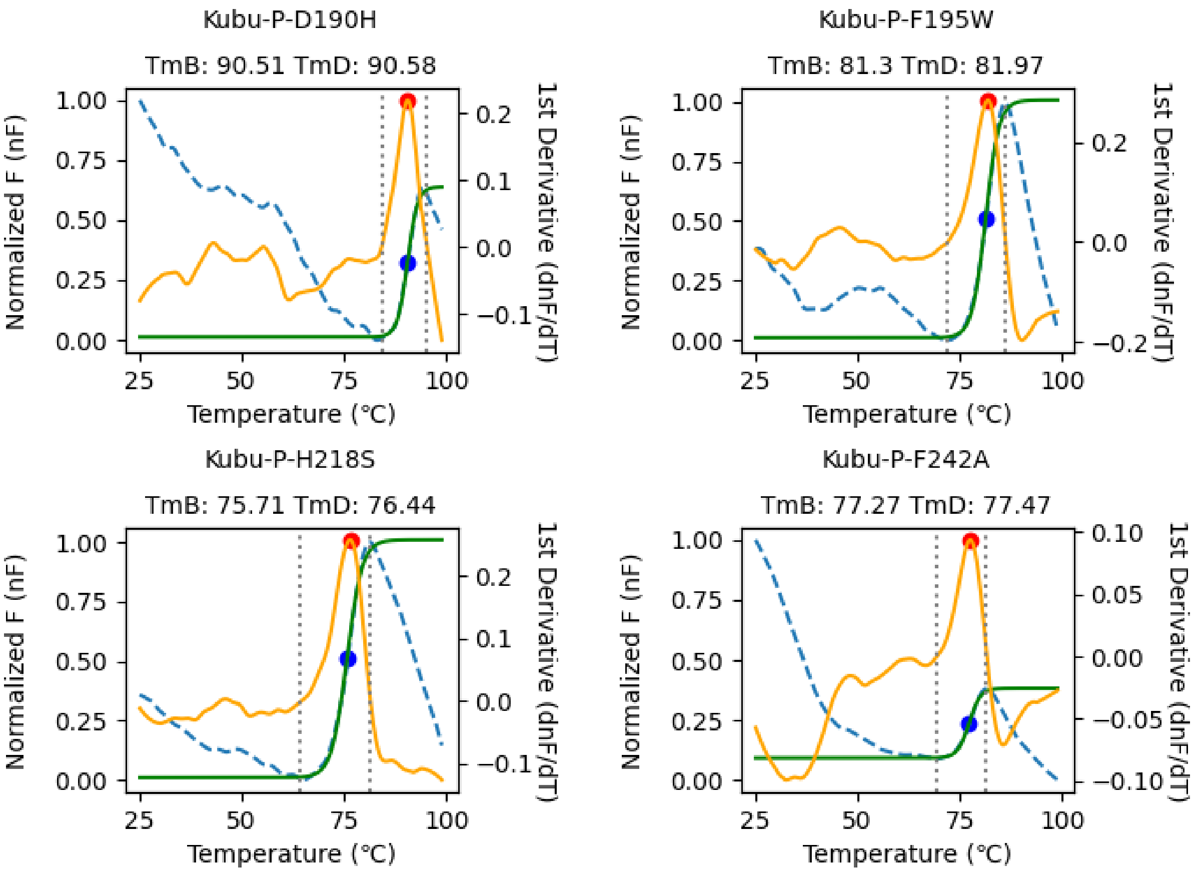


**Fig. S15. Melting temperature (Tm) determination of the structure-based replacement variants through differential scanning fluorimetry (3/3).** The normalized fluorescence (nF) and its 1st derivative is shown with a blue dashed and yellow solid lines, respectively. The Boltzmann curve fitted to the nF is shown with a green solid line. The determined $Tm_B$ and $Tm_D$ is shown with blue and red circles, respectively, and labeled.

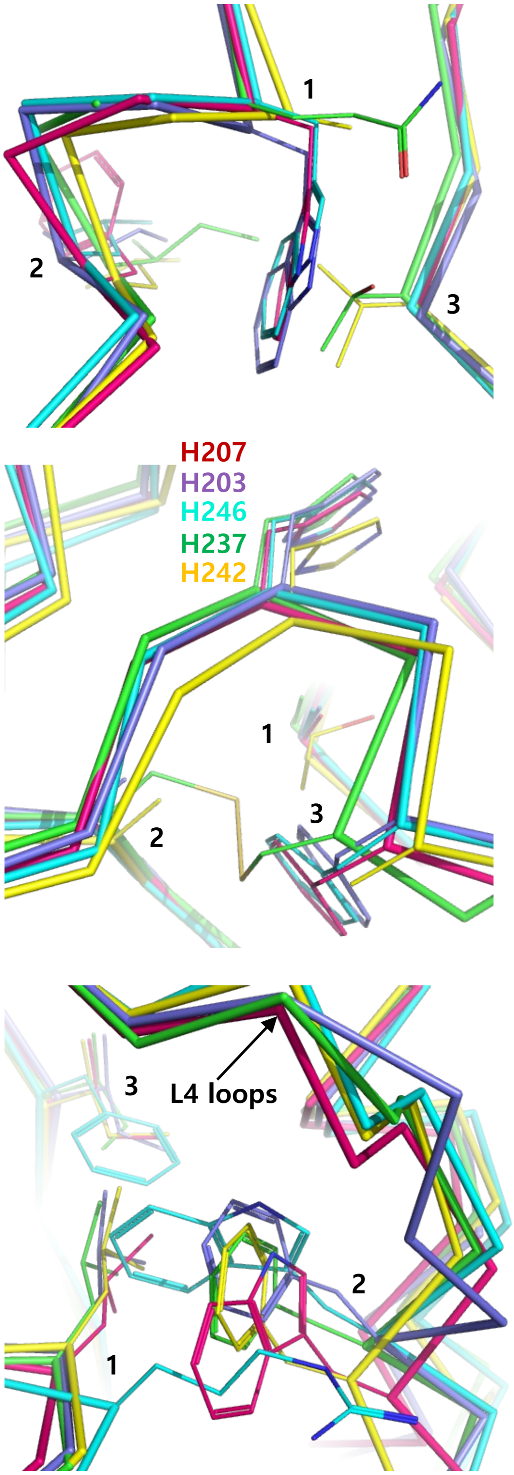


| | 1 2 3 |
|---|---|
| Mipa-P | W102-F105-G121 |
| Kubu-P | W96-I99-G115 |
| CaPETase | W105-L108-G124 |
| IsPETase | Q91-I94-T110 |
| LCC | A99-L102-V118 |

| | 1 2 3 |
|---|---|
| Mipa-P | A191-G211-F247 |
| Kubu-P | A187-A207-F242 |
| CaPETase | A192-G212-F248 |
| IsPETase | A183-C203-C239 |
| LCC | T188-A207-A244 |

| | 1 2 3 |
|---|---|
| Mipa-P | L175-W199-I208 |
| Kubu-P | L169-F195-I204 |
| CaPETase | R176-W200-F209 |
| IsPETase | L167-F191-I200 |
| LCC | L172-F196-I204 |

**Fig. S16. The local networks of CaPETase, and comparison with Mipa-P, Kubu-P, IsPETase, and LCC.** The networks consisting of triple residues are shown. Each residue position is indicated with a number. The distinguishable residual features are indicated with the enzymes' color scheme.

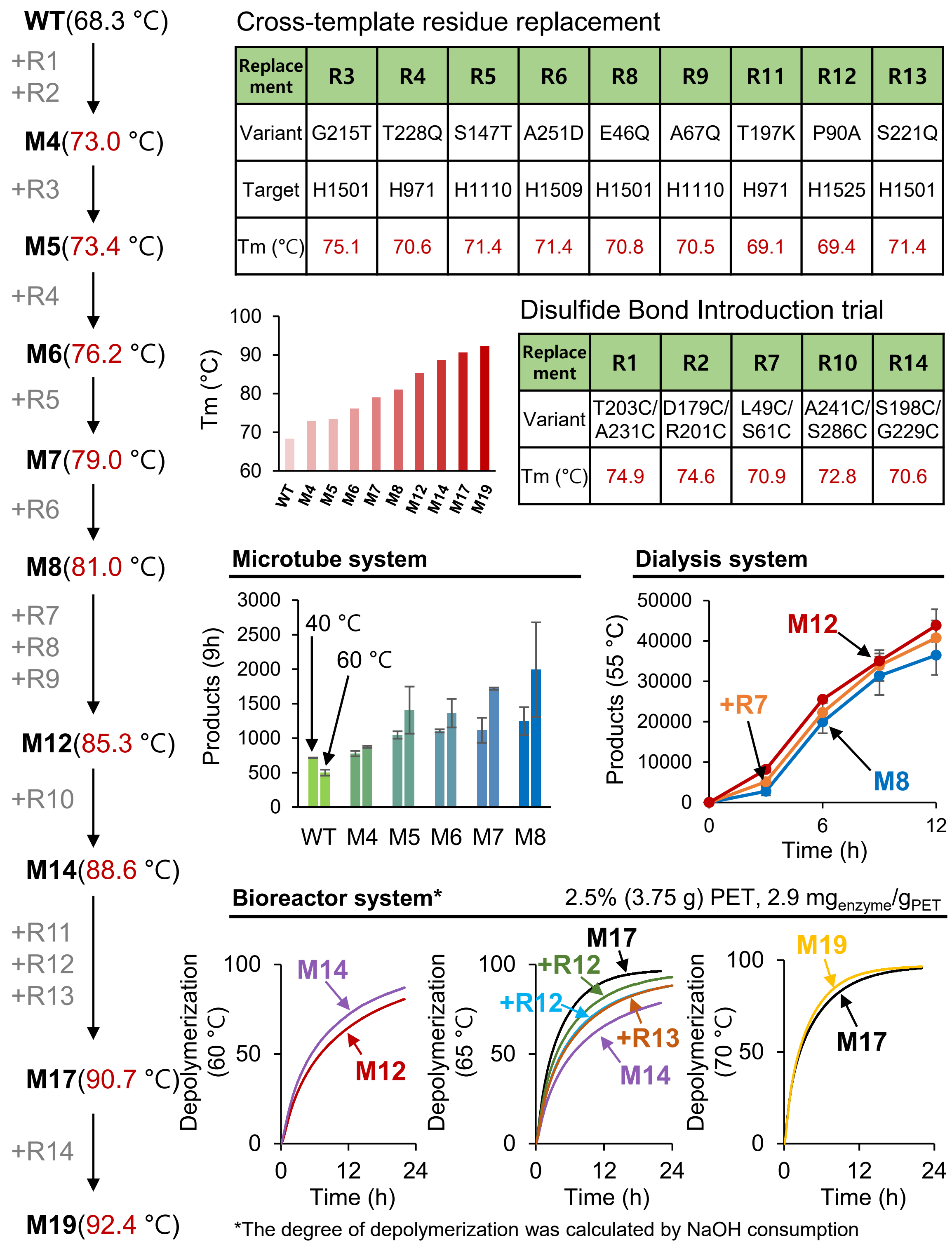


| Replacement | R3 | R4 | R5 | R6 | R8 | R9 | R11 | R12 | R13 |
|---|---|---|---|---|---|---|---|---|---|
| Variant | G215T | T228Q | S147T | A251D | E46Q | A67Q | T197K | P90A | S221Q |
| Target | H1501 | H971 | H1110 | H1509 | H1501 | H1110 | H971 | H1525 | H1501 |
| Tm (°C) | 75.1 | 70.6 | 71.4 | 71.4 | 70.8 | 70.5 | 69.1 | 69.4 | 71.4 |

| Replacement | R1 | R2 | R7 | R10 | R14 |
|---|---|---|---|---|---|
| Variant | T203C/ A231C | D179C/ R201C | L49C/ S61C | A241C/ S286C | S198C/ G229C |
| Tm (°C) | 74.9 | 74.6 | 70.9 | 72.8 | 70.6 |

**Fig. S17. Engineering procedures of Mipa-P$^{M19}$.**

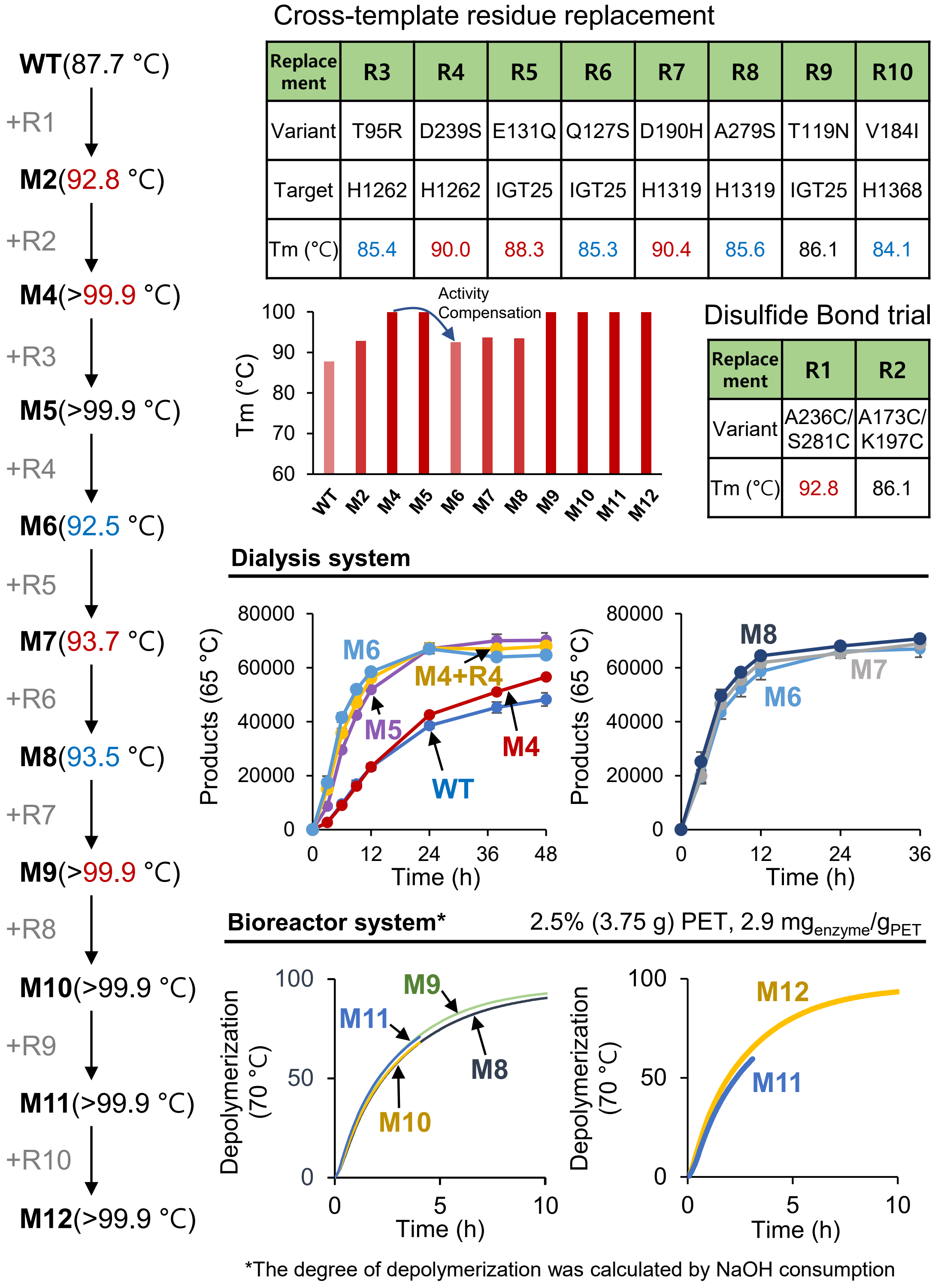


Cross-template residue replacement

| Replacement | R3 | R4 | R5 | R6 | R7 | R8 | R9 | R10 |
|---|---|---|---|---|---|---|---|---|
| Variant | T95R | D239S | E131Q | Q127S | D190H | A279S | T119N | V184I |
| Target | H1262 | H1262 | IGT25 | IGT25 | H1319 | H1319 | IGT25 | H1368 |
| Tm (°C) | 85.4 | 90.0 | 88.3 | 85.3 | 90.4 | 85.6 | 86.1 | 84.1 |

Disulfide Bond trial

| Replacement | R1 | R2 |
|---|---|---|
| Variant | A236C/S281C | A173C/K197C |
| Tm (°C) | 92.8 | 86.1 |

**Fig. S18. Engineering procedures of Kubu-P$^{M12}$.**

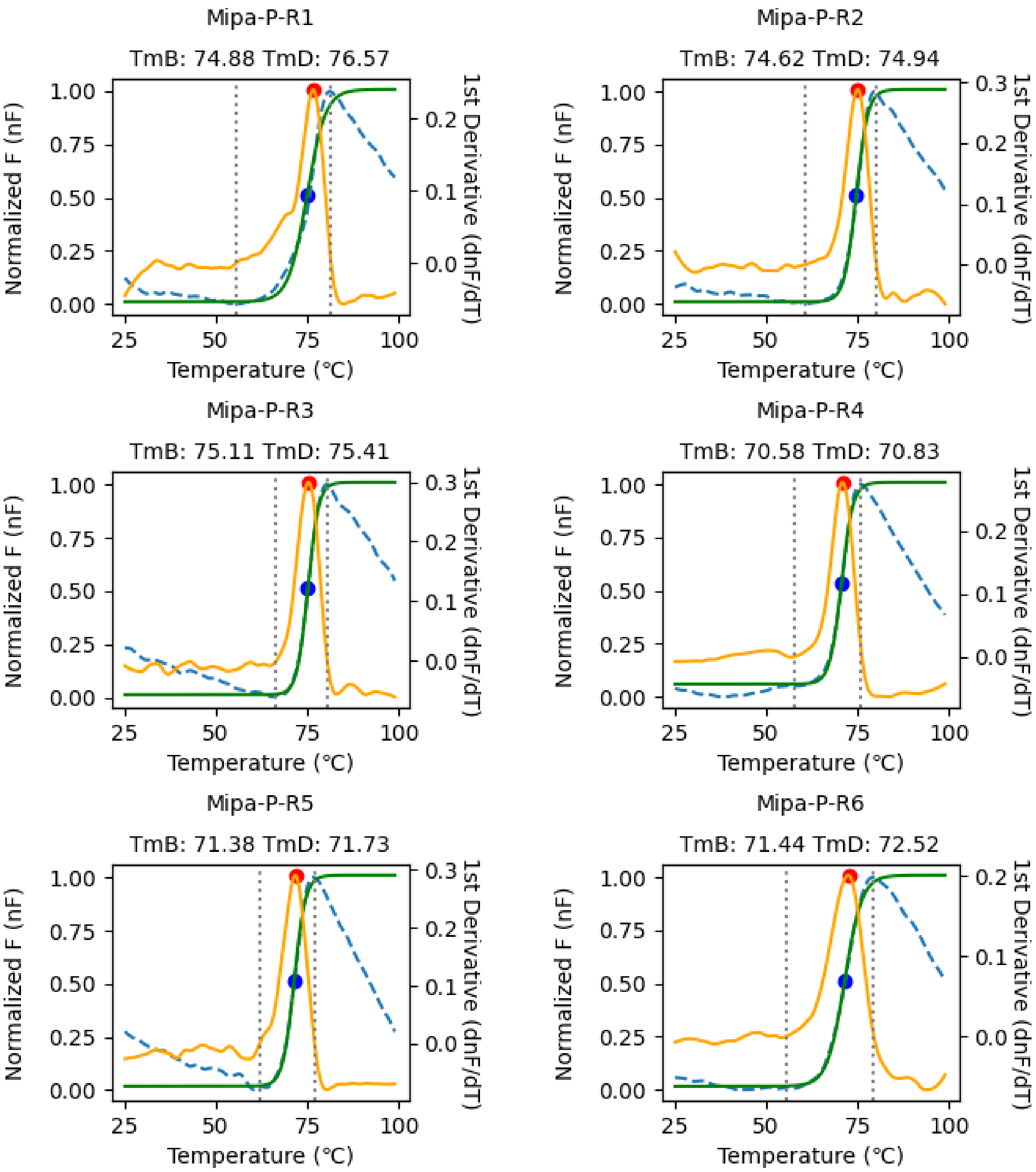


**Fig. S19. Melting temperature (Tm) determination of the engineered variants through differential scanning fluorimetry (continued, 1/9).** The normalized fluorescence (nF) and its 1st derivative is shown with a blue dashed and yellow solid lines, respectively. The Boltzmann curve fitted to the nF is shown with a green solid line. The determined $Tm_B$ and $Tm_D$ is shown with blue and red circles, respectively, and labeled.

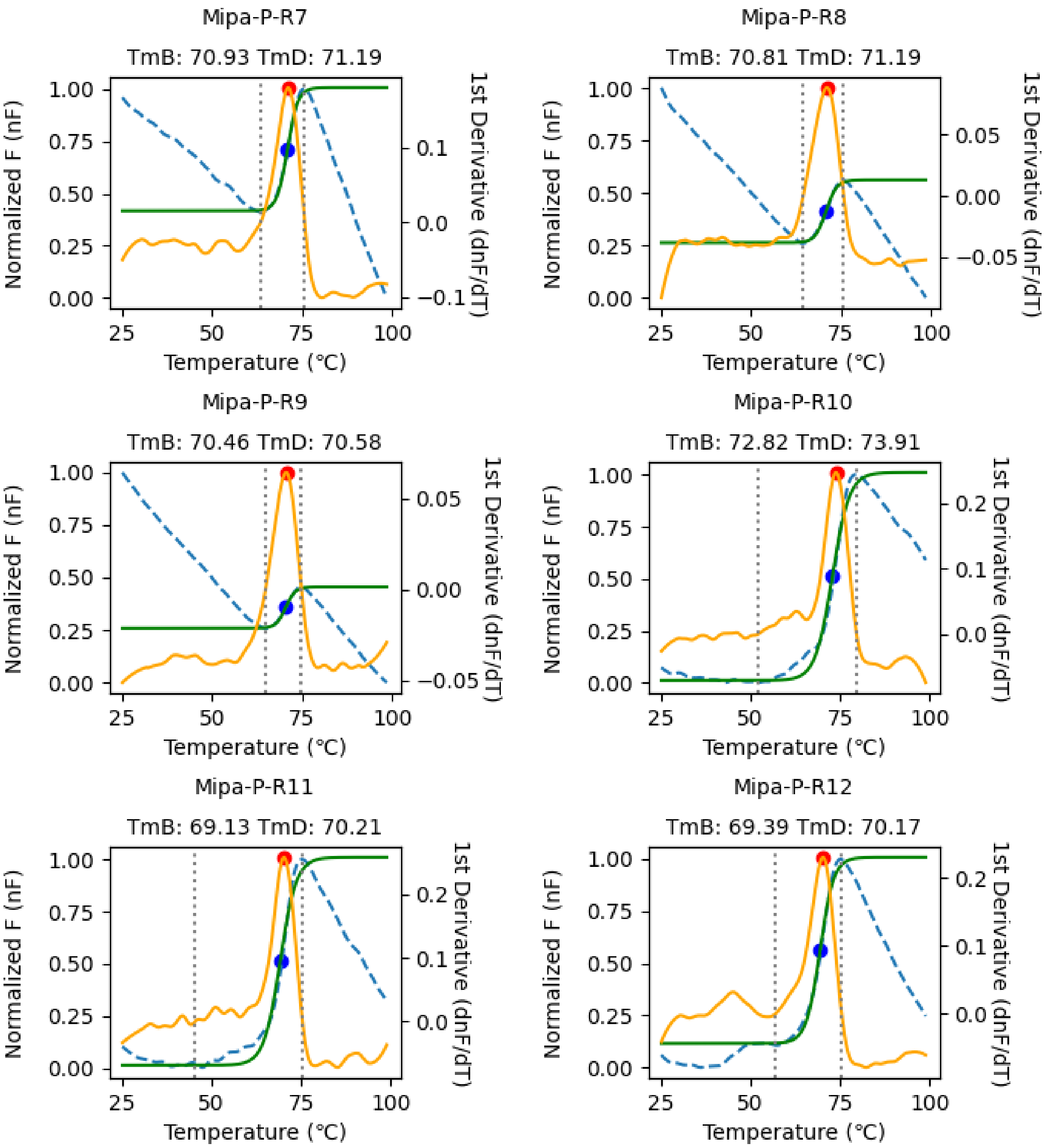


**Fig. S19. Melting temperature (Tm) determination of the engineered variants through differential scanning fluorimetry (continued, 2/9).** The normalized fluorescence (nF) and its 1st derivative is shown with a blue dashed and yellow solid lines, respectively. The Boltzmann curve fitted to the nF is shown with a green solid line. The determined $Tm_B$ and $Tm_D$ is shown with blue and red circles, respectively, and labeled.

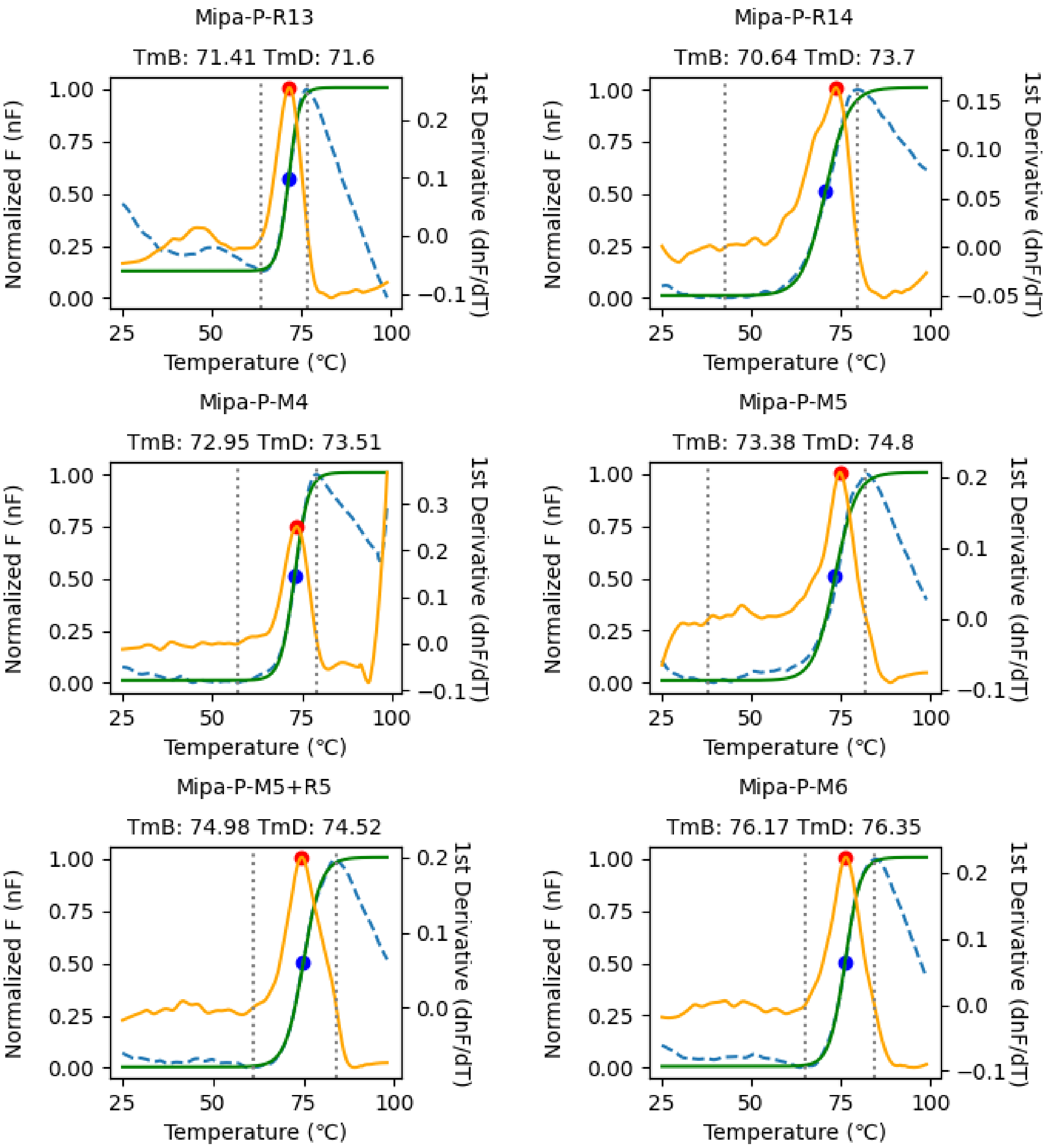


**Fig. S19. Melting temperature (Tm) determination of the engineered variants through differential scanning fluorimetry (continued, 3/9).** The normalized fluorescence (nF) and its 1st derivative is shown with a blue dashed and yellow solid lines, respectively. The Boltzmann curve fitted to the nF is shown with a green solid line. The determined $Tm_B$ and $Tm_D$ is shown with blue and red circles, respectively, and labeled.

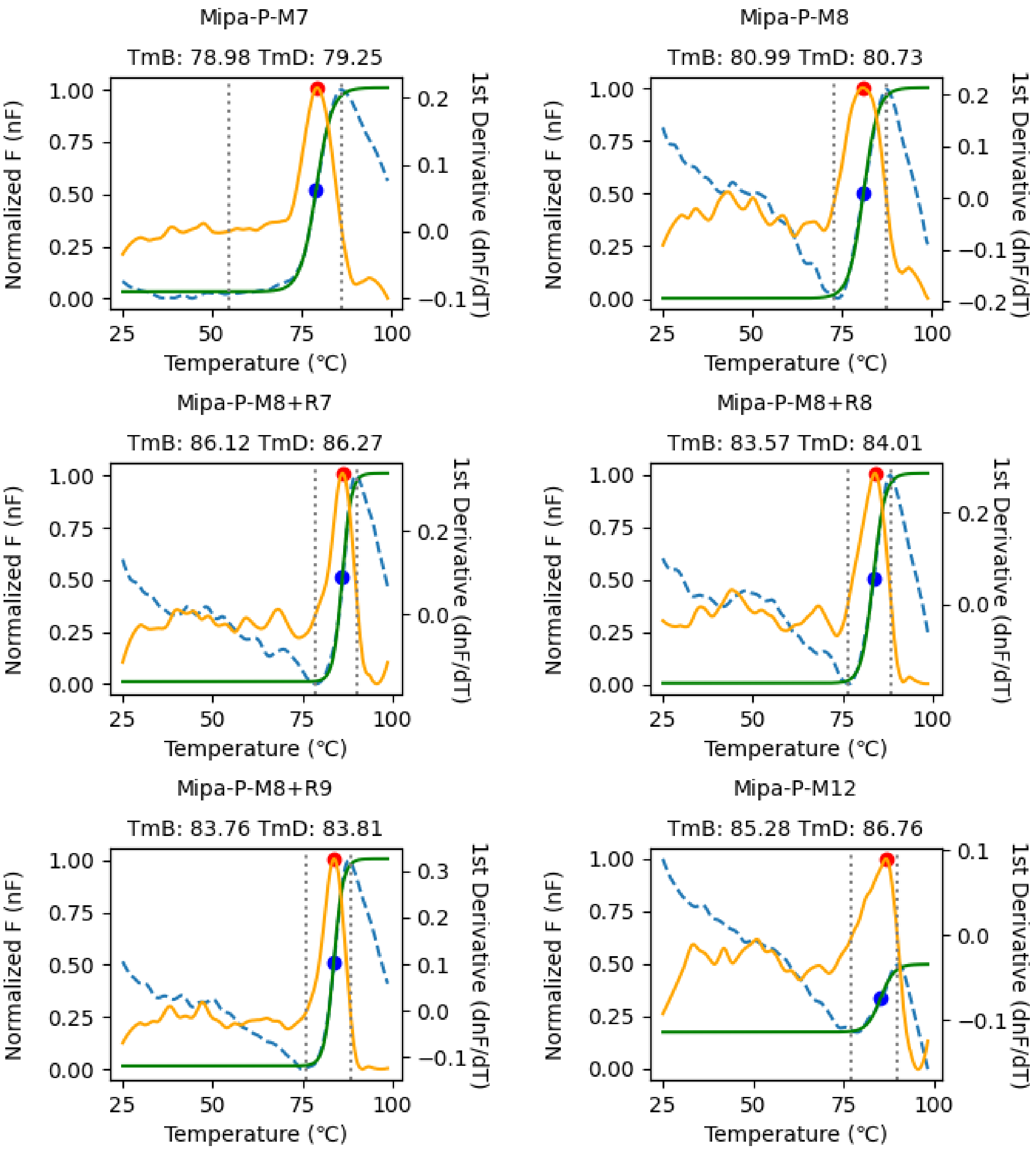


**Fig. S19. Melting temperature (Tm) determination of the engineered variants through differential scanning fluorimetry (continued, 4/9).** The normalized fluorescence (nF) and its 1st derivative is shown with a blue dashed and yellow solid lines, respectively. The Boltzmann curve fitted to the nF is shown with a green solid line. The determined $Tm_B$ and $Tm_D$ is shown with blue and red circles, respectively, and labeled.

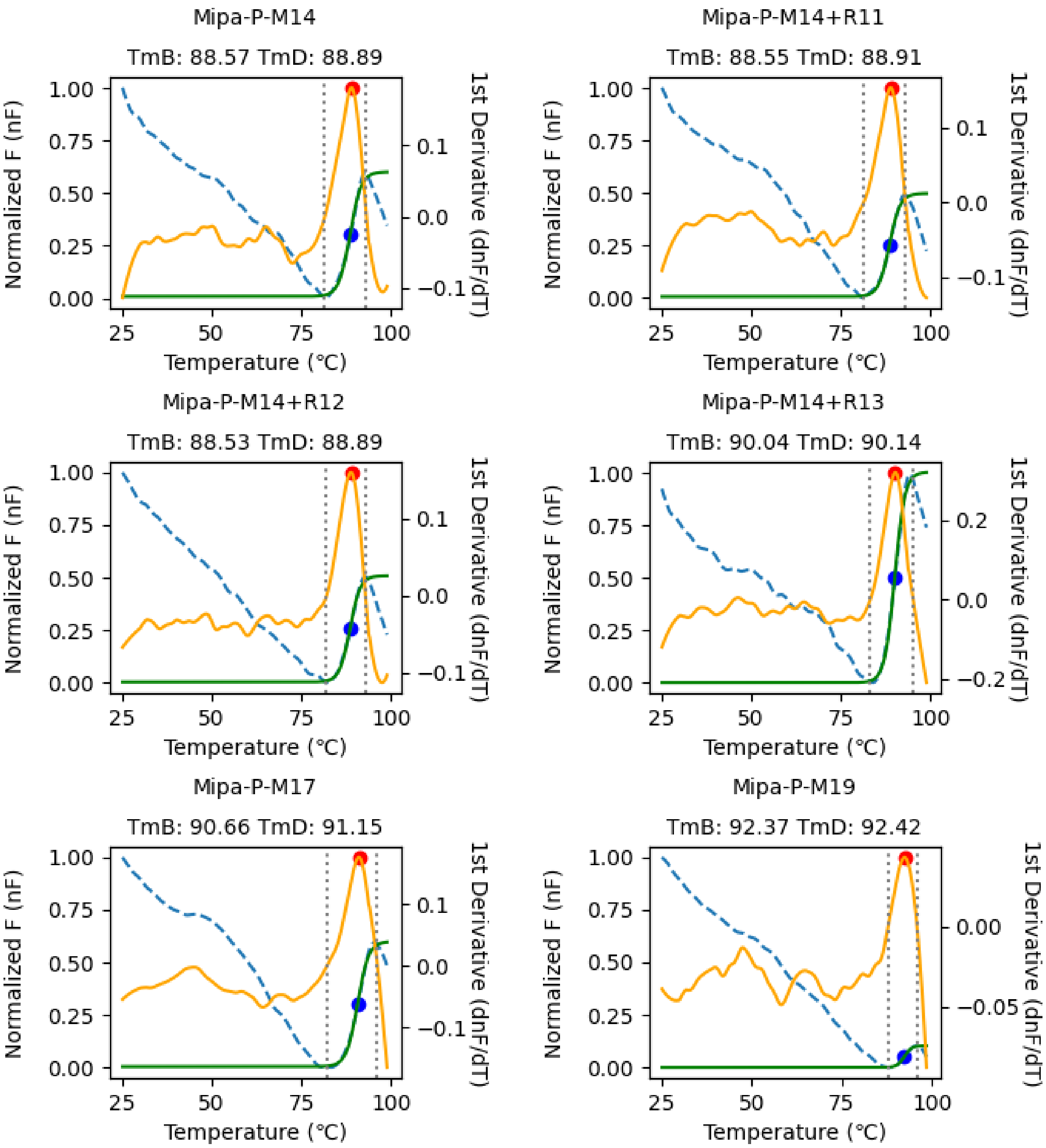


**Fig. S19. Melting temperature (Tm) determination of the engineered variants through differential scanning fluorimetry (continued, 5/9).** The normalized fluorescence (nF) and its 1st derivative is shown with a blue dashed and yellow solid lines, respectively. The Boltzmann curve fitted to the nF is shown with a green solid line. The determined $Tm_B$ and $Tm_D$ is shown with blue and red circles, respectively, and labeled.

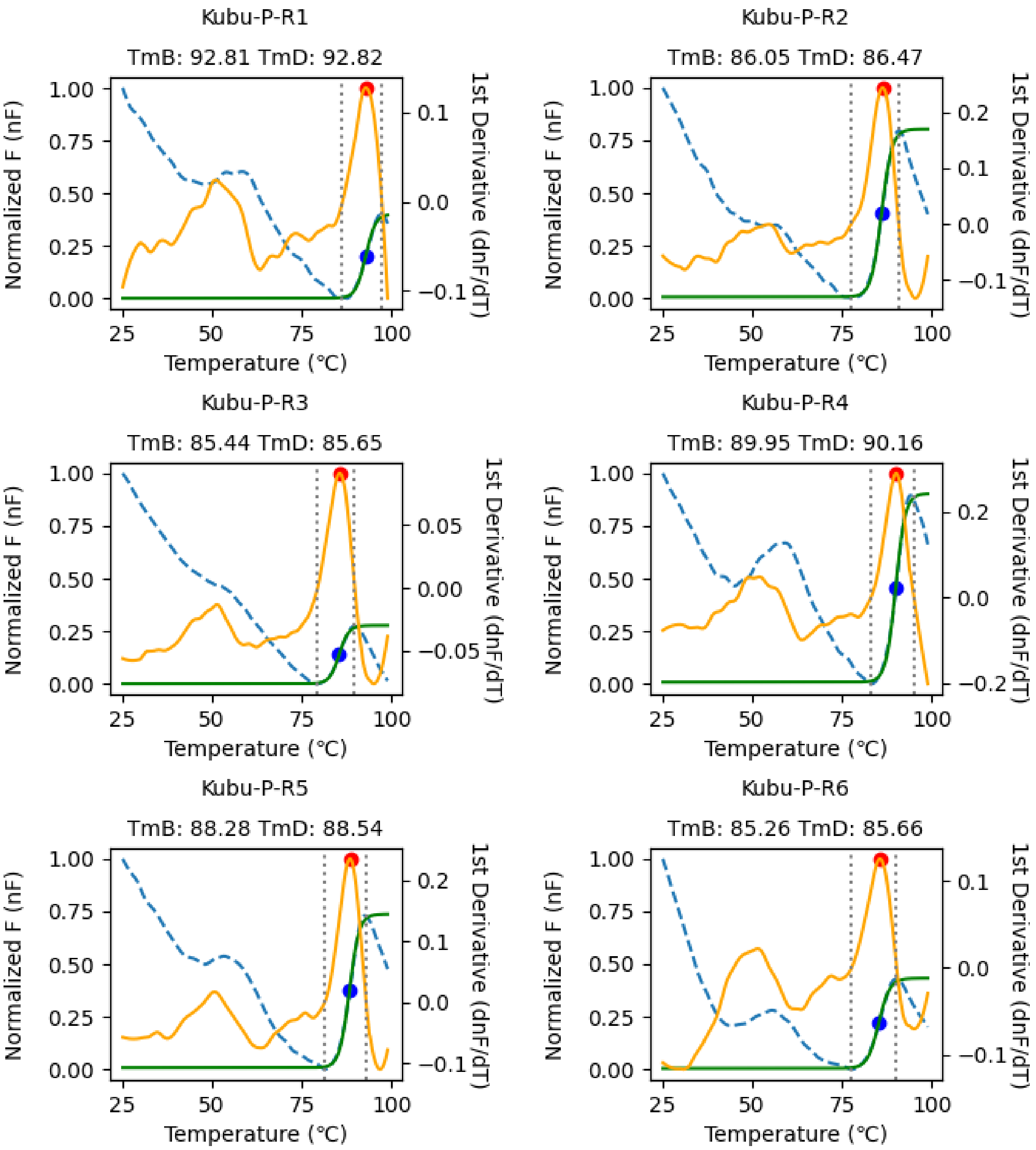


**Fig. S19. Melting temperature (Tm) determination of the engineered variants through differential scanning fluorimetry (continued, 6/9).** The normalized fluorescence (nF) and its 1st derivative is shown with a blue dashed and yellow solid lines, respectively. The Boltzmann curve fitted to the nF is shown with a green solid line. The determined $Tm_B$ and $Tm_D$ is shown with blue and red circles, respectively, and labeled.

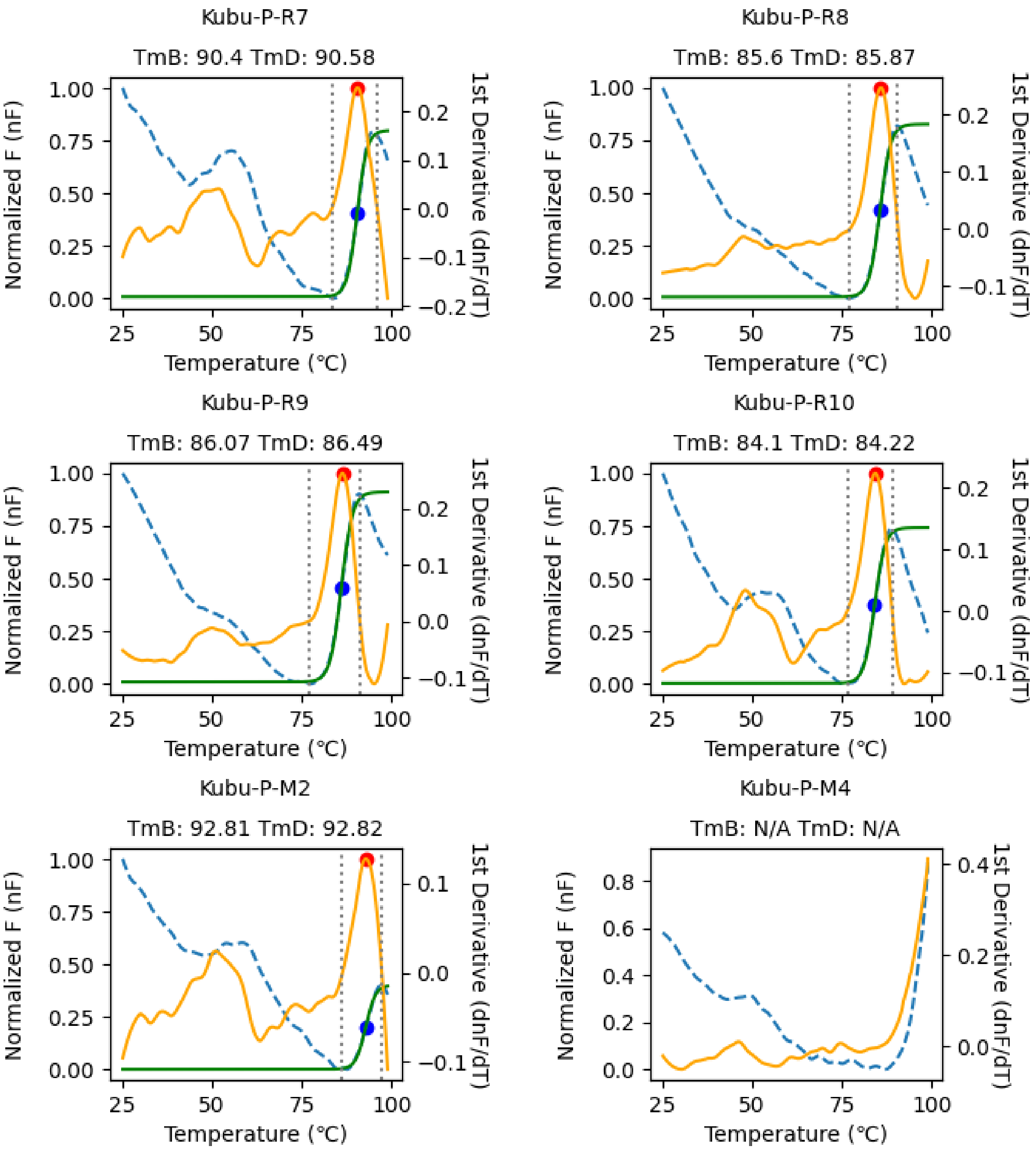


**Fig. S19. Melting temperature (Tm) determination of the engineered variants through differential scanning fluorimetry (continued, 7/9).** The normalized fluorescence (nF) and its 1st derivative is shown with a blue dashed and yellow solid lines, respectively. The Boltzmann curve fitted to the nF is shown with a green solid line. The determined $Tm_B$ and $Tm_D$ is shown with blue and red circles, respectively, and labeled.

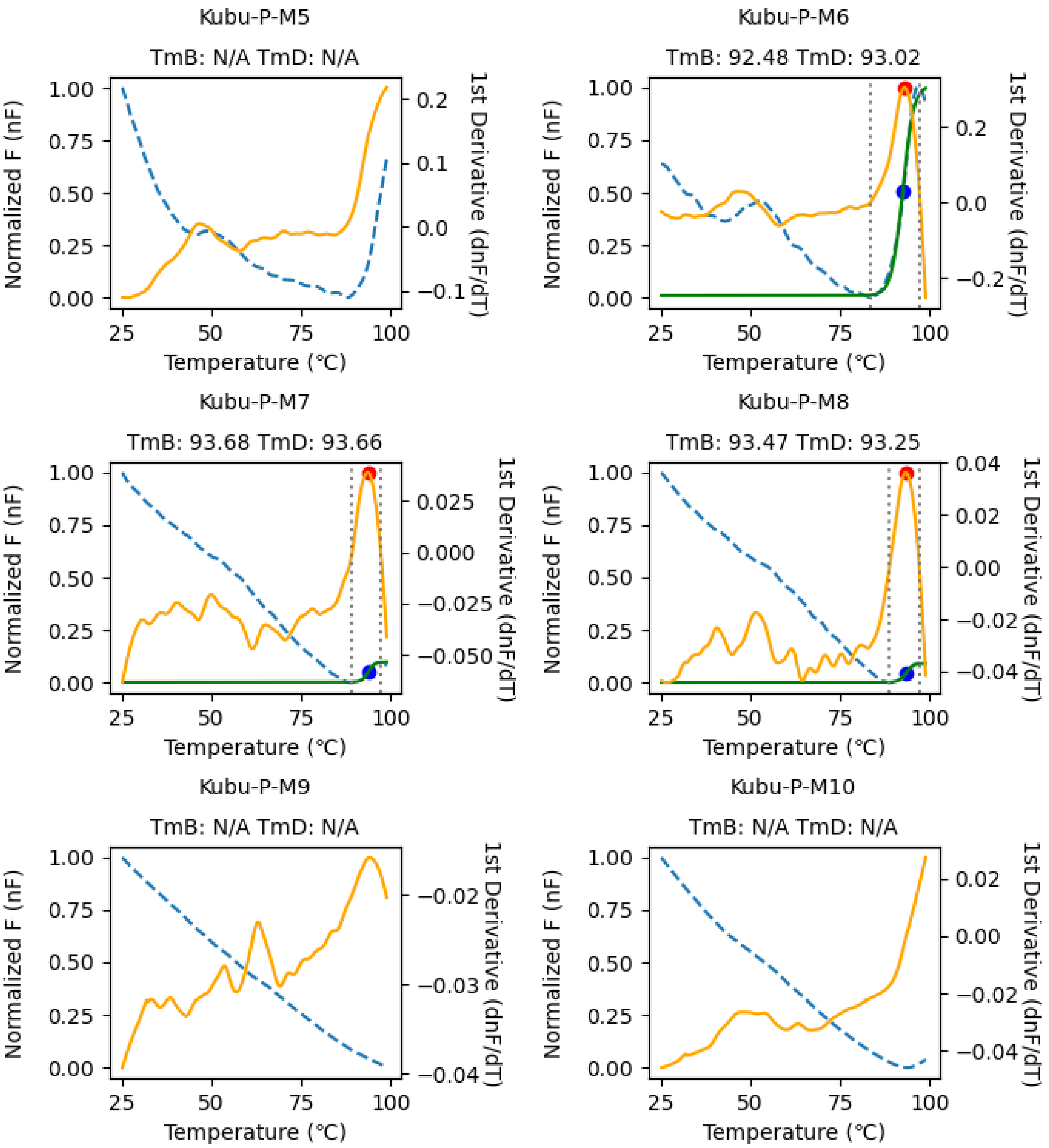


**Fig. S19. Melting temperature (Tm) determination of the engineered variants through differential scanning fluorimetry (continued, 8/9).** The normalized fluorescence (nF) and its 1st derivative is shown with a blue dashed and yellow solid lines, respectively. The Boltzmann curve fitted to the nF is shown with a green solid line. The determined $Tm_B$ and $Tm_D$ is shown with blue and red circles, respectively, and labeled.

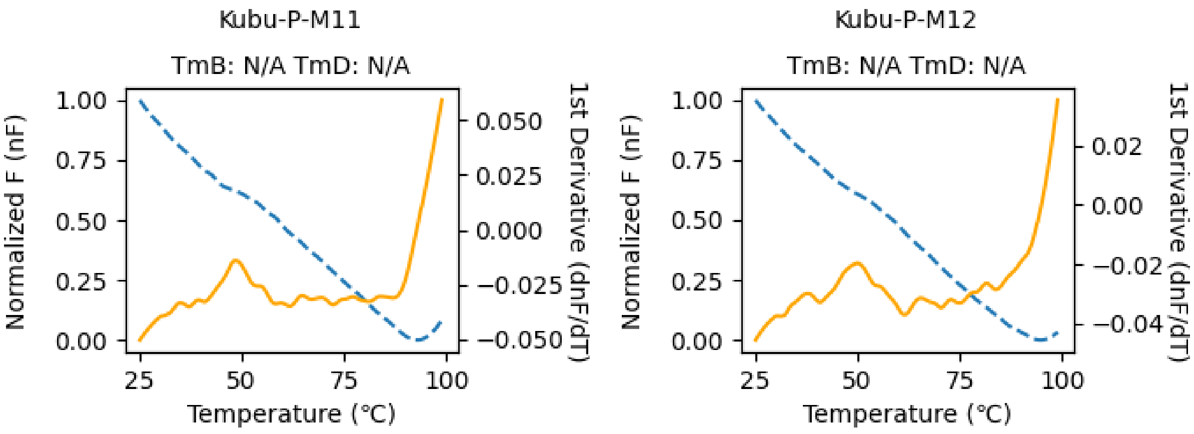


**Fig. S19. Melting temperature (Tm) determination of the engineered variants through differential scanning fluorimetry (9/9).** The normalized fluorescence (nF) and its 1$^{st}$ derivative is shown with a blue dashed and yellow solid lines, respectively. The Boltzmann curve fitted to the nF is shown with a green solid line. The determined $Tm_B$ and $Tm_D$ is shown with blue and red circles, respectively, and labeled.

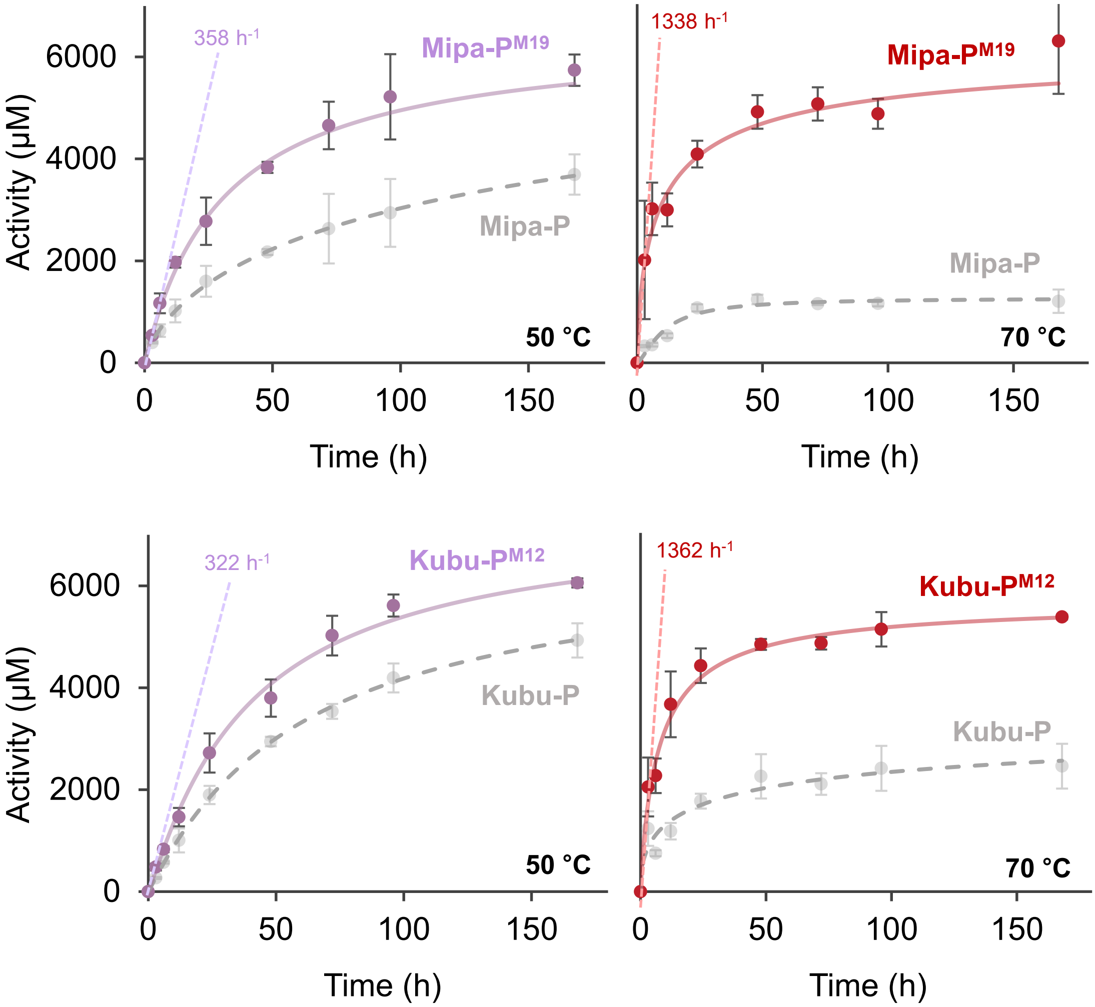


**Fig. S20. Comparisons of Mipa-P$^{M19}$ (A) and Kubu-P$^{M12}$ (B) with their wild-types.**

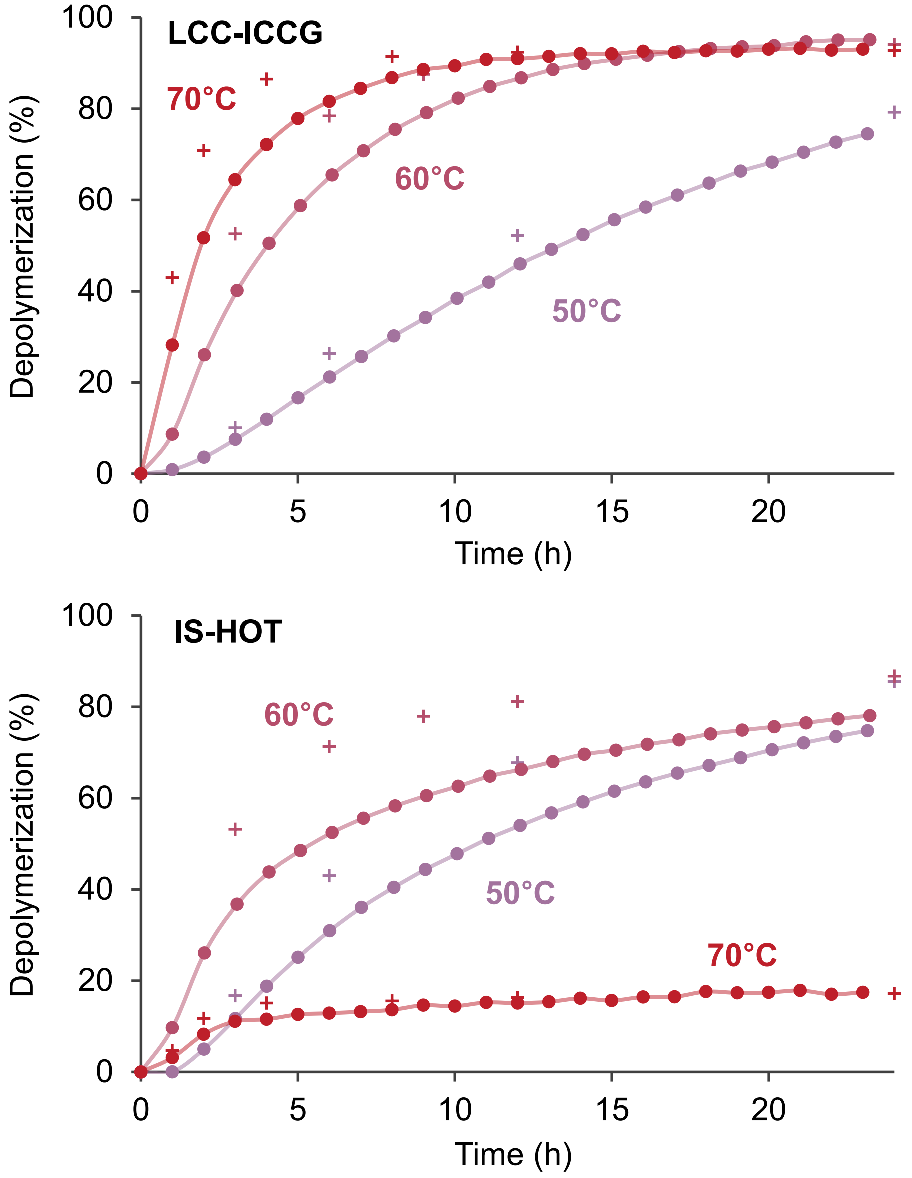


**Fig. S21. Depolymerizations in mild condition by LCC-ICCG and Hot-PETase in pH-stat bioreactor.** The graphs of linked circles or crosses indicate degree of depolymerization calculated by monomer release (HPLC sampling) or ester bond cleavage (NaOH consumption), respectively.

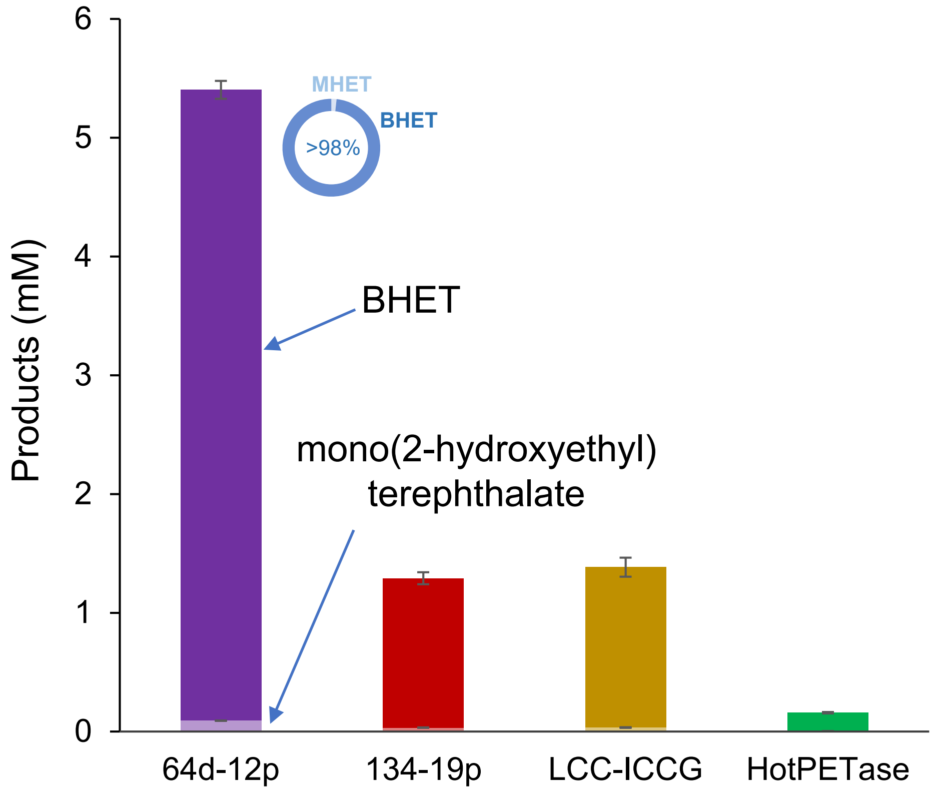


**Fig. S22. Monomer production by 1 d enzyme-catalyzed PET glycolysis under EG (95%) solvent with 4.5% lysis buffer.** The boxes of strongest color for each enzyme indicate BHET concentration, and those with the light color scheme show mono(2-hydroxyethyl) terephthalate concentration. Standard deviations are shown with bars.

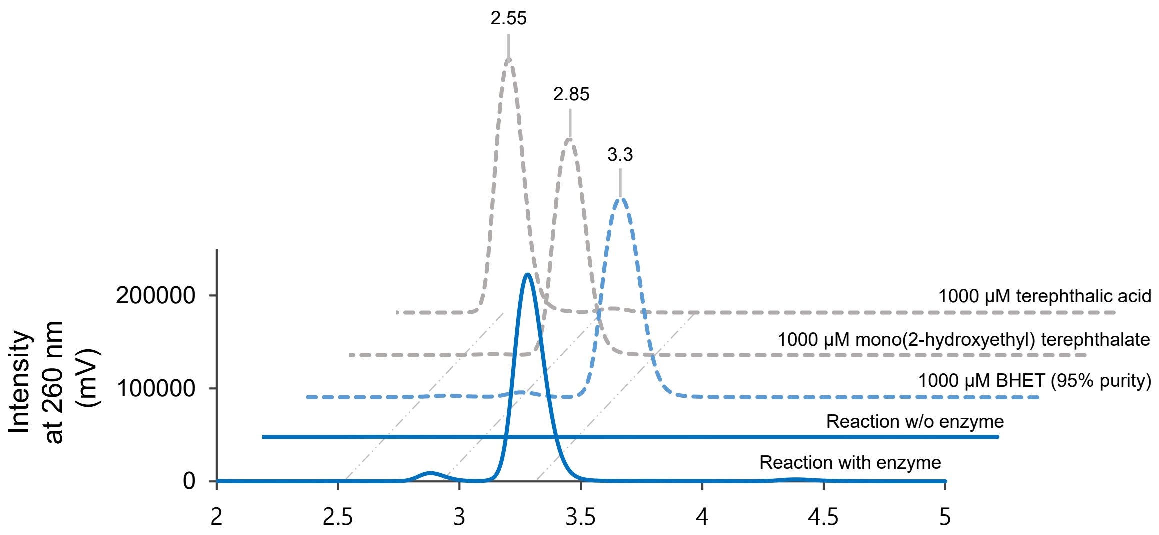


**Fig. S23. HPLC chromatograms in enzyme-catalyzed PET glycolysis.**

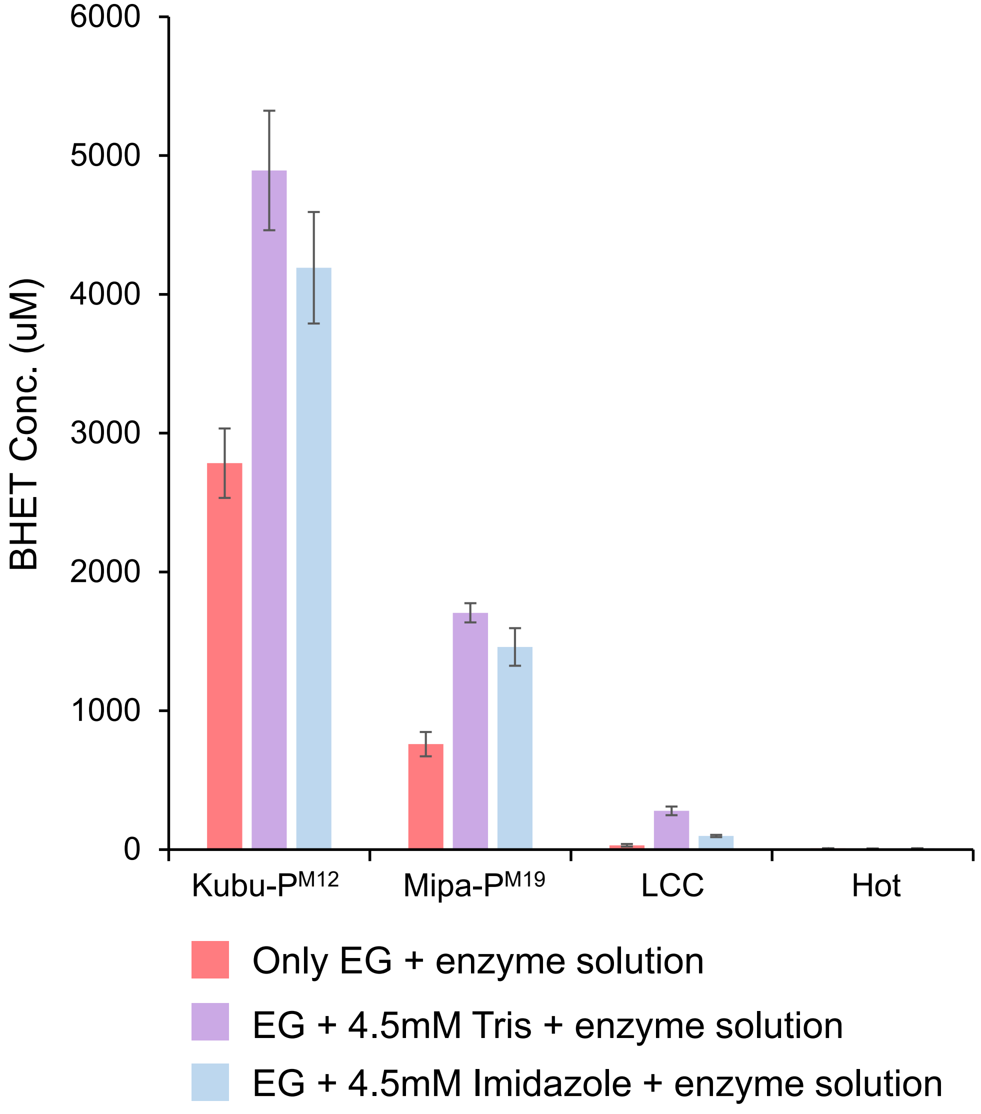


**Fig. S24. Buffer effect in enzyme-catalyzed PET glycolysis.**

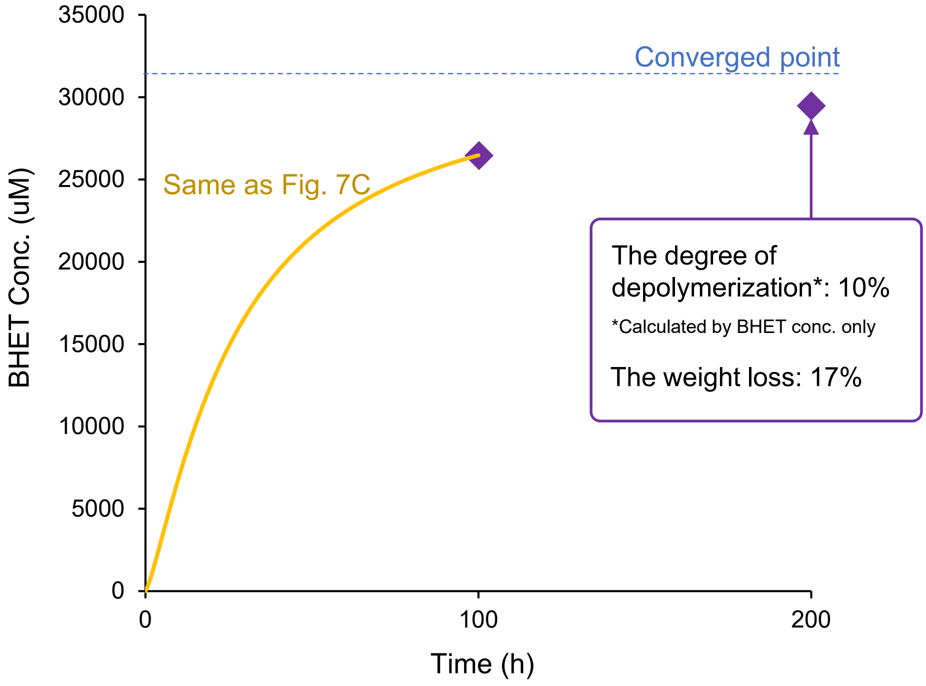


**Fig. S25. Enzyme-catalyzed PET glycolysis by Kubu-P$^{M12}$ at 40 °C.**

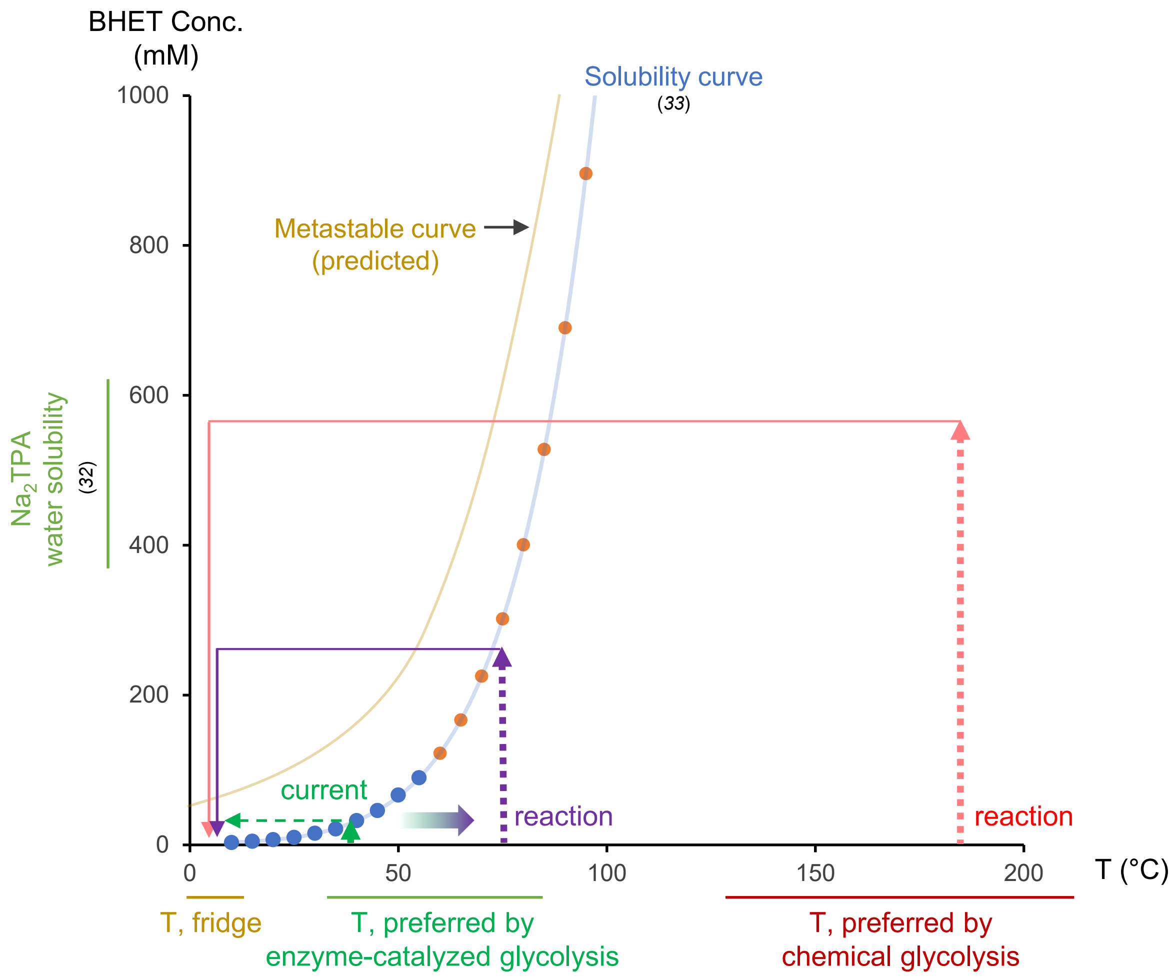


**Fig. S26. BHET recovery process and solubility problem.** The Na2TPA solubility from the reference 32 is shown. The solubility curve of BHET in EG is drawn with the blue (experimental) and orange (ideal solution model) circles in the reference 33. The dashed lines indicate BHET concentration increase during the reaction.

### Tables. S1 to S5

### Table S1. Search thresholds for each seed.

| Seed sequences | | | threshold | Ref. |
|---|---|---|---|---|
| Accession code | name | EPH Library forming | value | |
| XP_001817153 | AoC | X | 4e-22 | (*34*) |
| ADH43200.1 | BsEstB | X | 1e-85 | (*35*) |
| CAH17554.1 | BTA2 | O | 5e-33 | (*36*) |
| WP_015787089.1 | Cut190 | O | 2e-33 | (*37*) |
| BAI99230.2 | Est1 | O | 1e-23 | (*38*) |
| BAK48590.1 | Est2 | O | 1e-40 | (*38*) |
| EGU74658.1 | FoCut5a | X | 1e-18 | (*39*) |
| AAA33335.1 | FsC | X | 7e-20 | (*40*) |
| ASK40094.1 | HiC | X | 2e-16 | (*41*) |
| AEV21261.1 | LCC | O | 5e-34 | (*42*) |
| CCK74972.1 | PET5 | O | 1e-28 | (*43*) |
| ASA57064.1 | PET6 | O | 1e-30 | (*43*) |
| WP_047194864.1 | PET12 | O | 3e-36 | (*43*) |
| WP_054022242.1 | IsPETase | O | 2e-35 | (*44*) |
| WP_004373894.1 | PmC | X | 1e-30 | (*45*) |
| WP_012850775.1 | Tcur0390 | O | 1e-30 | (*46*) |
| PKK15152.1 | Tcur1278 | O | 1e-29 | (*46*) |
| P86325.1 | TfCa | X | 1e-64 | (*47*) |
| CBY05529.1 | TfCut1 | O | 6e-36 | (*36*) |
| PZN61876.1 | TfCut2 | O | 2e-22 | (*48*) |
| WP_011291330.1 | TfH | O | 5e-42 | (*36*) |
| AAZ54920.1 | Tfu_0882 | O | 5e-24 | (*36*) |
| WP_011291330.1 | Thc_Cut1 | O | 4e-37 | (*49*) |
| WP_016188393.1 | Thc_Cut2 | O | 7e-25 | (*49*) |
| WP_104613137.1 | Thf42_Cut1 | O | 3e-34 | (*50*) |
| AFA45122.1 | Thh_Est | O | 1e-40 | (*51*) |

### Tables S2. The statistics of the collected data and refinement.

| | | Mipa-P | Mipa-P$^{M19}$ | Kubu-P | Kubu-P$^{P185V}$ | Kubu-P$^{M12}_{+P185V}$ |
|---|---|---|---|---|---|---|
| **Data collection** | | | | | | |
| | Space group | $P2_12_12_1$ | $P2_1$ | $I4_132$ | $I4_132$ | $I222$ |
| Unit cell Param. | a, b, c (Å) | 49.07, 103.41, 131.88 | 42.62, 51.36, 107.65 | 159.10, 159.10, 159.10 | 157.22, 157.22, 157.22 | 64.96, 81.65, 83.73 |
| | α, β, γ (°) | 90.0, 90.0, 90.0 | 90, 94.3, 120.0 | 90.0, 90.0, 90.0 | 90.0, 90.0, 90.0 | 90.0, 90.0, 90.0 |
| | Resolution (Å) | 36.80-1.34 | 42.50-1.90 | 45.88-2.65 | 42.03-1.70 | 32.50-1.15 |
| | $R_{sym}$ or $R_{merge}$ | 12.5(36.4) | 8.2(32.7) | 12.3(32.8) | 8.1(30.6) | 13.5(29.5) |
| | I of sigma | 7.4(2.6) | 7.9(2.9) | 26.1(10.1) | 28.0(3.5) | 14.8(3.2) |
| | Completeness | 96.1(91.6) | 99.7(99.2) | 99.9(99.9) | 99.9(99.9) | 97.9(96.2) |
| | Redundancy | 2.6(2.5) | 8.6(4.6) | 14.9(11.4) | 4.8(3.1) | 14.5(13.0) |
| **Refinement** | | | | | | |
| | Resolution | 36.80-1.34 | 38.56-1.90 | 45.88-2.65 | 39.34-1.70 | 32.50-1.15 |
| | No. reflections | 136706 | 35259 | 10243 | 34762 | 74333 |
| | $R_{work}$ / $R_{free}$ | 21.1 / 23.4 | 18.8 / 24.1 | 21.7 / 28.3 | 16.1 / 19.5 | 16.0 / 17.8 |
| | *B*-factors | 25.4 | 31.8 | 60.3 | 27.0 | 8.9 |
| | Protein | 24.6 | 33.5 | 60.3 | 28.1 | 9.3 |
| | Water | 34.7 | 42.5 | - | 47.8 | 24.4 |
| | R.M.S.D | | | | | |
| | Bond lengths (Å) | 0.007 | 0.006 | 0.011 | 0.013 | 0.014 |
| | Bond angles (°) | 1.446 | 1.290 | 1.637 | 1.635 | 1.838 |

**Tables S3. The layout parameters and edge weight settings in Cytoscape.**

| parameter | | Value |
|---|---|---|
| Edge weight settings | The min edge weight to consider | 1E-1 |
| | The max edge weight to consider | 1E0 |
| | The default edge weight to consider | 1 |
| Layout parameters | numIterations | 100 |
| | numIterationsEdgeRepulsive | 5E1 |
| | defaultSpringCoefficient | 5E-5 |
| | defaultSpringLength | 10 |
| | defaultNodeMass | 3 |
| | isDeterministic | True |
| | fromScratch | False |
| | singlePartition | False |

**Tables S4. The primers used in this study (continued, 1/4).**

| Enz. | Category | Dir. | Sequence |
|---|---|---|---|
| Mipa-P$^{W102A}$ | Structure-based | F | 5’-GTTATACGGAGCGTGCGAGCAGTTTTGCATGG-3’ |
| | | R | 5’-CCATGCAAAACTGCTCGCACGCTCCGTATAAC-3’ |
| Mipa-P$^{W102Q}$ | Structure-based | F | 5’-GTTATACGGAGCGTCAGAGCAGTTTTGCATGG-3’ |
| | | R | 5’-CCATGCAAAACTGCTCTGACGCTCCGTATAAC-3’ |
| Mipa-P$^{H167W}$ | Structure-based | F | 5’-CAAGGCGTAAGTGGCTGGAGCATGGGTGGCGG-3’ |
| | | R | 5’-CACCGCCACCCATGCTCCAGCCACTTACGCCTTG-3’ |
| Mipa-P$^{H194D}$ | Structure-based | F | 5’-CATTAGCACCTTGGGATACGACTACTAGCTG-3’ |
| | | R | 5’-CAGCTAGTAGTCGTATCCCAAGGTGCTAATG-3’ |
| Mipa-P$^{W199F}$ | Structure-based | F | 5’-CATACGACTACTAGCTTTCCCCGCGTCACCAATC-3’ |
| | | R | 5’-GATTGGTGACGCGGGGAAAGCTAGTAGTCGTATG-3’ |
| Mipa-P$^{H222S}$ | Structure-based | F | 5’-CCTGTCTCTAGCAGCGCAATTCCAATGTATACG-3’ |
| | | R | 5’-CGTATACATTGGAATTGCGCTGCTAGAGACAGG-3’ |
| Mipa-P$^{F247A}$ | Structure-based | F | 5’-CGGGCCATAATGCTCCGAATTCGGCGAATCCC-3’ |
| | | R | 5’-GGGATTCGCCGAATTCGGAGCATTATGGCCCG-3’ |
| Mipa-P$^{E100S}$ | Structure-based | F | 5’-GCCCGGGTTATACGTCGCGTTGGAGCAGTTTTG-3’ |
| | | R | 5’-CAAAACTGCTCCAACGCGACGTATAACCCGGGC-3’ |
| Mipa-P$^{E100A}$ | Structure-based | F | 5’-CCCGGGTTATACGGCGCGTTGGAGCAGTTT-3’ |
| | | R | 5’-AAACTGCTCCAACGCGCCGTATAACCCGGG-3’ |
| Kubu-P$^{W96A}$ | Structure-based | F | 5’-GGTTTTGTAAGCACAGCGTCACAAATTAGTTGGC-3’ |
| | | R | 5’-GCCAACTAATTTGTGACGCTGTGCTTACAAAACC-3’ |
| Kubu-P$^{W96Q}$ | Structure-based | F | 5’-GGTTTTGTAAGCACACAGTCACAAATTAGTTGGC-3’ |
| | | R | 5’-GCCAACTAATTTGTGACTGTGTGCTTACAAAACC-3’ |
| Kubu-P$^{W161H}$ | Structure-based | F | 5’-CGCGGTGTAGCTGGTCACAGCATGGGTGGTGGT-3’ |
| | | R | 5’-ACCACCACCCATGCTGTGACCAGCTACACCGC-3’ |
| Kubu-P$^{D190H}$ | Structure-based, R7 | F | 5’-CGCTCGCACCGTGGCATATTGGTCAAGATTTTAG-3’ |
| | | R | 5’-CTAAAATCTTGACCAATATGCCACGGTGCGAGCG-3’ |
| Kubu-P$^{F195W}$ | Structure-based | F | 5’-GATATTGGTCAAGATTGGAGTAAAGTGACGAAAC-3’ |
| | | R | 5’-GTTTCGTCACTTTACTCCAATCTTGACCAATATC-3’ |
| Kubu-P$^{H218S}$ | Structure-based | F | 5’-CCACCGGCTCAATCTGCGGTTCCGTTTTATAATG-3’ |
| | | R | 5’-CATTATAAAACGGAACCGCAGATTGAGCCGGTGG-3’ |
| Kubu-P$^{F242A}$ | Structure-based | F | 5’-GGTGCGGATCATTTCGCTCCCACGACTGCTAATC-3’ |
| | | R | 5’-GATTAGCAGTCGTGGGAGCGAAATGATCCGCACC-3’ |

**Tables S4. The primers used in this study (continued, 2/4).**

| Enz. | Category | Dir. | Sequence |
|---|---|---|---|
| Mipa-$P^{T203C}$ | R1 component | F | 5'-CTGGCCCCGCGTCTGCAATCCAGTAATGATTTTG -3' |
| | | R | 5'-CAAAATCATTACTGGATTGCAGACGCGGGGCCAG-3' |
| Mipa-$P^{A231C}$ | R1 component | F | 5'-AATGTATACGGGTGTATGCAGTGGGGAGAAAGC-3' |
| | | R | 5'-CTTTCTCCCCACTGCATACACCCGTATACATT-3' |
| Mipa-$P^{D179C}$ | R2 component | F | 5'-CCCTTAGCGCGATGTGTCAACGTCCGAGCGTAC-3' |
| | | R | 5'-GTACGCTCGGACGTTGACACATCGCGCTAAGGG-3' |
| Mipa-$P^{R201C}$ | R2 component | F | 5'-CTACTAGCTGGCCCTGCGTCACCAATCCAGTAA-3' |
| | | R | 5'-TTACTGGATTGGTGACGCAGGGCCAGCTAGTAG-3' |
| Mipa-$P^{G215T}$ | R3 component | F | 5'-GTGGTCAAAATGATACCATTGCCCCTGTCTC-3' |
| | | R | 5'-GAGACAGGGGCAATGGTATCATTTTGACCAC-3' |
| Mipa-$P^{T228Q}$ | R4 component | F | 5'-GCAATTCCAATGTATCAGGGTGTAGCGAGTG-3' |
| | | R | 5'-CACTCGCTACACCCTGATACATTGGAATTGC-3' |
| Mipa-$P^{S147T}$ | R5 component | F | 5'-CTCGATTGGGCGTCAACAAGCGCGCCTGCGGCAG-3' |
| | | R | 5'-CTGCCGCAGGCGCGCTTGTTGACGCCCAATCGAG-3' |
| Mipa-$P^{A251D}$ | R6 component | F | 5'-ATAATTTTCCGAATTCGGACAATCCCATTGTATC -3' |
| | | R | 5'-GATACAATGGGATTGTCCGAATTCGGAAAATTAT-3' |
| Mipa-$P^{L49C}$ | R7 component | F | 5'-GCAACGGAACGTGGCTGTGCACCAACGGCAGC-3' |
| | | R | 5'-GCTGCCGTTGGTGCACAGCCACGTTCCGTTGC-3' |
| Mipa-$P^{S61C}$ | R7 component | F | 5'-TTACGGGTGATGGTTGTTATGGTGTAGTAAGTG-3' |
| | | R | 5'-CACTTACTACACCATAACAACCATCACCCGTAA-3' |
| Mipa-$P^{E46Q}$ | R8 component | F | 5'-CCAGCAAGTGCAACGCAACGTGGCTTAGCACC-3' |
| | | R | 5'-GGTGCTAAGCCACGTTGCGTTGCACTTGCTGG-3' |
| Mipa-$P^{A67Q}$ | R9 component | F | 5'-CTTATGGTGTAGTAAGTCAAACTATTACGGGTG-3' |
| | | R | 5'-CACCCGTAATAGTTTGACTTACTACACCATAAG -3' |
| Mipa-$P^{A241C}$ | R10 component | F | 5'-GCCTATGTGGAACTCTGCGGTGCGGGCCATAAT-3' |
| | | R | 5'-ATTATGGCCCGCACCGCAGAGTTCCACATAGGC-3' |
| Mipa-$P^{S286C}$ | R10 component | F | 5'-GGTGGTGCGAGCATCTGCCAATTTCGTAGTACG-3' |
| | | R | 5'-CGTACTACGAAATTGGCAGATGCTCGCACCACC-3' |
| Mipa-$P^{T197K}$ | R11 component | F | 5'-CCTTGGCATACGACTAAAAGCTGGCCCCGCGTC-3' |
| | | R | 5'-GACGCGGGGCCAGCTTTTAGTCGTATGCCAAGG-3' |
| Mipa-$P^{P90A}$ | R12 component | F | 5'-CTACGGAGCGTTTTGCGGTGGTTGCAATTAG-3' |
| | | R | 5'-CTAATTGCAACCACCGCAAAACGCTCCGTAG-3' |

**Tables S4. The primers used in this study (continued, 3/4).**

| Enz. | Category | Dir. | Sequence |
|---|---|---|---|
| Mipa-P$^{S221Q}$ | R13 component | F | 5’-TTGCCCCTGTCTCTCAGCATGCAATTCCAATG-3’ |
| | | R | 5’-CATTGGAATTGCATGCTGAGAGACAGGGGCAA-3’ |
| Mipa-P$^{S198C}$ | R14 component | F | 5’-CTTGGCATACGACTACTTGCTGGCCCCGCGTC-3’ |
| | | R | 5’-GACGCGGGGCCAGCAAGTAGTCGTATGCCAAG-3’ |
| Mipa-P$^{G229C}$ | R14 component | F | 5’-CAATTCCAATGTATACGTGTGTAGCGAGTGGG-3’ |
| | | R | 5’-CCCCACTCGCTACACACGTATACATTGGAATTG-3’ |
| Mipa-P$^{S61C}_{+A67Q}$ | R7+R9 | F | 5’-GTTATGGTGTAGTAAGTCAAACTATTACGGGTG-3’ |
| | | R | 5’-CACCCGTAATAGTTTGACTTACTACACCATAAC-3’ |
| Mipa-P$^{E46Q}_{+L49C}$ | R7+R8 | F | 5’-CCAGCAAGTGCAACGCAACGTGGCTGTGCACC-3’ |
| | | R | 5’-GGTGCACAGCCACGTTGCGTTGCACTTGCTGG-3’ |
| Mipa-P$^{T197K}_{+S198C+R201C+T203C}$ | R1+R2+R11+R14 | F | 5’-GGCATACGACTAAATGCTGGCCCTGCGTCTGC-3’ |
| | | R | 5’-GCAGACGCAGGGCCAGCATTTAGTCGTATGCC-3’ |
| Mipa-P$^{T228Q}_{+G229C+A231C}$ | R1+R4+R14 | F | 5’-ATTCCAATGTATCAGTGTGTATGCAGTGGGGAG-3’ |
| | | R | 5’-CTCCCCACTGCATACACACTGATACATTGGAAT-3’ |
| Kubu-P$^{A236C}$ | R1 component | F | 5’-CGTACCTTGAGTTATGCGGTGCGGATCATTTC-3’ |
| | | R | 5’-GAAATGATCCGCACCGCATAACTCAAGGTACG-3’ |
| Kubu-P$^{S281C}$ | R1 component | F | 5’-CGGGTGCAGCAGTATGCGCGTTTCGCTCGACC-3’ |
| | | R | 5’-GGTCGAGCGAAACGCGCATACTGCTGCACCCG-3’ |
| Kubu-P$^{A173C}$ | R2 component | F | 5’-CTGGAAGCATTATGCAAAGATACAACGGGTACCG-3’ |
| | | R | 5’-CGGTACCCGTTGTATCTTTGCATAATGCTTCCAG-3’ |
| Kubu-P$^{K197C}$ | R2 component | F | 5’-GGTCAAGATTTTAGTTGCGTGACGAAACCTG-3’ |
| | | R | 5’-CAGGTTTCGTCACGCAACTAAAATCTTGACC -3’ |
| Kubu-P$^{T95R}$ | R3 component | F | 5’-GGGTTTTGTAAGCCGTTGGTCACAAATTAGTTG-3’ |
| | | R | 5’-CAACTAATTTGTGACCAACGGCTTACAAAACCC-3’ |
| Kubu-P$^{D239S}$ | R4 component | F | 5’-TTAGCTGGTGCGAGCCATTTCTTTCCCACGACTG-3’ |
| | | R | 5’-CAGTCGTGGGAAAGAAATGGCTCGCACCAGCTAA -3’ |
| Kubu-P$^{E131Q}$ | R5 component | F | 5’-GCCAACGTGCGGATCAACTGCTGGCCGCCCTG-3’ |
| | | R | 5’-CAGGGCGGCCAGCAGTTGATCCGCACGTTGGC-3’ |
| Kubu-P$^{Q127S}$ | R6 component | F | 5’-CCCCCAGCTCACGTGCGGATGAACTGCTGGCCG-3’ |
| | | R | 5’-CGGCCAGCAGTTCATCCGCACGTGAGCTGGGGG-3’ |
| Kubu-P$^{A279S}$ | R8 component | F | 5’-CTTTGCGGGTGCAAGTGTATCCGCGTTTCGCTC-3’ |
| | | R | 5’- GAGCGAAACGCGGATACACTTGCACCCGCAAAG -3’ |

### Tables S4. The primers used in this study (4/4).

| Enz. | Category | Dir. | Sequence |
|---|---|---|---|
| Kubu-P$^{T119N}$ | R9 component | F | 5'-GTTGGTGCAGATACGAACTCTG GTTTTGATTCCC-3' |
| | | R | 5'-GGGAATCAAAACCAGAGTTCGTATCTGCACCAAC-3' |
| Kubu-P$^{V184I}$ | R10 | F | 5'-GTACCGTTAAAGCGGGAATACCGCTCGCACCGTG-3' |
| | | R | 5'-CACGGTGCGAGCGGTATTCCCGCTTTAACGGTAC-3' |
| Kubu-P$^{Q127S}_{+E131Q}$ | R5+R6 | F | 5'-CCCCCAGCTCACGTGCGGATCAACTGCTGGCCG-3' |
| | | R | 5'-CGGCCAGCAGTTGATCCGCACGTGAGCTGGGGG-3' |
| Kubu-P$^{A279S}_{+S281C}$ | R1+R8 | F | 5'-GGCTTTGCGGGTGCAAGTGTATGCGCGTTTCGC-3' |
| | | R | 5'-GCGAAACGCGCATACACTTGCACCCGCAAAGCC-3' |
| Kubu-P$^{V184I}_{+P185V+D190H}$ | Structure-determination | F | 5'-GTTAAAGCGGGAATAGTGCTCGCACCGTGGCAT-3' |
| | | R | 5'-ATGCCACGGTGCGAGCACTATTCCCGCTTTAAC-3' |
| Kubu-P$^{P185V}$ | Structure-determination | F | 5'-CCGTTAAAGCGGGAGTAGTGCTCGCACCGTGG-3' |
| | | R | 5'-CCACGGTGCGAGCACTACTCCCGCTTTAACGG-3' |
| I17-H64 | Primary Contest | F | 5'-TATATCATATGTCCAACCCCTACGAGCGTGG-3' |
| | | R | 5'-TATATCTCGAGGGTGGTGTGGGGGCAGGAG-3' |
| I8-H804 | Primary Contest | F | 5'-TATATCATATGCAGCAGACCGGCCCGAATCCC-3' |
| | | R | 5'-TATATCTCGAGGTAAGGGCAGTTCTGGCTGTAAC-3' |
| I123-H1110 | Primary Contest | F | 5'-TATATCATATGGCGGCCGACAACCCCTACCAAC-3' |
| | | R | 5'-TATATCTCGAGGAACGGGCAGGTGTTCATCGAC-3' |

### Tables S5. The codon-optimized sequences of Mipa-P, Kubu-P, LCC-ICCG and Hot-PETase.

| | Amino acid sequence | Nucleotide sequence |
|---|---|---|
| Mipa-P | MAPPASATERGLAPTAANITG<br>DGSYGVVSATITGASGFGGGV<br>VYYPNATERFPVVAISPGYTE<br>RWSSFAWLGRRLASWGFVVV<br>GIETNSLFDQPNSRGTQLLRAL<br>DWASSSAPAAVRDRVDATRQ<br>GVSGHSMGGGGTLSAMDQRP<br>SVRAGVPLAPWHTTTSWPRV<br>TNPVMILGGQNDGIAPVSSHA<br>IPMYTGVASGEKAYVELAGA<br>GHNFPNSANPIVSRAAVSWFK<br>RFLDDDTRFAPFACDFGGASIS<br>QFRSTCPVLEHHHHHH | CATATGGCACCCCCAGCAAGTGCAACGGAACGTGGCTTAGCACCAACGGCAGCAAAT<br>ATTACGGGTGATGGTTCTTATGGTGTAGTAAGTGCAACTATTACGGGTGCGTCGGGCT<br>TTGGCGGTGGGGTAGTTTATTATCCCAATGCTACGGAGCGTTTTCCTGTGGTTGCAATT<br>AGCCCGGGTTATACGGAGCGTTGGAGCAGTTTTGCATGGCTGGGTCGTCGCCTCGCA<br>AGCTGGGGCTTTGTTGTGGTGGGGATTGAAACCAATTCACTGTTTGATCAACCAAATT<br>CGCGCGGAACTCAATTACTCCGTGCTCTCGATTGGGCGTCATCAAGCGCGCCTGCGGC<br>AGTTCGCGATCGTGTGGATGCAACACGCCAAGGCGTAAGTGGCCATAGCATGGGTGG<br>CGGTGGCACCCTTAGCGCGATGGATCAACGTCCGAGCGTACGTGCTGGGGTCCCATT<br>AGCACCTTGGCATACGACTACTAGCTGGCCCCGCGTCACCAATCCAGTAATGATTTTG<br>GGTGGTCAAAATGATGGTATTGCCCCTGTCTCTAGCCATGCAATTCCAATGTATACGG<br>GTGTAGCGAGTGGGGAGAAAGCCTATGTGGAACTCGCAGGTGCGGGCCATAATTTTC<br>CGAATTCGGCGAATCCCATTGTATCACGTGCTGCAGTCTCCTGGTTTAAACGCTTTCTG<br>GATGATGATACGCGCTTTGCGCCATTTGCGTGTGATTTTGGTGGTGCGAGCATCAGTC<br>AATTTCGTAGTACGTGCCCCGTACTCGAGCACCACCACCACCACCACTGA |
| Kubu-P | MADQVGQAPTAANITGDGSF<br>ATASAPITNQTGFGGGTVYYP<br>TAAGTYPVVAVVPGFVSTWS<br>QISWLGPRVASWGFVVVGAD<br>TTSGFDSPSQRADELLAALNW<br>AVNSAPAAVRGKVDGTRRGV<br>AGWSMGGGGTLEALAKDTTG<br>TVKAGVPLAPWDIGQDFSKVT<br>KPVFIVGAQNDTIAPPAQHAV<br>PFYNAAAGPKSYLELAGADH<br>FFPTTANPTVSRAMVSWLKRF<br>VSSDDRFTPFTCGFAGAAVSA<br>FRSTACLEHHHHHH | CATATGGCTGACCAAGTGGGACAAGCACCGACAGCAGCCAATATTACAGGAGATGG<br>GTCGTTTGCGACCGCAAGCGCTCCTATTACGAATCAAACTGGTTTTGGTGGTGGTACT<br>GTTTATTATCCTACCGCCGCGGGAACCTACCCTGTCGTAGCGGTTGTGCCGGGTTTTGT<br>AAGCACATGGTCACAAATTAGTTGGCTTGGACCCCGCGTAGCCTCTTGGGGTTTTGTG<br>GTGGTTGGTGCAGATACGACATCTGGTTTTGATTCCCCCAGCCAACGTGCGGATGAAC<br>TGCTGGCCGCCCTGAATTGGGCCGTTAATTCAGCCCCCGCTGCGGTTCGTGGTAAAGT<br>AGATGGTACCCGTCGCGGTGTAGCTGGTTGGAGCATGGGTGGTGGTGGTACACTGGA<br>AGCATTAGCGAAAGATACAACGGGTACCGTTAAAGCGGGAGTACCGCTCGCACCGTG<br>GGATATTGGTCAAGATTTTAGTAAAGTGACGAAACCTGTATTTATTGTTGGTGCACAA<br>AATGATACGATTGCCCCACCGGCTCAACATGCGGTTCCGTTTTATAATGCAGCAGCGG<br>GTCCTAAATCGTACCTTGAGTTAGCTGGTGCGGATCATTTCTTTCCCACGACTGCTAAT<br>CCGACCGTTTCGCGTGCTATGGTTAGCTGGCTCAAACGTTTTGTTTCTTCAGATGATCG<br>TTTTACACCTTTTACTTGTGGCTTTGCGGGTGCAGCAGTATCCGCGTTTCGCTCGACCG<br>CGTGTCTCGAGCACCACCACCACCACCACTGA |
| LCC-ICCG | MQSNPYQRGPNPTRSALTAD<br>GPFSVATYTVSRLSVSGFGGG<br>VIYYPTGTSLTFGGIAMSPGYT<br>ADASSLAWLGRRLASHGFVV<br>LVINTNSRFDGPDSRASQLSA<br>ALNYLRTSSPSAVRARLDANR<br>LAVAGHSMGGGGTLRIAEQN<br>PSLKAAVPLTPWHTDKTFNTS<br>VPVLIVGAEADTVAPVSQHAI<br>PFYQNLPSTTPKVYVELCNAS<br>HIAPNSNNAAISVYTISWMKL<br>WVDNDTRYRQFLCNVNDPAL<br>CDFRTNNRHCQLEHHHHHH | CATATGCAATCCAACCCGTACCAGCGCGGTCCCAACCCCACGCGGAGCGCGCTCACG<br>GCCGACGGGCCGTTCTCGGTGGCAACCTACACCGTTTCGCGGCTCTCGGTGAGCGGCT<br>TCGGGGGCGGGGTGATCTACTACCCCACAGGCACCTCGCTGACCTTCGGCGGGATCG<br>CCATGTCGCCGGGGTACACGGCCGACGCCAGTTCGCTGGCGTGGCTCGGGCGGCGGT<br>TGGCATCGCACGGCTTCGTGGTGCTCGTCATCAACACGAACTCGCGATTCGACGGCCC<br>TGACTCCCGCGCAAGCCAGCTATCGGCGGCGCTGAACTACCTGCGGACGAGCAGCCC<br>CTCAGCCGTACGCGCCCGGCTCGATGCGAACCGCCTGGCCGTTGCGGGGCATTCGAT<br>GGGCGGCGGCGGGACCCTCCGCATCGCAGAGCAGAACCCGTCGCTGAAAGCGGCCG<br>TACCGCTCACGCCGTGGCACACCGACAAGACGTTCAACACGTCGGTACCGGTGCTCA<br>TCGTGGGAGCGGAAGCGGACACGGTCGCGCCCGTGAGCCAGCACGCCATTCCGTTCT<br>ACCAGAACTTGCCCTCGACCACGCCGAAGGTGTACGTGGAGCTCTGCAATGCGTCGC<br>ACATCGCGCCCAACAGCAACAACGCGGCGATCTCCGTGTACACCATCTCGTGGATGA<br>AGCTGTGGGTGGATAACGACACCCGCTACCGGCAGTTCCTCTGCAACGTGAACGATC<br>CGGCGCTGTGCGACTTCCGGACGAACAACCGCCACTGCCAGCTCGAGCACCACCACC<br>ACCACCACTGA |
| Hot-PETase | MQTNPYARGPNPTAASLEASA<br>GPFTVRSFTVARPVGYGAGTV<br>YYPTNAGGTVGAIAIVPGYTA<br>TQSSINWWGPRLASHGFVVITI<br>DTNSTLDKPESRSSQQMAALR<br>QVASLNGTSSSPIYGKVDTAR<br>GGVMGWSMGGGGSLISAAN<br>NPSLKAAAVMAPWHSSTNFS<br>SVTVPTLIFACENDRIAPVKEY<br>ALPIYDSMSLNAKQFLEICGGS<br>HSCACSGNSNQALIGMKGVA<br>WMKRFMDNDTRYSQFACENP<br>NSTAVCDFRTANCSLEHHHH<br>HH | CATATGCAGACTAACCCCTATGCTCGCGGGCCGAATCCTACAGCGGCCTCGTTGGAA<br>GCCAGTGCAGGTCCCTTCACCGTACGTTCGTTTACTGTTGCGCGTCCAGTGGGATATG<br>GGGCTGGCACCGTCTATTATCCGACTAACGCCGGTGGTACTGTGGGCGCAATCGCCAT<br>TGTCCCCGGCTACACTGCAACTCAATCCTCGATTAATTGGTGGGGACCACGCTTGGCT<br>AGCCACGGGTTTGTTGTGATCACCATCGATACTAACAGTACGTTGGACAAGCCAGAG<br>AGTCGCAGCTCTCAGCAGATGGCGGCATTACGCCAGGTGGCGAGCTTAAATGGGACG<br>AGTTCAAGTCCAATTTATGGCAAGGTCGATACCGCCCGTGGTGGAGTAATGGGCTGG<br>AGTATGGGTGGCGGCGGATCTCTTATCTCGGCAGCGAATAATCCAAGCCTGAAGGCA<br>GCAGCCGTTATGGCACCCTGGCATAGTAGTACGAACTTTTCGTCTGTTACGGTTCCTA<br>CGCTTATCTTTGCCTGTGAGAATGATAGGATCGCACCAGTGAAGGAGTACGCCTTGCC<br>AATTTACGACTCGATGTCGCTCAACGCTAAACAGTTTCTTGAGATTTGTGGGGGCTCT<br>CACTCTTGTGCCTGTAGTGGGAATTCCAATCAAGCCCTTATCGGAATGAAGGGCGTTG<br>CTTGGATGAAGCGCTTCATGGACAATGACACTCGTTATTCACAGTTCGCTTGCGAAAA<br>CCCAAACTCAACCGCCGTATGTGATTTTCGCACTGCTAACTGCAGCCTCGAGCACCAC<br>CACCACCACCACTGA |
